# Effective connectivity and spatial selectivity-dependent fMRI changes elicited by microstimulation of pulvinar and LIP

**DOI:** 10.1101/2020.09.16.298539

**Authors:** Igor Kagan, Lydia Gibson, Elena Spanou, Melanie Wilke

## Abstract

The thalamic pulvinar and the lateral intraparietal area (LIP) share reciprocal anatomical connections and are part of an extensive cortical and subcortical network involved in spatial attention and oculomotor processing. The goal of this study was to compare the effective connectivity of dorsal pulvinar (dPul) and LIP and to probe the dependency of microstimulation effects on task demands and spatial tuning properties of a given brain region. To this end, we applied unilateral electrical microstimulation in the dPul (mainly medial pulvinar) and LIP in combination with event-related BOLD fMRI in monkeys performing fixation and memory-guided saccade tasks. Microstimulation in both dPul and LIP enhanced task-related activity in monosynaptically-connected fronto-parietal cortex and along the superior temporal sulcus (STS) including putative face patch locations, as well as in extrastriate cortex. LIP microstimulation elicited strong activity in the opposite homotopic LIP while no homotopic activation was found with dPul stimulation. Both dPul and LIP stimulation also elicited activity in several heterotopic cortical areas in the opposite hemisphere, implying polysynaptic propagation of excitation. Despite extensive activation along the intraparietal sulcus evoked by LIP stimulation, there was a difference in frontal and occipital connectivity elicited by posterior and anterior LIP stimulation sites. Comparison of dPul stimulation with the adjacent but functionally dissimilar ventral pulvinar also showed distinct connectivity. On the level of single trial timecourses within each region of interest (ROI), most ROIs did not show task-dependence of stimulation-elicited response modulation. Across ROIs, however, there was an interaction between task and stimulation, and task-specific correlations between the initial spatial selectivity and the magnitude of stimulation effect were observed. Consequently, stimulation-elicited modulation of task-related activity was best fitted by an additive model scaled down by the initial response amplitude. In summary, we identified overlapping and distinct patterns of thalamocortical and corticocortical connectivity of pulvinar and LIP, highlighting the dorsal bank and fundus of STS as a prominent node of shared circuitry. Spatial task-specific and partly polysynaptic modulations of cue and saccade planning delay period activity in both hemispheres exerted by unilateral pulvinar and parietal stimulation provide insight into the distributed interhemispheric processing underlying spatial behavior.

**Highlights:**

- Electrical stimulation of pulvinar and LIP was used to study fMRI effective connectivity
- Both regions activated prefrontal cortex and the dorsal bank of superior temporal sulcus
- Activations within and across hemispheres suggest polysynaptic propagation
- Stimulation effects show interactions between task- and spatial selectivity
- Stimulation effects are best fitted by an additive model scaled by the initial response

## 1 Introduction

Spatial exploration and selection of visual stimuli for further analysis by goal-directed saccadic eye movements is an important feature of primate behavior. This behavior relies on a widely distributed subcortical-cortical brain circuitry including fronto-parietal cortices, superior colliculus (SC), basal ganglia and thalamus (Andersen and Cui, 2009; Glimcher, 2003; Gold and Shadlen, 2007; Wurtz et al., 2011; Wurtz and Hikosaka, 1986). While until recently most concepts of flexible visually-guided behavior emphasized interactions within cortical networks, direct connections between cortical regions are paralleled by indirect routes through higher-order thalamic nuclei such as pulvinar (dPul), raising the question what kind of information is contributed by the additional thalamic routes and how the processing compares between the functionally pertinent cortical and thalamic sites (Bridge et al., 2015; Grieve et al., 2000; Halassa and Kastner, 2017; Sherman and Guillery, 2011; Shipp, 2003). Two major nodes of the network that link vision with saccadic eye movements and where lesions lead to spatial neglect/extinction symptoms in monkeys and humans are the lateral intraparietal region and the dorsal pulvinar (Karnath et al., 2002; Mesulam, 1999; Storm et al., 2017; Wardak et al., 2002; M. Wilke et al., 2010). The causal influence of these two regions on interconnected brain circuitry is the focus of the current study.

In monkeys, the lateral intraparietal area (LIP) is located in the intraparietal sulcus (IPS) of the posterior parietal cortex (PPC), and its neurons encode the position and behavioral relevance of visual stimuli along with the direction of an upcoming saccade (Andersen et al., 1990; Barash et al., 1991; Bisley and Goldberg, 2010). Response fields in LIP are predominantly in the contralateral hemifield (Ben Hamed et al., 2001; Blatt et al., 1990). Furthermore, LIP neurons exhibit delay period activity in memory-guided saccade tasks that reflects saccade preparation and the spatial allocation of attention (Andersen and Cui, 2009; Bisley and Goldberg, 2010; Colby and Goldberg, 1999). Area LIP is reciprocally interconnected with the pulvinar (Asanuma et al., 1985; Blatt et al., 1990; Hardy and Lynch, 1992; Romanski et al., 1997), which is a heterogeneous structure that has been subdivided in various manners based on histochemical, cytoarchitectonic or anatomical connectivity tracer techniques (Baldwin et al., 2017; Bridge et al., 2015; Gutierrez et al., 2000; Jones, 2007). Although not reflecting all detailed division schemes, several anatomical and functional studies have motivated a functional distinction between ventral and dorsal pulvinar portions (Arcaro et al., 2015; Kaas and Lyon, 2007; Komura et al., 2013; Stepniewska et al., 1994). The ventral pulvinar (vPul) encompasses the inferior pulvinar (PI) and the ventral part of the lateral pulvinar (PLvl), has strong reciprocal connections to visual cortices and contains several retinotopic maps with mainly contralateral visual receptive fields (Arcaro et al., 2015; Arcaro and Livingstone, 2017; Bender, 1981; Petersen et al., 1985). Ventral pulvinar neurons are also modulated by visual attention (Bender and Youakim, 2001; Saalmann et al., 2012; Zhou et al., 2016) and show saccade-related activity that is mostly peri- or post-saccadic (Berman and Wurtz, 2011; Petersen et al., 1985; Robinson et al., 1986). The dorsal pulvinar (dPul) occupies the region above the level of the brachium of the superior colliculus and encompasses the medial pulvinar (PM) and the dorsal portion of the lateral pulvinar (PLdm) (Gutierrez et al., 2000; Kaas and Lyon, 2007). Similar to LIP and vPul, dPul neurons show enhancement for visual stimuli that are attended due to their behavioral relevance and/or indicate an upcoming saccade target (Bender and Youakim, 2001; Fiebelkorn et al., 2019) and discharge in cue and saccade execution phases with an overall contralateral preference (Benevento and Port, 1995; Dominguez-Vargas et al., 2017; Robinson et al., 1986; Schneider et al., 2020). But unlike vPul and to a certain extent LIP (Patel et al., 2010), dPul does not follow a clear retinotopic organization (Benevento and Miller, 1981; Benevento and Port, 1995; Petersen et al., 1987).

Providing causal evidence for involvement in spatial visuomotor behavior, reversible pharmacological inactivation in LIP or dPul leads to spatial neglect/extinction characterized by impaired responses to and exploration of the contralateral space (Christopoulos et al., 2018; Li et al., 1999; Liu et al., 2010; Petersen et al., 1987; Wardak et al., 2002, 2004; M. Wilke et al., 2010; Wilke et al., 2013). Particularly, employing visually-guided or memory-guided saccade tasks in monkeys, several studies have shown that local inactivation of LIP or the dorsal pulvinar leads to biased saccade choices towards the ipsilesional side while keeping saccade parameters largely intact, apart from a moderate increase of contralesional saccade latencies (Chen et al., 2016; Katz et al., 2016; Kubanek et al., 2015; Wardak et al., 2002; M. Wilke et al., 2010; Wilke et al., 2012, 2013). Similarly, electrical microstimulation in LIP or dorsal pulvinar induces a bias of perceptual and saccade choices towards contraversive (contralateral to the side of stimulation) spatial positions, and affects saccade latencies (Dai et al., 2014; Dominguez-Vargas et al., 2017; Hanks et al., 2006). Neglect-like deficits following stroke-induced parietal and pulvinar lesions in humans are also consistent with the critical contribution of both regions to visually-guided spatial behavior (Arend et al., 2008a, 2008b; Karnath, 2015; Karnath et al., 2002; Rafal et al., 2004; Van der Stigchel et al., 2010).

Area LIP and the dorsal pulvinar share a large number of afferent and efferent subcortical and cortical connections (Blatt et al., 1990; Lewis and Van Essen, 2000). Anatomical tracer studies have shown that both regions are reciprocally connected to other areas in the posterior parietal cortex (e.g. MIP, AIP, area 7) (Asanuma et al., 1985; Cappe et al., 2009, 2007; Grieve et al., 2000; Hardy and Lynch, 1992; Yeterian and Pandya, 1985), the lower and upper bank of the superior temporal sulcus (STS), eye movement-related areas in prefrontal cortex such as dorsolateral prefrontal cortex (dlPFC) and frontal eye fields (FEF, areas 8 and 45) (Homman-Ludiye et al., 2020; Lewis and Van Essen, 2000; Romanski et al., 1997) and extrastriate visual cortex (Blatt et al., 1990). Similar functional connectivity patterns have been reported in human fMRI studies (Arcaro et al., 2018, 2015). While LIP projects to the intermediate layers of the superior colliculus (Andersen et al., 1990; Pare and Wurtz, 1997), dPul only receives afferent projections from the intermediate and deep layers of the superior colliculus without sending a projection back (Baldwin and Bourne, 2017; Bender and Butter, 1987; Benevento and Standage, 1983; Harting et al., 1980).

The combination of electrical microstimulation and fMRI (es-fMRI) has emerged as a powerful technique to probe the effective connectivity of a local cortical or subcortical stimulation site to distal brain regions *in vivo* (Field et al., 2008; Logothetis et al., 2010; Matsui et al., 2011; Moeller et al., 2008; Murayama et al., 2011; Petkov et al., 2015; Sultan et al., 2012), as recently summarized in an extensive review (Klink et al., 2021). Es-fMRI allows the mapping of anatomical connections in a living animal with a precision that cannot be achieved with other in-vivo methods such as diffusion tensor imaging or resting state connectivity (Honey et al., 2009; Schilling et al., 2019). Furthermore, previous studies showed that es-fMRI provides functional information that goes beyond purely anatomical connections. Particularly, the strength of stimulation effects is modulated by the brain state (e.g. anaesthetized vs. awake) (Murris et al., 2020), the properties of the employed visual stimuli such as contrast or the visual responsiveness of activated voxels (Ekstrom et al., 2009, 2008) and task demands (Premereur et al., 2013). The first *goal* of this study was to identify the shared functional circuitry of dPul and LIP during oculomotor tasks. While es-fMRI effective connectivity of the pulvinar has not been investigated previously, es-fMRI effects of LIP have been studied as a control experiment in a work on grasp-related anterior intraparietal area AIP, but were not the main focus of this paper (Premereur et al., 2015). Second, we asked how the magnitude of stimulation effects of dPul and LIP on BOLD responses depends on task demands and/or spatial contingencies. A similar question has been addressed in a es-fMRI study on area FEF (Premereur et al., 2013), but not for parietal or thalamic regions. Based on our previous behavioral context-contingent effects of pulvinar microstimulation (Dominguez-Vargas et al., 2017) and es-fMRI studies in FEF showing visual stimulus-gated stimulation effects in early visual areas, and stronger effects in a saccade task compared to passive fixation (Ekstrom et al., 2009; Premereur et al., 2013), we hypothesized that microstimulation-elicited fMRI effects might be more pronounced in the saccade task compared to pure visual fixation. Third, we asked whether microstimulation in LIP and/or dPul interacts with the spatial tuning of a given brain region. For instance, we expected the largest microstimulation effect in the regions less optimally driven by the task (Ekstrom et al., 2008; Premereur et al., 2013), e.g. for contraversively tuned voxels in a ipsiversive saccade condition. To address these questions, we measured brain-wide fMRI BOLD activity while monkeys performed a sustained fixation or a memory saccade task towards ipsi- and contraversive spatial positions. This task, similarly to covert spatial attention tasks, elicits strong and often contralaterally-tuned activation in the fronto-parietal and parieto-temporal network (Bogadhi et al., 2018; Caspari et al., 2015; Kagan et al., 2010; Wilke et al., 2012). We employed a slow time-resolved event-related design to isolate BOLD responses from different task epochs (Kagan et al., 2010), and compared fixation, visual cue and delay period activity in interleaved trials with and without microstimulation, in an extensive dataset comprising microstimulation in several pulvinar and LIP locations.

## 2 Methods

### Data and code availability statement

The data used in this study will be made available to the community via **PRIMatE Data Exchange (PRIME-DE)** repository (http://fcon_1000.projects.nitrc.org/indi/indiPRIME.html), and all custom source code will be uploaded to a publically accessible GitHub repository (https://github.com/igorkagan/Kagan_et_al_2021).

### 2.1 Procedures

All experimental procedures complied with the ARRIVE guidelines (https://arriveguidelines.org) and were conducted in accordance with the European Directive 2010/63/EU, the corresponding German law governing animal welfare, and German Primate Center institutional guidelines. The procedures were approved by the responsible government agency LAVES (Niedersaechsisches Landesamt fuer Verbraucherschutz und Lebensmittelsicherheit, Oldenburg, Germany).

### 2.2 Animal preparation

Two adult male rhesus monkeys (*Macaca mulatta*), weighing approximately 9 kg (monkey C) and 10.5 kg (monkey B), served as subjects. Surgical procedures were similar for both animals. In an initial surgery, each animal was implanted with a MRI-compatible polyetheretherketone (PEEK) head post embedded in a bone cement head cap (Palacos with gentamicin; BioMet) anchored by ceramic screws (Rogue Research) under general anesthesia and aseptic conditions. A separate surgery was performed in each animal to implant a PEEK MRI-compatible chamber (inside diameter 22 mm) allowing access to the right pulvinar and LIP. To aid the MR-guided pre-surgical planning of the chamber’s location and angle in the stereotaxic space, performed in the Planner software package (https://github.com/shayo/Planner) (Ohayon and Tsao, 2012), several 1-2 mm indents that were filled with a MR-visible markers during scanning were embedded in the head cap. After confirming chamber positioning with post-surgical MRI scans, a partial craniotomy was made inside the chamber (chamber positioning monkey C, right hemisphere: center at 0.5A, 14.5R mm, tilted - 11P, 27R degrees; monkey B, right hemisphere: center at 3.5P, 17R mm, tilted 15P, 30R degrees; the coordinates are relative to stereotaxic zero, A – anterior, P – posterior, R – right; the tilts are relative to the vertical axis normal to stereotaxic plane: P – posterior, top of the chamber tilted towards the back of the head, R – right, top of the chamber tilted towards the right ear).

### 2.3 Electrical microstimulation

An S88X dual output square pulse stimulator (Grass Products, Natus Neurology, USA) triggered by a custom MATLAB-based task controller software (https://github.com/dagdpz/monkeypsych) generated 200 ms trains of twin pulses at 300 Hz, which in turn triggered a battery-powered constant current stimulus isolator A365 (World Precision Instruments, USA) to produce 60 biphasic pulses. These devices were placed in the control room outside of the scanner Faraday cage. The current (100-250 µA) was delivered to the target structure using single monopolar electrodes (platinum-iridium, 100 mm length, 125 *µ*m thick core, initial 2 cm glass coating with an exposed tip of 40 *µ*m, total thickness of 230 *µ*m including polyamide tubing coating, customer part ID: UEIK1, FHC Inc., USA). The electrode was connected to the lead wire using a small detachable stainless steel tubing connector made from 27G Spinocan Spinal needle (B. Braun). A return (reference) tungsten rod was placed in the recording chamber filled with saline. The two wires leading to the electrode and the reference were twisted together and fed into a shielded BNC cable that was connected, via a low pass filter (50 MHz cutoff) and the patch panel BNC or a waveguide of the scanner Faraday cage, to the output of the stimulus isolator. Voltage drop was monitored as the difference between voltage measured before and after a 10 kO resistor in series with the electrode using a 2 channel 2GS/s 200 MHz Tektronix TDS2022C oscilloscope. The manufacturer-specified impedance of the electrodes was 300-500 kO. The initial impedance measured in the lab at 1000 Hz before the experiment was 110-1400 kO. Since the impedance dropped dramatically after a few stimulation trains were applied, before each session 20-30 pulse trains were delivered to the electrode immersed in saline using 250 *µ*A current, in order to bring the electrode impedance to a more stable regime. Following this procedure, the impedance ranged from 10 kO to 70 kO in monkey C and from 10 kO to 120 kO in monkey B. In each stimulation trial, 10 consecutive stimulation trains were applied at a frequency of 1 Hz (see section 2.6 for behavioral task).

### 2.4 MR imaging

Both animals were scanned in a 3 T MRI scanner (monkey C: Magnetom Tim Trio; Siemens; monkey B: Magnetom Prisma; Siemens). For planning of chamber and electrode positioning for each animal, high-resolution full-head T1-weighted (3D magnetization-prepared rapid gradient-echo, MPRAGE, 0.5 mm isometric) and additional T2-weighted (rapid acquisition with relaxation enhancement, RARE, 0.25 mm in plane, 1 mm slice thickness) images with the slice package aligned to the chamber vertical axis were acquired before and after chamber implantation in an awake state or under anesthesia using the built-in gradient body transmit coil and a custom single-loop receive coil (Windmiller Kolster Scientific). T1- and T2-weighted scans were coregistered and transformed into “chamber normal” orientation (aligned to the chamber vertical axis) and into anterior commissure-posterior commissure (AC–PC) space for electrode targeting and visualization. These images were acquired with the chamber and the grid filled with gadolinium (Magnevist; Bayer)/saline solution (proportion 1:200) with tungsten rods inserted in predefined grid locations for alignment purposes.

Functional BOLD data were acquired using a custom-made 4-channel receive coil and 1-channel transmit coil (Windmiller Kolster Scientific). In each experimental session, the transmit coil and each individual channel of the receive coil were tuned and matched to the resonance frequency of the scanner (∼123.3 MHz) with manual knob adjustments, after being fixed in the final position around the monkey head, using portable probe tuning device (Morris Instruments Inc). In each session, advanced 3D shimming within a cuboid volume contained inside the brain followed by an interactive shimming was performed to maximize the T2* and minimize the spectral tuning curve full width at half maximum. To minimize motion-related distortions during shimming, monkeys performed a long fixation task and data was only acquired during stable fixation periods. After shimming and a localizer scan, T2-weighted images with the slice package aligned to the chamber vertical axis (32 slices) were acquired (**Figure 1A**). The first image was acquired with the electrode inserted into the brain but before lowering it to the targeted brain region. Based on that image, we measured the 2D-distance between the electrode tip and the center of the target structure and adjusted the electrode position accordingly. The second image was acquired after adjusting the electrode position to confirm proper placement within the target structure, allowing for the online control over electrode positioning. In addition, an in-plane (with the functional EPI slices) anatomical scan was acquired (T2-weighted, Turbo-Spin-Echo sequence, TR: 5 s, TE: 18 ms, flip angle: 180°, 0.75 mm x 0.75 mm in-plane resolution, slice thickness: 1.2 mm) for later coregistration with the functional images. Blood-oxygen-level dependent (BOLD) functional images were acquired with a T2*-weighted echo-planar imaging (EPI) single-shot sequence (TR: 2 s, TE: 27 ms, flip angle: 76°, bandwidth: 1302 Hz/pixel, 80×80 matrix, anterior-posterior phase-encoding direction, no acceleration in monkey C and image parallel acceleration factor (iPAT) 2 in monkey B, percent sampling 100% in monkey C and 80% in monkey B, field of view: 96 mm, 1.2 mm x 1.2 mm in-plane resolution, 30 axial slices in monkey C and 32 axial slices in monkey B, slice thickness: 1.2 mm, inter slice time: 66.7 ms for monkey C and 61.8 ms for monkey B). In each session, data were acquired in several separate, consecutive runs of 454 volumes (15 minutes) each. Before functional data acquisition, the received signal was checked for RF artifacts and the level of background noise using the same echo-planar imaging sequence with the transmitter RF pulse amplitude set to zero, without and with the microstimulation circuit connections and the running microstimulation.

**Figure 1.**
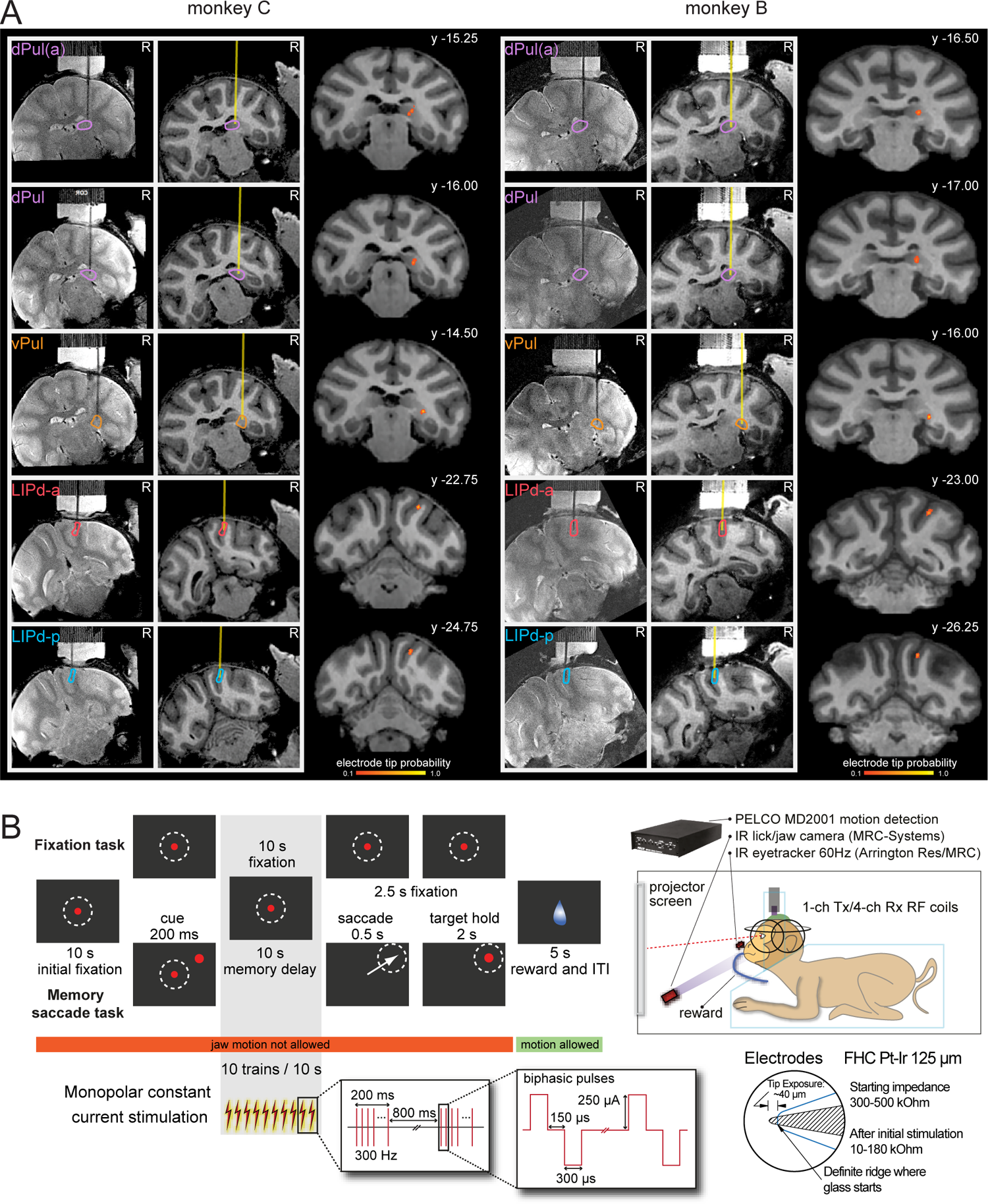
Microstimulation targeting, task and microstimulation timing. **(A)** Electrode localization in dPul, vPul and LIPd. Electrode positions localized in T2-weighted MR images (left column in each monkey) and reconstructed in T1-weighted images (yellow line, middle column in each monkey), both aligned to the chamber vertical axis, in example sessions. Color outlines mark the respective target region. The right column for each monkey shows an example coronal section through the center of a probability map reconstructing the electrode tip position across all sessions, displayed on a T1-weighted MR image aligned to AC-PC space. R - right, y - distance from AC-PC origin along the anterior/posterior plane in mm. **(B)** Timing of one successful trial of the fixation task and the memory saccade task. Dashed circles illustrate the eye position fixation window but were not displayed on the screen. Correct trials were rewarded with a fluid reward. Jaw motion was only allowed during reward delivery and the inter-trial interval (ITI). In half of the trials current stimulation was applied in the indicated time period. The timing and the properties of the microstimulation are depicted below the trial timing. On the right inset, the custom specifications used for the manufacturing the stimulating electrodes are depicted.

### 2.5 Pulvinar and LIP targeting

A custom-made MR-compatible polyetherimide (Ultem) grid (0.8 mm hole spacing, 0.43 mm hole diameter) and a custom-made plastic XYZ manipulator drive (design courtesy of Dr. Sebastian Moeller, (Moeller et al., 2008)) were used to position platinum-iridium electrodes (FHC, see section 2.3 for detailed specifications) in the corresponding grid hole. Grid hole planning and penetration trajectory estimation was based on anatomical MRI using Planner (Ohayon and Tsao, 2012) and BrainVoyager (Version 2.4.2, 64-bit; Brain Innovation). During penetration, the electrode was protected by a custom-made MR compatible guide tube (polyimide coated fused silica, 430 *µ*m outer diameter, 320 *µ*m inner diameter; Polymicro Technologies). An MR compatible stopper (polyimide coated fused silica, 700 *µ*m outer diameter, 530 *µ*m inner diameter; Polymicro Technologies) ensured that the guide tube only penetrated the dura and minimally the cortex below. Before penetration, the electrode tip was aligned to the guide tube tip and was held in place by a drop of melted petroleum jelly. For each experimental session, the final electrode location was determined based on the coronal T2-weighted scan (see section 2.4). In each animal, we stimulated several different sites, some located more anterior and the other located more posterior, both in the right dPul and the right dorsal LIP.

For targeting the pulvinar stimulation sites during the experiments we used online and downloadable NeuroMaps atlas (Rohlfing et al., 2012). The coronal template outlines were adapted from http://braininfo.rprc.washington.edu/PrimateBrainMaps/atlas/Mapindex.html, also available at https://scalablebrainatlas.incf.org/macaque/DB09 (Wu et al., 2000). For all analyses we utilized the recent SARM atlas in NMT v2 template space that uses anterior, medial, lateral and inferior delineation (APul, MPul, LPul, IPul) (Hartig et al., 2020; Jung et al., 2021). Since various atlases and research papers adopt different pulvinar nomenclature and abbreviations (e.g. PM or MPul for the medial pulvinar, and PLdm/PLvl or LPul for the lateral pulvinar), we summarize the nomenclature schematically in **Figure 2C**. The anterior pulvinar, the medial pulvinar and the dorsal part of the lateral pulvinar (often denoted PLdm, e.g. (Bridge et al., 2015; Kaas and Lyon, 2007)) constitute the dorsal pulvinar (dPul); the ventral part of the lateral pulvinar (often denoted PLvl, e.g. (Baldwin et al., 2017)) and the inferior pulvinar comprise the ventral pulvinar (vPul).

**Figure 2.**
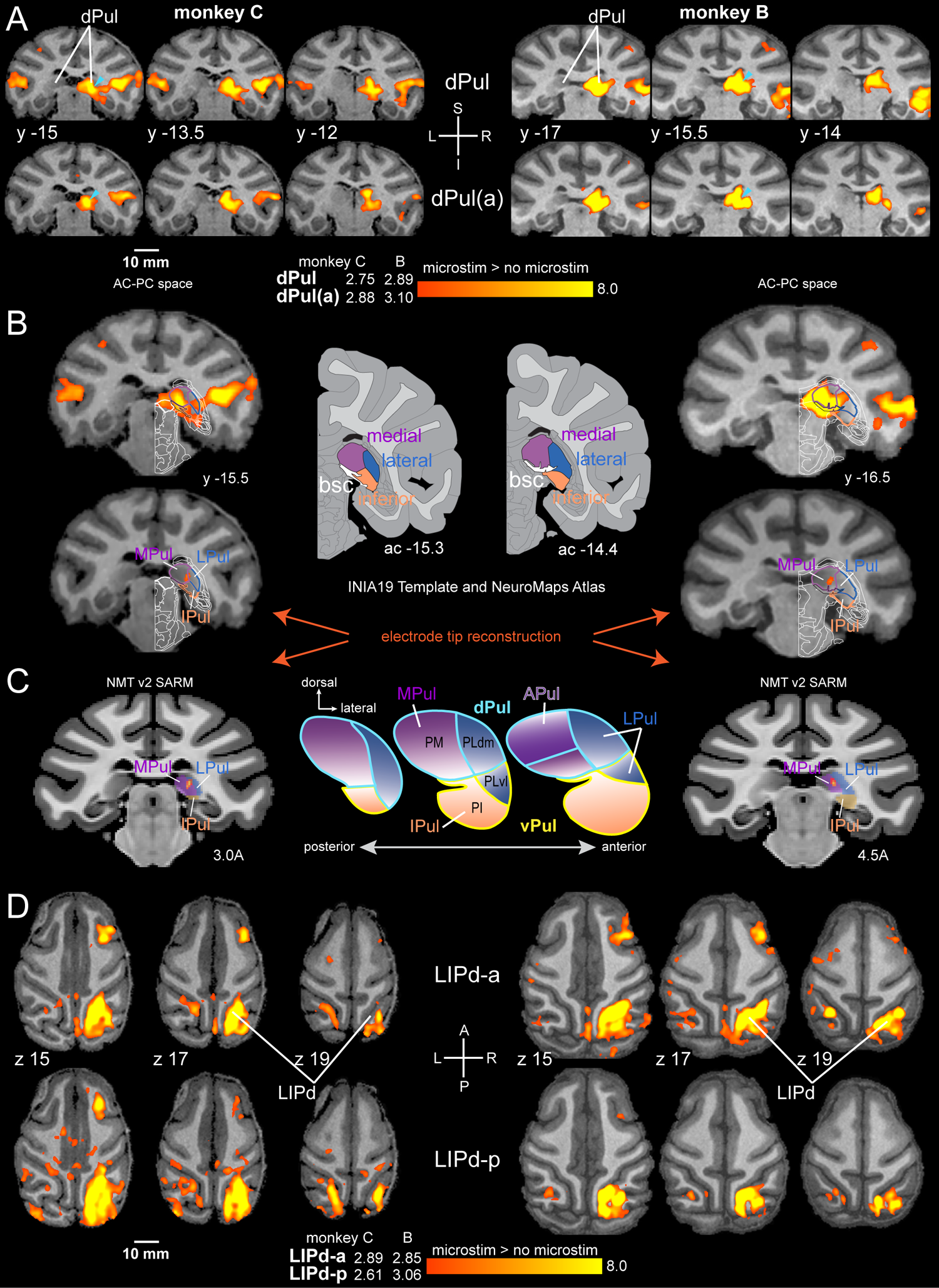
Local activation at the stimulation sites. **(A)** Coronal sections showing statistical t-maps of BOLD activation elicited by microstimulation in the two dPul sites (microstimulation > no microstimulation contrast, all tasks combined). In this and all other figures (unless stated otherwise), the activation maps were thresholded at q(FDR)<0.05 level and with a cluster threshold of 20 functional voxels, producing individual threshold t-value for each dataset and each monkey (see Methods), indicated next to the color bar: here and in other figures, left value is for monkey C, right value for monkey B. Cyan arrowheads indicate the location of the electrode tip, and an approximate angle. **(B)** Coronal sections showing activation (top image, dPul dataset) through the center of the electrode tip reconstruction map (bottom image). Schematic outlines of the pulvinar nuclei (medial, lateral, inferior) and neighboring structures were adapted from the NeuroMaps atlas (Rohlfing et al., 2012), shown in the middle. **(C)** The center of the electrode tip reconstruction map warped to NMT v2 template, with SARM pulvinar regions. The schematic in the middle shows pulvinar parcellation nomenclature. **(D)** Axial sections showing statistical t-maps of BOLD activation elicited by microstimulation in the two LIPd sites. L – left, R – right, S – superior, I – inferior, A – anterior, P – posterior, y - distance from AC-PC origin along the anterior/posterior plane, z - distance from AC-PC origin along the superior/inferior axis, in millimeters.

**Figure 1A** shows the electrode positions in example sessions of dPul stimulation measured in T2-weighted MR images with the slice package aligned to the chamber vertical axis and reconstructed in T1-weighted MR images for each animal, and one coronal section through the center of the electrode tip probability map in AC-PC plane (see section 2.7 for a description of the generation of the displayed probability maps). **Figure S1** (upper two rows in each monkey) shows the full extent of probability maps of the estimated electrode tip positions across dPul stimulation sessions. As can be seen, the dPul stimulation sites corresponded to MPul and the dorsal part of LPul (PLdm). The brachium of the SC (bsc) and other neighboring structures such as the reticular thalamic nucleus and the tail of the caudate nucleus were avoided. Slightly more anterior dPul(a) and slightly more posterior dPul sites were very close: nominally – based on grid locations and assuming straight electrode trajectories – only ∼1.1 mm and ∼0.8 mm apart from each other in monkey C and monkey B, respectively. Since we used grids with 0.43 mm holes that tightly matched the outer diameter of the guide tubes, which in turn snugly held the electrodes, and the online anatomical control for the penetration depth using MRI in each session, the final position of electrode tips was quite reproducible within each intended stimulation site (**Table S1**). Note that dPul(a) label in each animal does not imply that these sites were in the anterior (sometimes called oral) pulvinar, which is located further anterior than the majority of our dPul(a) sites, as can be seen in the NMT v2 SARM atlas space (**Figure S2**). In monkey B, however, both dPul(a) and dPul stimulation sites were by ∼1.5-2 mm more anterior than in monkey C in the normalized atlas space, and extended into anterior pulvinar according to the SARM atlas (Hartig et al., 2020).

**Table 1.** Datasets. Number of EPI runs and successful trials per stimulation site, monkey, and session for stimulation in dorsal pulvinar (more anterior and more posterior sites), ventral pulvinar, anterior dorsal lateral intraparietal area (LIPd-a), and posterior dorsal lateral intraparietal area (LIPd-p).

| Stimulation site | Session number | Runs | Monkey C |  | Runs | Monkey B |  |
| --- | --- | --- | --- | --- | --- | --- | --- |
| | | | Current (μA) | Successful trials | | Current (μA) | Successful trials |
| dPul(a) | 1 | 3 | 200 | 69 | 7 | 150-250 | 133 |
|  | 2 | 7 | 150 | 165 | 8 | 150-200 | 126 |
|  | 3 | 8 | 150 | 158 | 7 | 150-250 | 124 |
|  | 4 | 7 | 150 | 140 | 7 | 250 | 112 |
|  | 5 | - | - | - | 7 | 200-250 | 124 |
|  | 6 | - | - | - | 8 | 250 | 147 |
|  | 7 | - | - | - | 6 | 250 | 106 |
|  | 8 | - | - | - | 6 | 250 | 120 |
|  | 9 | - | - | - | 7 | 250 | 136 |
| dPul | 1 | 3 | 250 | 62 | 7 | 250 | 142 |
|  | 2 | 6 | 250 | 120 | 8 | 250 | 165 |
|  | 3 | 6 | 250 | 137 | 7 | 250 | 132 |
|  | 4 | 6 | 250 | 143 | 8 | 250 | 167 |
|  | 5 | 5 | 250 | 90 | 6 | 250 | 128 |
|  | 6 | 6 | 250 | 107 | 7 | 250 | 133 |
|  | 7 | 9 | 250 | 163 | 9 | 250 | 168 |
|  | 8 | - | - | - | 6 | 250 | 114 |
|  | 9 | - | - | - | 8 | 250 | 120 |
| dPul<br>low current<br>stimulation | 1 | 7 | 100 | 129 | - | - | - |
|  | 2 | 6 | 100 | 102 | - | - | - |
|  | 3 | 6 | 100 | 138 | - | - | - |
|  | 4 | 5 | 100 | 108 | - | - | - |
|  | 5 | 6 | 100 | 134 | - | - | - |
|  | 6 | 6 | 100 | 150 | - | - | - |
|  | 7 | 10 | 100 | 240 | - | - | - |
| vPul<br>low current<br>stimulation | 1 | 3 | 100 | 52 | 6 | 100 | 122 |
|  | 2 | 9 | 100 | 239 | 4 | 100 | 81 |
|  | 3 | 5 | 100 | 117 | 7 | 100 | 123 |
|  | 4 | 8 | 100 | 187 | 6 | 100 | 96 |
|  | 5 | 6 | 100 | 137 | 8 | 100 | 168 |
|  | 6 | 7 | 100 | 174 | 8 | 100 | 149 |
|  | 7 | 6 | 100 | 147 | 6 | 100 | 129 |
|  | 8 | - | - | - | 7 | 100 | 115 |
| LIPd-a | 1 | 8 | 100, 250 | 195 | 6 | 250 | 112 |
|  | 2 | 6 | 250 | 157 | 8 | 250 | 129 |
|  | 3 | 6 | 250 | 162 | 8 | 250 | 155 |
|  | 4 | 8 | 250 | 211 | 11 | 250 | 216 |
|  | 5 | 6 | 250 | 154 | 3 | 250 | 24 |
|  | 6 | - | - | - | 10 | 250 | 181 |
|  | 7 | - | - | - | 8 | 250 | 143 |
|  | 8 | - | - | - | 9 | 250 | 131 |
|  | 9 | - | - | - | 7 | 250 | 137 |
|  | 10 | - | - | - | 7 | 250 | 128 |
|  | 11 | - | - | - | 8 | 250 | 153 |
| LIPd-p | 1 | 6 | 250 | 157 | 6 | 250 | 123 |
|  | 2 | 8 | 250 | 199 | 8 | 250 | 156 |
|  | 3 | 5 | 250 | 131 | 7 | 250 | 122 |
|  | 4 | 9 | 250 | 246 | 8 | 250 | 138 |
|  | 5 | 6 | 250 | 155 | 7 | 250 | 128 |
|  | 6 | - | - | - | 6 | 250 | 121 |
|  | 7 | - | - | - | 6 | 250 | 123 |
|  | 8 | - | - | - | 6 | 250 | 111 |
|  | 9 | - | - | - | 6 | 250 | 123 |
|  | 10 | - | - | - | 7 | 250 | 139 |

Note that in both monkeys in several sessions the more anterior dPul site was stimulated with currents lower than 250 μA (**Table 1**). Therefore, dPul(a) datasets serve as an additional controls, and main conclusions about the dorsal pulvinar stimulation effects are derived from the slightly more posterior dPul datasets.

In each animal we also targeted one location in the ventral pulvinar (vPul) for direct comparison with the dorsal pulvinar stimulation. **Figure 1A** and **Figure S3** show the electrode position in an example session of vPul stimulation in a T2-weighted MR image with the slice package aligned to the chamber vertical axis and reconstructed in a T1-weighted MR image, as well as the probability maps of reconstructed electrode tip positions across all vPul sessions in comparison to dPul stimulation sessions, in the AC-PC space. The probability maps indicate that the vPul stimulation sites were located in the ventral part of lateral pulvinar known as PLvl (Kaas and Lyon, 2007) and inferior pulvinar. The local microstimulation-elicited BOLD fMRI activation maps for vPul (and dPul for comparison), together with overlaid electrode probability maps demonstrate that the activation spread around and below the estimated electrode tip, affecting the entire vPul (**Figure S3**).

For localization of the LIP stimulation sites we used the segregation of LIP into a dorsal (LIPd) and a ventral (LIPv) zone (Jung et al., 2021; Saleem and Logothetis, 2006). **Figure 1A** shows examples of measured and reconstructed electrode positions and one coronal section through the center of the electrode tip probability map in AC-PC plane. **Figure S1** (bottom two rows in each monkey) shows the full extent of probability maps of the estimated electrode tip positions across LIP stimulation sessions. In monkey C anterior and posterior LIPd sites were nominally ∼3.4 mm apart from each other. In monkey B the two LIPd sites were nominally ∼5 mm apart.

### 2.6 Behavioral paradigm

For training and scanning, monkeys sat or lay in custom-made horizontal MR compatible primate chairs (Rogue Research, Canada) in a sphinx position with their heads rigidly attached to the chair with a PEEK headholder. Visual stimuli (800×600 pixels resolution) were back-projected onto a custom-made MR compatible tangential screen, placed in front of a monkey (**Figure 1B**). Visual cues and targets were displayed at one of three locations per hemifield (six locations in total) with an eccentricity of 12° of visual angle. Stimulus locations were arranged concentrically around the fixation spot at 0° (mid left), 30° (up left), 150° (up right), 180° (mid right), 210° (down right), and 330°(down left). Trials were presented in a pseudorandomized order to ensure a similar distribution of trial types throughout the duration of each fMRI run (15 min). Monocular eye position was monitored at 60 Hz with a MR-compatible infrared camera (Resonance Technology/Arrington Research, or MRC-Systems 60-M) and was recorded simultaneously with stimulus and timing parameters and digital triggers from the scanner. Stimulus presentation, all behavioral control functions, and synchronization of the behavioral task with the scanner were programmed in MATLAB (R2014a, 64-bit; The MathWorks, Inc., USA) using the Psychophysics Toolbox (Brainard, 1997) and in-house developed software (https://github.com/dagdpz/monkeypsych).

To isolate the BOLD fMRI activity associated with the cue and the memory delay phases during stable fixation from the peri- and post-saccadic responses, we used a slow time-resolved event-related design with long trials. In 33% of the trials, the animals performed a central eye fixation task. In the remaining 66% of trials, they performed delayed memory-guided saccades. **Figure 1B** shows a schematic of the behavioral tasks and the timing of microstimulation. In training sessions the tasks were very similar except for a more variable timing to discourage that animals learn to anticipate the occurrence of trial events. Trials were initiated by fixating the central fixation spot (red dot, 0.25° diameter). Memory-guided saccade trials continued with an eye fixation period (monkey C: 10 s in experimental sessions, 12 - 14 s in training sessions; monkey B: 10 s in experimental sessions, 9.25 - 10.25 s in training sessions). Subsequently, a red filled circle (1° diameter) representing a visuospatial cue was presented for 200 ms either in the contraversive (left, contralateral to the side of stimulation, which was always in the right hemisphere in this study) or the ipsiversive (right, ipsilateral to the side of stimulation) hemifield while the animals were maintaining central eye fixation. The offset of the cue determined the beginning of the memory period (monkey C: 10 s in experimental sessions, 12 - 14 s in training sessions; monkey B: 10 s in experimental sessions and 9.5 - 10.5 s in training sessions) in which the animals were required to remember the cued spatial location and could plan a saccadic eye movement towards this location while maintaining central eye fixation. After the memory period, the fixation spot disappeared (“Go signal”), signaling that the animals were allowed to execute the saccadic eye movement towards the remembered location. If the animals performed a saccade towards the correct location within a radius of 5° to 7° around the center of the cued location within 500 ms, a target stimulus (red, filled circle, 1° diameter) appeared at the cued location to indicate to the animal that the saccade had been performed correctly. After the animal fixated the target stimulus for few seconds (monkey C: 2 s in experimental sessions, 2 - 3 s in training sessions; monkey B: 1.5 s in experimental sessions, 1 - 1.5 s in training sessions) a fluid reward was delivered. In the fixation task, the animals were required to maintain central eye fixation until the end of the trial was signaled by the fixation spot offset (monkey C: 22.2 s in experimental sessions, 26 - 31 s in training sessions; monkey B: 21.7 s in experimental sessions, 20 - 22.5 s in training sessions) in order to get a fluid reward. In training sessions the fluid reward was preceded by a feedback sound. In both tasks, blink allowance time – a period when a fixation break was permitted - was 0.3 s. Trials with fixation breaks exceeding an allowance window of 4° to 5° radius around the fixation spot and trials with incorrect or too slow saccade execution were aborted and not rewarded. In addition to the online eye tracking, a MR-compatible IR camera (MRC-Systems) connected to video-based motion-detection system (Pelco MD2001) were used to train the animals to minimize their jaw movements during the trials and to track jaw movements during scanning. Jaw movements were only allowed in the inter-trial interval (2 s after aborted trials with no reward, 5 s after correct trials with reward). Trials compromised by detected jaw motion were aborted and not rewarded.

In half of the trials, selected pseudo-randomly, the microstimulation (ten 200 ms trains separated by 800 ms) was delivered throughout the 10 s memory period in the memory-guided saccade task, starting at the time of the visual cue offset, or in the corresponding time window in the fixation task (**Figure 1B**). **Table 1** gives an overview of the number of sessions, EPI runs, and successful trials per stimulation site, animal, and sessions for all stimulation sites.

### 2.7 Data analysis

#### 2.7.1 Behavioral analysis

For behavioral analysis, all trials of all experimental sessions were pooled for each stimulation site separately. To test for the effects of microstimulation on task performance, first the overall hit rate for all control trials without stimulation and for all stimulation trials was calculated, respectively, and a Chi-squared test was performed to determine the effect of microstimulation on the frequency of successful trials. If this test revealed a significant difference between the frequency of successful trials in control and stimulation conditions, additional Chi-squared tests were performed for each task (fixation, contraversive memory saccade, ipsiversive memory saccade) separately to test whether microstimulation affected task performance in a task-specific manner. Since trials in control and stimulation conditions were only different starting from stimulation onset, similar analyses were performed on the frequency of trials aborted during and after the stimulation period, in order to detect changes in task performance that were specific to the delivery of current pulse trains and to assess the effect of stimulation on subsequent saccade execution.

All eye movements with a minimum velocity of and a minimum duration of 10 ms were included in the analysis, including small saccades and eye blinks. The time when eye velocity passed the 15 °/s threshold determined eye movement onset. Movement offset was defined as the point in time when eye velocity dropped below 10 °/s. Using these detection criteria, the number of eye movements during the stimulation period was extracted for each trial and averaged across all trials of each experimental condition (control and stimulation for fixation, ipsiversive memory saccade, and contraversive memory saccade). To test whether stimulation influenced fixation behavior, the number of eye movements during the stimulation period was submitted to a two-way ANOVA with the factors task (fixation, ipsiversive saccade, contraversive saccade) and stimulation (control, stimulation). Further post-hoc t-tests were performed if the ANOVA revealed significant effects of stimulation. To investigate the effects of microstimulation on saccade execution, saccade latencies were extracted from all saccade trials and tested in a two-way ANOVA design with factors task (ipsiversive saccade, contraversive saccade) and stimulation (control, stimulation). Significant effects of stimulation on saccade latencies were further tested using post-hoc t-tests.

#### 2.7.2 NMT v2 template and CHARM/SARM atlas utilization

To facilitate the comparability between the results in two individual animals, and to promote standardization for data sharing, we utilized recently published NMT v2 template and associated hierarchical cortical CHARM and subcortical SARM atlases (Hartig et al., 2020; Jung et al., 2021; Seidlitz et al., 2018). As compared to the AC-PC space with the anterior commissure (AC) as an origin, NMT v2 is in a stereotaxic space, with the origin at the intersection of the midsagittal plane and the interaural line. We used the approach described in AFNI’s MACAQUE_DEMO that uses @animal_warper to nonlinearly coregister each animal’s skull-stripped T1-weighted volume (0.5 mm resolution) in the individual AC-PC space to the symmetric NMT template (https://afni.nimh.nih.gov/pub/dist/doc/htmldoc/nonhuman/macaque_demos/demo_task_fmri.html). Using resulting forward and inverse warps, CHARM and SARM atlases were transformed to the individual AC-PC space and exported to BrainVoyager VOI format (volume-of-interest) for further processing – such as the extraction of the ROI timecourses using atlas regions (section 2.7.5) and visualization. Additionally, electrode localization probability maps and statistical activation maps in the individual AC-PC space were warped into the NMT template space (with 3dAllineate and 3dNwarpApply), and NMT v2 template-aligned atlas ROIs were converted to BrainVoyager VOI format for visualization purposes.

For conversion to NIfTI (for use with AFNI) and back to the BrainVoyager VMR (volume anatomical) and VMP (volume statistical map) formats we used NeuroElf toolbox (v1.0, http://neuroelf.net/), custom MATLAB code, and the NIFTI toolbox in MATLAB (Jimmy Shen 2021. Tools for NIfTI and ANALYZE image, https://www.mathworks.com/matlabcentral/fileexchange/8797-tools-for-nifti-and-analyze-image, MATLAB Central File Exchange. Retrieved April 4, 2021).

#### 2.7.3 Estimation of electrode tip positions

For assessment of the variability in electrode tip positions across sessions, for each experimental session the location of the electrode tip was estimated based on the respective T2-weighted image (0.25 mm resolution in plane) acquired with the slice package aligned to the chamber vertical axis (see section 2.4) using BrainVoyager. For each session, a sphere (radius 0.5 mm) was created around the estimated electrode tip position and a probability map across all sessions for this stimulation site was created based on the resulting volumes of interest showing the probability of overlapping 0.25 mm voxels. The resulting probability maps were then transformed into individual AC-PC space and overlaid onto the high-resolution, full-brain T1-weighted anatomical image of each monkey. For better comparability, to avoid being hampered by the individual brain anatomy and size differences, these maps were also warped into a common NMT v2 template space (**Figure S2**), using AFNI, NeuroElf, and NIFTI toolbox, as described in the section above.

#### 2.7.4 Functional data processing

The first four EPI volumes were excluded from functional data analysis in order to eliminate transient effects of magnetic saturation. Preprocessing was performed using MATLAB (R2014a, 64-bit; The MathWorks, Inc., USA) and the NeuroElf toolbox (v1.0, http://neuroelf.net/). EPI data of each run was preprocessed using slice time correction and a high-pass temporal filter with a cut-off of three cycles per 15 min run. In addition, 3D motion correction was performed using the first functional volume of the first run included into the analysis as a reference. Coregistration and volume time course computation was done using BrainVoyager. First, the T2-weighted anatomical image acquired in-plane with EPI slices in each session was coregistered to the high-resolution full-brain T1-weighted anatomical scan in the AC-PC space. Then, EPI runs were coregistered to the respective T2-weighted anatomical image in the AC-PC space using rigid body transformations with automated initial alignment followed by careful manual fine-tuning of the resulting alignment based on anatomical sulcal landmarks. The use of within-session in-plane anatomy covering the same volume as the EPI facilitated the coregistration. **Figures S4-S7** demonstrate the correspondence between T1-, T2- and T2*-weighted EPI images. Finally, volume time courses were computed in the individual subject AC-PC space using 1 mm x 1 mm x 1 mm voxel size, intersecting the volume time course data a with a mask only including voxels within the brain; this resulted in ∼89,000 voxels in monkey C and ∼110,000 voxels in monkey B as the total number of voxels considered for FDR correction. A 3D 1.5 mm x 1.5 mm x 1.5 mm Gaussian kernel was applied for spatial smoothing of the volume time courses.

#### 2.7.5 GLM, ROI definition, and event-related averaging

General linear models (GLMs) were computed in MATLAB using the NeuroElf toolbox. For successful trials, all trial events except the initial fixation period – (1-4) contraversive/ipsiversive cue and memory delay periods with and without stimulation, (5-6) contraversive/ipsiversive saccades and subsequent peripheral target fixation, (7-9) the corresponding time windows in fixation trials, (10) reward delivery and the subsequent inter-trial interval – were extracted and used as predictors for the GLM. In addition, there was one predictor for aborted trials (11). The initial fixation served as GLM implicit baseline condition. Based on these 11 task event predictors, design matrices were created using convolution with the macaque hemodynamic response function ((Kagan et al., 2010), time to positive peak: 3 s, time to negative peak: 10 s) in order to compute GLMs. For each run, 6 motion correction parameters were included as confound predictors. Volume time courses were z-score normalized. For each stimulation site, data from all sessions and runs were combined and analyzed using fixed-effects GLM, using volumetric data in the individual subject AP-PC space.

For each animal and stimulation site the event-related statistical t-maps contrasting beta values in all three task conditions during the stimulation in the cue/memory or the matching fixation epochs with the corresponding control (no stimulation) conditions [“microstim > no microstim” contrast: contraversive saccade microstim + ipsiversive saccade microstim + fixation microstim - contraversive saccade - ipsiversive saccade – fixation] were used to identify regions activated by the microstimulation irrespective of the task. Based on comparing individual task condition contrasts and the results of ROI analysis, the combined contrast approximates the “OR” combination between tasks, i.e. a region is often significant in the map if there is robust stimulation-elicited activation in at least one task condition. We applied FDR-corrected threshold of *q* < 0.05 and a cluster threshold of k ≥ 20 1-mm^3^ functional voxels. For conjunction analysis, the highest t-value of the two contrast statistical maps corresponding to a FDR-corrected threshold *q* < 0.05 was used.

We defined two types of ROIs. 1) Functionally defined regions of interest (“stimulation effect ROIs”) were derived automatically using an intersection of the combined CHARM-SARM atlas ROIs warped to each individual AC-PC brain (section 2.7.2) and the “microstim > no microstim” activation maps. Note that the ROI selection based on the combined contrast does not introduce any specific bias towards any task condition, and hence is not circular for assessing task-dependent effects. We limited the analysis to 149 regions (79 cortical and 70 subcortical labels) in each hemisphere from the most detailed level 6 parcellation, resulting in 298 anatomical ROIs (**Table S2**, **Figure S8**). These ROIs covered the occipital, temporal, parietal and frontal cortical lobes relevant for the visuomotor tasks, and subcortical structures such as basal ganglia, the thalamus and the cerebellum. For each atlas ROI, the activation map was masked by the ROI volume, and activation clusters of at least 24 0.5-mm^3^ anatomical voxels (i.e. 3 1-mm^3^ functional voxels) were detected using NeuroElf ClusterTable method (https://neuroelf.net/wiki/doku.php?id=vmp.clustertable). When more than one activation cluster was found within the atlas ROI, they were combined, so that only a single resulting activation ROI per atlas ROI was defined. Out of 149 cortical and subcortical labels, 124 stimulation effect ROIs were found at least once across all 10 datasets – 5 stimulation sites in each monkey (76 cortical, 48 subcortical labels, **Table S2**, **Table S3**). 2) In addition, purely atlas-based individually-warped ROIs were defined, regardless of the stimulation effect, as a supplemental analysis approach (called “atlas ROIs”). For these ROIs, few deep subcortical labels that only partially overlapped with the functional volume coverage for a given dataset were excluded. The EPI dropout regions due to electrode-induced susceptibility artifacts were defined on the mean EPI volume for each dataset using signal intensity thresholding (**Figure S9**, **Figure S10**) and were also excluded from further ROI analysis, for both types of ROIs.

**Table 2.** Summary of activated ROIs. Number of ROIs activated by the microstimulation in each stimulation site, and stimulation effect correlation between hemispheres, across stimulation effect ROIs activated in either hemisphere, or across all atlas ROIs.

| Subject | Stimulation site | Left hemisphere (LH) | Right hemisphere (stimulated, RH) | ROIs in LH not present in RH | LH-RH correlation in stim. effect ROIs R (p) | LH-RH correlation in atlas ROIs R (p) |
| --- | --- | --- | --- | --- | --- | --- |
| Monkey C | dPul(a) | 34 | 67 | 1 | **0.41 (<0.001)** | **0.47 (p<0.001)** |
|  | dPul | 35 | 65 | 4 | **0.39 (<0.001)** | **0.57 (p<0.001)** |
|  | LIPd-a | 23 | 46 | 6 | **0.56 (<0.001)** | **0.38 (p<0.001)** |
|  | LIPd-p | 78 | 81 | 20 | **0.35 (<0.001)** | **0.62 (p<0.001)** |
|  | vPul | 14 | 35 | 2 | **0.51 (<0.001)** | **0.36 (p<0.001)** |
| Monkey B | dPul(a) | 9 | 48 | 1 | 0.15 (0.102) | **0.32 (p<0.001)** |
|  | dPul | 11 | 70 | 3 | -0.05 (0.576) | **0.34 (p<0.001)** |
|  | LIPd-a | 34 | 54 | 4 | **0.54 (<0.001)** | **0.45 (p<0.001)** |
|  | LIPd-p | 20 | 43 | 3 | **0.62 (<0.001)** | **0.49 (p<0.001)** |
|  | vPul | 17 | 28 | 2 | **0.47 (<0.001)** | **0.25 (0.002)** |
| Mean |  | 27.5 | 53.7 | 4.6 | 0.39 | 0.42 |
| Median |  | 21.5 | 51 | 3 | 0.44 | 0.42 |
| SD |  | 20.2 | 16.8 | 5.6 | 0.21 | 0.12 |

**Table 3.** Stimulation effects in different task conditions. Significant p-values (p<0.05) for the two-way repeated measures ANOVAs and t-tests are in bold font. p – p-value, F – F-statistics, t – t-value. See **Table S4** for the statistics on atlas ROIs.

|  | Main<br>effect<br>task | Main<br>effect<br>stim. | Task<br>×<br>stim. | Contra<br>vs ipsi | Contra<br>vs fix | Ipsi vs<br>fix | Main<br>effect<br>task | Main<br>effect<br>stim. | Task<br>×<br>stim. | Contra<br>vs ipsi | Contra<br>vs fix | Ipsi vs<br>fix |
| --- | --- | --- | --- | --- | --- | --- | --- | --- | --- | --- | --- | --- |
| Format | F<br>p | F<br>p | F<br>p | t<br>p | t<br>p | t<br>p | F<br>p | F<br>p | F<br>p | t<br>p | t<br>p | t<br>p |
|  | Left hemisphere |  |  |  |  |  | Right hemisphere (stimulated) |  |  |  |  |  |
| Monkey C |  |  |  |  |  |  |  |  |  |  |  |  |
| dPul(a) | 6.94 | 289.87 | 3.19 | 0.85 | 4.23 | 1.35 | 18.59 | 233.88 | 0.95 | 0.63 | 1.73 | 0.65 |
|  | **0.002** | **0.000** | 0.048 | 0.400 | **0.000** | 0.187 | **0.000** | **0.000** | 0.390 | 0.533 | 0.088 | 0.517 |
| dPul | 20.42 | 34.58 | 8.66 | 3.78 | 2.05 | -2.40 | 15.82 | 172.97 | 6.73 | 3.61 | 1.69 | -2.09 |
|  | **0.000** | **0.000** | **0.000** | **0.001** | **0.049** | **0.023** | **0.000** | **0.000** | **0.002** | **0.001** | 0.096 | **0.041** |
| LIPd-a | 1.98 | 49.44 | 4.57 | 2.88 | 1.99 | -1.40 | 31.69 | 126.61 | 2.23 | 2.19 | 0.79 | -1.34 |
|  | 0.152 | **0.000** | **0.016** | **0.009** | 0.060 | 0.178 | **0.000** | **0.000** | 0.114 | 0.034 | 0.434 | 0.186 |
| LIPd-p | 134.81 | 267.84 | 30.76 | 7.45 | 3.06 | -5.17 | 31.44 | 180.51 | 17.29 | 5.08 | 1.28 | -5.05 |
|  | **0.000** | **0.000** | **0.000** | **0.000** | **0.003** | **0.000** | **0.000** | **0.000** | **0.000** | **0.000** | 0.205 | **0.000** |
| vPul | 31.12 | 76.14 | 0.73 | -1.82 | 0.15 | 0.90 | 11.29 | 97.36 | 2.71 | 2.40 | 1.13 | -1.17 |
|  | **0.000** | **0.000** | 0.492 | 0.092 | 0.886 | 0.387 | **0.000** | **0.000** | 0.074 | **0.023** | 0.269 | 0.251 |
| Monkey B |  |  |  |  |  |  |  |  |  |  |  |  |
| dPul(a) | 0.20 | 550.07 | 1.18 | -2.40 | -1.02 | 0.01 | 10.80 | 138.61 | 2.66 | 0.66 | 2.58 | 1.71 |
|  | 0.818 | **0.000** | 0.336 | **0.048** | 0.341 | 0.995 | **0.000** | **0.000** | 0.075 | 0.515 | **0.013** | 0.094 |
| dPul | 16.52 | 23.47 | 10.69 | -1.72 | 3.00 | 4.12 | 24.23 | 97.73 | 21.28 | -2.07 | 4.18 | 6.38 |
|  | **0.000** | **0.001** | **0.001** | 0.116 | **0.013** | **0.002** | **0.000** | **0.000** | **0.000** | **0.042** | **0.000** | **0.000** |
| LIPd-a | 43.27 | 38.63 | 4.13 | -2.66 | -2.50 | -0.77 | 40.92 | 70.60 | 14.62 | -7.09 | -4.31 | -2.36 |
|  | **0.000** | **0.000** | **0.021** | **0.012** | **0.018** | 0.448 | **0.000** | **0.000** | **0.000** | **0.000** | **0.000** | **0.022** |
| LIPd-p | 6.17 | 61.79 | 6.71 | 1.20 | -1.95 | -6.52 | 34.18 | 114.87 | 10.58 | -2.44 | -3.70 | -3.08 |
|  | **0.005** | **0.000** | **0.003** | 0.246 | 0.066 | **0.000** | **0.000** | **0.000** | **0.000** | **0.020** | **0.001** | **0.004** |
| vPul | 11.19 | 138.76 | 0.61 | 0.68 | -0.43 | -1.17 | 9.95 | 150.82 | 1.74 | 1.62 | -0.21 | -1.84 |
|  | **0.000** | **0.000** | 0.548 | 0.507 | 0.677 | 0.263 | **0.000** | **0.000** | 0.185 | 0.118 | 0.839 | 0.078 |

For each stimulation site dataset, BOLD time courses of each ROI were extracted using MATLAB and the NeuroElf toolbox. For event-related averaging, BOLD time course data were interpolated to the temporal resolution of 1 s. Then, for each trial event-related time course in %BOLD signal change was computed relative to the individual trial baseline, which was defined as the mean BOLD activity in the last 4 s of the initial fixation period before cue presentation in memory-saccade trials or the corresponding time window in fixation trials. Then, BOLD time courses were averaged across trials per each task and stimulation condition.

#### 2.7.6 Region of interest analysis within and across ROIs

For ROI analysis, the mean response amplitude in %BOLD signal change between 2 and 9 s of the memory period or the corresponding time epoch in fixation trials was calculated in each ROI. The start of the analysis epoch was selected based on the observed fast responses to visual cues and the stimulation onset (time to peak 3-4 s); the end of the epoch was selected so that the last second of the delay period, which can be potentially contaminated by the saccade response, was omitted.

Within each ROI, we calculated the main effect of task (memory saccade right - ipsiversive, memory saccade left – contraversive, fixation), the main effect of stimulation (no stimulation - control, stimulation) and the task × stimulation interaction, using a two-way ANOVA across single trial response amplitudes. To investigate whether the magnitude of stimulation effect on BOLD responses depended on the task, we calculated the stimulation effect for each task and ROI as the difference between the mean BOLD response in the respective stimulation and control condition. Stimulation effects were then analyzed across ROIs using a two-way repeated measures ANOVA (rmANOVA) with the factors task (ipsiversive memory saccade, contraversive memory saccade, and fixation) and stimulation (no stimulation, stimulation) for each hemisphere separately. Significant effects were further investigated using paired-sample t-tests. Outlier ROIs with a mean BOLD response deviating more than 2.5 standard deviations from the mean BOLD response in control conditions across all ROIs for a given dataset were excluded from further across ROI analysis.

For each ROI we assessed the degree of contraversive selectivity, i.e., whether the BOLD response of an ROI was stronger to visual cues/saccade preparation presented in/towards spatial locations in the hemifield contralateral to the side of stimulation (contraversive memory saccade). To calculate contraversive selectivity index (CSI), we subtracted the mean BOLD response in ipsiversive trials from the mean BOLD response in contraversive trials for control and stimulation conditions for each ROI. Positive CSI values reflect contraversive tuning (contralateral for the stimulated hemisphere). Negative CSI values reflect ipsiversive tuning, i.e. preference towards the side of space that is ipsilateral to the side of stimulation (ipsiversive memory saccade condition). Note that the ipsiversive tuning is contralateral tuning for the non-stimulated (left) hemisphere.

To test whether stimulation effects on BOLD responses depended on the initial spatial tuning of ROIs, we categorized ROIs of both hemispheres according to their contraversive selectivity as indicated by CSI values for control trials (positive vs. negative values) and tested the differences between the mean BOLD response in the respective stimulation and control condition of each task in a two-way mixed ANOVA with between-subjects factor spatial tuning: contraversive selectivity, ipsiversive selectivity – and within-subjects factor task condition: ipsiversive memory saccade, contraversive memory saccade, fixation (the “subject” here being each ROI). Interaction effects were further investigated using paired-sample t-tests to compare stimulation effects between tasks for ROIs with contraversive and ipsiversive tuning, separately.

To test if microstimulation-elicited enhancement was best described by adding activity to the initial BOLD response in the control condition, or whether stimulation leads to multiplication of the BOLD response in the control condition, we fitted the relationship between BOLD response amplitudes (RA) with and without stimulation with an additive model: *RA_stim_ = RA_cont_ + a*, an additive model scaled by the control (no stimulation) response amplitude: *RA_stim_ = RA_cont_ + a*(1 – b*RA_cont_)*, and a multiplicative model *RA_stim_ = a*RA_cont_*, for each task and hemisphere separately, using the *fit.m* function with *NonlinearLeastSquares* method in MATLAB. We then evaluated the resulting adjusted R-squared values as indicators of goodness of fit.

#### 2.7.7 Surface maps

The individual monkeys’ left and right hemisphere cortical surfaces were reconstructed using standard BrainVoyager segmentation and surface reconstruction steps. The surface maps show the activity patterns extracted from the volume-based GLM contrasts (voxel-based map, “VMP” in BrainVoyager terminology), sampled along the dilated cortical gray matter/white matter boundary mesh vertex normal, from −0.5 mm to 1.5 mm, and mapped on the reconstructed and inflated surface boundary (“VMP”->”SMP” surface map conversion). Note that some volume activations above and below this 2 mm layer (e.g. in the gray matter in the arcuate sulcus and in the superior temporal sulcus) may not be visible on surface maps.

NMT v2 template surfaces were created by converting the GIFTI files rh.white_surface.rsl.gii and lh.white_surface.rsl.gii provided with the atlas (Jung et al., 2021) using BrainVoyager GIFTI converter 1.7 plugin (https://support.brainvoyager.com/brainvoyager/available-tools/86-available-plugins/63-gifti-conversion-plugin), by realigning and rescaling the resulting surface files to NMT v2 volume template and then inflating them in BrainVoyager.

## 3 Results

### 3.1 Task performance and eye movements

During scanning, monkeys performed two oculomotor tasks: central fixation and delayed memory saccades (**Figure 1B**). The memory trials were separated into left, i.e. contraversive (to the side of stimulation) and right (ipsiversive) saccades for further analysis. In half of the trials, the microstimulation was applied for 10 s, corresponding to the memory delay period, or in the corresponding 10 s in fixation only trials. Stimulation of several pulvinar and LIP sites was conducted in separate sessions, resulting in 5 datasets per animal (dPul(a), dPul, LIPd-a, LIPd-p, vPul, see section 3.2).

Details of the behavioral analysis are shown in the Supplementary Results and **Figures S11-S14**. In short, both pulvinar and LIP stimulation did not lead to major impairments in the fixation or memory saccade task. Only LIPd-p stimulation was associated with a slightly lower hit rate for memory saccades to the ipsiversive hemifield in monkey C. In monkey B, only dPul(a) stimulation led to a statistically significant but modest impairment in the execution of memory saccades to the contraversive hemifield (from 4% to 12%). Most consistently, both pulvinar and LIP stimulation led to a modest increase in saccade latencies (∼20-30 ms). More specifically, dPul and LIPd-p stimulation in monkey C were associated with significantly longer latencies only for contraversive saccades, and a similar effect was found for LIPd-a and vPul stimulation in monkey B, while dPul(a) stimulation in monkey B increased saccade latencies towards both the contraversive and the ipsiversive hemifield (**Figure S14**).

For the fMRI analysis, aborted trials were modeled in the GLM to account for the introduced variance, and hence might contribute to the statistics of activation patterns, but for the main statistical contrasts we only used successfully completed and rewarded trials (**Table 1**). The aborted trials were also excluded from event-related averaging of BOLD signal timecourses in the region-of-interest (ROI) analysis. This approach, the overall mild and not always consistent effects of the microstimulation on behavior, as well as asymmetrical (stronger in the stimulated hemisphere) patterns of microstimulation-elicited activation (see section 3.3) all support the notion that the fMRI results were not majorly affected by the changes in performance and eye movements.

### 3.2 BOLD activity induced by pulvinar and LIP microstimulation

#### 3.2.1 Activation at and around stimulated sites

Microstimulation elicited strong local BOLD activity in the targeted dPul or LIP (**Figure 2**). The extent of activation around and below the electrode tip was substantially larger than expected from a passive current spread within 0.5-1 mm radius at 100-250 μA (Klink et al., 2021; Thier and Andersen, 1998). Dorsal pulvinar stimulation affected mainly the medial pulvinar (MPul, including the anterior portion of the medial pulvinar, APul), but the edges of activation extended ventromedially to the superior colliculus (SC), and a weaker activity was found in the adjacent lateral pulvinar and ventral pulvinar (vPul) (**Figure 2A,B,C**, see **Figure S15** for dPul activation close-up with overlaid atlas regions). Stimulation of LIPd led to an extensive activation along the intraparietal sulcus, including dorsal and ventral LIP (LIPd and LIPv) and adjacent cortical regions (**Figure 2D**). Note that the BOLD activity above the electrode tip in the vicinity of the stimulating electrode shank (which was present both in stimulation and no stimulation trials) cannot be measured due to signal dropouts caused by the magnetic susceptibility artefacts around metallic objects (**Figure S9**, **Figure S10**).

#### 3.2.2 Distal activation patterns

Both the pulvinar and parietal unilateral stimulation elicited widespread distal activations, in both hemispheres. **Figure 3A,B** shows statistical t-maps reflecting BOLD activity enhancement elicited by stimulation on the inflated brain surface based on the animal’s individual anatomy. **Figure 3C** and **Figure S16** show the same maps in the individual AC-PC volume space on coronal sections. **Figure S17** shows that activations derived from the combined microstimulation > no microstimulation contrast across all three task conditions were similar to the maps derived from the fixation trials only. **Figure S18** shows the combined contrast activations in the NMT v2 space, to facilitate the comparison between monkeys. In the stimulated hemisphere, consistently across the two animals dPul stimulation activated primary and extrastriate visual cortex, an extensive swath encompassing dorsal and ventral superior temporal sulcus (STS) areas (dorsal/fundus: IPa, TAs, FST, PGa, TPO; ventral: TEO, TE, MT), insula, parietal cortex (area 7a, MST and several regions in the parieto-occipital and intraparietal sulcus: PO, LOP, MIP, LIP, VIP), and frontal and prefrontal cortex (areas 12, 46d/v, 8A/Bs, 44, 45a/b, dorsal premotor area PMd, ventral premotor area F5) (**Figure 3A**, **Table S2**). Subcortical regions such as superior colliculus, amygdala, putamen and cerebellum, as well as caudate nucleus and ventral thalamus in monkey C, and lateral and mediodorsal thalamus in monkey B, were also activated. Enhanced activity was also found in a subset of these cortical and subcortical regions in the opposite hemisphere, although to a lesser extent: visual cortex (both monkeys), frontal cortex (mainly monkey C), STS and IPS (monkey C), and caudate nucleus (monkey B). The activation in the opposite hemisphere suggests a polysynaptic transmission of microstimulation-elicited excitation, since to our knowledge there are no direct connections from the pulvinar to cortical regions in the contralateral hemisphere.

**Figure 3.**
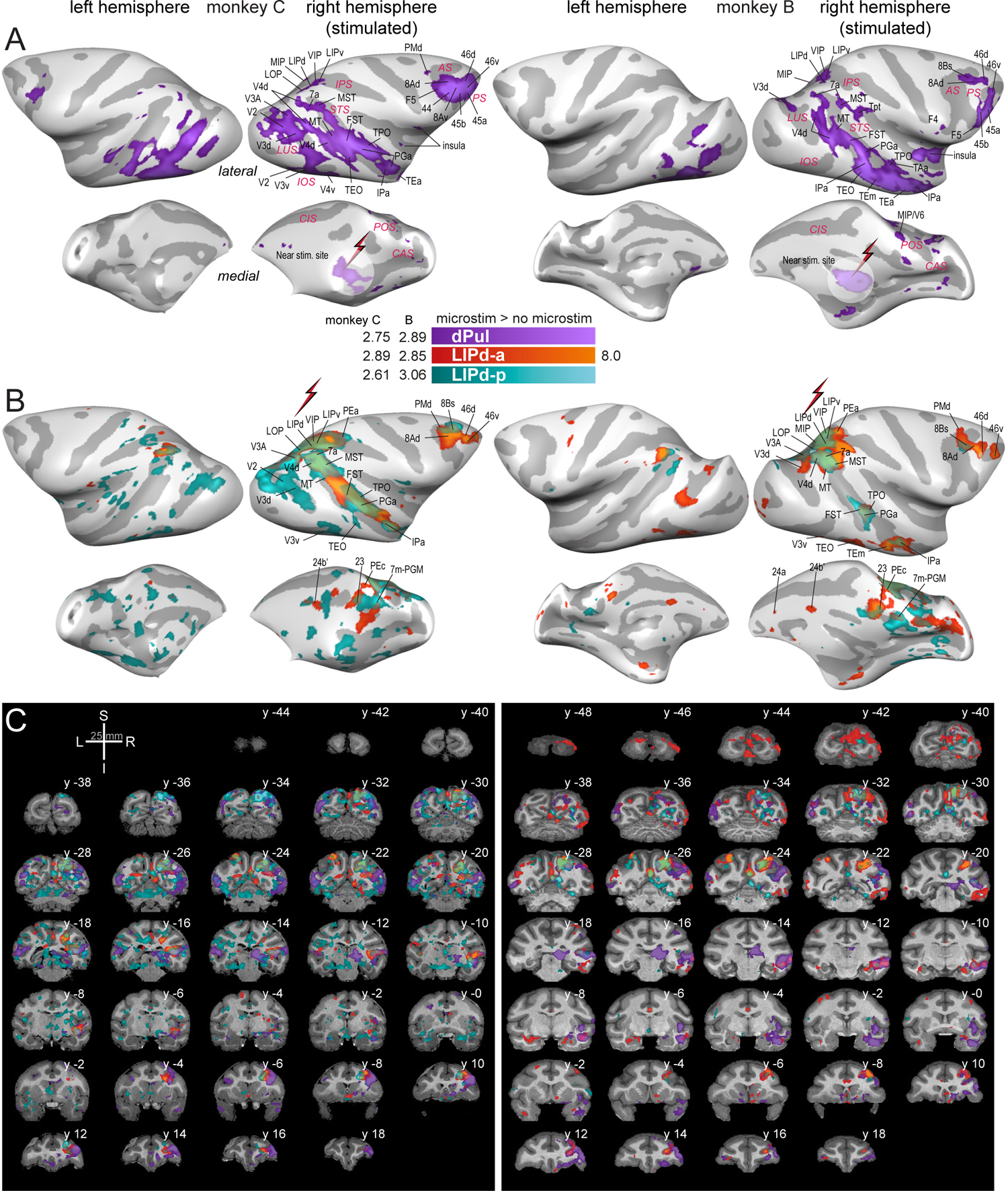
Dorsal pulvinar and LIPd stimulation effects. Statistical t-maps showing BOLD activation elicited by microstimulation on the inflated cortical surface (lateral and medial view) for **(A)** dPul stimulation and **(B)** LIPd stimulation (LIPd-a and LIPd-p maps overlaid). Major sulci and activated regions are labeled. **(C)** dPul and LIPd stimulation activation maps in the volume space, coronal sections (2 mm spacing). L – left, R – right, S – superior, I – inferior, y - distance from AC-PC origin along the anterior/posterior plane in mm. See **Figure S16** for the same maps plotted separately for dPul and LIPd sites.

The above summary is derived from the main pulvinar datasets (dPul). Note that in dPul(a) datasets in many sessions lower 150 – 200 μA currents were used and in monkey C only 4 sessions were included, therefore we treat these datasets as supplementary (but present summary ROI statistics in subsequent analyses for completeness). **Figure S19** compares the stimulation effects of dPul and dPul(a) sites. In both monkeys, the two sites activated a large number of frontal, parietal, STS and extrastriate visual regions. At the same time, in each monkey there was a tendency for more extensive activity in the extrastriate visual cortex and inferotemporal areas TEO and TE with the slightly more posterior stimulation site (Methods), while parietal and frontal (the latter only in monkey C) regions were more strongly activated by dPul(a).

Similar to dPul stimulation, in both animals LIPd stimulation activated a large number of cortical areas in the stimulated hemisphere: primary (V1) and extrastriate visual cortex (V2, V3, V4), STS (IPa, PGa, TPO, FST, MT, MST), other areas along and around the IPS (area 7, PO, LOP, LIP, MIP, VIP, PEa), prefrontal cortex (45, 8A/B, 46), and PMd (**Figure 3B**), as well as putamen and cerebellum (**Figure S16B**). Additionally, LIPd stimulation caused strong activation in the medial posterior parietal area 7m/PGM, as well as in the dorso-caudal area PEc, V6, the posterior (area 23b) and the anterior cingulate (areas 24a/b/c). In the opposite hemisphere, homotopic IPS regions, medial parietal regions, several areas in the STS and anterior cingulate were activated, as well as the insula in monkey C and the dorsal premotor area PMd in monkey B. The homotopic activation of the opposite IPS regions stands in a contrast to the dPul stimulation, which did not elicit the activation in the opposite pulvinar.

#### 3.2.3 Overlap of pulvinar- and LIP-elicited effective connectivity

We next analyzed the overlap of dPul- and LIPd-elicited activations. **Figure 4** presents the overlaid dPul and LIPd activation maps (**4A**) and the conjunction maps (**4B**) in the stimulated hemisphere, together with the individually-warped CHARM atlas regions in and around STS. **Figure S20** shows similar activation maps derived from the fixation trials only. The left (non-stimulated) hemisphere is not shown since there was only a minor overlap. Beyond LIP and adjacent parietal regions, the shared effective connectivity between dPul and either of the two LIPd sites was strongest in the fundus and the dorsal bank of STS, in caudal, middle, and anterior parts of the sulcus (areas MST, MT, FST, IPa, PGa, TPO) and, only for the LIPd-a, in pre(frontal) cortex (areas 8A, 8Bs, 46d/v). These results implicate the dorsal bank and the fundus of STS as a main shared “node” of the dorsal pulvinar-parietal circuitry.

**Figure 4.**
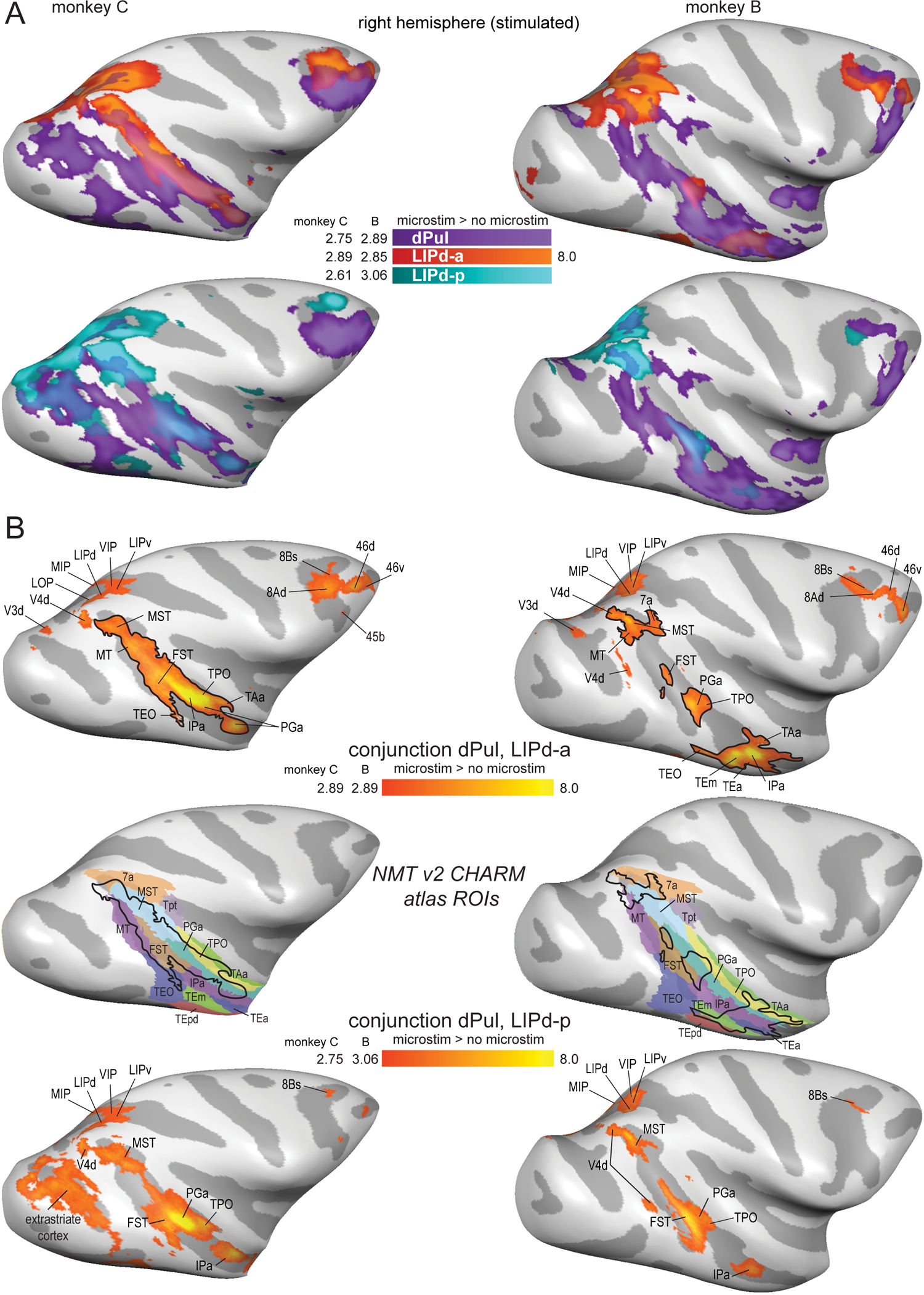
Overlap of thalamic and parietal activation. **(A)** Statistical t-maps showing BOLD activation on the inflated cortical lateral surface for dPul stimulation overlaid together with LIPd-a or dPul-LIPd-p stimulation. **(B)** Conjunction maps showing the overlap of activation elicited by the stimulation of dPul with LIPd-a (top row) and dPul with LIPd-p (bottom row). Major overlap regions are labeled. For reference, middle row shows the NMT v2 CHARM regions in STS, warped to the individual brain anatomy together with the overlap contours from the top row (dPul & LIPd-a conjunction).

The widespread activation along the ventral and dorsal bank of STS elicited by pulvinar and LIP stimulation might have encompassed several brain regions (“face patches”) that have previously been functionally characterized to be face-specific (Moeller et al., 2008; Tsao et al., 2006). A representative face patch pattern from an example monkey (Schwiedrzik et al., 2015) partially overlapped with these activations (most consistently the middle face patch MF, as well as lateral patches PL, ML and frontal patch PO; **Figure 5**).

**Figure 5.**
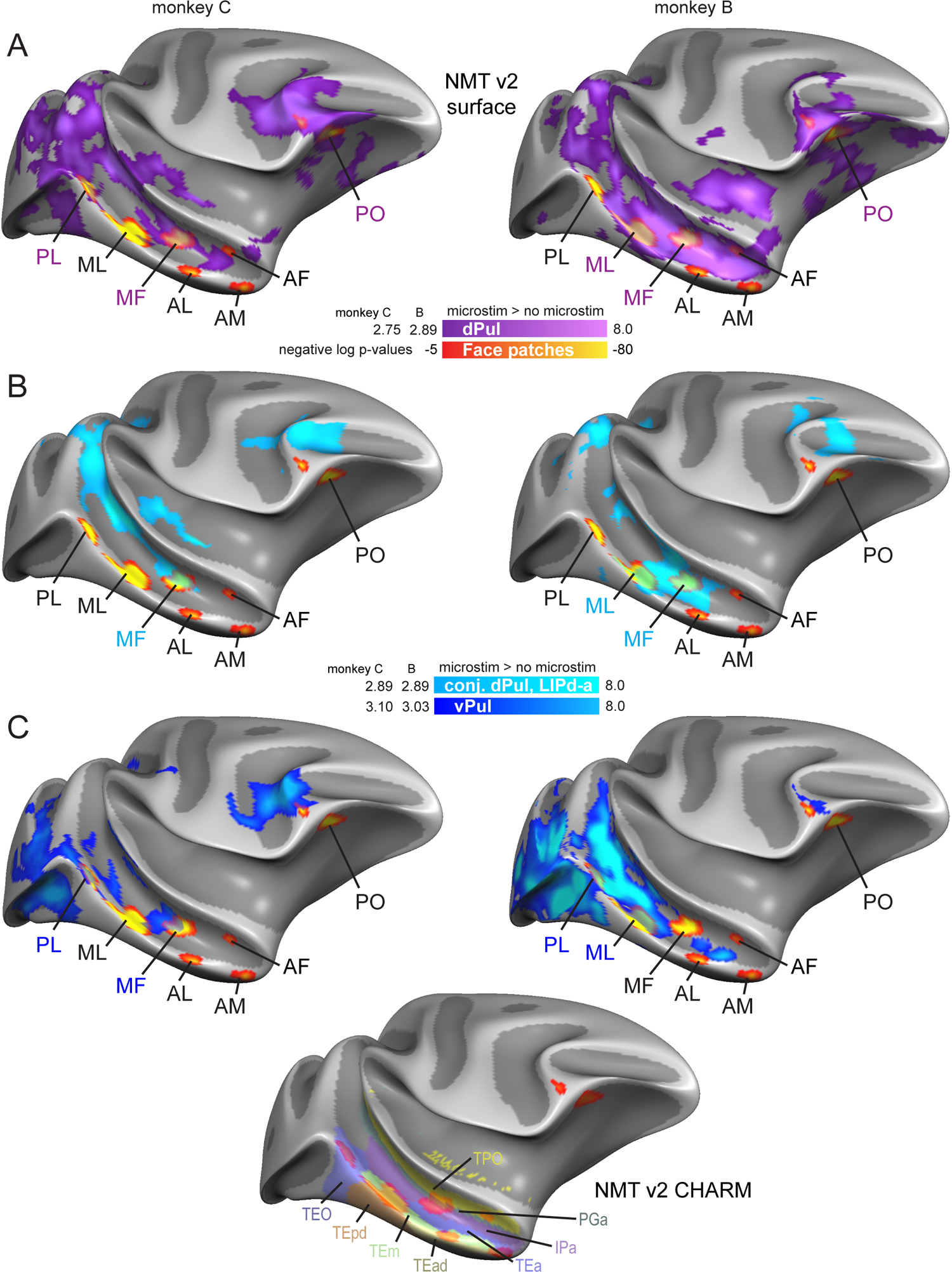
Putative relationship to the face patch system. Example face patches (Schwiedrzik et al., 2015) displayed on partially inflated NMT v2 right hemisphere white matter surfaces, together with **(A)** Dorsal pulvinar stimulation-elicited map, **(B)** Conjunction of dPul and LIPd-a stimulation map, **(C)** Ventral pulvinar stimulation map. The inset below shows CHARM atlas STS regions. Face patch labels overlapping with stimulation-elicited activations are colored correspondingly.

#### 3.2.4 Localized specificity of stimulation and distal connectivity

Despite a substantial local spread of BOLD activation outside of the immediately stimulated vicinity - that might be partially due to fMRI signal smearing or subthreshold excitability changes (Tehovnik et al., 2006), there are indications that the distal stimulation effects were quite site-specific. Firstly, the posterior and anterior LIP sites elicited partially distinct distal effects, especially in prefrontal regions (please refer back to **Figure 3B**). Namely, LIPd-a stimulation caused extensive activity in prefrontal cortex (8A, 45, and 46), but activated only few areas in visual cortex. Conversely, prefrontal activation during LIPd-p stimulation was mainly restricted to medial area 8B and area 46 whereas activation in visual cortex was more extensive. Secondly, we directly compared the activation patterns resulting from stimulation of dPul and adjacent, but anatomically and functionally distinct ventral pulvinar, vPul (which encompasses the ventral division of lateral pulvinar, PLvl, and inferior pulvinar). Since vPul is smaller compared to dPul, and since we wanted to limit the spread of activation into the dorsal pulvinar, we stimulated vPul with a lower current strength (100 μA). Despite a partial overlap of local activations around the stimulation sites (but with clearly separated peaks of activation), the comparison of dPul and vPul distal stimulation effects showed divergent effective connectivity, largely consistent with their known anatomical connectivity. Specifically, vPul showed strong es-fMRI connectivity to early ventral visual areas and the ventral bank of STS (**Figure 6**, **Figure S21**). The stimulation of vPul also led to a distinct activation peak in SC, unlike the more diffuse dPul activation spread encompassing SC on the fringe but not exhibiting a separate SC peak.

**Figure 6.**
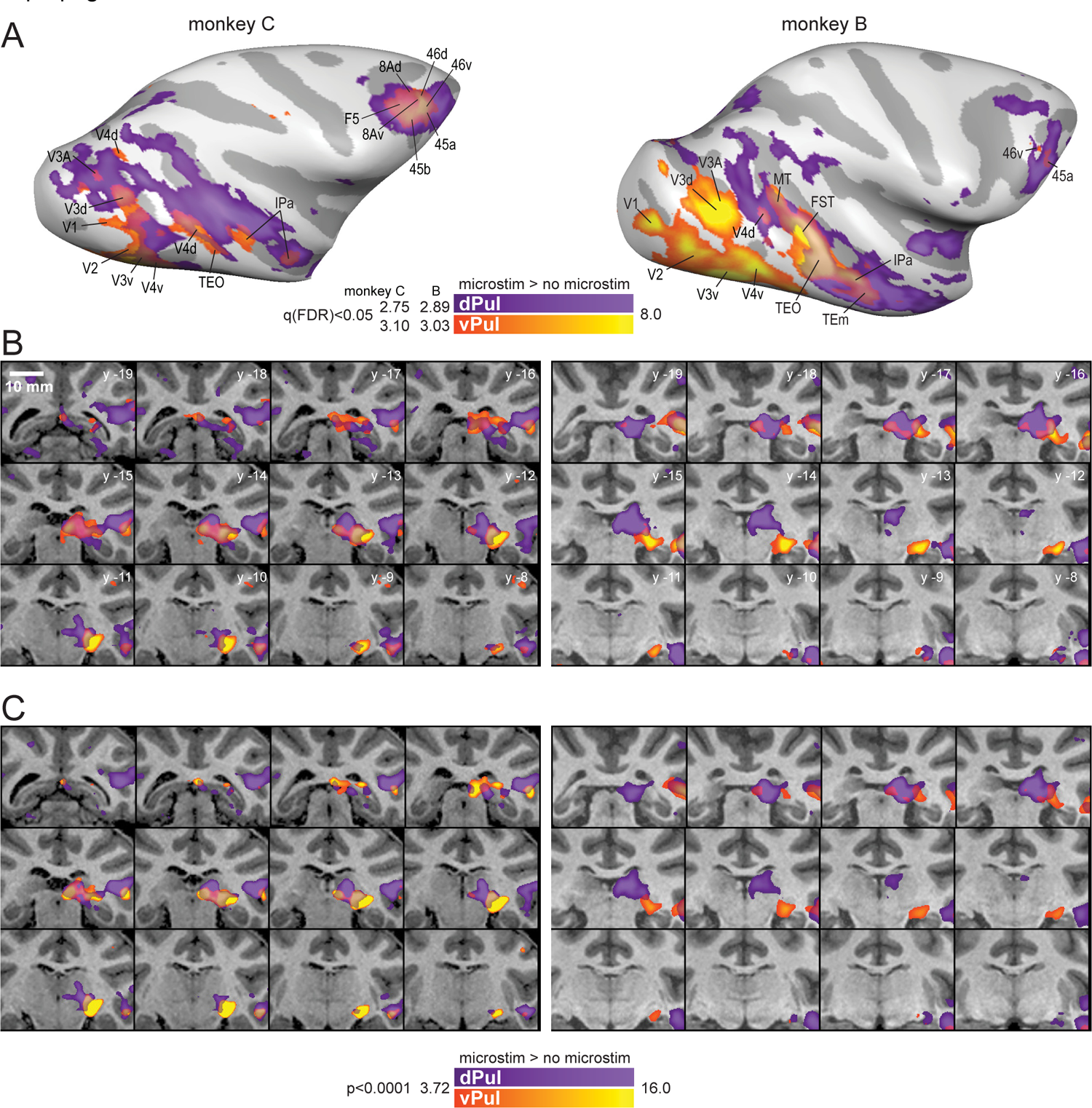
Comparison of dPul and vPul stimulation effects. **(A)** Activation maps on the inflated brain surface, lateral view. Ventral pulvinar-elicited activated regions are labeled. **(B)** Zoomed in sections through the thalamus (1 mm spacing). **(C)** Same as (B), but with a higher statistic threshold (p<0.0001 uncorrected). L – left, R – right, S – superior, I – inferior, y - distance from AC-PC origin along the anterior/posterior plane in mm.

Since there was a partial overlap between cortical connectivity of the dPul and the vPul, we compared the stimulation effect strength (see section 3.3.1) in the overlapping regions (**Figure S22**). In monkey C, only V3v clearly showed stronger activity due to vPul stimulation. In monkey B, occipital ROIs as well as MT, MST, and FST showed a stronger activation by vPul. In both monkeys, IPa, PGa and TEO/TE showed similar or stronger dPul stimulation effects. Within the limits of interpretability due to the difference of stimulation current strength, and possible gradations of anatomical projection density, it cannot be ruled out that some early visual areas were activated via adjacent vPul (or SC) co-activation during dPul stimulation. Yet, a weaker direct connectivity or polysynaptic transmission via cortex is also plausible. Additional control experiments comparing a lower current strength (100 μA), and hence more local dPul stimulation that barely encroached on the vPul (**Figure S3**), supported an effective connectivity gradient from ventral occipital and STS areas (vPul stimulation) to the fundus and the dorsal STS (dPul stimulation; see **Figure S23**). This datatset also confirmed dPul-elicited activation of 7a, MT, FST, PGa and IPa. Thus, even when the stimulation sites are only 3-4 mm apart, the distal activations show distinct patterns.

One unexpected finding was the activation of the frontal cortex following stimulation of vPul, extensively in areas 8Ad/v, 8Bs, 45a/b and 46d/v in monkey C, and in areas 45a and 46v in monkey B (**Figure S24**). To the best of our knowledge, direct anatomical connections from vPul to these regions have not been reported, further suggesting polysynaptic propagation of microstimulation-elicited fMRI activity, e.g. from extrastriate cortex. Likewise, the activation of SC both after dPul and vPul stimulation can be polysynaptic via thalamo-cortical-collicular pathways or via antidromic propagation.

### 3.3 Effects of microstimulation on BOLD responses during different task conditions

Up to this point, we considered stimulation-elicited activations pooling across all task conditions (except in control analyses presented in **Figures S17** and **S20** for the fixation only condition). We next used such combined microstimulation > no microstimulation maps and CHARM/SARM atlases to define a set of regions of interests (ROIs), individually for each dataset and each monkey (**Table S3**). We performed event-related averaging of single trial timecourses extracted from those ROIs (Methods), to compare activations in the two hemispheres and to explore task-specific effects.

#### 3.3.1 Comparison of stimulation effects between hemispheres

As expected, stimulation effects were stronger in the stimulated hemisphere compared to the opposite hemisphere, based on both the number of significantly activated ROIs (Wilcoxon matched-pairs signed-ranks test, p=0.002; **Figure S25**, **Table 2**) and on the statistical comparison of the stimulation effect strength, derived from single trial response amplitudes averaged across trials from all task conditions. Combining across ROIs, 8/10 datasets showed higher stimulation-elicited enhancement in the stimulated hemisphere (two-sided t-test, p<0.001; dPul(a) datasets in both monkeys did not show this effect). Weaker stimulation effect in the opposite hemisphere was also observed for all 10 datasets using atlas ROIs (all p<0.001, except monkey C, LIPd-a p=0.004, vPul p = 0.015).

Most ROIs that were activated in the non-stimulated hemisphere had a homotopic counterpart in the stimulated hemisphere, suggesting a transcallosal interhemispheric connectivity route (**Table 2**). This interpretation is further supported by the significant correlation between the activation strength profiles in the left and the right hemisphere, across ROIs activated in either hemisphere (significant in 8/10 datasets, besides the two dorsal pulvinar datasets in monkey B where the correlation analysis was limited by only 9 or 11 left hemisphere ROIs; **Table 2**). Furthermore, the comparable interhemispheric correlation was present in all datasets when calculated across all atlas ROIs, irrespective of a significant activation by the stimulation (**Table 2**).

#### 3.3.2 Task-specific stimulation effects

Example timecourses separated by task conditions are shown in **Figure 7**. The effect of microstimulation typically started 2 to 3 s after the stimulation onset (the first of 10 stimulation trains) and continued throughout the memory delay period or the corresponding fixation period. The stimulation modulated the task-related activity, which often showed a contralateral spatial selectivity of transient cue and sustained delay activity in control trials (i.e. stronger responses in the contraversive memory task condition in the stimulated hemisphere, and stronger responses in the ipsiversive condition in the opposite hemisphere). In many ROIs, the stimulation tended to enhance the initial task-related activity additively by a similar amount; in some other ROIs, the stimulation effect seemed to vary between task conditions. To quantify these effects, single trial response amplitudes were extracted using an analysis window of 2 – 9 s after the onset of stimulation, and were submitted to a two-way ANOVA with factors task and stimulation, in each ROI. **Figure S26** and **Figure S27** show the ANOVA results for pulvinar and LIPd sites, respectively. Since these ROIs were selected on the basis of the positive stimulation effect in the statistical contrast map, main effect of the stimulation in most ROIs merely confirmed trial epoch-specific enhancement derived directly from BOLD signal timecourses. Here, the important question was whether the stimulation effect interacted with the task the animal was performing. At the level of single ROIs, however, there was very little evidence for task-dependence of the stimulation effect. Even when the stimulation effect varied considerably between task conditions based on visual inspection of the timecourses, the variability of single trial response amplitudes was too large to result in a statistically significant task × stimulation interaction. Consequently, only few ROIs showed such interaction effect.

**Figure 7.**
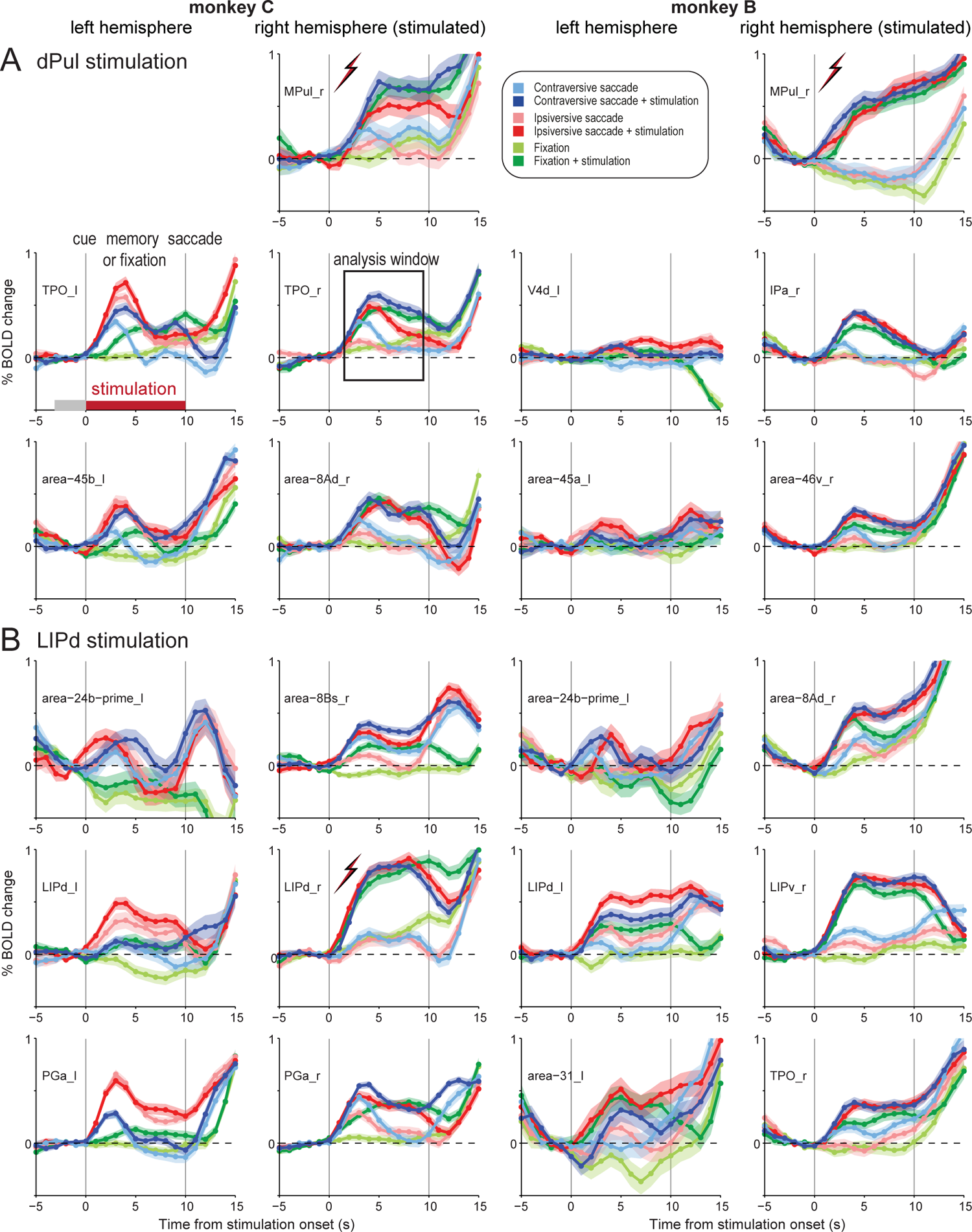
Example trial event-related average timecourses. Trial averages and standard error of %BOLD signal change relative to the last 4 s of the initial fixation are plotted for the last 5 s of the initial fixation, 10 s memory / fixation period coinciding with the stimulation epoch, and 5 s of saccadic response / fixation for the three task conditions, in control and stimulation trials. Timecourses were aligned to the onset of the stimulation that started immediately after the offset of 200 ms peripheral saccade cue, or an end of initial fixation period. The timing of trial events and the stimulation is indicated on the first column, second row panel (gray – fixation baseline, dark red – stimulation period). The analysis window (2 – 9 s) is indicated in the second row, second column panel. **(A)** dPul stimulation. **(B)** LIPd stimulation.

We next investigated average stimulation effects for each task condition, *across ROIs*. **Figure 8** shows the average magnitude of the stimulation effect in all five stimulation sites for each task and hemisphere, and **Table 3** lists the corresponding statistics. We used repeated measures ANOVA across ROIs on the mean BOLD response amplitudes during stimulation and the corresponding periods in trials without stimulation to assess the main effect of task (contraversive saccades, ipsiversive saccades, fixation), the main effect of stimulation (no stimulation, stimulation), and the interaction between the task and the stimulation. Post-hoc paired t-tests contrasted stimulation effects between task conditions. In the right (stimulated) hemisphere both monkeys showed a significant main effect of task (and trivially, a main effect of stimulation) for all stimulation sites. More importantly, a significant interaction between the stimulation effect and the task was found in the dPul and LIPd datasets (except in LIPd-a in monkey C). In the left (non-stimulated) hemisphere the interaction was significant for dPul and LIPd stimulation sites in both monkeys. The post-hoc paired t-tests between task conditions showed many significant differences but the direction of the effect was not consistent between monkeys. In keeping with our initial prediction, in monkey C the contraversive saccade condition showed the strongest stimulation effect compared to fixation and ipsiversive saccade trials, while in monkey B, the fixation or ipsiversive task condition showed the strongest stimulation effect.

**Figure 8.**
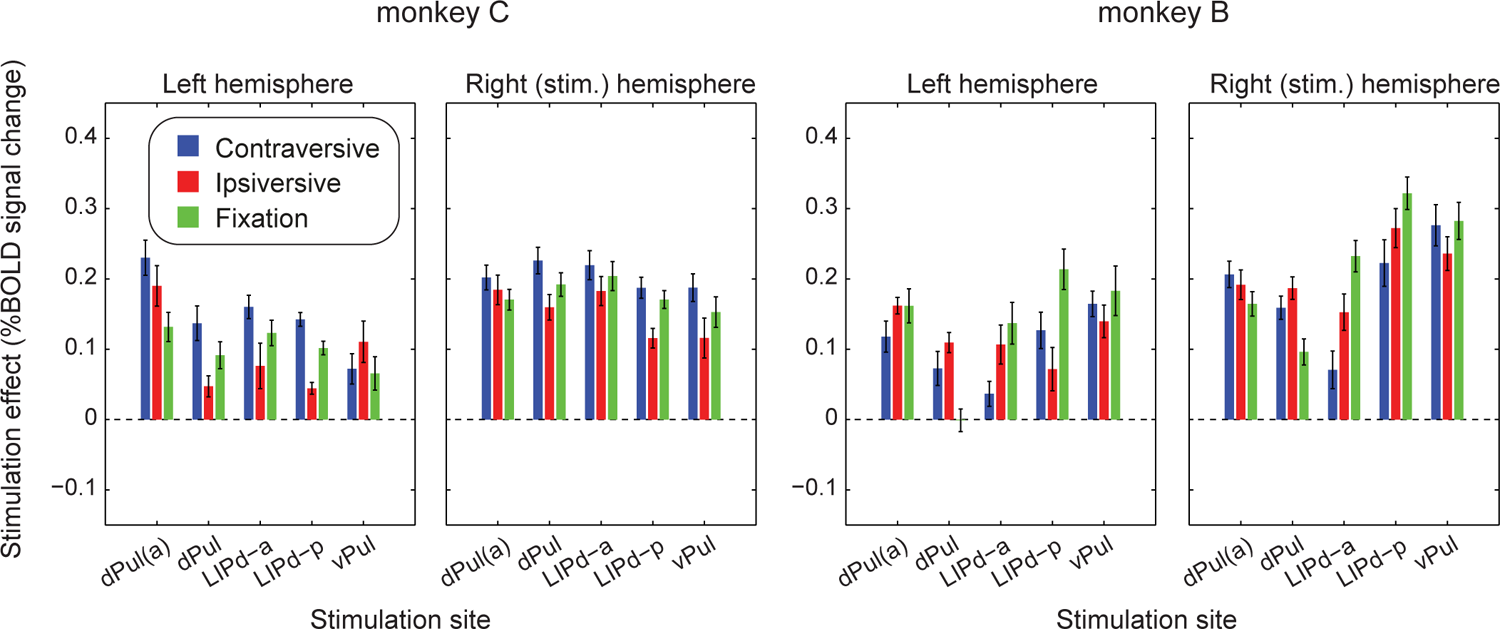
Effects of stimulation in dPul(a), dPul, LIPd-a, LIPd-p and vPul on trial-averaged BOLD responses. Means and standard errors across ROIs for stimulation effects calculated as the difference in %BOLD signal change between trials with and without stimulation, for the contraversive memory saccade task (blue bars), the ipsiversive memory saccade task (red bars), and the fixation task (green bars). See **Table 2** for the corresponding statistics.

### 3.4 Relationship between spatial selectivity and the magnitude of stimulation effects on BOLD responses

To further explore the dependence of stimulation effects on the task, and to understand the discrepancies between the two monkeys regarding the relative strength and direction of such effects, we next assessed whether the magnitude of stimulation effects on task-related activity depended on the initial spatial selectivity of the activated ROIs. To this end, we correlated contraversive selectivity index (CSI) derived from the control trials with the stimulation effects across all ROIs, separated by task condition. Note that for the ROIs in the stimulated right hemisphere the contraversive selectivity corresponds to the contralateral selectivity, while for the left hemisphere the contraversive selectivity is the ipsilateral selectivity. A significant correlation would suggest that the stimulation effect depends on initial spatial tuning, within a given task condition. Furthermore, if there is a difference in the above relationships between task conditions (i.e. different regression slopes), this would suggest a degree of task-dependence. To assess whether the relationship between the stimulation effect and CSI differed between task conditions, we grouped ROIs into two categories: with contraversive tuning (CSI > 0) or ipsiversive tuning (CSI < 0), and tested whether the magnitude of stimulation effects differed between tasks within each group using a two-way mixed ANOVA (see Methods). Further post-hoc t-tests were used to assess pairwise differences between tasks for each of the two ROI groups.

The resulting correlations for the contraversive and ipsiversive memory saccade task conditions are shown as scatter plots with least-square fits (**Figure 9**), and the corresponding statistics are shown in **Table 4**. The plots for all pair-wise task comparisons are shown in **Figures S28** and **S29**. To summarize the dependency of stimulation effects on spatial selectivity, we consistently found positive correlations across ROIs between contraversive selectivity and stimulation-elicited enhancement in the ipsiversive memory saccade task (significant in 8 out of 10 datasets). In other words, stronger contraversive tuning of the respective ROIs was associated with a stronger microstimulation-elicited enhancement of BOLD responses to visual cues and motor preparation directed towards the ipsiversive hemifield. Conversely, significant correlations for the contraversive task were negative, except for the LIPd-a site in monkey B. In the fixation task the correlations were less consistent: when significant, positive correlations were found in monkey C but a negative correlation in monkey B (**Figure S28**, **S29**).

**Figure 9.**
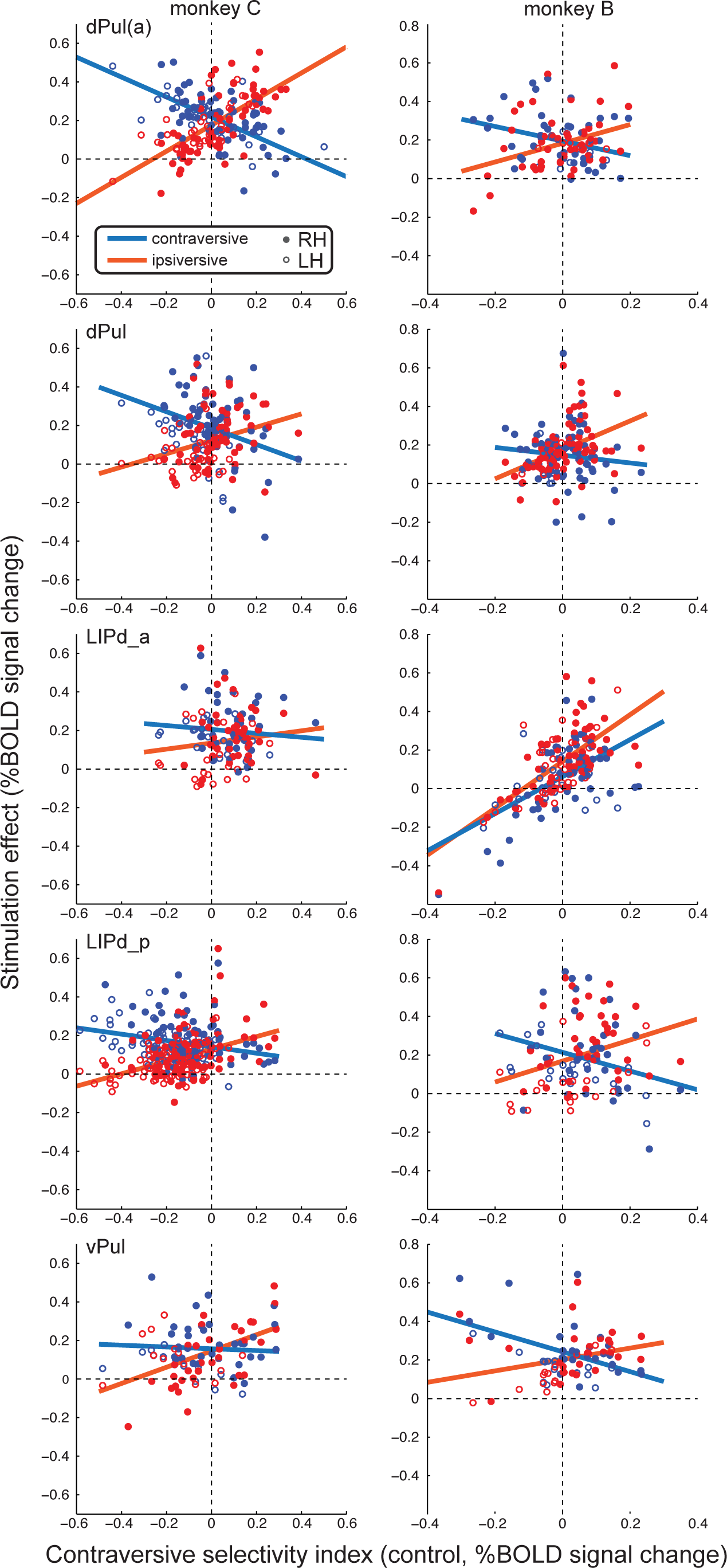
Microstimulation effect dependence on contraversive spatial selectivity and task. For each ROI, the task-specific effect of stimulation is plotted against ROI’s initial (in trials without stimulation) contraversive selectivity index for contraversive (blue) and ipsiversive (red) memory saccade task conditions. Each dot represents one ROI (filled dots – right hemisphere, empty dots – left hemisphere); solid lines show linear fits of stimulation effects across ROIs. See **Table 4** for the corresponding statistics, and **Figure S28** and **S29** for data on all pairwise task condition comparisons.

**Table 4.**
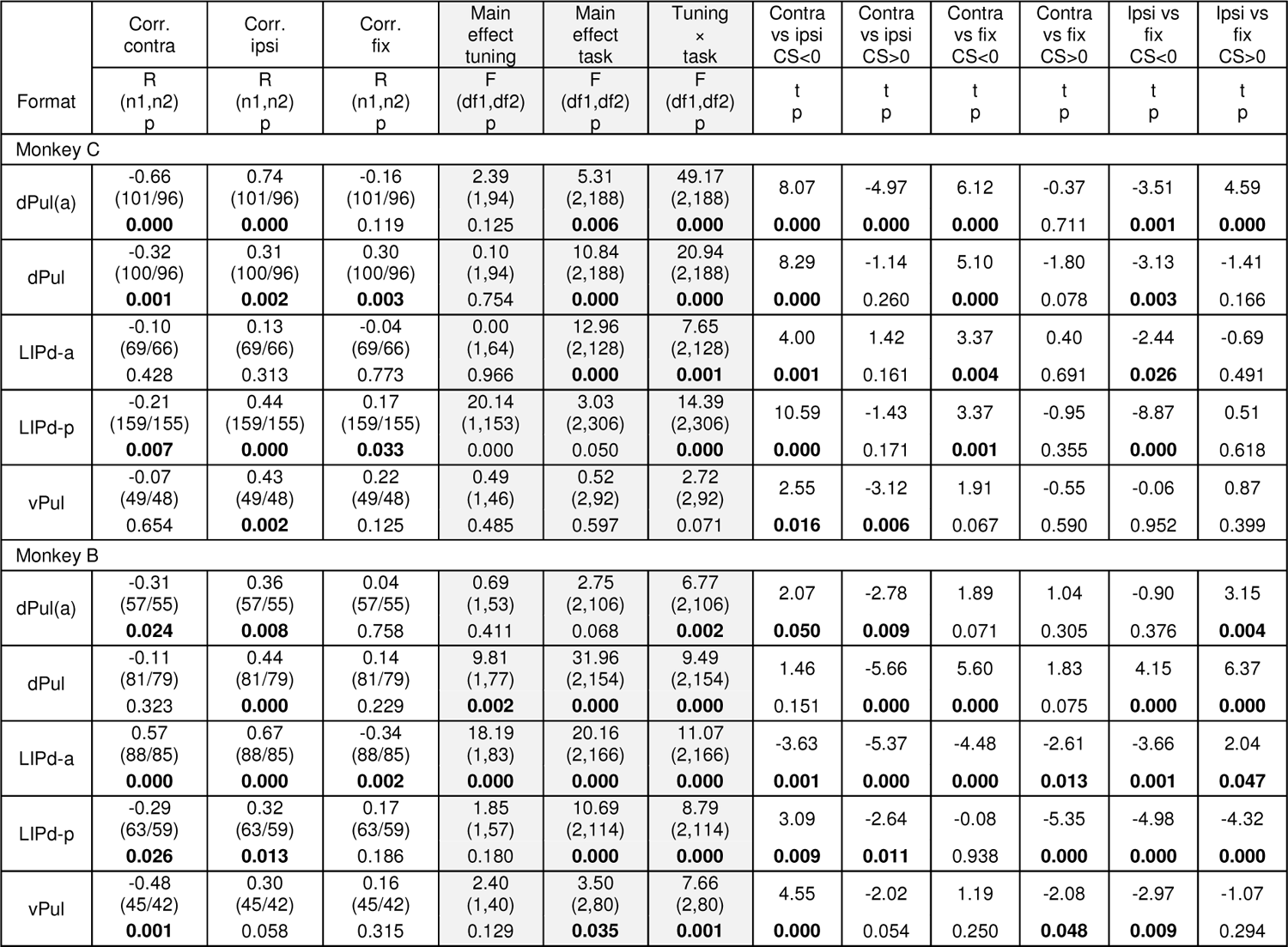
Relationship between stimulation effect, spatial selectivity and task condition. Significant p-values (p<0.05) for ANOVAs and t-tests are in bold font. For this analysis, ROIs from both hemispheres were combined. R – Pearson’s linear correlation coefficient for the three different task conditions, N – number of ROIs before / after exclusion of outliers (see Methods), p – p-value, F – F-statistics (two-way mixed ANOVA), t – t-value for the t-test on the difference of stimulation effect strength between the three task conditions (contra, ipsi, fix).

| Format | Corr. contra | Corr. ipsi | Corr. fix | Main effect tuning | Main effect task | Tuning × task | Contra vs ipsi CS<0 | Contra vs ipsi CS>0 | Contra vs fix CS<0 | Contra vs fix CS>0 | Ipsi vs fix CS<0 | Ipsi vs fix CS>0 |
| --- | --- | --- | --- | --- | --- | --- | --- | --- | --- | --- | --- | --- |
|  | R (n1,n2)<br>p | R (n1,n2)<br>p | R (n1,n2)<br>p | F (df1,df2)<br>p | F (df1,df2)<br>p | F (df1,df2)<br>p | t<br>p | t<br>p | t<br>p | t<br>p | t<br>p | t<br>p |
| Monkey C |  |  |  |  |  |  |  |  |  |  |  |  |
| dPul(a) | -0.66<br>(101/96)<br>**0.000** | 0.74<br>(101/96)<br>**0.000** | -0.16<br>(101/96)<br>0.119 | 2.39<br>(1,94)<br>0.125 | 5.31<br>(2,188)<br>**0.006** | 49.17<br>(2,188)<br>**0.000** | 8.07<br>**0.000** | -4.97<br>**0.000** | 6.12<br>**0.000** | -0.37<br>0.711 | -3.51<br>**0.001** | 4.59<br>**0.000** |
| dPul | -0.32<br>(100/96)<br>**0.001** | 0.31<br>(100/96)<br>**0.002** | 0.30<br>(100/96)<br>**0.003** | 0.10<br>(1,94)<br>0.754 | 10.84<br>(2,188)<br>**0.000** | 20.94<br>(2,188)<br>**0.000** | 8.29<br>**0.000** | -1.14<br>0.260 | 5.10<br>**0.000** | -1.80<br>0.078 | -3.13<br>**0.003** | -1.41<br>0.166 |
| LIPd-a | -0.10<br>(69/66)<br>0.428 | 0.13<br>(69/66)<br>0.313 | -0.04<br>(69/66)<br>0.773 | 0.00<br>(1,64)<br>0.966 | 12.96<br>(2,128)<br>**0.000** | 7.65<br>(2,128)<br>**0.001** | 4.00<br>**0.001** | 1.42<br>0.161 | 3.37<br>**0.004** | 0.40<br>0.691 | -2.44<br>**0.026** | -0.69<br>0.491 |
| LIPd-p | -0.21<br>(159/155)<br>**0.007** | 0.44<br>(159/155)<br>**0.000** | 0.17<br>(159/155)<br>**0.033** | 20.14<br>(1,153)<br>0.000 | 3.03<br>(2,306)<br>0.050 | 14.39<br>(2,306)<br>**0.000** | 10.59<br>**0.000** | -1.43<br>0.171 | 3.37<br>**0.001** | -0.95<br>0.355 | -8.87<br>**0.000** | 0.51<br>0.618 |
| vPul | -0.07<br>(49/48)<br>0.654 | 0.43<br>(49/48)<br>**0.002** | 0.22<br>(49/48)<br>0.125 | 0.49<br>(1,46)<br>0.485 | 0.52<br>(2,92)<br>0.597 | 2.72<br>(2,92)<br>0.071 | 2.55<br>**0.016** | -3.12<br>**0.006** | 1.91<br>0.067 | -0.55<br>0.590 | -0.06<br>0.952 | 0.87<br>0.399 |
| Monkey B |  |  |  |  |  |  |  |  |  |  |  |  |
| dPul(a) | -0.31<br>(57/55)<br>**0.024** | 0.36<br>(57/55)<br>**0.008** | 0.04<br>(57/55)<br>0.758 | 0.69<br>(1,53)<br>0.411 | 2.75<br>(2,106)<br>0.068 | 6.77<br>(2,106)<br>**0.002** | 2.07<br>**0.050** | -2.78<br>**0.009** | 1.89<br>0.071 | 1.04<br>0.305 | -0.90<br>0.376 | 3.15<br>**0.004** |
| dPul | -0.11<br>(81/79)<br>0.323 | 0.44<br>(81/79)<br>**0.000** | 0.14<br>(81/79)<br>0.229 | 9.81<br>(1,77)<br>**0.002** | 31.96<br>(2,154)<br>**0.000** | 9.49<br>(2,154)<br>**0.000** | 1.46<br>0.151 | -5.66<br>**0.000** | 5.60<br>**0.000** | 1.83<br>0.075 | 4.15<br>**0.000** | 6.37<br>**0.000** |
| LIPd-a | 0.57<br>(88/85)<br>**0.000** | 0.67<br>(88/85)<br>**0.000** | -0.34<br>(88/85)<br>**0.002** | 18.19<br>(1,83)<br>**0.000** | 20.16<br>(2,166)<br>**0.000** | 11.07<br>(2,166)<br>**0.000** | -3.63<br>**0.001** | -5.37<br>**0.000** | -4.48<br>**0.000** | -2.61<br>**0.013** | -3.66<br>**0.001** | 2.04<br>**0.047** |
| LIPd-p | -0.29<br>(63/59)<br>**0.026** | 0.32<br>(63/59)<br>**0.013** | 0.17<br>(63/59)<br>0.186 | 1.85<br>(1,57)<br>0.180 | 10.69<br>(2,114)<br>**0.000** | 8.79<br>(2,114)<br>**0.000** | 3.09<br>**0.009** | -2.64<br>**0.011** | -0.08<br>0.938 | -5.35<br>**0.000** | -4.98<br>**0.000** | -4.32<br>**0.000** |
| vPul | -0.48<br>(45/42)<br>**0.001** | 0.30<br>(45/42)<br>0.058 | 0.16<br>(45/42)<br>0.315 | 2.40<br>(1,40)<br>0.129 | 3.50<br>(2,80)<br>**0.035** | 7.66<br>(2,80)<br>**0.001** | 4.55<br>**0.000** | -2.02<br>0.054 | 1.19<br>0.250 | -2.08<br>**0.048** | -2.97<br>**0.009** | -1.07<br>0.294 |

Importantly, we consistently found interactions between the main effect of spatial tuning and the main effect of task condition on the stimulation effect magnitude in 9 out of 10 sites (the only exception being a trend in the vPul site in monkey C (p=0.071)) (**Table 4**). Similar interactions were observed when ROIs were separated by hemisphere (**Table S5**) and in atlas ROIs (**Table S6**). This indicates that across ROIs, the stimulation effects differed between ROIs with contraversive tuning and ROIs with ipsiversive tuning, depending on task condition. Specifically, in monkey C in *ROIs with ipsiversive tuning* (CS<0), both pulvinar and LIP stimulation led to the strongest enhancement in the contraversive memory task and weakest effects in the ipsiversive memory task. Monkey B showed a similar pattern with one exception: while in the pulvinar and LIPd-p datasets, the stimulation effect was strongest for the contraversive task (significant in dPul(a), LIPd-p and vPul), the LIPd-a dataset showed the strongest effect in the fixation task. Conversely, in both monkeys *contraversively-tuned ROIs* (CS>0) consistently showed a stronger stimulation effect in the ipsiversive task compared to the contraversive task (stimulation effect contra<ipsi in 9 out of 10 datasets, significant in 6 out of 10 datasets, **Table 4**) and fixation task (stimulation effect contra<fix in 7 out of 10 datasets, significant in 3 datasets in monkey B, **Table 4**). Taken together, our results indicate *inverse (“X-shaped”) pattern* of task-dependent effects for ROIs with contraversive tuning and ROIs with ipsiversive tuning. Furthermore, variations in the relative frequency of contraversively- and ipsiversively-tuned ROIs, as well as the slope and the intercept of the observed stimulation effect fits, could explain differences between average task-specific stimulation effects across monkeys and datasets (cf. **Table 3** and **Figure 8**).

### 3.5 Modeling effects of microstimulation on BOLD responses

Finally, we asked which model accounts best for the observed stimulation effects on BOLD signal amplitudes. The two basic models represent an additive or a multiplicative mechanism, i.e. an enhancement of the initial (no stimulation) response by a constant added term, or a multiplication by a certain factor. Additionally, since we observed a dependence of the microstimulation effect on the initial tuning of a region (ROIs with a negative contraversive selectivity showed more enhancement in contraversive memory saccade trials, while ROIs with a positive contraversive selectivity showed more enhancement in ipsiversive memory saccade trials), we also tested a variant of an additive model where the additive factor is scaled down by the initial response amplitude. Because of the stronger stimulation effect in the stimulated hemisphere, the fits were done separately for each hemisphere. The resulting adjusted R-squared values signifying the goodness of fit are shown in **Figure 10A**. In both monkeys, across datasets and the two hemispheres, the best model was the additive scaled model, followed by the additive and then the multiplicative model (all pairwise comparisons p<0.001, **Table S7**). This further confirms that while most individual ROIs did not show a significant task-dependence of the microstimulation effect across trials – as would otherwise be expected from a multiplicative model or an additive model with a strong difference between the additive factors across task conditions – the spatial selectivity, reflected in the initial response amplitude, plays a significant role in determining the strength of stimulation effect. The fitted additive coefficients matched well the actual magnitude of stimulation effects on BOLD responses (compare **Figure 10B** with **Figure 8**). Furthermore, a simple qualitative simulation implementing the additive scaled effect could reproduce the “X-shaped” pattern of interaction between the task condition and the stimulation effect strength in relationship to contraversive spatial selectivity (**Figure S30**).

**Figure 10.**
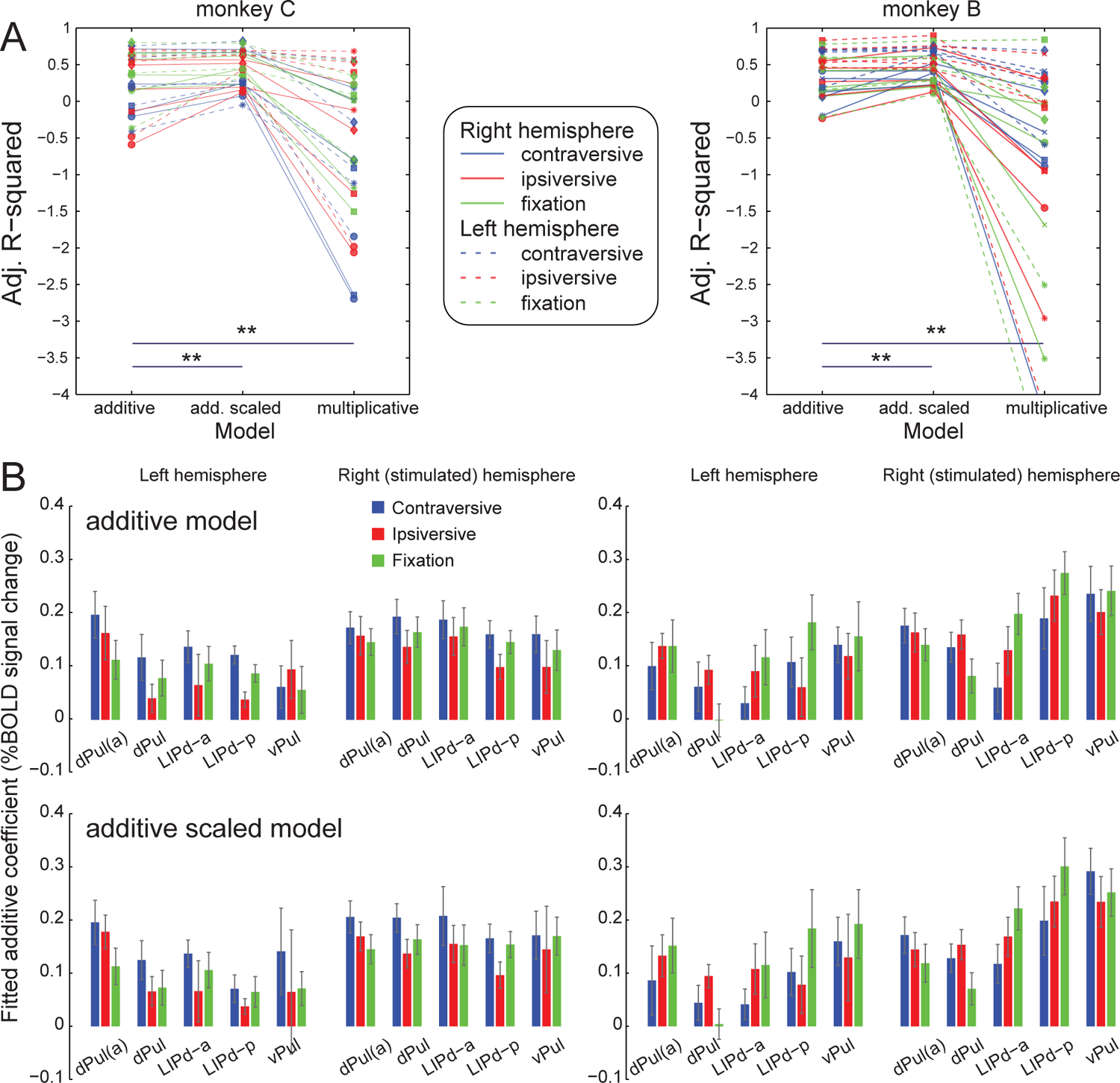
Stimulation effects are best approximated by additive scaled model. **(A)** Goodness of fit for modeling effects of microstimulation in different sites: dPul(a) (dots), dPul (squares), LIPd-a (diamonds), LIPd-p (crosses), vPul (stars). Adjusted R-squared values are shown separately for the right, stimulated hemisphere (solid lines) and the left hemisphere (dashed lines). **(B)** Fitted additive coefficients with 95% confidence intervals for the two additive models.

## 4 Discussion

Unilateral microstimulation in both dPul and LIPd caused widespread fMRI BOLD activity enhancement, predominantly in prefrontal and parietal cortex, the dorsal bank and fundus of superior temporal sulcus (STS) and in extrastriate visual cortex, in the stimulated hemisphere. Multiple regions along the dorsal bank of the STS stood out as a prominent shared “node” of es-fMRI connectivity for all dPul and LIPd stimulation sites. Weaker es-fMRI activity was also present in the opposite hemisphere, suggesting transcallosal and polysynaptic propagation. The comparison between the more anterior and posterior LIPd stimulation sites showed extensive activation overlap along the intraparietal sulcus and in STS, but partially differing activation patterns in frontal and occipital regions. The comparison of dPul and vPul stimulation also showed partly distinct effective connectivity, mainly consistent with known anatomical projections. For both pulvinar and parietal es-fMRI we observed task-specific correlations between the initial spatial selectivity and the magnitude of stimulation effects, across ROIs. The effects of microstimulation on BOLD signal amplitudes were best fitted by an additive model with an additive factor scaled down by response amplitude in control trials.

### 4.1 Pulvinar effective connectivity and comparison with anatomy

Microstimulation in the dPul elicited BOLD activity that was largely consistent with the reciprocal connections with visuomotor and polymodal cortical regions known from anatomical tracer studies in monkeys (Kaas and Lyon, 2007; Shipp, 2003). Thus, the extensive activity along the superior temporal sulcus (e.g. MT, MST, FST, TPO, PGa, TE, TEO), including anterior STS (IPa, TAa) (Yeterian and Pandya, 1991, 1989), posterior parietal cortex (e.g. LOP, LIP, MIP, VIP, 7a) (Asanuma et al., 1985; Cappe et al., 2009, 2007), frontal cortex (e.g. 8A/B, 45, 46), insula (Romanski et al., 1997), cingulate and premotor cortices (Baleydier and Mauguiere, 1985) was expected. The effective connectivity of dPul is conceptually in line with a number of recording and lesion studies in monkeys and humans that highlight its critical contribution to higher-order visual, oculomotor and skeletomotor functions involving spatial attention (Arend et al., 2008a; Fiebelkorn et al., 2019; Karnath et al., 2002; Petersen et al., 1987; Rafal et al., 2004; Van der Stigchel et al., 2010), decision making and action selection (Dominguez-Vargas et al., 2017; Komura et al., 2013; M. Wilke et al., 2010; Wilke et al., 2013) and sensorimotor transformations for visually-guided reaching and grasping (Mundinano et al., 2018; Schneider et al., 2020; M. Wilke et al., 2010; Wilke et al., 2018, 2017). The elicited activations are also largely consistent with human studies that investigated intracerebral-evoked responses to dorso-medial pulvinar stimulation in epileptic patients (Rosenberg et al., 2009) or used fMRI functional connectivity analyses (Arcaro et al., 2018). It is interesting however that whereas our stimulation results as well as the stimulation in humans highlight STS/temporal regions in addition to the fronto-parietal and cingulate connectivity (Rosenberg et al., 2009), functional connectivity work emphasizes strong coupling of dPul to fronto-parietal and cingulate cortex (Arcaro et al., 2018).

While the activation patterns elicited by the dorsal pulvinar stimulation were largely consistent in the two animals and the two stimulation sites, there were also some differences. For instance, in the stimulated hemisphere monkey B exhibited more extensive activity in inferotemporal cortex (TEO, TEm, TEa) and insula, while monkey C showed more extrastriate and frontal activation. Given the diverse connectivity of pulvinar subportions we cannot exclude that some differences in stimulated pulvinar populations account for this variability. For example, paired injection studies in the inferotemporal cortex (TE/TEO) and inferior parietal cortex (POa/PGc, corresponding to LIP and area 7) in macaques suggest that while both regions receive inputs from the medial pulvinar, the projection neurons to each of the areas are largely separated (Baizer et al., 1993; Baleydier and Morel, 1992): temporal projections originate from the most posterior medial pulvinar portion (caudal pole) while parietal projections are located more anteriorly within the dorsal pulvinar. In agreement with this, there was a tendency of more activity in the extrastriate and inferotemporal cortex with the more posterior dorsal stimulation site *in each monkey*, whereas parietal and frontal cortices (except the frontal areas in monkey B) were more strongly activated by slightly more anterior (∼ 1 mm) stimulation. However, while overall the stimulation was ∼ 2 mm more anterior in monkey B than in monkey C, monkey B showed more extensive ventral temporal and less extensive frontal activation, contrary to the expectations. Thus, the small anterior-posterior offsets in stimulation locations between monkeys cannot fully account for the observed differences. It is important to note that our dPul stimulation sites were mostly in the medial pulvinar but encroached on the border between the medial and the lateral pulvinar. In anatomical tracer studies, this region (referred to as “PMl” for “Pulvinar Medial lateral”) showed an overlap of temporal and parietal connectivity (Baleydier and Morel, 1992; Gutierrez et al., 2000).

In contrast to dPul, stimulation in vPul elicited stronger activation in extrastriate cortex and ventral STS, but weaker or no activation in parietal cortex and dorsal STS, as expected from the monkey anatomical literature (Kaas and Lyon, 2007) and functional connectivity studies in humans (Arcaro et al., 2018, 2015). We point out however that the partial overlap between the connectivity of dPul and vPul in extrastriate cortex and STS that we observed is present even in tracer studies, suggesting only a gradual partitioning to classical dorsal/ventral visual streams (Gutierrez et al., 2000).

Importantly, some activations must have been of antidromic and/or polysynaptic origin. For example, the superior colliculus (SC) was activated, and although it projects to both dPul and vPul, it does not receive direct projections from the pulvinar. Theoretically, this activation could due to antidromic propagation or second-order activation through cortex (Baldwin and Bourne, 2017; Bender and Butter, 1987; Benevento and Standage, 1983; Harting et al., 1980; Murayama et al., 2011). Likewise, V1/V2 were activated by microstimulation in the dPul, but retrograde tracer studies typically do not report corresponding projections from the dorsal pulvinar (Adams et al., 2000). We applied a relatively high current strength (250 µA) (as typically used in es-fMRI studies (Matsui et al., 2011; Moeller et al., 2008; Tolias et al., 2005)). Therefore, we cannot rule out that evoked activity in early visual cortex and MT that is typically associated with SC and vPul connectivity might also be due to co-activation of these adjacent regions (and conversely, prefrontal activation by vPul stimulation could have resulted from the co-activation of dPul). This possibility is reinforced by the absence of V1 and very weak V2 activity in the control experiment where dPul was stimulated with lower (100 µA) current, which elicited very minor ventral pulvinar co-activation. On the other hand, the extent of established direct dPul cortical connectivity also decreased dramatically with lower current strength. Notably, MT and FST were still weakly activated with the lower current, in line with their known anatomical connectivity to parts of dPul (Gutierrez et al., 2000). Furthermore, differences of distal activation patterns due to vPul or dPul stimulation even with higher currents suggest that notwithstanding the apparent partial overlap of the local activation spread, typically more extensive when measured with fMRI than with other techniques (Klink et al., 2021; Tehovnik et al., 2006), es-fMRI connectivity is strongly site-specific. Such specificity was also reported for the behavioral effects of dPul vs. vPul stimulation with comparable currents (Dominguez-Vargas et al., 2017). Therefore, we favor the view that the intra-hemispheric activations not expected from the monosynaptic connectivity are due to polysynaptic propagation.

Since the pulvinar is known to project exclusively to ipsilateral cortical regions, the strongest evidence for polysynaptic propagation of es-fMRI effects via inter-hemispheric transcallosal connections is activity in cortical regions in the hemisphere opposite to the stimulation. Similar evidence for polysynaptic spread has been reported in previous es-fMRI studies with stimulation of somatosensory cortex, FEF, and LGN (Matsui et al., 2012, 2011; Murayama et al., 2011).

The activation in the cerebellum was not expected from anatomical studies in primates. Nonetheless, in cats, several anatomical tracer studies reported projections from the cerebellum to the contralateral LP nucleus, a structure partially homologous to the primate pulvinar (Itoh and Mizuno, 1979; Rodrigo-Angulo and Reinoso-Suarez, 1984), and a previous macaque es-fMRI study in the deep nuclei of the cerebellum found strong BOLD enhancement in the dorsal pulvinar (Sultan et al., 2012). Additionally, a patient with a selective (bilateral) lesion in the medial pulvinar exhibited a combination of parietal and cerebellar deficits (Wilke et al., 2018). Thus, the functional significance of pulvino-cerebellar pathway in monkeys/humans deserves further investigation.

### 4.2 LIPd effective connectivity and comparison with anatomy

The LIPd stimulation-elicited distal activation patterns were largely predicted by anatomical tracer studies (Blatt et al., 1990): both banks and the fundus of STS, regions in the posterior parietal cortex (PIP, LOP, MIP, 7a, area 5), medial parietal and posterior cingulate regions, prefrontal areas 45a/b, 8A/B and 46d/v, striate and extrastriate visual cortex, and cerebellum. Since anatomical tracer, electrophysiological and functional fMRI studies indicate that LIP is a heterogeneous area, even beyond the well-established ventral/dorsal subdivision (Blatt et al., 1990; Chen et al., 2016; Lewis and Van Essen, 2000; Liu et al., 2010; Patel et al., 2010), we had stimulated two different portions along the dorsal LIP. The anterior stimulation site LIPd-a was similar to a previous es-fMRI study in monkeys that stimulated an anterior portion of the LIP as control for stimulation in the grasp-related area AIP (Premereur et al., 2015). This study reported LIP-elicited activation in MIP, FST, V4, V3A and to a lesser extent in FEF. In comparison, the activation patterns in our study were more extensive and included more brain regions (e.g. cerebellum) and notably also homotopic visuomotor regions in the non-stimulated hemisphere. Generally, it is difficult to compare the stimulation results across studies due to many methodological variations. Since current strength was comparable and both studies used awake animals, this might is due to differences in other stimulation parameters such as longer stimulation durations in our study. While the macaque literature already makes an anatomical and functional division between ventral LIPd and LIPv subdivision (Lewis and Van Essen, 2000; Liu et al., 2010), the anterior-posterior axis is less well described. While the stimulation of the more posterior vs. anterior sites in LIPd yielded largely overlapping patterns in parietal cortex and STS, there was more extensive activation of PMd, FEF (area 8A/B) and PFC (area 46d/v) with anterior LIPd-a stimulation. This anterior-posterior gradient should be followed up with systematic studies of electrophysiological properties (Premereur et al., 2015).

Unlike the dPul stimulation effects in the opposite hemisphere, the activation of heterotopic regions in the opposite parietal and parieto-temporal cortex by the LIP stimulation cannot be taken as a definitive proof of polysynaptic propagation, due to a presence of heterotopic transcallosal connections between these regions (Hedreen and Yin, 1981). It remains to be investigated if these connections are dense enough to support monosynaptic transcallosal activation, as an alternative to the polysynaptic transmission via homotopic parietal regions. Small activations in the premotor or cingulate cortex that we observed are unlikely to be mediated by the direct transcallosal connectivity (Lanz et al., 2017).

The lack of consistent and convincing activation of dPul and superior colliculus by LIP stimulation was surprising. The anatomical and functional connections between LIP and those subcortical regions are well established (Andersen et al., 1990; Asanuma et al., 1985; Field et al., 2008; Lynch et al., 1985), and dorsal pulvinar stimulation did elicit significant activity in LIP in the present study. One reason could be lower BOLD fMRI sensitivity in subcortical structures related to penetration of the RF signal due to surface RF coils, higher susceptibility to residual motion and/or a scanning protocol that was not optimized for subcortical functional imaging (Arcaro et al., 2015; Miletić et al., 2020). Another explanation could be asymmetric anatomical and functional LIP-pulvinar vs. pulvinar-LIP connectivity, which remains to be elucidated in future work.

### 4.3 Similarities and differences of LIPd vs. dPul microstimulation effects

Given the considerable differences between thalamic and cortical local organization and projections, it was not *a priori* obvious how much similarity of microstimulation effects should be expected. The comparison of effective connectivity patterns of dPul and LIPd revealed many similarities across prefrontal, parietal, and most extensively, STS areas. This confirms that both regions belong to the same functional brain network supporting the visuospatial guidance of eye movements and attention (Bridge et al., 2015; Gold and Shadlen, 2007; Grieve et al., 2000; Halassa and Kastner, 2017). In support of this notion, inactivation of either region results in a saccade choice bias towards the ipsilesional space (Christopoulos et al., 2018; Wardak et al., 2002; Wilke et al., 2013, 2012). Nonetheless, dPul plays a role in reach as well as saccade choices, while LIP inactivation results mainly in an effector-specific bias for eye movements (Christopoulos et al., 2018; M. Wilke et al., 2010). While the prefrontal and parietal regions have been classically associated with spatial attention and decisions, one interesting location of overlap of the dPul and LIPd stimulation effect was the middle and the anterior portion of STS, encompassing areas FST, IPa, PGa and TPO. This finding is consistent with anatomy, since these areas receive strong input from both LIP (Blatt et al., 1990; Seltzer and Pandya, 1994) and the dorsal pulvinar (Gutierrez et al., 2000; Yeterian and Pandya, 1991, 1989). This underlines the importance of these STS regions in the control of eye movements and visuospatial functions found in earlier electrophysiological and lesion studies (Bruce et al., 1981; Karnath, 2001; Luh et al., 1986; Scalaidhe et al., 1995; Watson et al., 1994). Confirming and extending these findings, these STS regions and the adjacent area called PITd have recently emerged as consistently being activated in monkey fMRI studies employing visual attention and delayed saccade tasks (Bogadhi et al., 2018; Caspari et al., 2015; Kagan et al., 2010; Patel et al., 2015). It is worth noting that inactivation-induced changes in these STS regions were also a common denominator of LIP, dPul and superior colliculus inactivation studies, accompanying contralesional saccade selection and attentional deficits (Bogadhi et al., 2019; Klink et al., 2021; Melanie Wilke et al., 2010; Wilke et al., 2013, 2012).

Some activations in STS elicited by pulvinar and LIP stimulation seem to correspond to regions that have previously been characterized as “face patches” (Moeller et al., 2008; Tsao et al., 2006). While we lacked the functional localizer for faces in our study and cannot disentangle direct from indirect activation pathways, the connectivity of the pulvinar with face patches is consistent with previous studies in macaques. A seminal es-fMRI study showed activity in the inferior pulvinar evoked by stimulation of the middle and anterior face patches (ML, AL, AM) (Moeller et al., 2008), an anatomical tracing study reported connections between several face patches (PL, ML, AL, AM) and the ventral (lateral and inferior) pulvinar (Grimaldi et al., 2016), and a resting-state fMRI study detected functional connectivity between face patches and dorsal and ventral pulvinar (Schwiedrzik et al., 2015). Although the co-activation along the STS elicited by pulvinar and LIP stimulation was not specific for face patches, we speculate that this shared connectivity might facilitate attention to faces by routing information from early visual areas to IT cortex and/or help to filter the important (e.g. facial) information from behaviorally irrelevant features (Bridge et al., 2015; Hesse and Tsao, 2020; Kastner et al., 2020). These data are also compatible with the notion of pulvinar involvement in face processing and social cognition, e.g. via a projection to amygdala (Nguyen et al., 2013; Saalmann and Kastner, 2013), but the exact connectivity and contribution of different pulvinar subnuclei remain to be elucidated.

Major differences between the distal activation patterns following dPul vs. LIPd stimulation were the LIPd-elicited activations in posterior IPS / PO sulci, the medial posterior parietal area 7m-PGM (Thier and Andersen, 1998), as well as in the homotopic LIP and adjacent IPS areas in the opposite hemisphere. In contrast, the pulvinar in the opposite hemisphere was not activated by the pulvinar stimulation.

### 4.4 Task-dependent effects of microstimulation on BOLD activity

While structural connectivity is relatively fixed, dynamic synaptic connectivity can vary as a function of external inputs and behavioral demands. Thus, the combination of es-fMRI with a behavioral task can be used to investigate the contribution of a given brain region to underlying brain and behavioral processes. Only a few es-fMRI studies investigated microstimulation effects as a function of external inputs or behavioral task contingencies (Ekstrom et al., 2009, 2008; Premereur et al., 2013). A pioneering study using low current FEF microstimulation (26-58 µA) showed that higher order extrastriate areas directly connected to the FEF exhibited microstimulation effect irrespective of visual stimulation (mimicking attention-driven additive “baseline shifts”), while polysynaptically connected V1 showed microstimulation-induced activation only in the presence of visual stimulus, mostly in voxels not directly responsive to the visual stimulus (Ekstrom et al., 2008). Similarly, FEF stimulation induced strong fMRI signal increase in visual areas for the low contrast stimuli, and no or even negative effects for high contrast stimuli that already strongly drove fMRI activity (Ekstrom et al., 2009). Another study from the same group (Premereur et al., 2013) specifically addressed the task-dependence of FEF stimulation effects (83-316 µA) by comparing activity between fixation and saccade tasks with matched visual inputs. This study reported higher es-fMRI effects in visual areas in the more demanding saccade task as compared to passive fixation, but again in the voxels that were not strongly activated by the visual stimulus.

Based on these studies, and the predominately contralateral tuning of LIPd and dPul cue and delay period activity both on the level of single neurons and fMRI signals (Blatt et al., 1990; Caspari et al., 2015; Dominguez-Vargas et al., 2017; Fiebelkorn et al., 2019; Kagan et al., 2010; Patel et al., 2010; Schneider et al., 2020), we hypothesized that the effects of dPul and/or LIP stimulation might depend on the visual stimulation and the direction of the upcoming saccade. For instance, if microstimulation acts as a multiplicative gain mechanism we might expect the highest fMRI-elicited enhancement in the contraversive saccade task condition, at least in the stimulated hemisphere. On the other hand, it could be that the microstimulation effect is strongest when the initial (no microstimulation) task activity is weakest. These effects could be manifested both at the level of single ROIs, and/or across ROIs.

On the level of single trial BOLD response amplitudes within each ROI, there was little indication of interaction between the stimulation and the task, in part due to a large variability of single trial timecourses. The primary ROIs in the present study were selected on the basis of activity enhancement by the microstimulation pooled over all task conditions, regardless of their task responsivity and other properties such as spatial tuning. Although it might appear that such selection, while unbiased for any specific task condition, favors ROIs without task-dependence of stimulation effect, the inspection of ROI timecourses and response amplitudes indicated that a strong positive activation in at least one of the three task conditions was enough to produce a significant “combined” t-statistics. It is thus unlikely that we missed the major regions with task-dependent stimulation effects. Across ROIs, however, the interaction was observed in both pulvinar and LIP stimulation datasets. The direction of the effect was not consistent however between the two animals – while in monkey C according to our initial prediction the contraversive saccade task condition produced the strongest microstimulation enhancement, it was not the case for monkey B (ipsiversive or fixation > contraversive).

We thus investigated how pulvinar and LIP microstimulation effects in the different tasks might be influenced by the initial spatial selectivity of stimulation-activated ROIs. Here we found a consistent relationship between the degree of contraversive selectivity and the strength of microstimulation-elicited activation, across ROIs. Specifically, the higher the contraversive selectivity of an ROI, the higher is the stimulation effect when monkeys are cued to perform a saccade into the ipsiversive hemifield. Conversely, such relationship was often inverted for the contraversive saccade task, during which ipsiversively-selective ROIs exhibited a higher stimulation effect. This “X-shape” interaction between the task condition and the initial spatial selectivity is reminiscent of FEF stimulation studies described above, showing that microstimulation effects in visual cortices are stronger with weaker visual stimuli and in task conditions not favored by the microstimulation-activated ROI (Ekstrom et al., 2009; Premereur et al., 2013). A plausible explanation is that strong initial task-related response leaves less “room” for modulation of neuronal activity (or a hemodynamically-coupled BOLD signal) by microstimulation before saturation is reached.

In line with these observations, the model that best described microstimulation effect amplitudes across ROIs was the additive mechanism, but with strength of the additive factor scaled down by the initial response amplitude. Across datasets, this model provided a better fit than a simple additive model, and was by far better than the multiplicative model. A simple qualitative simulation was able to reproduce the observed “X-shape” interaction pattern between the task condition and the initial spatial selectivity.

It remains to be elucidated how these microstimulation-elicited modulations of neural signals during the delay period might be relevant for ensuing spatial behavior. The present experiment was not designed to address stimulation-contingent changes of behavior. We used cognitively simple, single target instructed memory saccade task with a long and fixed delay, and the repeated microstimulation paradigm with the last stimulation train ending 800 ms before the end of the delay period, more than 1 s before the saccade onset (given ∼300 ms saccade latency). Hence, the modest behavioral effects were expected. Our previous study in the pulvinar found large latency increases only when the stimulation was delivered around the time of saccade onset (Dominguez-Vargas et al., 2017). Furthermore, in the present study contraversive saccades were delayed more than ipsiversive saccades while the opposite was true for the stimulation around the Go signal in the previous study. This suggests that stimulating repeatedly and temporally dissociated from the motor action might affect behavior via a different mechanism. Similarly, two previous LIP microstimulation studies that investigated its influence on perceptual decisions stimulated just before the saccade and reported modest saccade latency effects (reaction time increase for out-receptive-field choices and decrease for in-receptive-field choices) (Dai et al., 2014; Hanks et al., 2006). Therefore, faster trials with stimulation periods temporally closer to the action, and more demanding tasks, e.g. involving attentional selection or a choice between different response options might be required to study behaviorally-relevant effects of stimulation (Arsenault et al., 2014; Dominguez-Vargas et al., 2017).

Further work is also needed to derive conclusions about brain-wide changes in spatial tuning due to microstimulation, given the asymmetric effect in the two hemispheres and its task-dependence. The stronger effect in the stimulated hemisphere that also involved more activated regions could lead to changes of interhemispheric balance, and an overall stronger potentiation of contraversive spatial representations. Using a symmetrical set of task-based ROIs across the two hemispheres and analyzing microstimulation-induced changes in spatial tuning and task-related functional connectivity might help addressing these outstanding questions.

## Authors contributions

**Igor Kagan:** Conceptualization, Methodology, Software, Formal analysis, Resources, Data Curation, Writing – Original Draft, Writing – Review & Editing, Visualization, Project administration, Supervision, Funding acquisition; **Lydia Gibson:** Conceptualization, Methodology, Software, Formal analysis, Investigation, Writing – Original Draft, Visualization; **Elena Spanou:** Methodology, Software, Investigation, **Melanie Wilke:** Conceptualization, Methodology, Resources, Writing – Original Draft, Writing – Review & Editing, Supervision, Funding acquisition

## Acknowledgements

We thank Dr. Vladislav Kozyrev for helping to collect a subset of the MRI data, Dr. Sebastian Moeller for sharing the design of the MR-compatible electrode drive and helping with the microstimulation setup, Dr. Peter Dechent and Dr. Juergen Baudewig for helping with setting up MRI data acquisition, Dr. Jochen Weber for providing support for NeuroElf, Alexandra Witt for help with CHARM and SARM atlas processing, and Dr. Caspar Schwiedrzik for sharing the face patch data. We thank Ira Panolias, Dr. Daniela Lazzarini, Sina Plümer, Klaus Heisig, Leonore Burchardt, and Dirk Pruesse for technical support. We also thank Stefan Treue, Alexander Gail, Hansjoerg Scherberger, members of the Decision and Awareness Group, the Sensorimotor Group, and the Cognitive Neuroscience Laboratory for helpful discussions.

