## Supplementary Material for "Effective connectivity and spatial selectivity-dependent fMRI changes elicited by microstimulation of pulvinar and LIP"

### Supplementary Figures

**Figure S1.** Electrode tip reconstruction probability maps in dorsal pulvinar and LIP, individual AC-PC space.
**Figure S2.** Electrode tip reconstruction probability maps in dorsal pulvinar, NMT v2 space.
**Figure S3.** Comparison of electrode localization and local activations in dPul and vPul.
**Figure S4.** Axial T1-weighted and T2-weighed (acquired in-plane with EPI) scans, resliced in AC-PC space, dPul stimulation site.
**Figure S5.** Axial T2-weighed (acquired in-plane with EPI) and T2*-w. EPI scans, resliced in AC-PC space, dPul stimulation site.
**Figure S6.** Overlaid axial T2-weighed and EPI scans, and separate EPI images, resliced in AC-PC space, dPul stimulation site.
**Figure S7.** Overlaid axial T2-weighed and EPI scans, resliced in AC-PC space, LIPd-p and vPul stimulation sites.
**Figure S8.** NMT v2 CHARM and SARM atlas ROIs.
**Figure S9.** Electrode susceptibility dropout ROI, dPul stimulation site.
**Figure S10.** Electrode susceptibility dropout ROI, vPul stimulation site.
**Figures S11 – S14.** Behavioral performance and eye movements.
**Figure S15.** Activation in and around the pulvinar, with overlaid warped NMT v2 SARM atlas, dPul stimulation site.
**Figure S16.** dPul and LIPd stimulation activation maps in the volume space.
**Figure S17.** dPul and LIPd stimulation activation maps, fixation only task condition.
**Figure S18.** Pulvinar and LIPd stimulation activation maps in the NMT v2 space.
**Figure S19.** Comparison of dPul(a) and dPul stimulation effects.
**Figure S20.** Overlap of dPul and LIPd stimulation effects, fixation only task condition.
**Figure S21.** Stimulation in vPul (current strength 100 µA).
**Figure S22.** Comparison of stimulation effect strength in cortical areas after dPul and vPul stimulation.
**Figure S23.** Comparison of low current 100 µA dPul and vPul stimulation effects, monkey C.
**Figure S24.** Activation of frontal regions by a low 100 µA current vPul stimulation.
**Figure S25.** Stimulation effects derived from event-related average response amplitudes.
**Figure S26.** Stimulation effects for each task and within-ROI ANOVA results, pulvinar datasets.
**Figure S27.** Stimulation effects for each task and within-ROI ANOVA results, LIPd datasets.
**Figure S28.** Microstimulation effect dependence on contraversive spatial selectivity and task. Monkey C.
**Figure S29.** Microstimulation effect dependence on contraversive spatial selectivity and task. Monkey B.
**Figure S30.** Qualitative simulation of additive-scaled by initial response amplitude stimulation enhancement.

### Supplementary Tables

**Table S1.** Electrode tip positions in the dPul pulvinar sites.
**Table S2.** Summary of selected regions of interest.
**Table S3.** Regions of interest selected for each animal and stimulation site dataset.
**Table S4.** Stimulation effects in different task conditions, *NMT atlas ROIs*.
**Table S5.** Relationship between stimulation effect, spatial selectivity and task condition, *separately for each hemisphere*, stimulation effect ROIs.
**Table S6.** Relationship between stimulation effect, spatial selectivity and task condition, *NMT atlas ROIs.*
**Table S7.** Stimulation effect model fitting.

### Supplementary Results

#### Task performance and eye movements

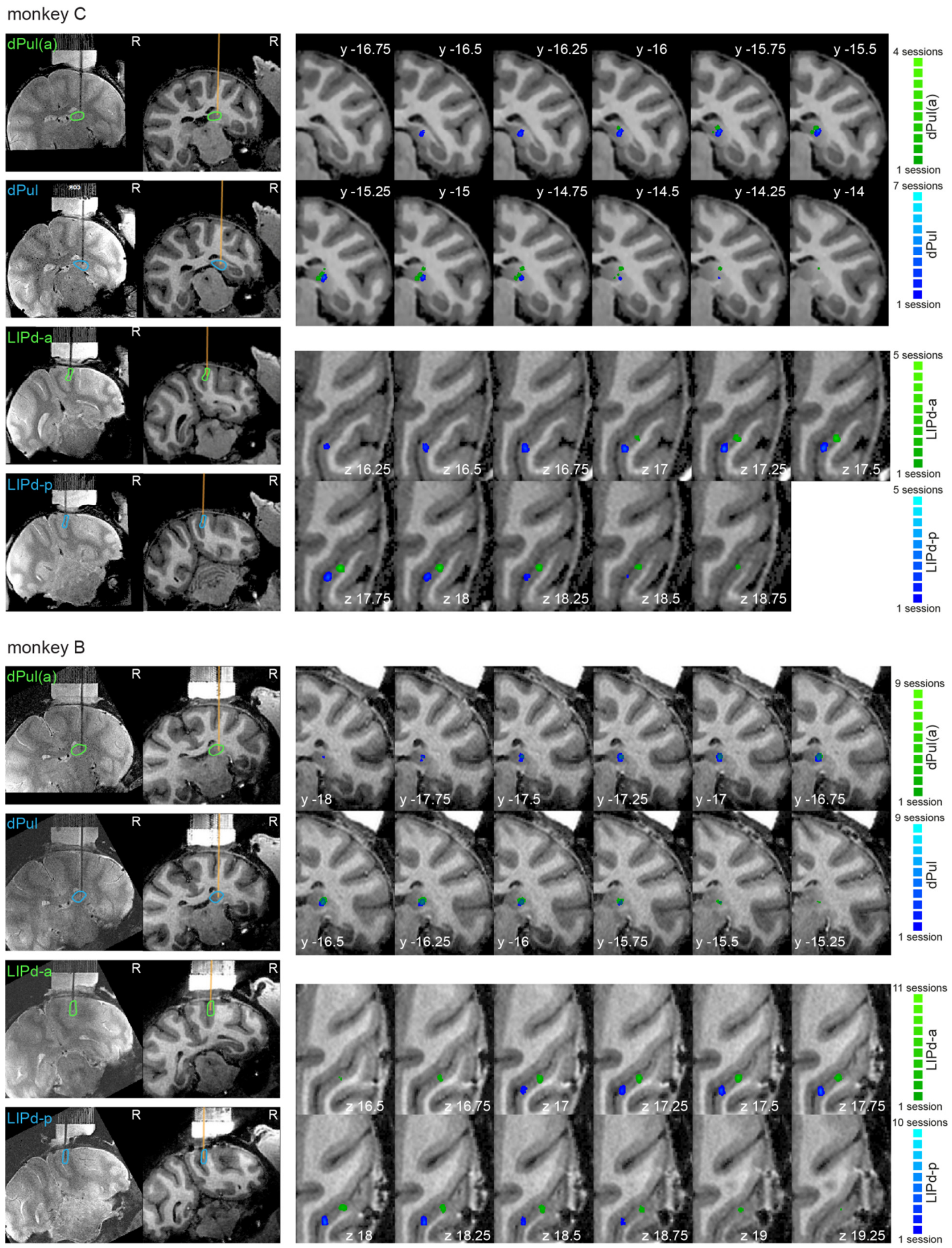

**Figure S1. (A)** Electrode localization in dPul and LIPd. Left panels: electrode positions measured in T2-weighted MR images and reconstructed in T1-weighted images, both aligned to the chamber vertical axis, in example sessions with microstimulation in dPul and vPul. Outlines mark the respective target region. Right panels: probability maps of electrode tip positions across sessions displayed on a T1-weighted MR image aligned to standard AC-PC space (dPul – coronal sections, LIP – axial sections, only a part of the right hemisphere focusing on the pulvinar or the intraparietal sulcus is shown). R - right, y - distance from AC-PC origin along the anterior/posterior plane in mm, z: distance from AC-PC plane along the dorsal-ventral dimension in mm.

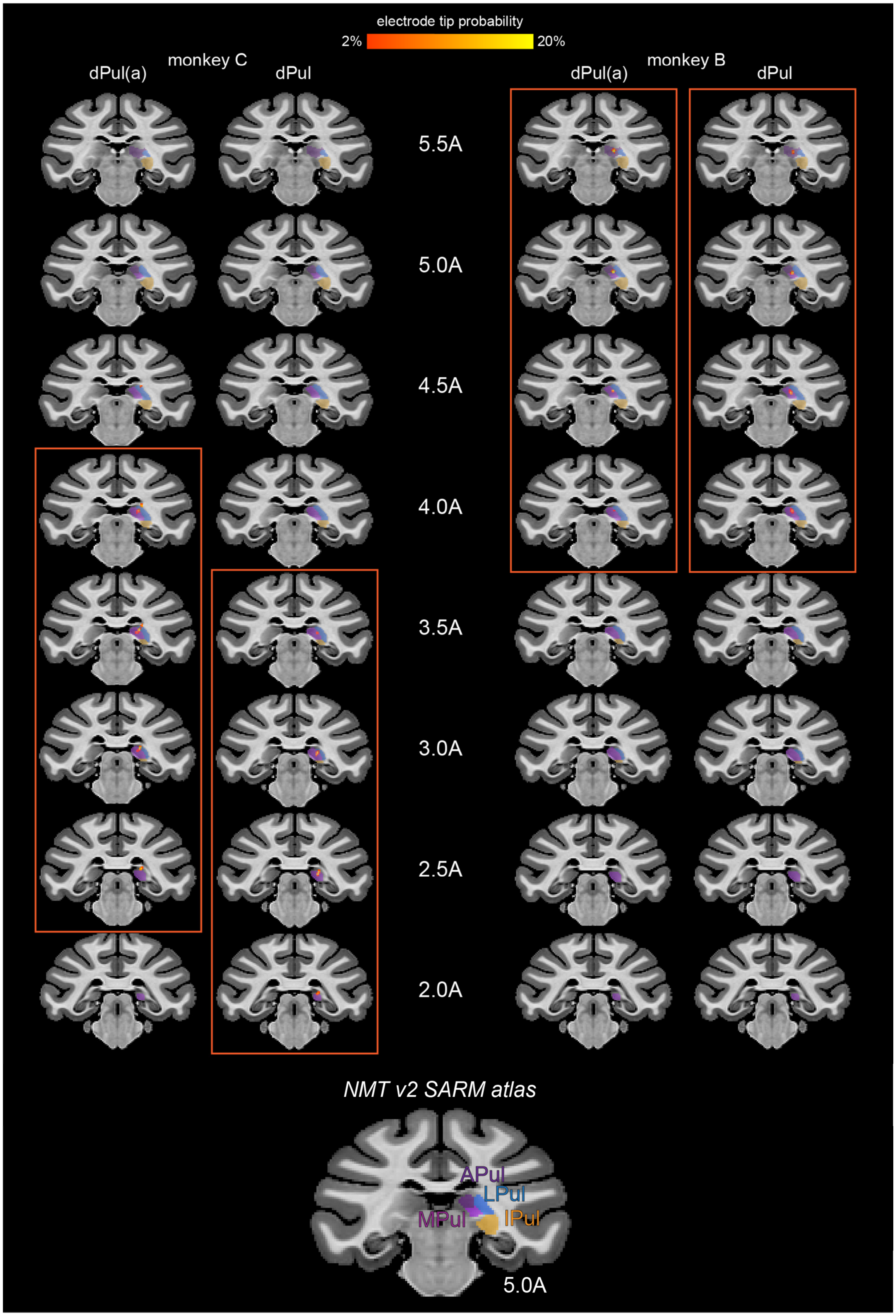

**Figure S2.** Electrode tip reconstruction probability maps in dPul(a) and dPul, NMT v2 space. The inset below shows the SARM pulvinar parcellation. Coronal sections are labeled in respect to the template stereotaxic origin (A – anterior), in mm.

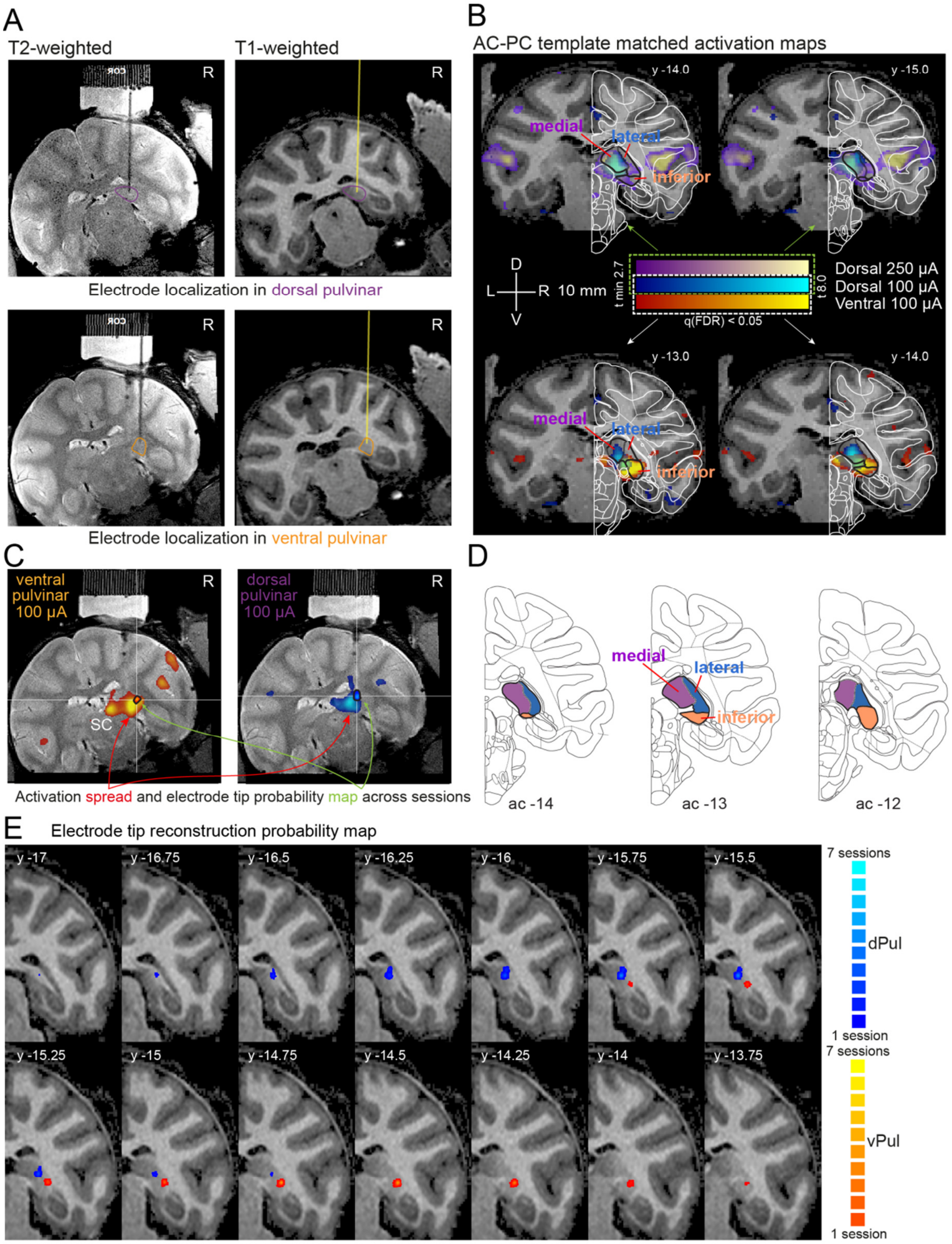

**Figure S3.** Comparison of electrode localization and local activations in dPul and vPul. **(A)** Electrode positions in T2-weighted MR images and reconstructed in T1-weighted images, aligned to the chamber vertical axis, in example dPul and vPul stimulation sessions. Outlines mark the respective target region. **(B)** Coronal sections showing microstimulation activation maps at the stimulation sites during high (250 µA) and low (100 µA) current stimulation of dPul (purple-yellow and blue/cyan, respectively) and vPul (red/yellow). Schematic outlines were adapted from the NeuroMaps atlas downloaded from BrainInfo (http://braininfo.rprc.washington.edu/PrimateBrainMaps/atlas/Mapcorindex.html). **(C)** The same chamber-normal coronal sections as in (A), with activation maps and overlaid probability map of electrode tip positions (outlined in black, see green arrows) across all 100 µA sessions. **(D)** Three example atlas sections (relative to the AC origin) depicting the medial, lateral and inferior pulvinar nuclei, as well as the brachium of the superior colliculus (bsc). **(E)** Probability maps of electrode tip positions across sessions in dPul (blue) and vPul (red) displayed on a T1-weighted MR image aligned to standard AC-PC space. R - right, y - distance from AC-PC origin along the anterior/posterior plane in mm.

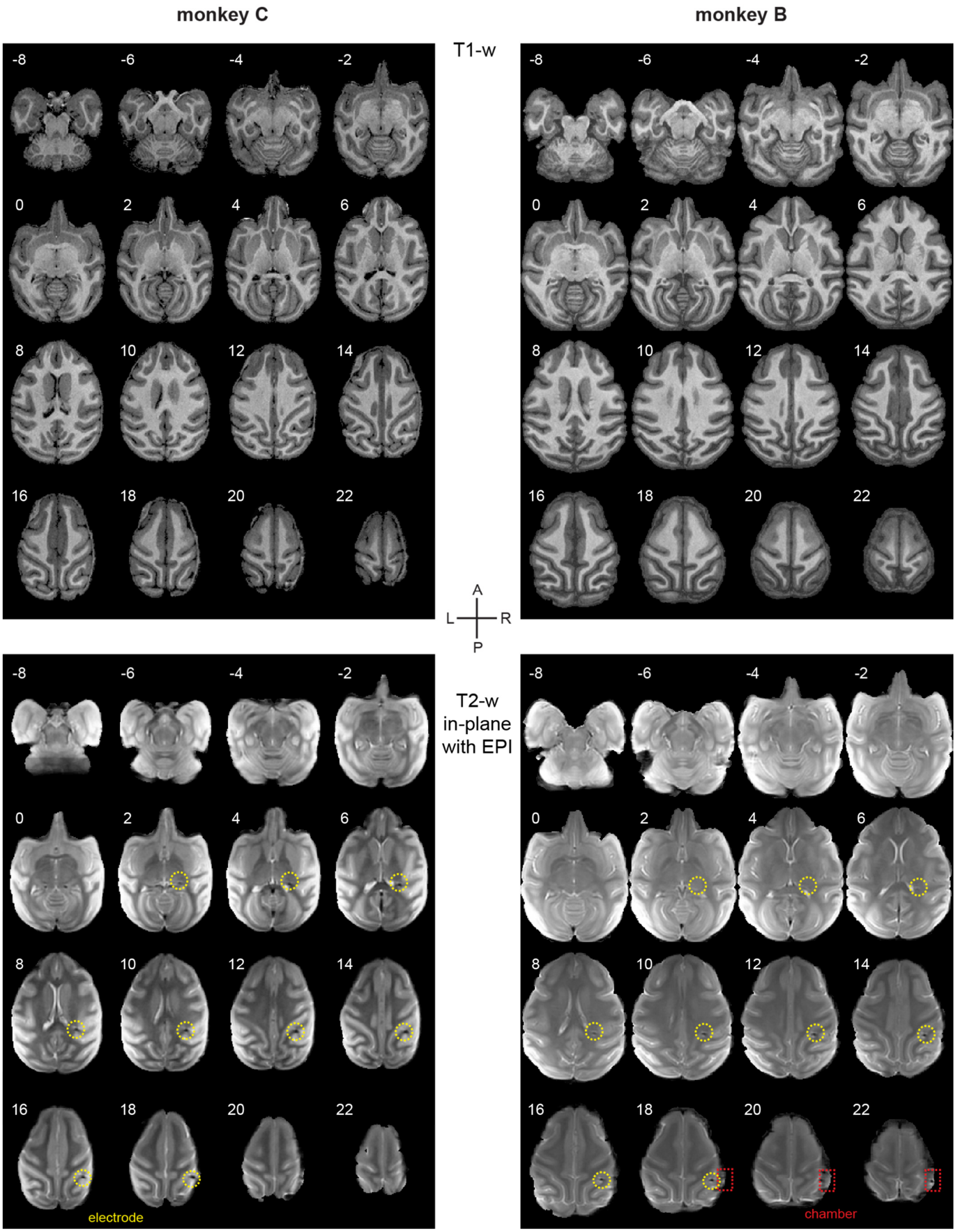

**Figure S4.** Axial T1-weighted and T2-weighed (acquired in-plane with EPI) scans, resliced in AC-PC space, dPul stimulation site. Axial sections are labeled relative to AC-PC plane. The T2-weighed scans were acquired in plane with EPI slices in each session; the image here shows the mean image across dPul stimulation sessions.

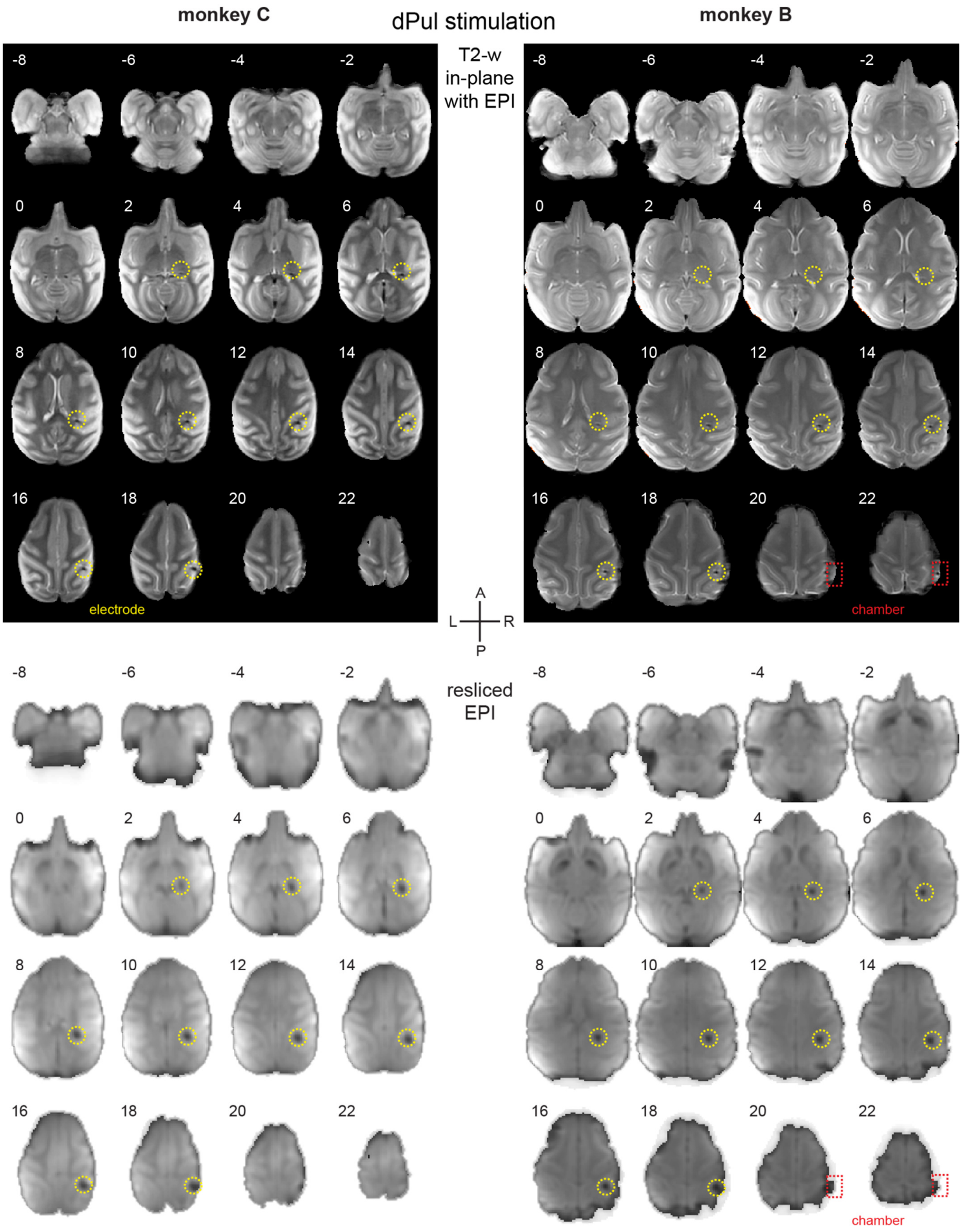

**Figure S5.** Axial T2-weighed (acquired in-plane with EPI, as in Figure S4) and T2*-weighed EPI scans, resliced in AC-PC space, dPul stimulation site. Most posterior occipital cortex shows EPI dropout artifacts due to magnetic susceptibility introduced by the edge of the bone cement headcap implant. Additionally, there is a dropout along the electrode shank. The EPI image shows the mean image across dPul stimulation sessions.

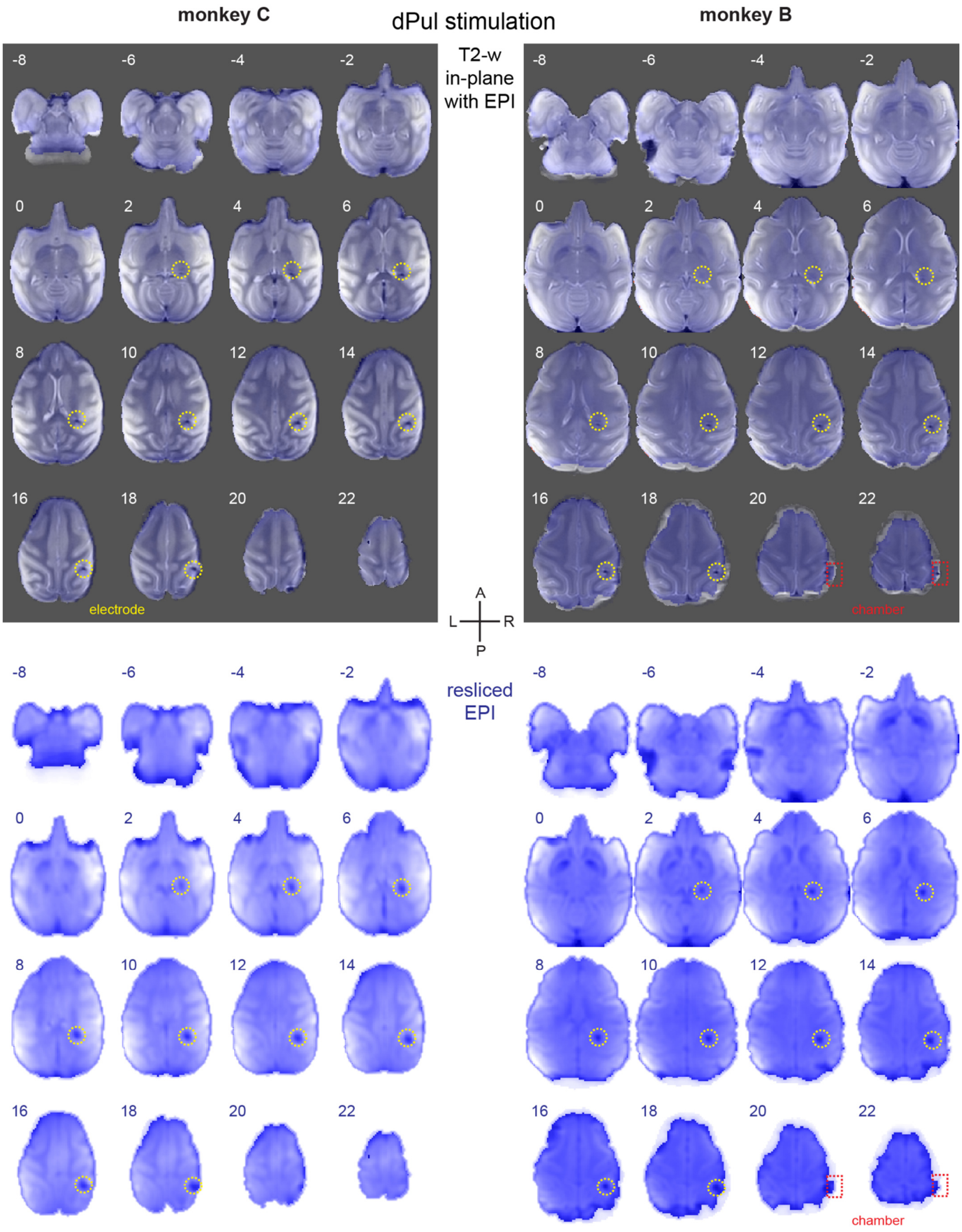

**Figure S6.** Overlaid axial T2-weighed and EPI scans, and separate EPI images, resliced in AC-PC space, dPul stimulation site. The comparison between T2-weighted anatomy and EPI highlights some occipital edge distortions, but otherwise a good match between EPI and underlying anatomy.

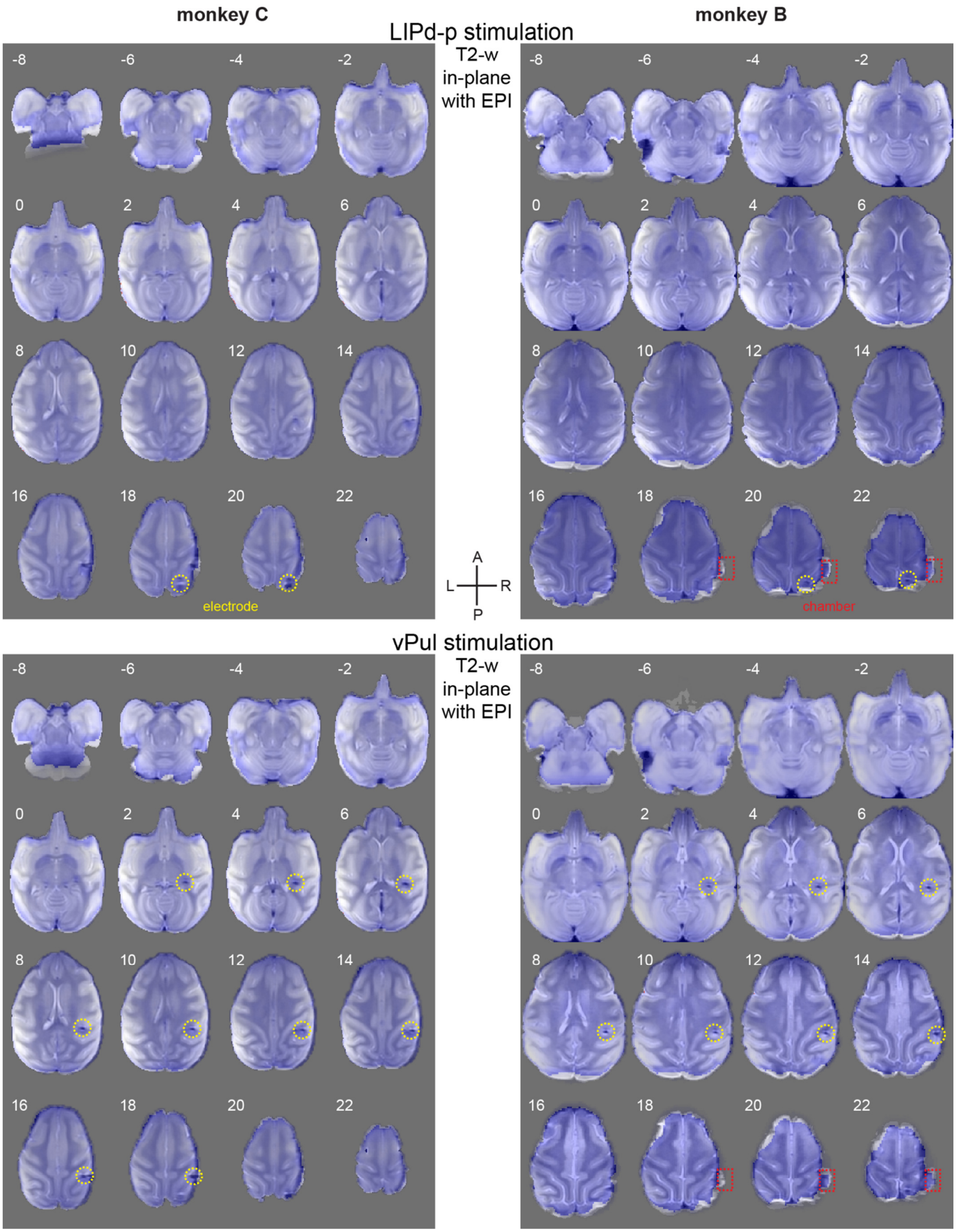

**Figure S7.** Overlaid axial T2-weighed and EPI scans, resliced in AC-PC space, LIPd-p and vPul stimulation sites. The comparison between T2-weighted anatomy and EPI highlights some occipital edge distortions, but otherwise a good match between EPI and underlying anatomy.

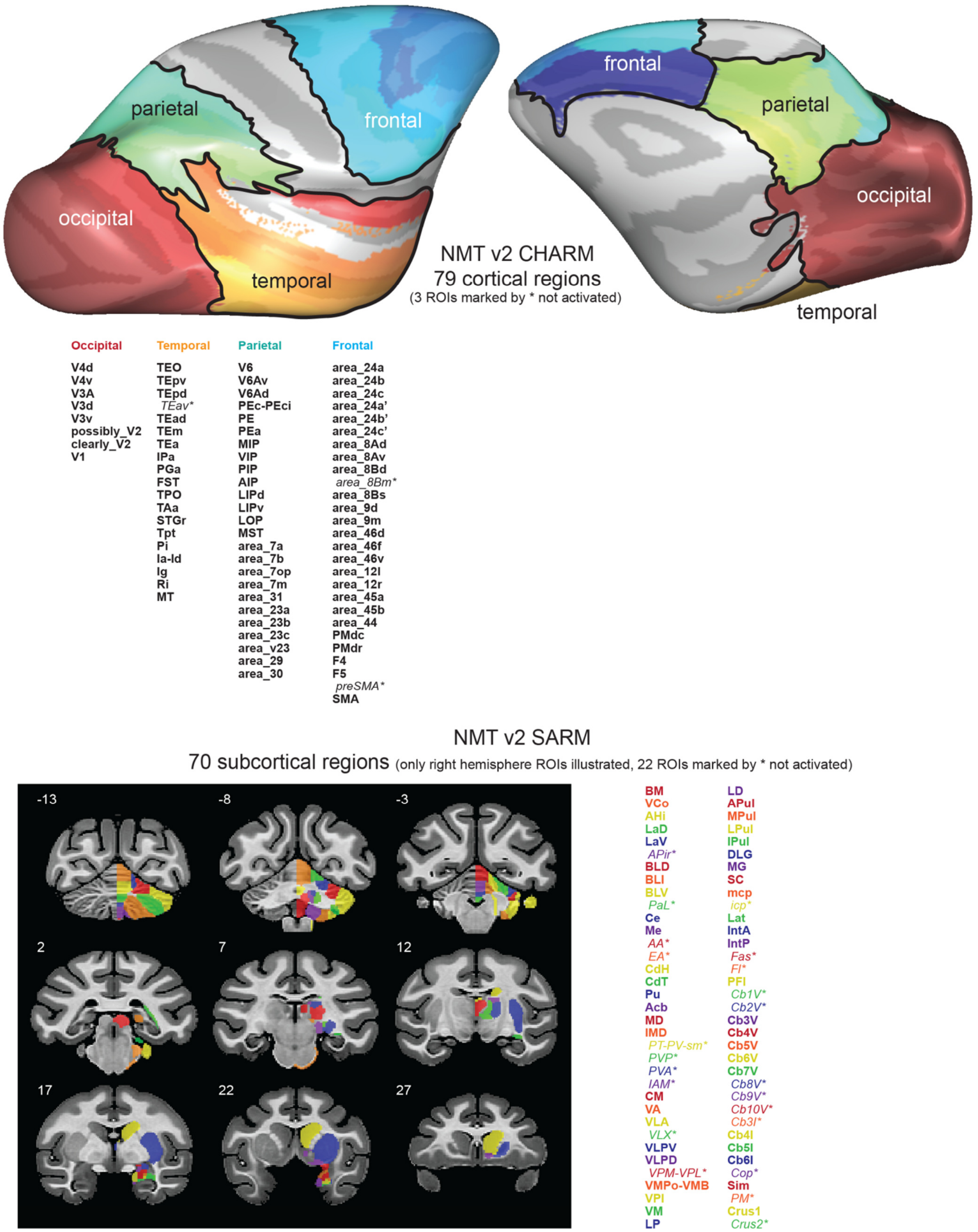

**Figure S8.** NMT v2 CHARM and SARM atlas ROIs, selected for the ROI analysis. See **Table S2** for details.

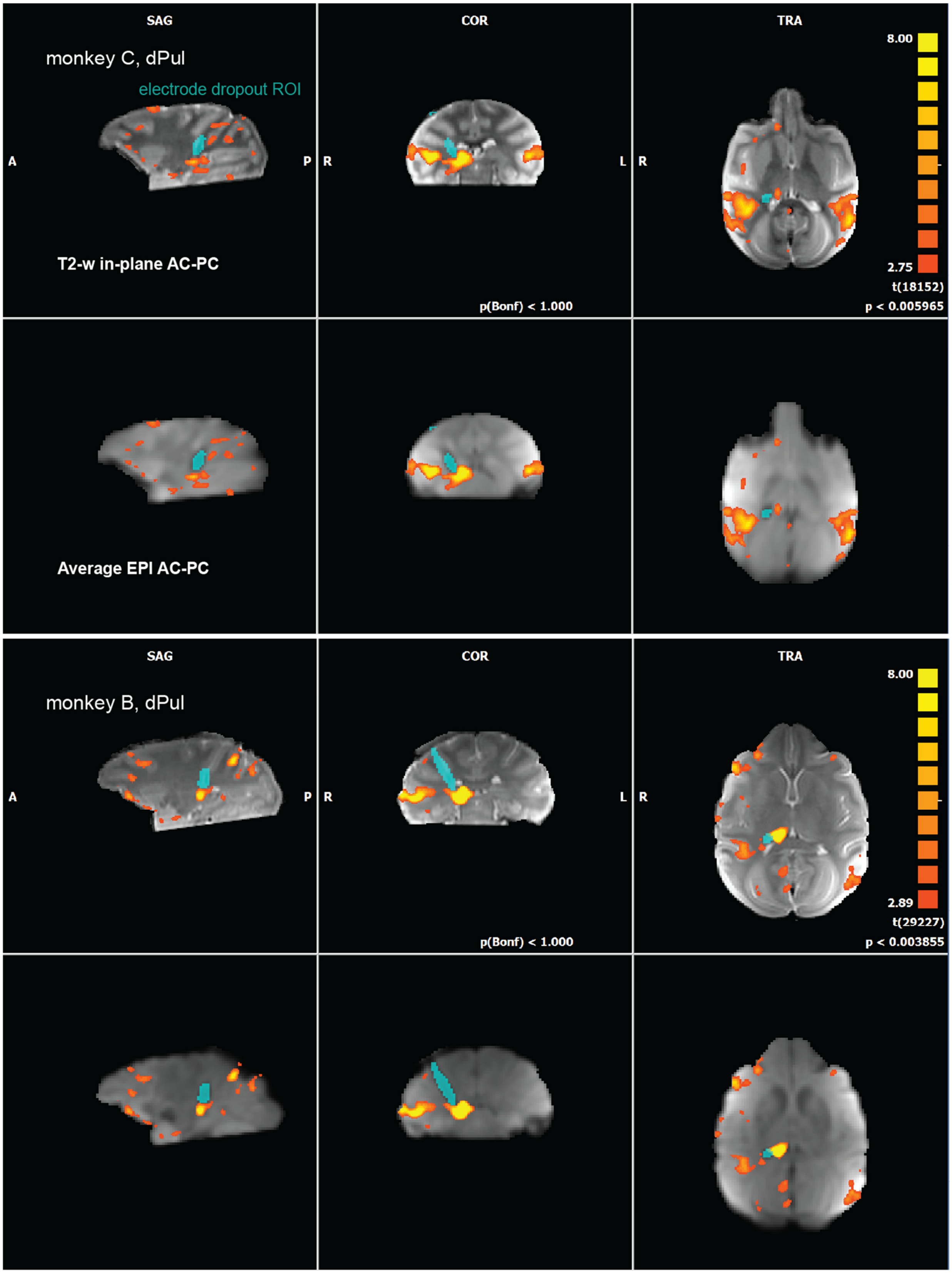

**Figure S9.** Electrode susceptibility EPI dropout ROI (shown on both T2-weighted – top row, and EPI – bottom row, for each monkey), and adjacent stimulation-elicited activation, in sagittal, coronal and axial sections. The dropout ROI was identified using intensity threshold on a mean EPI image across all sessions for each stimulation site: here dPul stimulation.

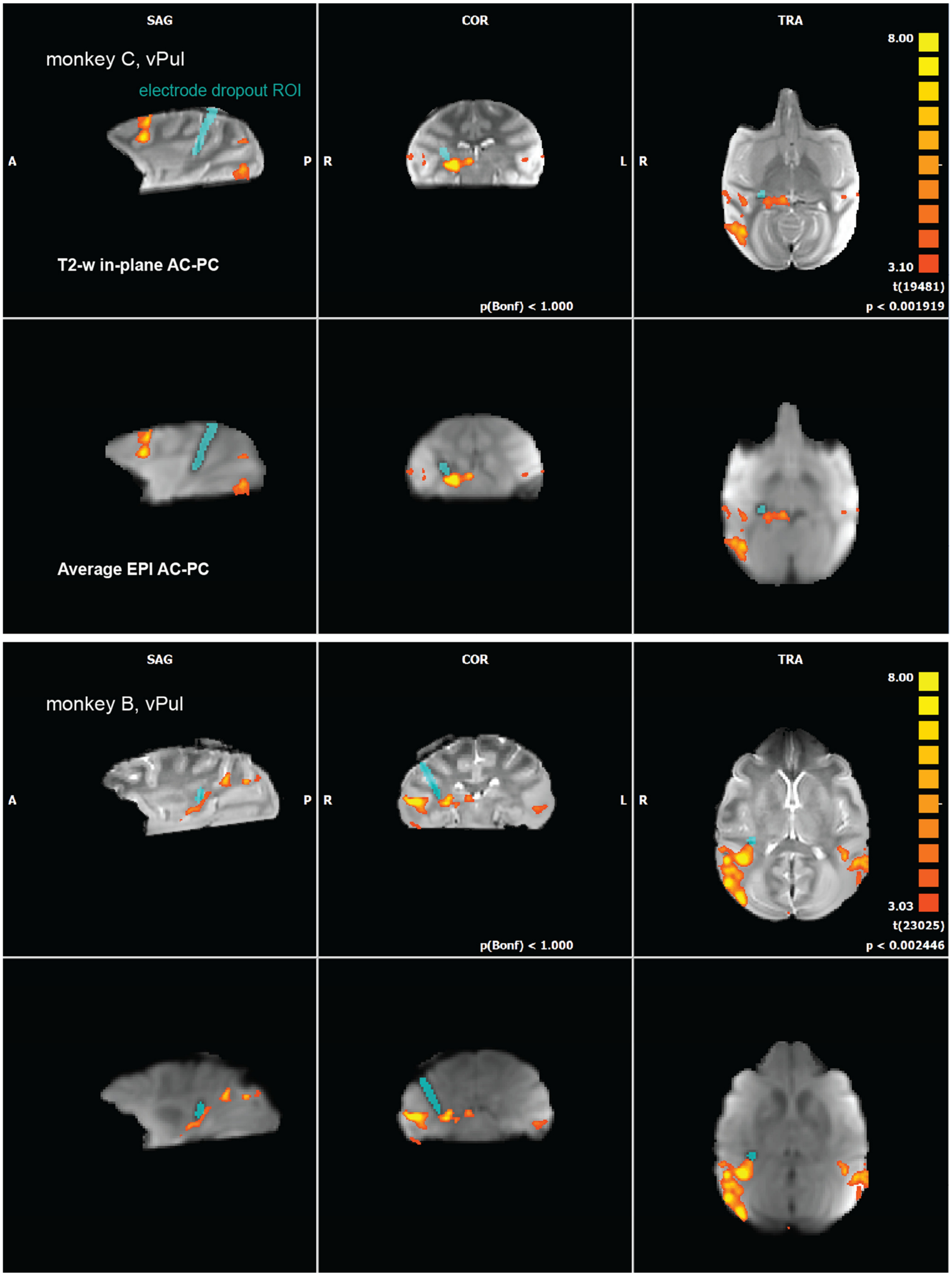

**Figure S10.** Electrode susceptibility dropout ROI (shown on both T2-weighted – top row, and EPI – bottom row, for each monkey), and adjacent stimulation-elicited activation, in sagittal, coronal and axial sections. The dropout ROI was identified using intensity threshold on a mean EPI image across all sessions for each stimulation site: here vPul stimulation.

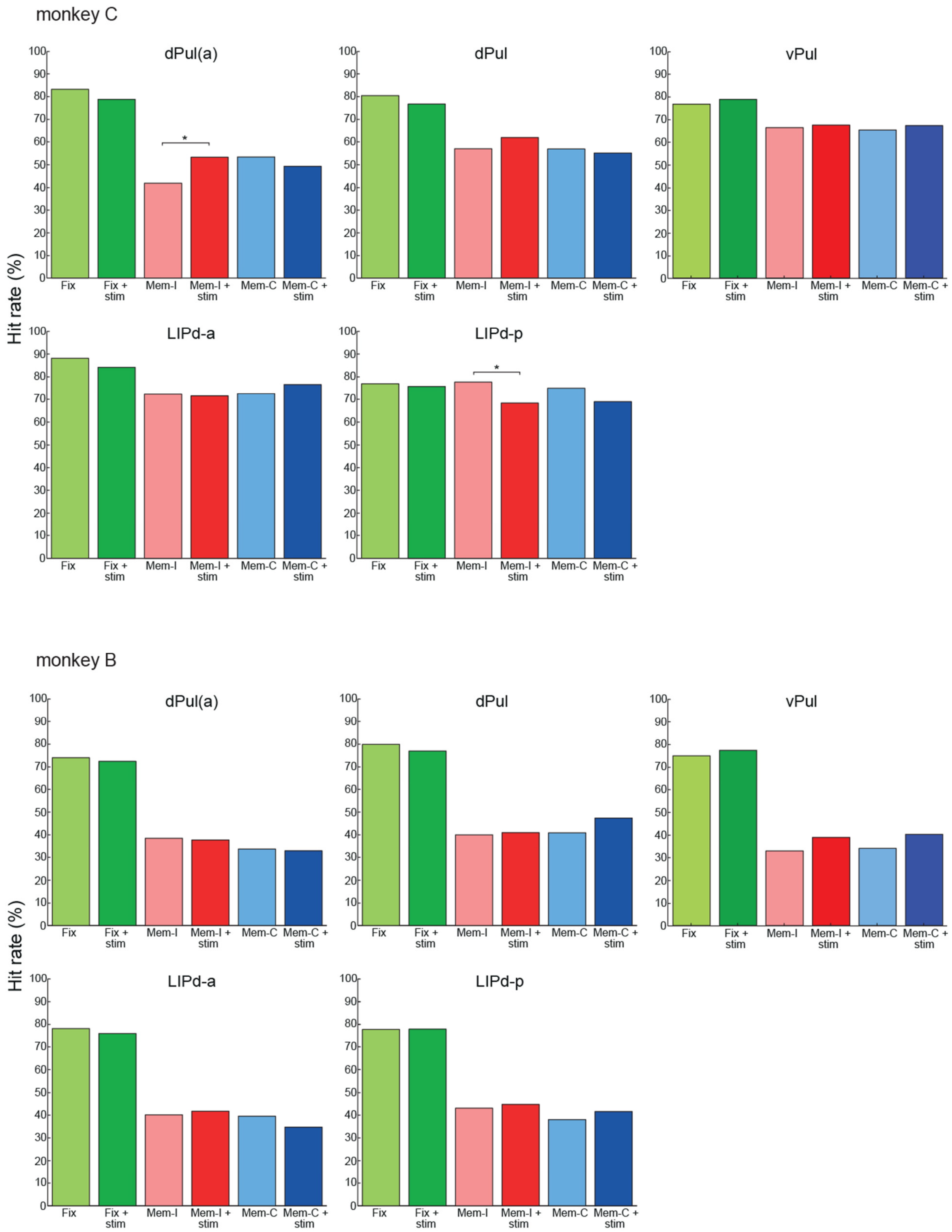

**Figure S11.** Hit rates (proportion of successful trials) across all trials from all sessions combined in the fixation, ipsiversive memory saccade, and contraversive memory saccade task in control (Fix, Mem-I, and Mem-C) and stimulation trials (Fix + stim, Mem-I + stim, Mem-C + stim) for stimulation in dPul(a), dPul, vPul, LIPd-a, and LIPd-p. * $p < 0.05$.

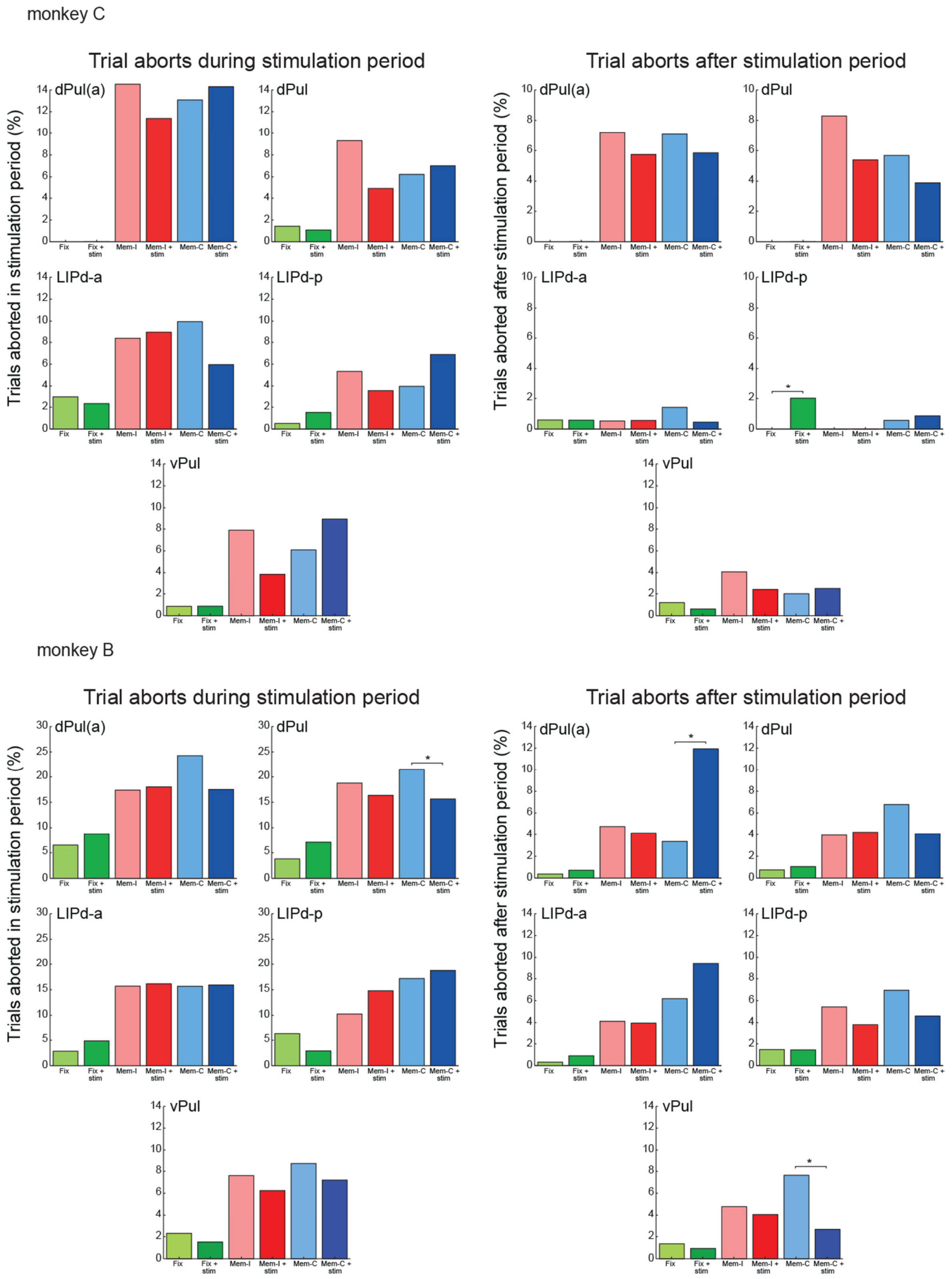

**Figure S12.** *Left two columns:* proportions of trials aborted *in the stimulation period* (or a corresponding period in no stimulation trials) in the fixation, ipsiverive memory saccade, and contraversive memory saccade task in control (Fix, Mem-I, Mem-C) and stimulation trials (Fix + stim, Mem-I + stim, Mem-C + stim) for stimulation in dPul(a), dPul, vPul, LIPd-a, and LIPd-p. *Right two columns:* proportions of trials aborted *after the stimulation period* (or a corresponding period in no stimulation trials) in the fixation, ipsiversive memory saccade, and contraversive memory saccade task in control (Fix, Mem-I, Mem-C) and stimulation trials (Fix + stim, Mem-I + stim, Mem-C + stim) for stimulation in dPul(a), dPul, vPul, LIPd-a, and LIPd-p. * $p < 0.05$.

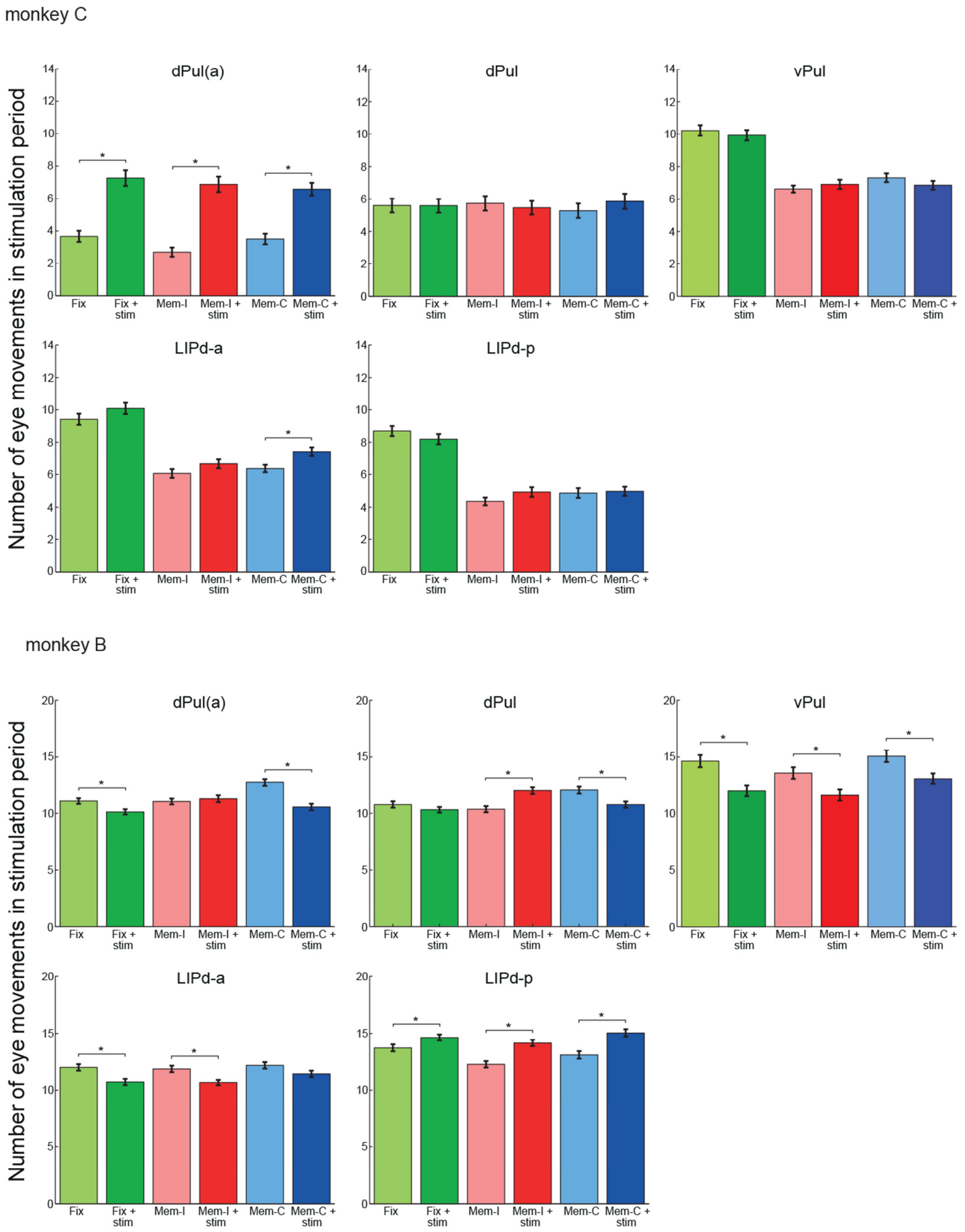

**Figure S13.** Mean number of eye movements in the stimulation period and standard errors of means across trials in the fixation, ipsiversive memory saccade, and contraversive memory saccade task in control (Fix, MemI, Mem-C) and stimulation trials (Fix + stim, Mem-I + stim, Mem-C + stim) for stimulation in dPul(a), dPul, vPul, LIPd-a, and LIPd-p. * $p < 0.05$.

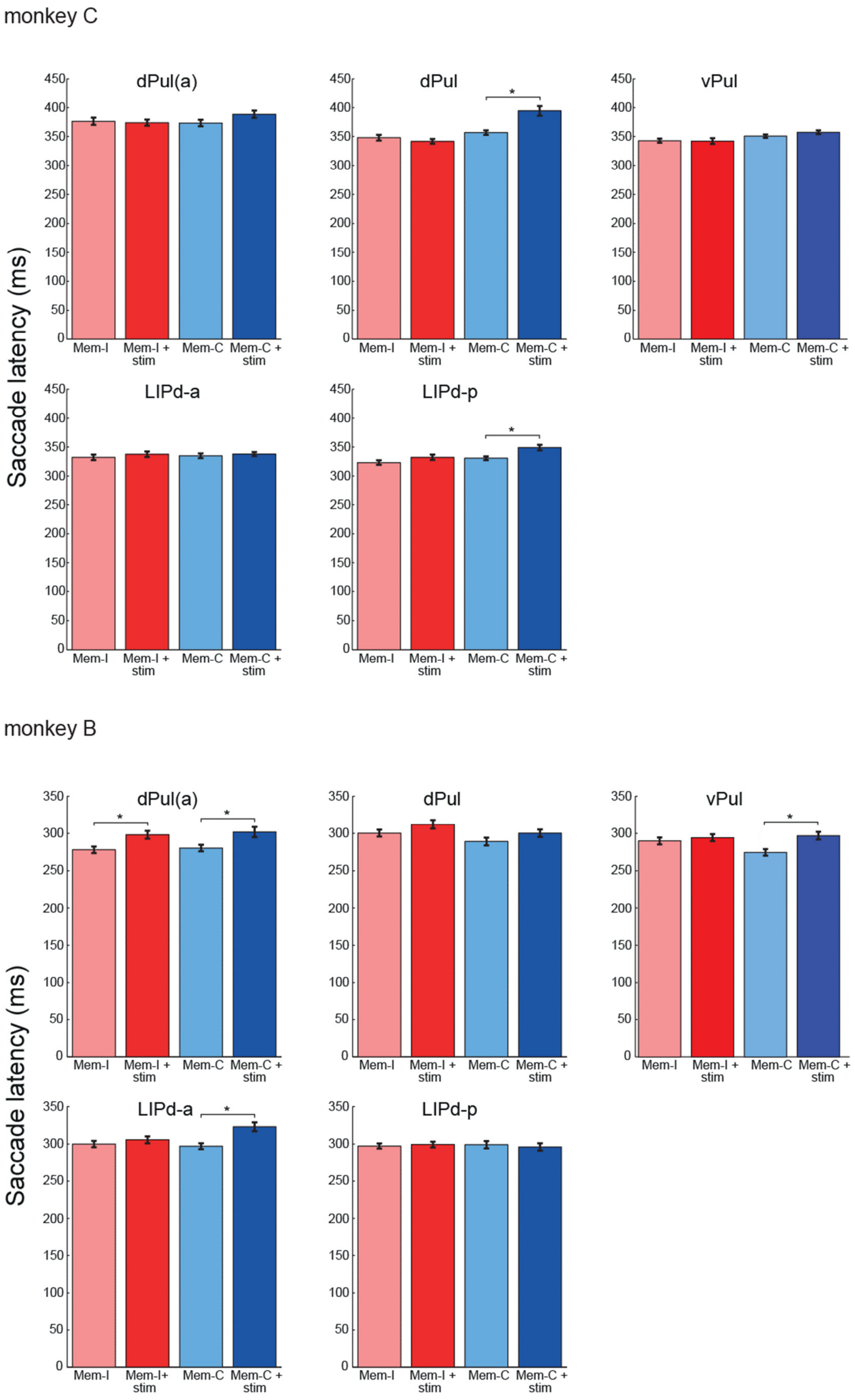

**Figure S14.** Mean saccade latencies and standard errors of means across trials in the ipsiversive memory saccade and contraversive memory saccade task in control (Mem-I, Mem-C) and stimulation trials (Mem-I + stim, Mem-C + stim) for stimulation in dPul(a), dPul, vPul, LIPd-a, and LIPd-p. * $p < 0.05$.

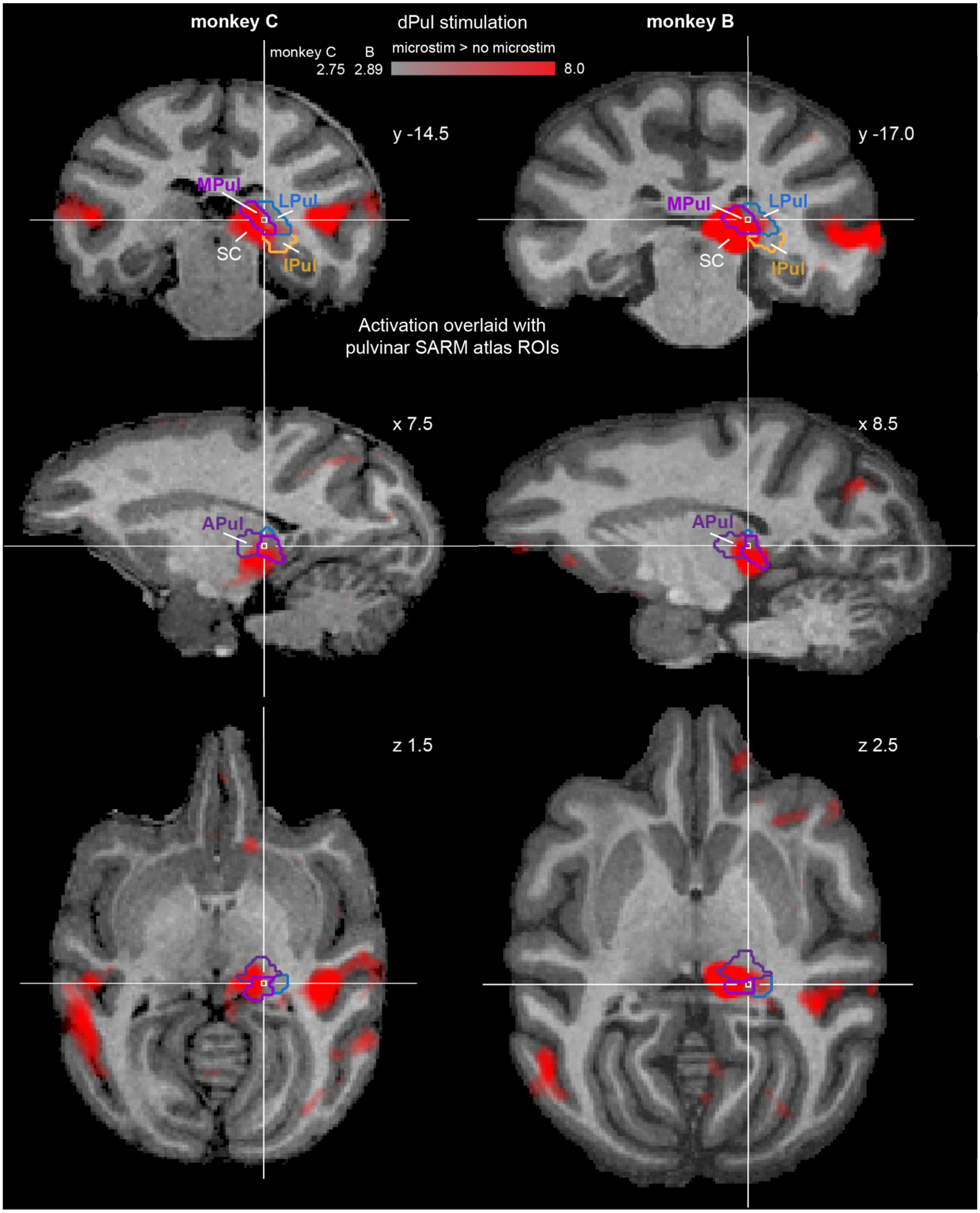

**Figure S15.** Activation in and around the pulvinar, with overlaid warped NMT v2 SARM atlas, dPul stimulation site. Here transparency of the activation map scales with significance.

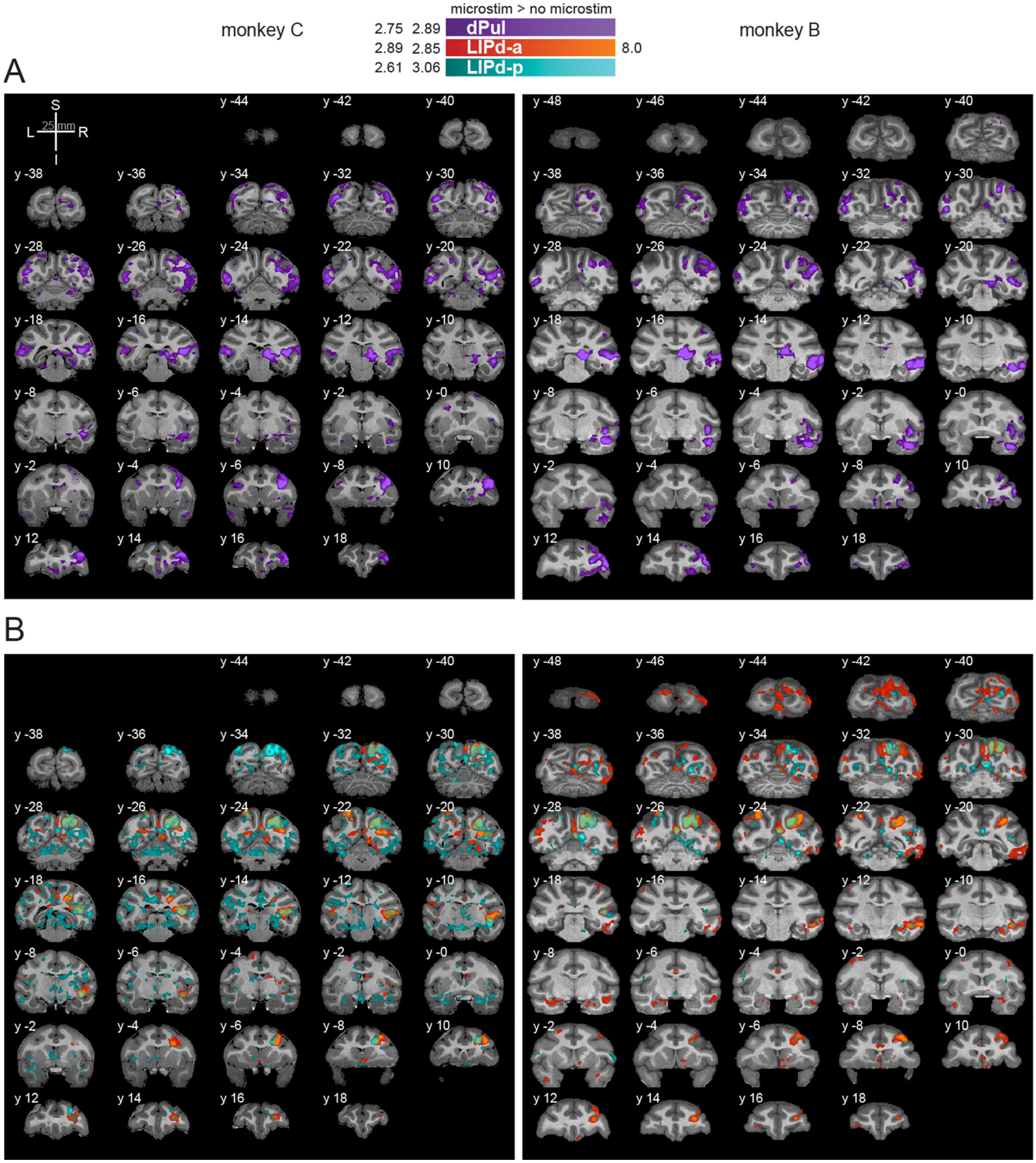

**Figure S16.** dPul and LIPd stimulation activation maps in the volume space, coronal sections (2 mm spacing). L – left, R – right, S – superior, I – inferior, y - distance from AC-PC origin along the anterior/posterior plane in mm.

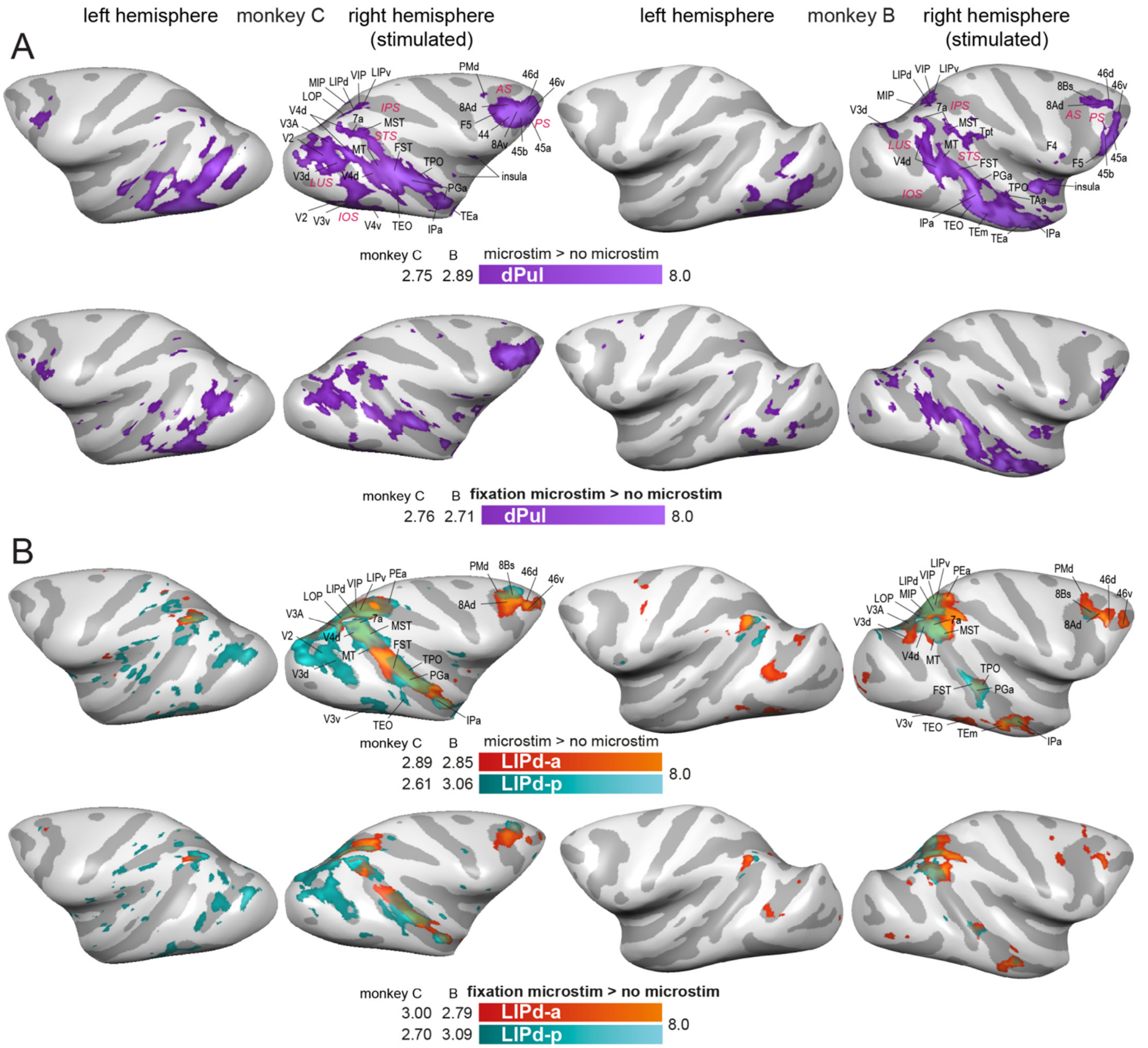

**Figure S17.** dPul and LIPd stimulation activation maps, fixation only task condition (bottom row in each panel), and all task conditions (top row), for direct comparison. **(A)** Stimulation in dPul. **(B)** Stimulation in LIPd-a and LIPd-p.

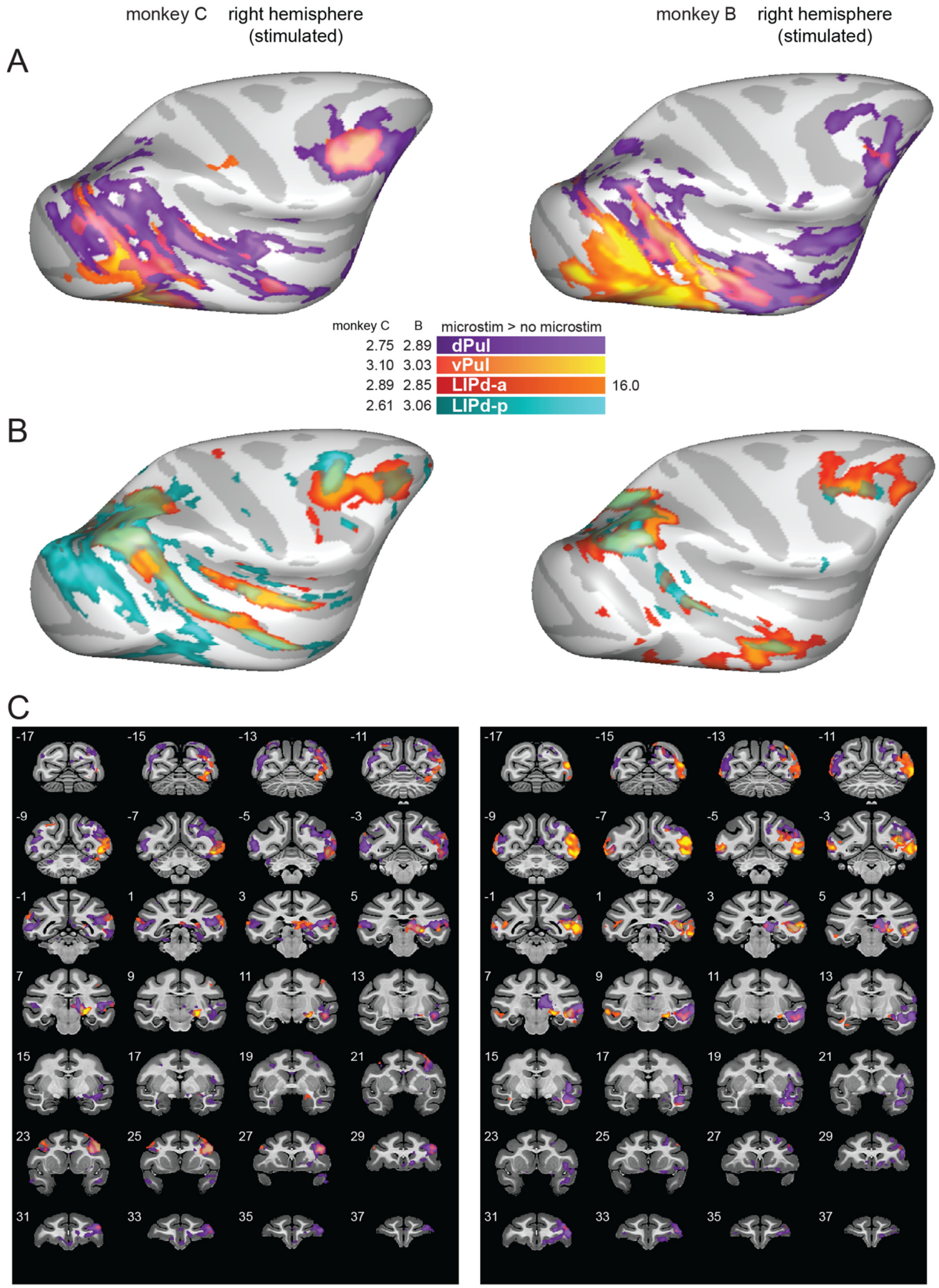

**Figure S18.** Pulvinar and LIPd stimulation activation maps in the NMT v2 space. **(A)** Dorsal and ventral pulvinar stimulation. **(B)** LIPd-a and LIPd-p stimulation. **(C)** Coronal sections, relative to the NMT v2 stereotaxic zero origin, 2 mm spacing.

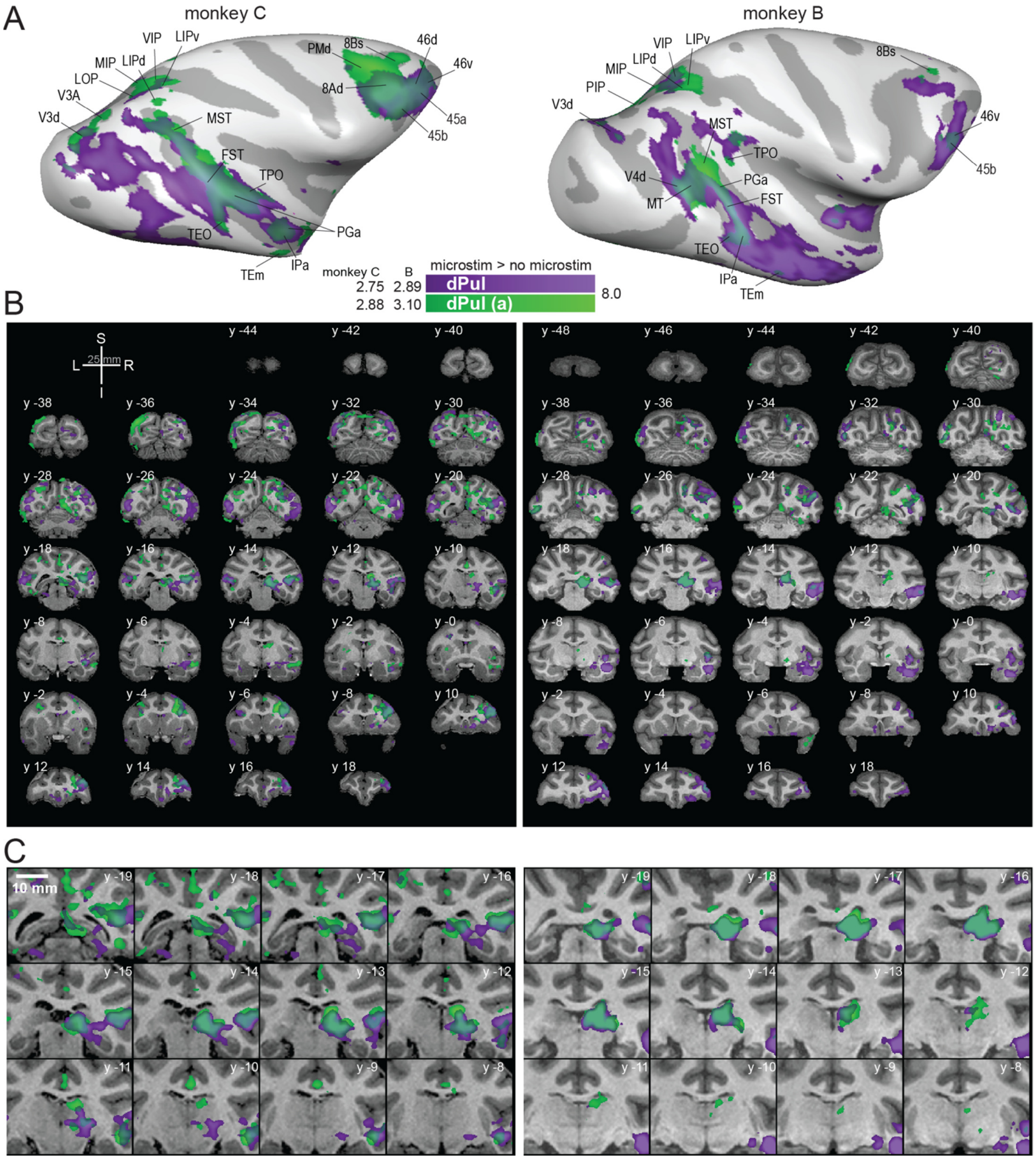

**Figure S19.** Comparison of dPul(a) and dPul stimulation effects. **(A)** Activation maps on the inflated cortical surface (stimulated hemisphere), lateral view. Cortical areas activated by dPul(a) stimulation are labeled. In both monkeys, distal activations due to dPul(a) and dPul stimulation coincided in V4d, V3A, V3d, V3v, V2, the dorsal bank and the fundus of STS (FST, IPa, PGa, TPO), ventral STS (MT, TEO, TEm), insula, parietal areas LIP, VIP, MIP, 7a, MST, and prefrontal areas 46v, 8Bs, 45A. **(B)** Same maps in the volume space, coronal sections (2 mm spacing). **(C)** Zoomed in sections through the thalamus (1 mm spacing). L – left, R – right, S – superior, I – inferior, y - distance from AC-PC origin along the anterior/posterior plane in mm.

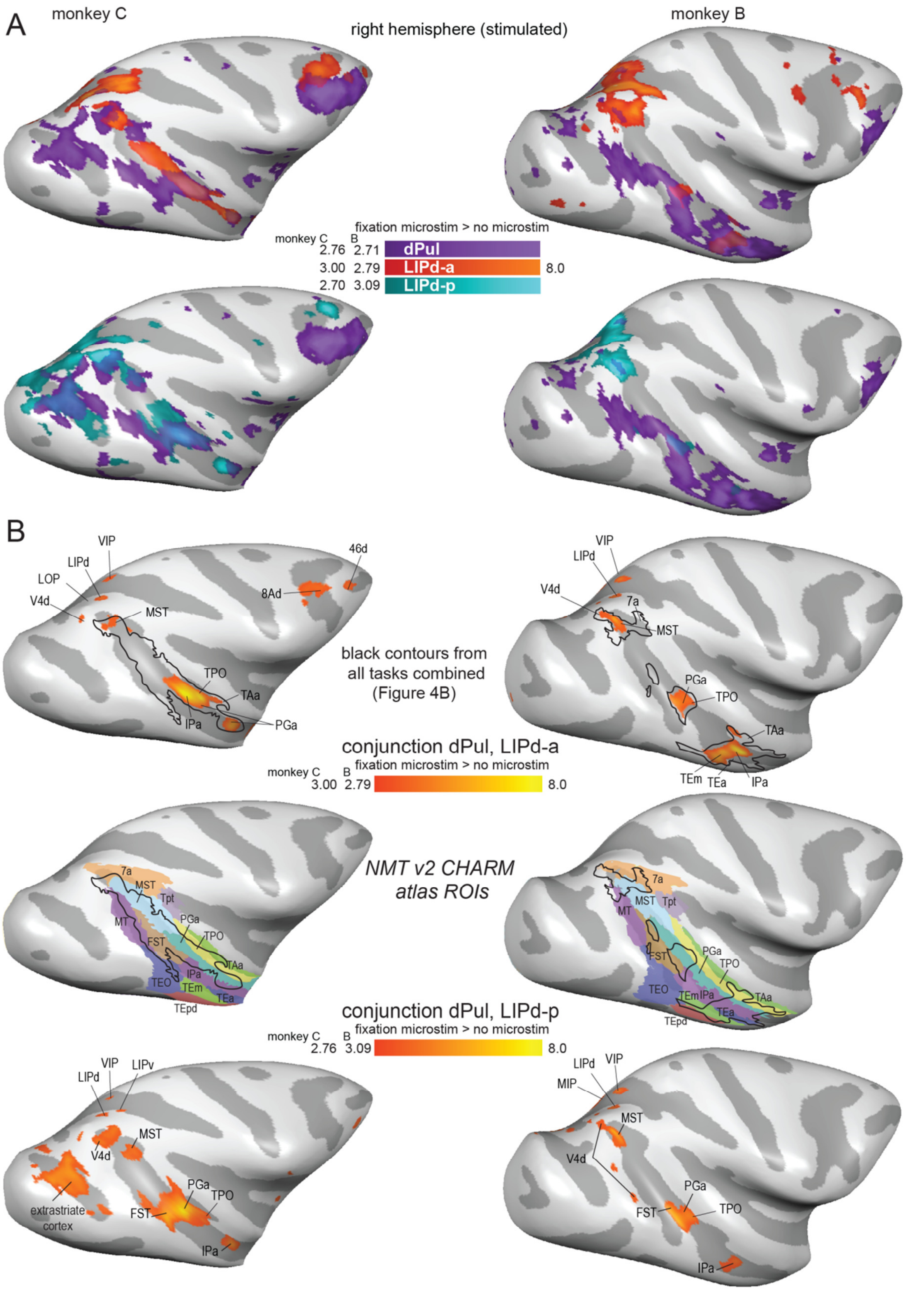

**Figure 20. Overlap of thalamic and parietal activation, fixation only task condition. (A)** Statistical t-maps showing BOLD activation on the inflated cortical lateral surface for dPul stimulation overlaid together with LIPd-a or dPul-LIPd-p stimulation. **(B)** Conjunction maps showing the overlap of activation elicited by the stimulation of dPul with LIPd-a (top row) and dPul with LIPd-p (bottom row). Major overlap regions are labeled. For reference, middle row shows the NMT v2 CHARM regions in STS, warped to the individual brain anatomy together with the combined contrast overlap contours from the **Figure 4B** (dPul & LIPd-a conjunction).

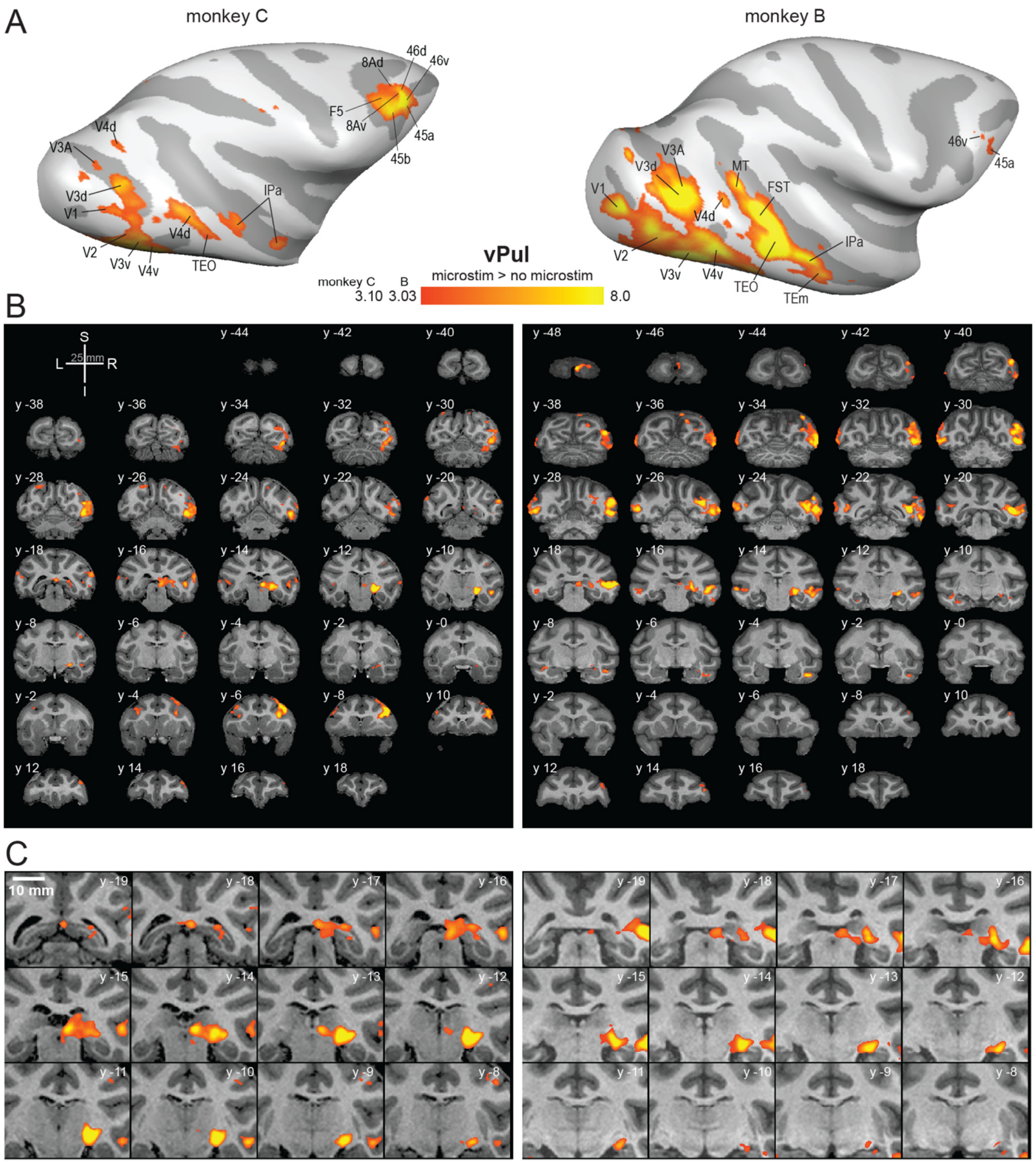

**Figure S21.** Stimulation effects in vPul (current strength 100 µA). **(A)** Activation maps on the inflated cortical surface, lateral view. **(B)** Same maps in the volume space, coronal sections (2 mm spacing). **(C)** Zoomed in sections through the thalamus (1 mm spacing). L – left, R – right, S – superior, I – inferior, y - distance from AC-PC origin along the anterior/posterior plane in mm.

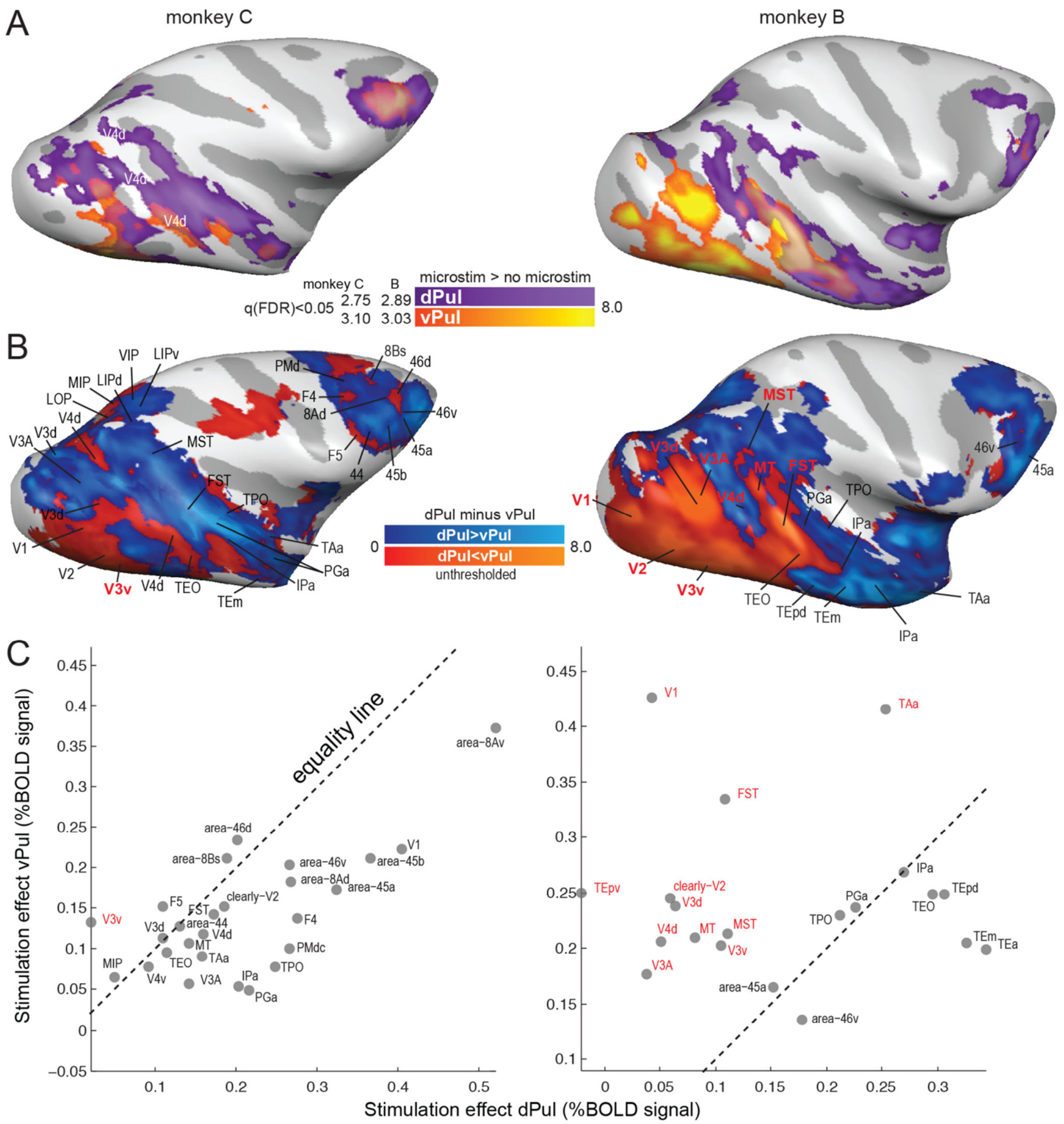

#### D ROIs activated both by dPul and vPul stimulation

| | |
|---|---|
| area-8Ad | IPa |
| area-8Av | PGa |
| area-8Bs | FST |
| area-46d | TPO |
| area-46v | TAa |
| area-45a | MT |
| area-45b | V4d |
| area-44 | V4v |
| PMdc | V3A |
| F4 | V3d |
| F5 | V3v |
| MIP | clearly-V2 |
| TEO | V1 |

Paired t-test dPul 0.21 vs. vPul 0.14, p<0.001

| | |
|---|---|
| area-46v | FST |
| area-45a | TPO |
| MST | TAa |
| TEO | MT |
| TEpv | V4d |
| TEpd | V3A |
| TEm | V3d |
| TEa | V3v |
| IPa | clearly-V2 |
| PGa | V1 |

Paired t-test dPul 0.16 vs. vPul 0.24, p=0.014

**Figure S22.** Comparison of stimulation effect strength in cortical areas elicited by dPul and vPul stimulation. **(A)** dPul and vPul activation maps on the inflated cortical surface (stimulated hemisphere), lateral view. Multiple V4d clusters are labeled. **(B)** Unthresholded difference of the dPul and vPul activation maps. Overlapping cortical activation ROIs are labeled, red color denotes ROIs that had stronger vPul stimulation effect derived from the ROI analysis, as seen in **(C)**. **(D)** The list of ROIs activated by stimulation of both sites, and results of paired t-test comparing stimulation effect strength elicited by dPul and vPul stimulation.

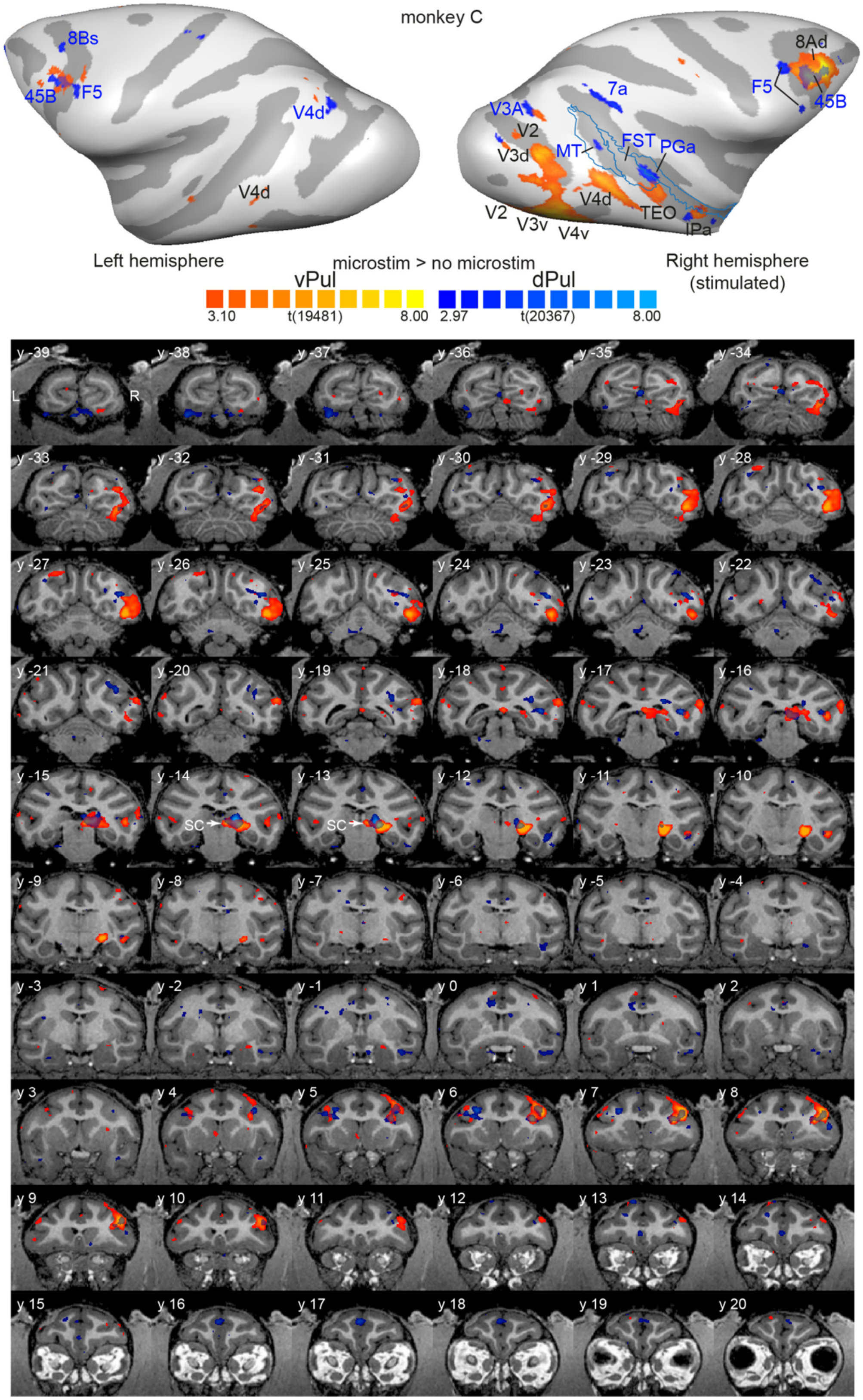

**Figure S23.** Comparison of low current 100 µA dPul and vPul stimulation effects, monkey C. **(A)** Activation maps on the inflated cortical surface, lateral view. Cortical regions activated by dPul stimulation are labeled in blue color, vPul – black. The CHARM template outlines of MT, FST and PGa areas shown as blue contours. **(B)** Same maps in the volume space, coronal sections (1 mm spacing). L – left, R – right, y - distance from AC-PC origin along the anterior/posterior plane in mm.

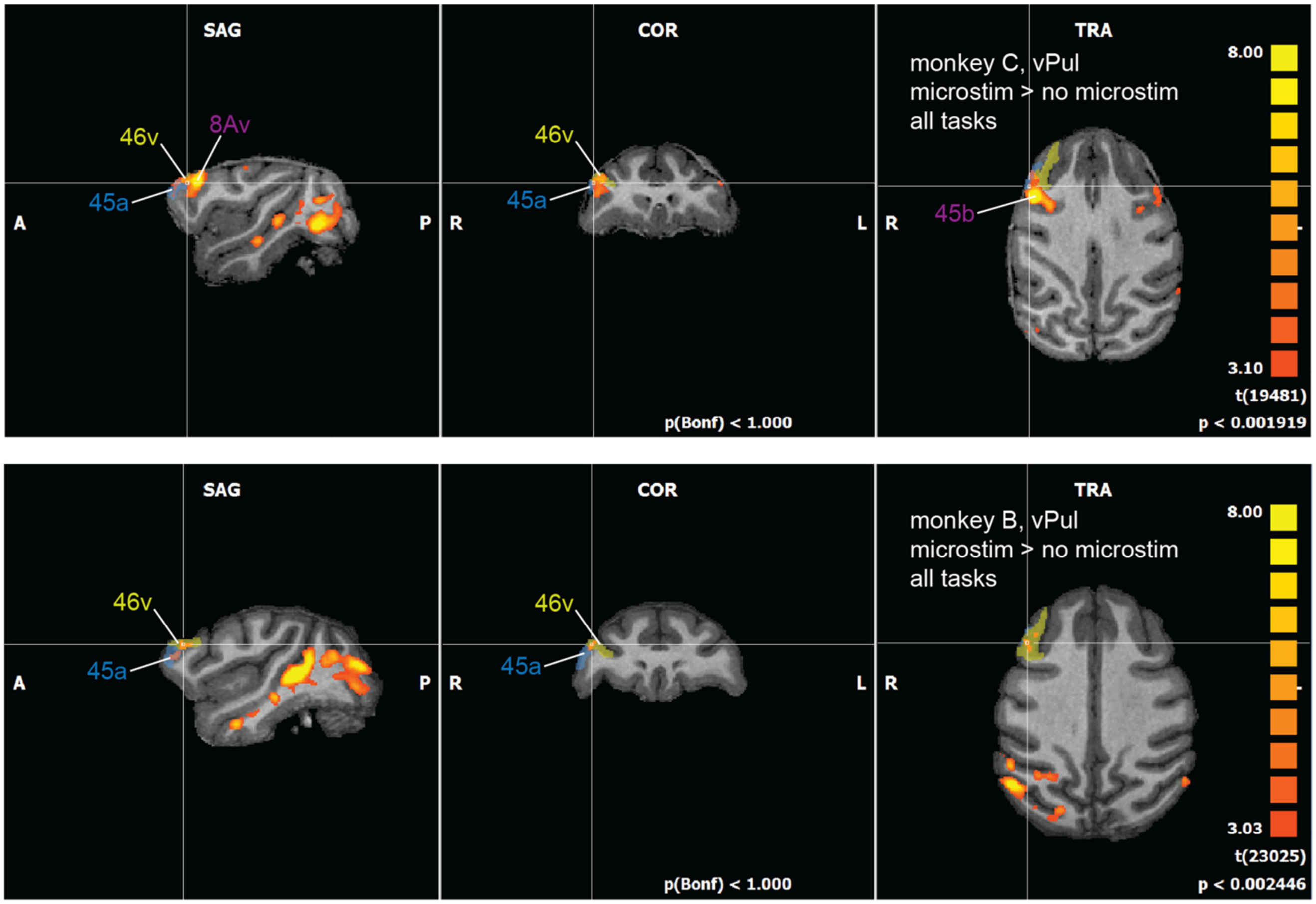

**Figure S24.** Activation of frontal regions by a low 100 µA current vPul stimulation. Coronal, axial and sagittal sections through the frontal activation locus are shown for each monkey (monkey C, top row, monkey B, bottom row), together with CHARM atlas areas 45a, 45b, and 46v.

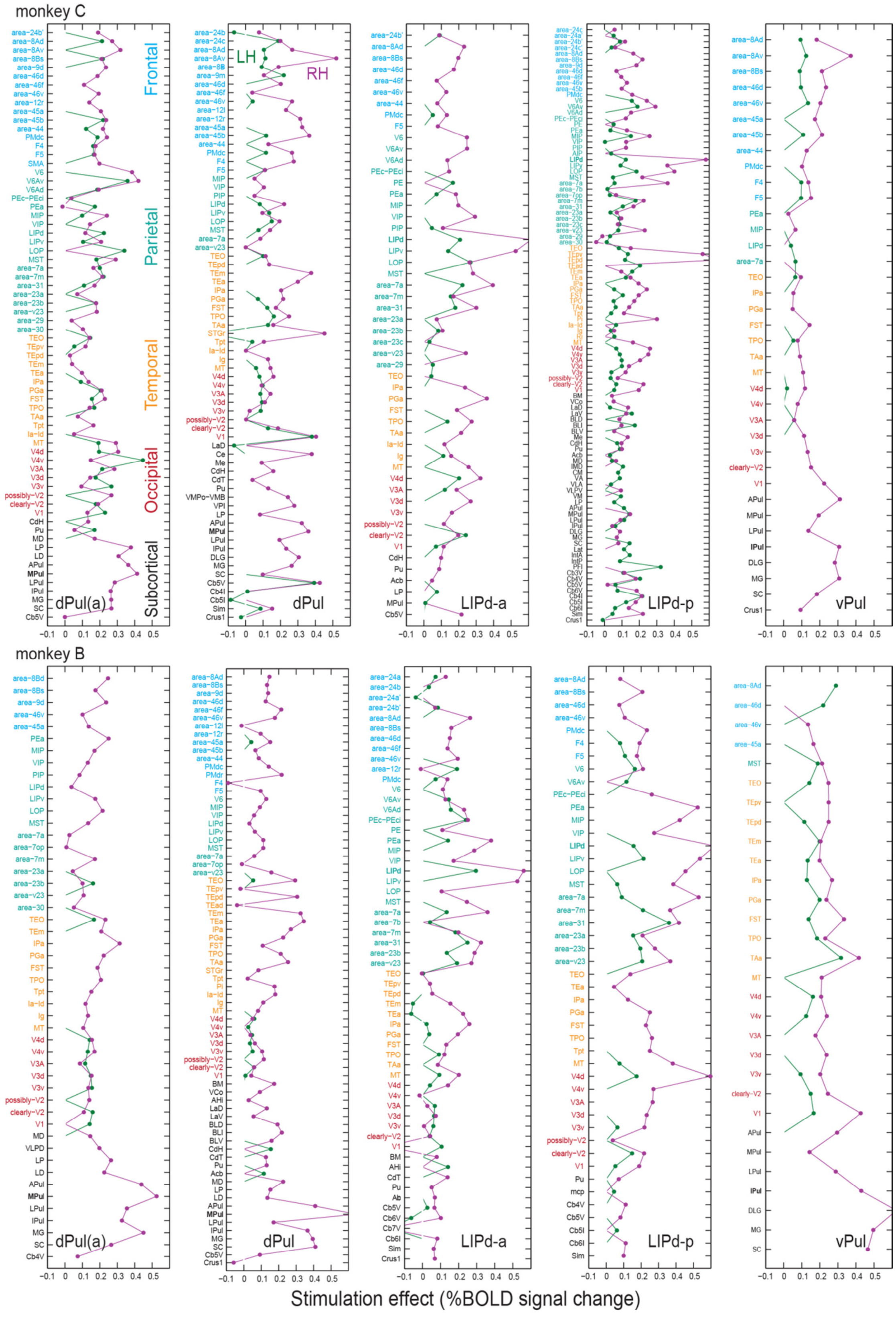

**Figure 25. Stimulation effect profiles derived from event-related average response amplitudes, for each stimulation site.** Magenta – stimulated right hemisphere (RH), green – left hemisphere (LH).

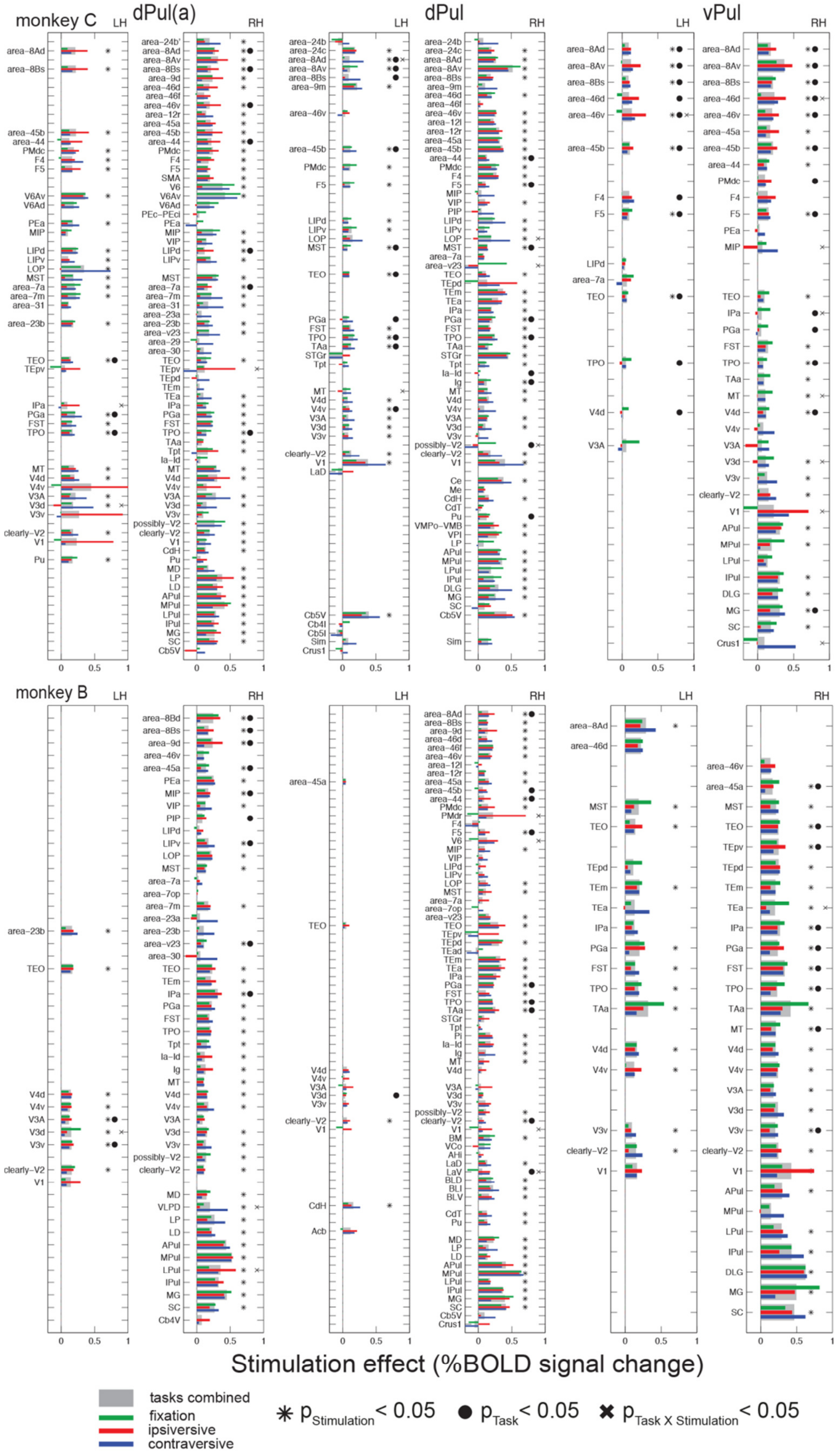

**Figure S26.** Stimulation effects derived from event-related average response amplitudes, pulvinar sites. For each ROI, the mean stimulation effect calculated as a difference between stimulation and no stimulation response amplitudes (gray bar) and task condition-specific stimulation effects (green, red and blue bars) are shown together with the results of two-way ANOVA across trials (asterisk –significant effect of stimulation, filled circle –significant effect of task, cross –significant interaction, p<0.05).

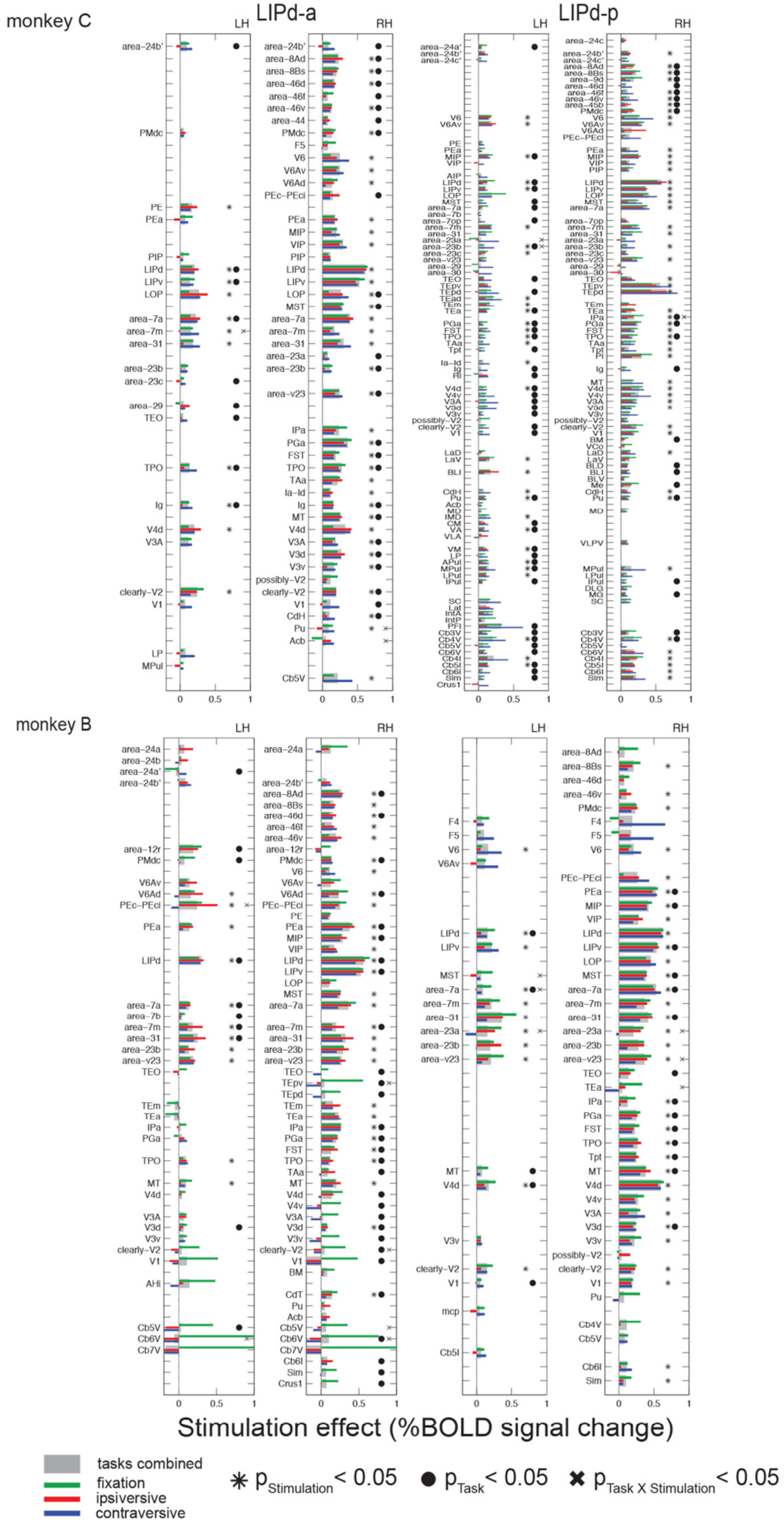

**Figure S27.** Stimulation effects derived from event-related average response amplitudes, LIPd sites. For each ROI, the mean stimulation effect calculated as a difference between stimulation and no stimulation response amplitudes (gray bar) and task condition-specific stimulation effects (green, red and blue bars) are shown together with the results of two-way ANOVA across trials (asterisk –significant effect of stimulation, filled circle –significant effect of task, cross –significant interaction, p<0.05).

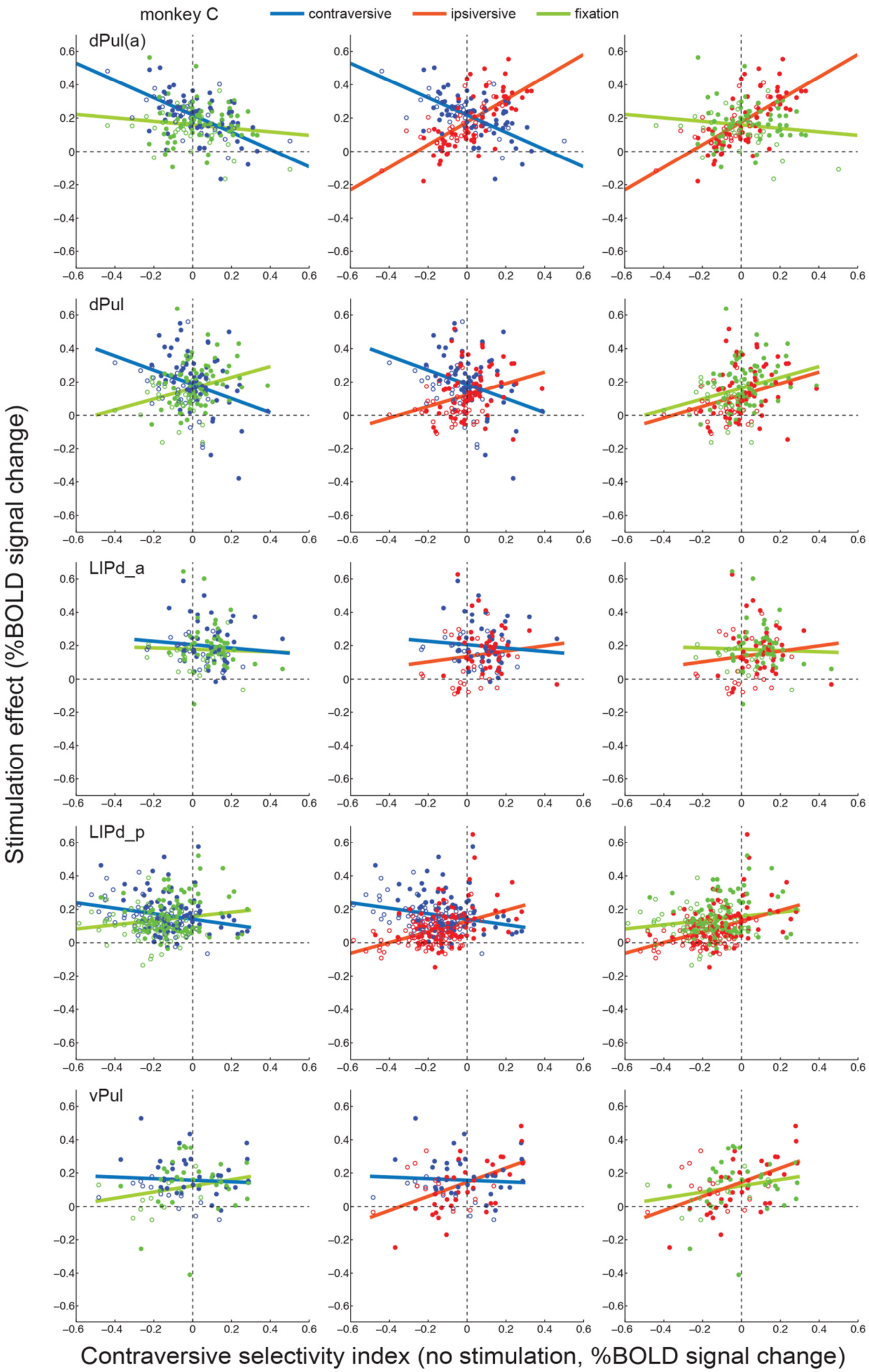

**Figure S28.** Microstimulation effect dependence on contraversive spatial selectivity and task. Monkey C. For each ROI, the effect of stimulation, defined as the difference between the BOLD response in stimulation and no stimulation trials, is plotted against its initial contraversive selectivity index (CSI in no stimulation trials) for contraversive memory saccade (blue), ipsiversive memory saccade (red), and fixation (green) task conditions. Each dot represents one ROI (filled circles – right hemisphere, empty circles – left hemisphere); solid lines show linear fits of stimulation effects across ROIs. See **Table 4** for the corresponding statistics.

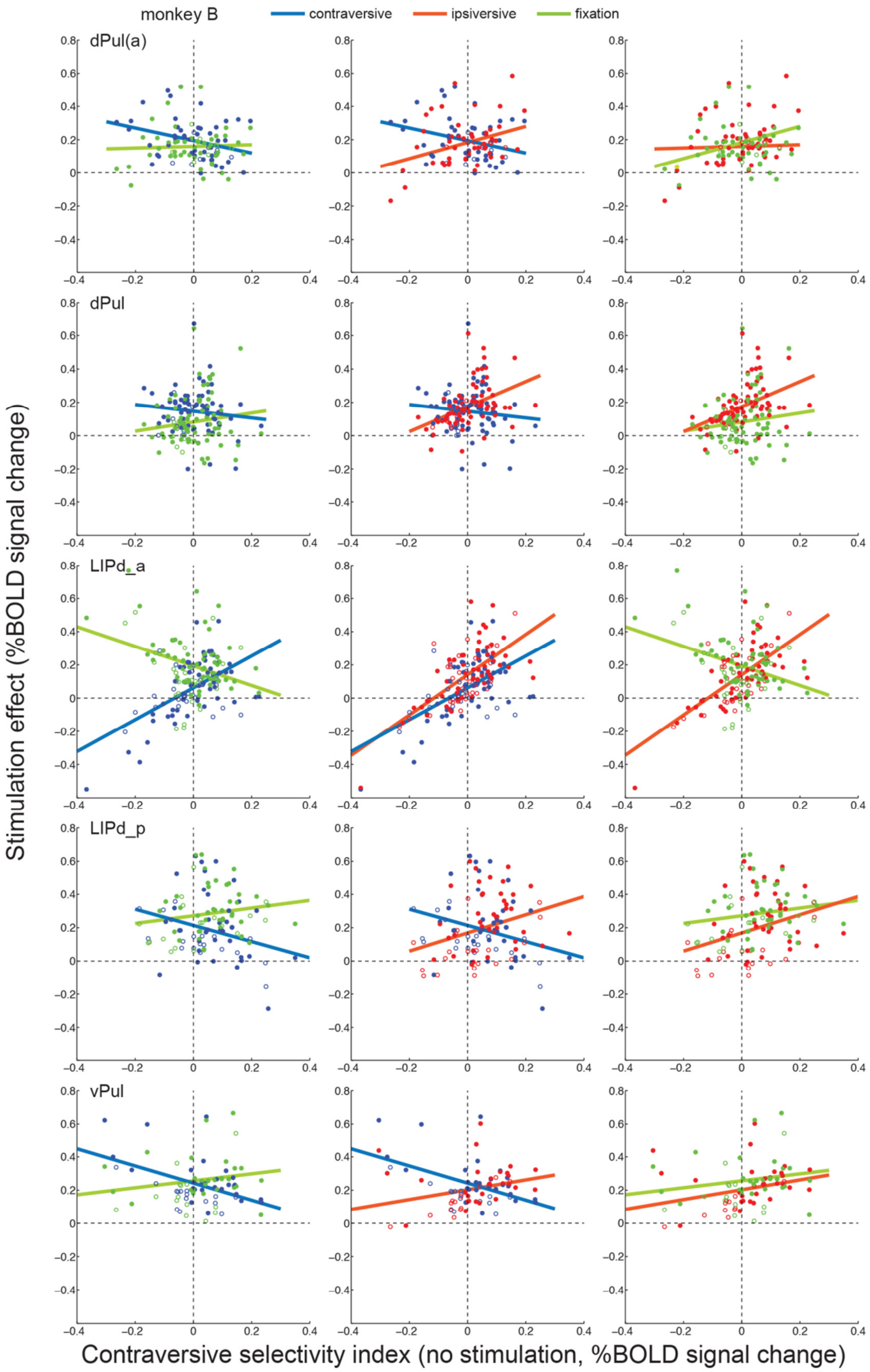

**Figure S29.** Microstimulation effect dependence on contraversive spatial selectivity and task. Monkey B. For each ROI, the effect of stimulation, defined as the difference between the BOLD response in stimulation and no stimulation trials, is plotted against its initial contraversive selectivity index (CSI in no stimulation trials) for contraversive memory saccade (blue), ipsiversive memory saccade (red), and fixation (green) task conditions. Each dot represents one ROI (filled circles – right hemisphere, empty circles – left hemisphere); solid lines show linear fits of stimulation effects across ROIs. See **Table 4** for the corresponding statistics.

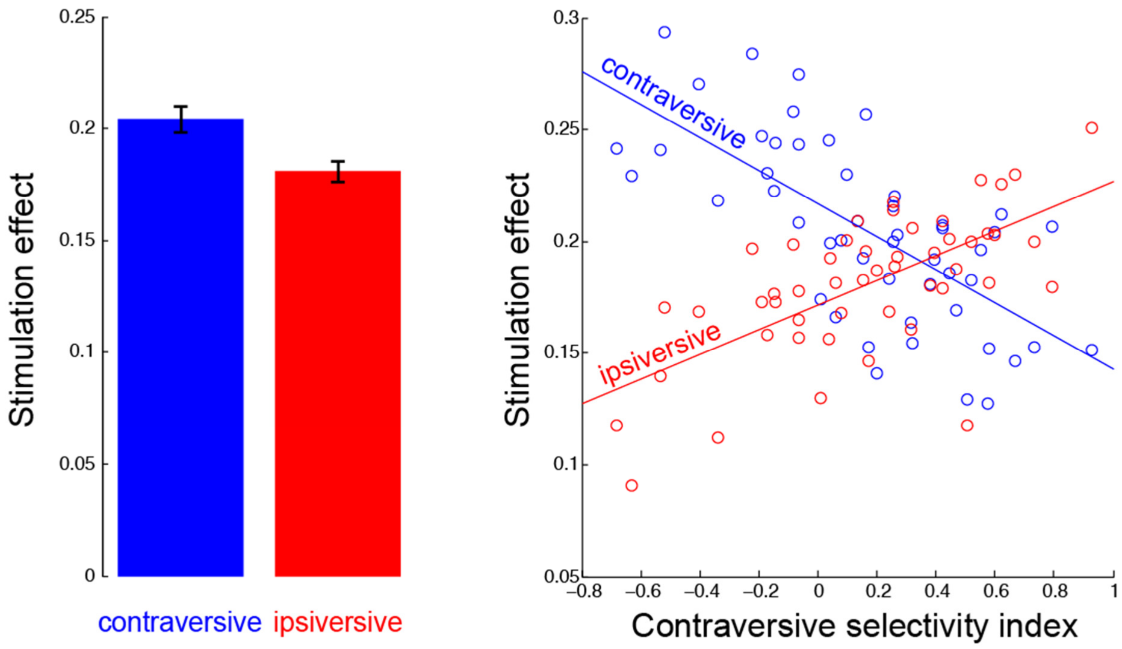

**Figure S29.** A simple qualitative model of additive microstimulation enhancement scaled down by the initial response amplitude reproduced the "X-shaped" interaction between the task condition-specific stimulation effect and contraversive selectivity, across ROIs. *Left panel:* average stimulation effect for contraversive and ipsiversive task conditions, across ROIs. *Right panel:* stimulation effect per ROI as a function of initial contraversive selectivity in control trials, for the two task conditions.

Simulation parameters:

```
RA_noise_level            = 0.25;
Stim_effect_contra        = 0.25;
Stim_effect_ipsi          = 0.2;
Stim_effect_noise_level   = 0.02;
RA_c_contra               = randn(50,1)*RA_noise_level+[0.4]; % control contraversive
RA_c_ipsi                 = randn(50,1)*RA_noise_level+[0.15]; % control ipsiversive
RA_s_contra = RA_c_contra + Stim_effect_contra*(1 - 0.5*RA_c_contra) + randn(50,1)*Stim_effect_noise_level;
RA_s_ipsi   = RA_s_ipsi   + Stim_effect_ipsi*(1 - 0.5*RA_c_ipsi)   + randn(50,1)*Stim_effect_noise_level;
```

### 1 Supplementary Results

#### 1.1 Task performance and eye movements

##### 1.1.1 Overall hit rate and trial aborts during and after stimulation

In monkey C, dPul(a) stimulation neither affected hit rate nor the number of trials aborted in and after the stimulation period, respectively (all $\chi^2$s(3) ≤ 0.83, all *p*s ≥ 0.3624). Similar results were found for dPul stimulation (all $\chi^2$s(3) ≤ 1.31, all *p*s ≥ 0.2523). In monkey B, overall hit rate and the number of trials aborted in the stimulation period were also not affected by dPul(a) stimulation (both $\chi^2$s(3) ≤ 2.494, all *p*s ≥ 0.1143). However, dPul(a) stimulation led to an overall increase in the number of trials aborted *after* the stimulation period ($\chi^2(3) = 15.642$, $p < 0.001$), which was mainly driven by an impairment in making saccades to cued locations in the contraversive (left) hemifield ($\chi^2(3) = 29.03$, $p < 0.001$); dPul stimulation did not have an effect on the number of trials aborted after the stimulation period ($\chi^2(3) = 1.48$, $p = 0.2240$). Overall hit rate was not affected by dPul stimulation ($\chi^2(3) = 2.21$, $p = 0.1370$) but there was a significant effect on the number of trials aborted in the stimulation period ($\chi^2(3) = 3.67$, $p < 0.05$) with a significantly decreased number of aborted trials in the contraversive memory saccade task ($\chi^2(3) = 5.37$, $p < 0.05$).

The stimulation in vPul did not lead to any changes in the overall hit rate or the number of trials aborted during or after the stimulation period in monkey C (although there was a weak tendency of increased hit rates in all task conditions), but in monkey B, both the hit rate overall increased ($\chi^2$s(3)=8.91, p=0.003) and the trials were aborted less frequently during and after the stimulation ($\chi^2$s(3)=4.30, p<0.05 and $\chi^2$s(3)=8.14, p<0.01), significant for the contraversive memory saccade trials after the stimulation.

LIPd-a stimulation did not have a significant effect on hit rate or the number of aborted trials (monkey C: all $\chi^2$s(3) ≤ 1.11, all *p*s ≥ 0.2930; monkey B: all $\chi^2$s(3) ≤ 3.63, all *p*s ≥ 0.0568). LIPd-p stimulation led to a significantly lower overall hit rate in monkey C ($\chi^2(3) = 5.03$, $p < 0.05$), which was mainly driven by a lower number of successful trials in the ipsiversive memory saccade task with stimulation compared to the control condition ($\chi^2(3) = 4.48$, $p < 0.05$). However, there was no significant increase in the number of trials aborted during or after the stimulation period ($\chi^2(3) = 0.59$, $p = 0.4416$ and 2(3) = 3.18, $p = 0.0747$, respectively). In monkey B, LIPd-p stimulation did not affect overall hit rate $\chi^2(3) = 1.84$, $p = 0.1753$) or the number of trials aborted in the stimulation period ($\chi^2(3) = 1.37$, $p = 0.2424$) but led to a significant decrease in the overall number of trials aborted after the stimulation period ($\chi^2(3) = 4.07$, $p < 0.05$). None of the comparisons between stimulation and control trials for each task separately reached significance (all $\chi^2$s(3) ≤ 2.70, all *p*s ≥ 0.1005).

**Figure S11** and **S12** summarize these data as bar plots.

#### 1.1.2 Frequency of eye movements

In brief, the effect on the frequency of eye movements was not consistent across datasets and animals: the stimulation of dPul(a) in monkey C and LIPd-p in monkey B led to a significant increase in the number of small eye movements during the fixation in stimulation period in all three tasks, but other datasets showed no change or a small increase and/or decrease in saccade frequency.

For dPul(a) dataset in monkey C, the two-way ANOVA on the number of eye movements in the stimulation period revealed a significant main effect of stimulation ($F(1, 526) = 139.41$, $p < 0.001$) with an increased number of eye movements in stimulation trials compared to the control condition in all three tasks (all *p*s < 0.001). In contrast, there were no significant effects of dPul stimulation on the number of eye movements in the stimulation period (main effect stimulation: $F(1, 823) = 0.07$, $p = 0.7881$, task × stimulation interaction: $F(2, 823) = 0.46$, $p = 0.6324$). In monkey B dPul(a) stimulation also significantly affected the number of eye movements in the stimulation period (main effect stimulation: $F(1, 1122) = 18.49$, task × stimulation interaction: $F(2, 1122) = 9.50$, both *p*s < 0.001), with a significantly lower number of eye movements in the fixation and the contraversive memory saccade task ($t(392) = 2.80$ and $t(364) = 5.09$, respectively, both *p*s < 0.001). dPul stimulation also significantly influenced the number of eye movements in the stimulation period as shown by a significant task × stimulation interaction effect ($F(2, 1263) = 14.71$, $p < 0.001$): there was a significantly higher number of eye movements in the ipsiversive memory saccade task ($t(424) = 4.13$, $p < 0.001$) whereas the number of eye movements was decreased by dPul stimulation in the contraversive memory saccade task ($t(412) = 3.15$, $p < 0.001$).

The stimulation in vPul did not lead to any changes in the number of eye movements in the stimulation period in monkey C, but in monkey B there was a main effect of the stimulation on decreasing the number of eye movements ($F(1, 983)=29.33$, $p<0.000$), significant also separately in all tasks (*p*s < 0.01).

The ANOVA on the number of eye movements in the stimulation period revealed a significant main effect of stimulation in LIPd-a in monkey C ($F(1, 873) = 10.74$, $p < 0.01$). Further post-hoc t tests showed that LIPd-a stimulation led to a significantly higher number of eye movements only in the contraversive memory saccade task ($t(318) = 3.09$, $p < 0.01$). LIPd-p stimulation did not affect the number of eye movements in the stimulation period (main effect stimulation: $F(1, 882) = 0.06$, $p = 0.8063$, task × stimulation interaction: $F(2, 882) = 1.73$, $p = 0.1773$). In monkey B, LIPd-a stimulation also affected the number of eye movements in the stimulation period as shown by a significant main effect of stimulation ($F(1, 1503) = 22.89$, $p < 0.001$), but in contrast to monkey C, it led to less eye movements with the difference between stimulation and control trials reaching significance in the fixation ($t(495) = 3.24$, $p < 0.01$) and the ipsiversive memory saccade task ($t(504) = 3.24$, $p < 0.01$). Stimulation in LIPd-p led to a higher number of eye movements in the stimulation period as shown by a significant main effect of stimulation ($F(1, 1278) = 41.94$, $p < 0.001$)

and significant differences between stimulation and control trials for all three tasks (fixation: $t(423) = 2.25$, $p < 0.05$; memory saccade right: $t(432) = 4.69$, $p < 0.001$; memory saccade left: $t(423) = 4.26$, $p < 0.001$).

**Figure S13** summarizes these data as bar plots.

##### 1.1.3 Saccade latencies

In monkey C, the two-way ANOVA on saccade latencies did not reveal significant effects of dPul(a) stimulation ($F(1, 336) = 0.16$, $p = 0.69$), but a significant effects of dPul stimulation ($F(1, 514)$s $\geq$ 8.04, $p$s < 0.001) with significantly longer latencies for contraversive saccades following stimulation compared to the control condition ($t(244) = 4.17$, $p < 0.001$). In monkey B for dPul(a) stimulation the two-way ANOVA on saccade latencies revealed a significant main effect of stimulation ($F(1, 727) = 15.70$, $p < 0.001$). Further post-hoc t tests showed that saccades to cued locations in both the ipsiversive and the contraversive hemifield were significantly delayed compared to the control conditions ($t(366) = 2.95$ and $t(361) = 2.68$, respectively, both $p$s < 0.01). Similar to dPul(a) stimulation, the two-way ANOVA revealed a significant main effect of dPul stimulation with longer latencies of saccades to both the contraversive and the ipsiversive hemifield ($F(1, 833) = 4.95$, $p < 0.05$). However, for neither of the saccade tasks the difference in saccade latencies between stimulation and control trials reached significance in post-hoc t tests ($t(423) = 1.60$ and $t(410) = 1.54$, both $p$s $\geq$ 0.1048).

The stimulation in vPul did not lead to any changes in the saccade latency in monkey C, but in monkey B there was a main effect of the stimulation on increasing the latency ($F(1, 651) = 8.12$, p < 0.01), significant also separately for the contraversive task ($p < 0.01$).

Saccade latencies were not affected by LIPd-a stimulation in monkey C (main effect stimulation: $F(1, 581) = 1.05$, $p = 0.3068$, task × stimulation interaction: $F(1, 581) = 0.10$, $p = 0.7557$), but the two-way ANOVA on saccade latencies revealed a significant main effect of LIPd-p stimulation ($F(1, 584) = 10.42$, $p < 0.01$) mainly driven by an increased latency due to stimulation in the contraversive memory saccade task ($t(290) = 3.03$, $p < 0.01$). Saccade latencies were also affected by LIPd-a stimulation in monkey B as shown by a significant main effect of stimulation ($F(1, 1004) = 11.11$, $p < 0.001$) and a significant task × stimulation interaction effect ($F(1, 1004) = 4.59$, $p < 0.05$). Post-hoc t tests showed that saccade latencies were significantly longer due to stimulation only in the contraversive memory saccade task ($t(502) = 3.66$, $p < 0.001$). LIPd-p stimulation did not have significant effects on saccade latencies (main effect stimulation: $F(1, 854) = 0.01$, $p = 0.9190$, task × stimulation interaction: $F(1, 854) = 0.36$, $p = 0.5493$).

**Figure S14** summarizes these data as bar plots.

**Table S1.** Electrode tip positions in the dPul pulvinar sites, relative to the chamber normal orientation (vertical z axis along the chamber and the grid), in mm. Horizontal x (negative left to positive right) and y (negative posterior to positive anterior) are relative to the chamber center: grid hole 0,0; z is relative to the brain surface entry point (negative – inside the brain). Mean and standard deviation (SD) are across sessions.

| Site | Estimate | x | y | z |
|---|---|---|---|---|
| Monkey C, dPul(a) | Mean | 2.60 | 3.10 | -19.00 |
| | SD | 0.22 | 0.52 | 1.17 |
| Monkey C, dPul | Mean | 3.75 | 2.32 | -19.79 |
| | SD | 0.00 | 0.47 | 0.49 |
| Monkey B, dPul(a) | Mean | 3.36 | 0.86 | -20.39 |
| | SD | 0.31 | 0.31 | 0.33 |
| Monkey B, dPul | Mean | 3.33 | 0.44 | -20.78 |
| | SD | 0.22 | 0.39 | 0.48 |

**Table S2.** Summary of selected regions of interest. For each monkey and stimulation site dataset, 0 and 1 denote absence/presence of significant stimulation effect in the ROI. Out of 79 pre-selected cortical atlas labels, microstimulation-elicited activation was not detected in 3 areas (area_8Bm, preSMA, TAav; gray rows, last column "FALSE"). Out of 70 pre-selected subcortical atlas labels, microstimulation-elicited activation was not detected in 22 areas (gray rows).

| | | | | | | Monkey C | | | | | | | | | | Monkey B | | | | | | | | | | |
|---|---|---|---|---|---|---|---|---|---|---|---|---|---|---|---|---|---|---|---|---|---|---|---|---|---|---|
| ROI number | Index | Abbreviation | Lobe | Full_Name | Atlas | dPul(a) LH | dPul(a) RH | dPul LH | dPul RH | LIPd-a LH | LIPd-a RH | LIPd-p LH | LIPd-p RH | vPul LH | vPul RH | dPul(a) LH | dPul(a) RH | dPul LH | dPul RH | LIPd-a LH | LIPd-a RH | LIPd-p LH | LIPd-p RH | vPul LH | vPul RH | Exists once or more |
| 1 | 8 | area_24a | frontal | area_24a | CHARM | 0 | 0 | 0 | 0 | 0 | 0 | 0 | 0 | 0 | 0 | 0 | 0 | 0 | 0 | 1 | 1 | 0 | 0 | 0 | 0 | TRUE |
| 2 | 9 | area_24b | frontal | area_24b | CHARM | 0 | 0 | 1 | 1 | 0 | 0 | 0 | 0 | 0 | 0 | 0 | 0 | 0 | 0 | 1 | 0 | 0 | 0 | 0 | 0 | TRUE |
| 3 | 10 | area_24c | frontal | area_24c | CHARM | 0 | 0 | 1 | 1 | 0 | 0 | 0 | 1 | 0 | 0 | 0 | 0 | 0 | 0 | 0 | 0 | 0 | 0 | 0 | 0 | TRUE |
| 4 | 13 | area_24a' | frontal | area_24a' | CHARM | 0 | 0 | 0 | 0 | 0 | 0 | 1 | 0 | 0 | 0 | 0 | 0 | 0 | 0 | 1 | 0 | 0 | 0 | 0 | 0 | TRUE |
| 5 | 14 | area_24b' | frontal | area_24b' | CHARM | 0 | 1 | 0 | 0 | 1 | 1 | 1 | 1 | 0 | 0 | 0 | 0 | 0 | 0 | 1 | 1 | 0 | 0 | 0 | 0 | TRUE |
| 6 | 15 | area_24c' | frontal | area_24c' | CHARM | 0 | 0 | 0 | 0 | 0 | 0 | 1 | 1 | 0 | 0 | 0 | 0 | 0 | 0 | 0 | 0 | 0 | 0 | 0 | 0 | TRUE |
| 7 | 52 | area_8Ad | frontal | dorsal_periarcuate_area_8A | CHARM | 1 | 1 | 1 | 1 | 0 | 1 | 0 | 1 | 1 | 1 | 0 | 0 | 0 | 1 | 0 | 1 | 0 | 1 | 1 | 0 | TRUE |
| 8 | 53 | area_8Av | frontal | ventral_periarcuate_area_8A | CHARM | 0 | 1 | 1 | 1 | 0 | 0 | 0 | 0 | 1 | 1 | 0 | 0 | 0 | 0 | 0 | 0 | 0 | 0 | 0 | 0 | TRUE |
| 9 | 56 | area_8Bd | frontal | dorsal_area_8B | CHARM | 0 | 0 | 0 | 0 | 0 | 0 | 0 | 0 | 0 | 0 | 0 | 1 | 0 | 0 | 0 | 0 | 0 | 0 | 0 | 0 | TRUE |
| 10 | 57 | area_8Bm | frontal | medial_area_8B | CHARM | 0 | 0 | 0 | 0 | 0 | 0 | 0 | 0 | 0 | 0 | 0 | 0 | 0 | 0 | 0 | 0 | 0 | 0 | 0 | 0 | FALSE |
| 11 | 58 | area_8Bs | frontal | arcuate_sulcus_area_8B | CHARM | 1 | 1 | 1 | 1 | 0 | 1 | 0 | 1 | 1 | 1 | 0 | 1 | 0 | 1 | 0 | 1 | 0 | 1 | 0 | 0 | TRUE |
| 12 | 61 | area_9d | frontal | dorsal_area_9 | CHARM | 0 | 1 | 0 | 0 | 0 | 0 | 0 | 1 | 0 | 0 | 0 | 1 | 0 | 1 | 0 | 0 | 0 | 0 | 0 | 0 | TRUE |
| 13 | 62 | area_9m | frontal | medial_area_9 | CHARM | 0 | 0 | 1 | 1 | 0 | 0 | 0 | 0 | 0 | 0 | 0 | 0 | 0 | 0 | 0 | 0 | 0 | 0 | 0 | 0 | TRUE |
| 14 | 63 | area_46d | frontal | dorsal_area_46 | CHARM | 0 | 1 | 0 | 1 | 0 | 1 | 0 | 1 | 1 | 1 | 0 | 0 | 0 | 1 | 0 | 1 | 0 | 1 | 1 | 0 | TRUE |
| 15 | 67 | area_46f | frontal | area_46_in_fundus_of_the_principal_sulcus | CHARM | 0 | 1 | 0 | 1 | 0 | 1 | 0 | 1 | 0 | 0 | 0 | 0 | 0 | 1 | 0 | 1 | 0 | 0 | 0 | 0 | TRUE |
| 16 | 68 | area_46v | frontal | ventral_area_46 | CHARM | 0 | 1 | 1 | 1 | 0 | 1 | 0 | 1 | 1 | 1 | 0 | 1 | 0 | 1 | 0 | 1 | 0 | 1 | 0 | 1 | TRUE |
| 17 | 70 | area_12l | frontal | lateral_area_12 | CHARM | 0 | 0 | 0 | 1 | 0 | 0 | 0 | 0 | 0 | 0 | 0 | 0 | 0 | 1 | 0 | 0 | 0 | 0 | 0 | 0 | TRUE |
| 18 | 71 | area_12r | frontal | rostral_area_12 | CHARM | 0 | 1 | 0 | 1 | 0 | 0 | 0 | 0 | 0 | 0 | 0 | 0 | 0 | 1 | 1 | 1 | 0 | 0 | 0 | 0 | TRUE |
| 19 | 74 | area_45a | frontal | area_45a | CHARM | 0 | 1 | 0 | 1 | 0 | 0 | 0 | 0 | 0 | 1 | 0 | 1 | 1 | 1 | 0 | 0 | 0 | 0 | 0 | 1 | TRUE |
| 20 | 75 | area_45b | frontal | area_45b | CHARM | 1 | 1 | 1 | 1 | 0 | 0 | 0 | 1 | 1 | 1 | 0 | 0 | 0 | 1 | 0 | 0 | 0 | 0 | 0 | 0 | TRUE |
| 21 | 76 | area_44 | frontal | area_44 | CHARM | 1 | 1 | 0 | 1 | 0 | 1 | 0 | 0 | 0 | 1 | 0 | 0 | 0 | 1 | 0 | 0 | 0 | 0 | 0 | 0 | TRUE |
| 22 | 82 | PMdc | frontal | caudal_dorsal_premotor_cortex | CHARM | 1 | 1 | 1 | 1 | 1 | 1 | 0 | 1 | 0 | 1 | 0 | 0 | 0 | 1 | 1 | 1 | 0 | 1 | 0 | 0 | TRUE |
| 23 | 83 | PMdr | frontal | rostral_dorsal_premotor_cortex | CHARM | 0 | 0 | 0 | 0 | 0 | 0 | 0 | 0 | 0 | 0 | 0 | 0 | 0 | 1 | 0 | 0 | 0 | 0 | 0 | 0 | TRUE |
| 24 | 85 | F4 | frontal | area_F4_of_ventral_premotor_cortex | CHARM | 1 | 1 | 0 | 1 | 0 | 0 | 0 | 0 | 1 | 1 | 0 | 0 | 0 | 1 | 0 | 0 | 1 | 1 | 0 | 0 | TRUE |
| 25 | 86 | F5 | frontal | area_F5_of_ventral_premotor_cortex | CHARM | 1 | 1 | 1 | 1 | 0 | 1 | 0 | 0 | 1 | 1 | 0 | 0 | 0 | 1 | 0 | 0 | 1 | 1 | 0 | 0 | TRUE |
| 26 | 88 | preSMA | frontal | presupplementary_motor_area | CHARM | 0 | 0 | 0 | 0 | 0 | 0 | 0 | 0 | 0 | 0 | 0 | 0 | 0 | 0 | 0 | 0 | 0 | 0 | 0 | 0 | FALSE |
| 27 | 89 | SMA | frontal | supplementary_motor_area | CHARM | 0 | 1 | 0 | 0 | 0 | 0 | 0 | 0 | 0 | 0 | 0 | 0 | 0 | 0 | 0 | 0 | 0 | 0 | 0 | 0 | TRUE |
| 28 | 98 | V6 | parietal | visual_area_V6 | CHARM | 0 | 1 | 0 | 0 | 0 | 1 | 1 | 1 | 0 | 0 | 0 | 0 | 0 | 1 | 0 | 1 | 1 | 1 | 0 | 0 | TRUE |
| 29 | 100 | V6Av | parietal | ventral_visual_area_6A | CHARM | 1 | 1 | 0 | 0 | 0 | 1 | 1 | 1 | 0 | 0 | 0 | 0 | 0 | 0 | 1 | 1 | 1 | 0 | 0 | 0 | TRUE |
| 30 | 101 | V6Ad | parietal | dorsal_visual_area_6A | CHARM | 1 | 1 | 0 | 0 | 0 | 1 | 0 | 1 | 0 | 0 | 0 | 0 | 0 | 0 | 1 | 1 | 0 | 0 | 0 | 0 | TRUE |
| 31 | 104 | PEc-PEci | parietal | areas_PEc_and_PEci | CHARM | 0 | 1 | 0 | 0 | 0 | 1 | 0 | 1 | 0 | 0 | 0 | 0 | 0 | 0 | 1 | 1 | 0 | 1 | 0 | 0 | TRUE |
| 32 | 105 | PE | parietal | area_PE | CHARM | 0 | 0 | 0 | 0 | 1 | 0 | 1 | 0 | 0 | 0 | 0 | 0 | 0 | 0 | 0 | 1 | 0 | 0 | 0 | 0 | TRUE |
| 33 | 106 | PEa | parietal | area_PEa | CHARM | 1 | 1 | 0 | 0 | 1 | 1 | 1 | 1 | 0 | 1 | 0 | 1 | 0 | 0 | 1 | 1 | 0 | 1 | 0 | 0 | TRUE |
| 34 | 108 | MIP | parietal | medial_intraparietal_area | CHARM | 1 | 1 | 0 | 1 | 0 | 1 | 1 | 1 | 0 | 1 | 0 | 1 | 0 | 1 | 0 | 1 | 0 | 1 | 0 | 0 | TRUE |
| 35 | 110 | VIP | parietal | ventral_intraparietal_area | CHARM | 0 | 1 | 0 | 1 | 0 | 1 | 1 | 1 | 0 | 0 | 0 | 1 | 0 | 1 | 0 | 1 | 0 | 1 | 0 | 0 | TRUE |
| 36 | 111 | PIP | parietal | posterior_intraparietal_area | CHARM | 0 | 0 | 0 | 1 | 1 | 1 | 0 | 1 | 0 | 0 | 0 | 1 | 0 | 0 | 0 | 0 | 0 | 0 | 0 | 0 | TRUE |
| 37 | 114 | AIP | parietal | anterior_intraparietal_area | CHARM | 0 | 0 | 0 | 0 | 0 | 0 | 1 | 0 | 0 | 0 | 0 | 0 | 0 | 0 | 0 | 0 | 0 | 0 | 0 | 0 | TRUE |

*Table S2, continued*

| | | | | | | | | | | | | | | | | | | | | | | | | | | |
|---|---|---|---|---|---|---|---|---|---|---|---|---|---|---|---|---|---|---|---|---|---|---|---|---|---|---|
| 38 | 116 | LIPd | parietal | dorsal_lateral_intraparietal_area | CHARM | 1 | 1 | 1 | 1 | 1 | 1 | 1 | 1 | 1 | 0 | 0 | 1 | 0 | 1 | 1 | 1 | 1 | 1 | 0 | 0 | TRUE |
| 39 | 117 | LIPv | parietal | ventral_lateral_intraparietal_area | CHARM | 1 | 1 | 1 | 1 | 1 | 1 | 1 | 1 | 0 | 0 | 0 | 1 | 0 | 1 | 0 | 1 | 1 | 1 | 0 | 0 | TRUE |
| 40 | 118 | LOP | parietal | lateral_occipital_parietal_area | CHARM | 1 | 0 | 1 | 1 | 1 | 1 | 1 | 1 | 0 | 0 | 0 | 1 | 0 | 1 | 0 | 1 | 0 | 1 | 0 | 0 | TRUE |
| 41 | 119 | MST | parietal | medial_superior_temporal_area | CHARM | 1 | 1 | 1 | 1 | 0 | 1 | 1 | 1 | 0 | 0 | 0 | 1 | 0 | 1 | 0 | 1 | 1 | 1 | 1 | 1 | TRUE |
| 42 | 122 | area_7a | parietal | caudal_inferior_parietal_lobule_area_7a_(Opt/PG) | CHARM | 1 | 1 | 0 | 1 | 1 | 1 | 1 | 1 | 1 | 0 | 0 | 1 | 0 | 1 | 1 | 1 | 1 | 1 | 0 | 0 | TRUE |
| 43 | 123 | area_7b | parietal | rostral_inferior_parietal_lobule_area_7b_(PFG/PF) | CHARM | 0 | 0 | 0 | 0 | 0 | 0 | 1 | 0 | 0 | 0 | 0 | 0 | 0 | 0 | 1 | 0 | 0 | 0 | 0 | 0 | TRUE |
| 44 | 124 | area_7op | parietal | parietal_operculum | CHARM | 0 | 0 | 0 | 0 | 0 | 0 | 1 | 1 | 0 | 0 | 0 | 1 | 0 | 1 | 0 | 0 | 0 | 0 | 0 | 0 | TRUE |
| 45 | 126 | area_7m | parietal | area_7_(PGm)_on_the_medial_wall | CHARM | 1 | 1 | 0 | 0 | 1 | 1 | 1 | 1 | 0 | 0 | 0 | 1 | 0 | 0 | 1 | 1 | 1 | 1 | 0 | 0 | TRUE |
| 46 | 129 | area_31 | parietal | area_31 | CHARM | 1 | 1 | 0 | 0 | 1 | 1 | 1 | 1 | 0 | 0 | 0 | 0 | 0 | 0 | 1 | 1 | 1 | 1 | 0 | 0 | TRUE |
| 47 | 131 | area_23a | parietal | area_23a | CHARM | 0 | 1 | 0 | 0 | 0 | 1 | 1 | 1 | 0 | 0 | 0 | 1 | 0 | 0 | 0 | 0 | 1 | 1 | 0 | 0 | TRUE |
| 48 | 132 | area_23b | parietal | area_23b | CHARM | 1 | 1 | 0 | 0 | 1 | 1 | 1 | 1 | 0 | 0 | 1 | 1 | 0 | 0 | 1 | 1 | 1 | 1 | 0 | 0 | TRUE |
| 49 | 133 | area_23c | parietal | area_23c | CHARM | 0 | 0 | 0 | 0 | 1 | 0 | 1 | 1 | 0 | 0 | 0 | 0 | 0 | 0 | 0 | 0 | 0 | 0 | 0 | 0 | TRUE |
| 50 | 134 | area_v23 | parietal | area_v23 | CHARM | 0 | 1 | 0 | 1 | 0 | 1 | 1 | 1 | 0 | 0 | 0 | 1 | 0 | 1 | 1 | 1 | 1 | 1 | 0 | 0 | TRUE |
| 51 | 136 | area_29 | parietal | area_29 | CHARM | 0 | 1 | 0 | 0 | 1 | 0 | 1 | 1 | 0 | 0 | 0 | 0 | 0 | 0 | 0 | 0 | 0 | 0 | 0 | 0 | TRUE |
| 52 | 137 | area_30 | parietal | area_30 | CHARM | 0 | 1 | 0 | 0 | 0 | 0 | 1 | 1 | 0 | 0 | 0 | 1 | 0 | 0 | 0 | 0 | 0 | 0 | 0 | 0 | TRUE |
| 53 | 176 | TEO | temporal | area_TEO | CHARM | 1 | 1 | 1 | 1 | 1 | 0 | 1 | 1 | 1 | 1 | 1 | 1 | 1 | 1 | 1 | 1 | 0 | 1 | 1 | 1 | TRUE |
| 54 | 180 | TEpv | temporal | posterior_ventral_area_TE | CHARM | 1 | 1 | 0 | 0 | 0 | 0 | 1 | 1 | 0 | 0 | 0 | 0 | 0 | 1 | 0 | 1 | 0 | 0 | 0 | 1 | TRUE |
| 55 | 181 | TEpd | temporal | posterior_dorsal_area_TE | CHARM | 0 | 1 | 0 | 1 | 0 | 0 | 1 | 1 | 0 | 0 | 0 | 0 | 0 | 1 | 0 | 1 | 0 | 0 | 1 | 1 | TRUE |
| 56 | 183 | TEav | temporal | anterior_ventral_area_TE | CHARM | 0 | 0 | 0 | 0 | 0 | 0 | 0 | 0 | 0 | 0 | 0 | 0 | 0 | 0 | 0 | 0 | 0 | 0 | 0 | 0 | FALSE |
| 57 | 184 | TEad | temporal | anterior_dorsal_area_TE | CHARM | 0 | 0 | 0 | 0 | 0 | 0 | 1 | 0 | 0 | 0 | 0 | 0 | 0 | 1 | 0 | 0 | 0 | 0 | 0 | 0 | TRUE |
| 58 | 186 | TEm | temporal | area_TEm | CHARM | 0 | 1 | 0 | 1 | 0 | 0 | 1 | 1 | 0 | 0 | 0 | 1 | 0 | 1 | 1 | 1 | 0 | 0 | 1 | 1 | TRUE |
| 59 | 187 | TEa | temporal | area_TEa | CHARM | 0 | 1 | 0 | 1 | 0 | 0 | 1 | 1 | 0 | 0 | 0 | 0 | 0 | 1 | 1 | 1 | 0 | 1 | 1 | 1 | TRUE |
| 60 | 190 | IPa | temporal | area_IPa | CHARM | 1 | 1 | 0 | 1 | 0 | 1 | 0 | 1 | 0 | 1 | 0 | 1 | 0 | 1 | 1 | 1 | 0 | 1 | 1 | 1 | TRUE |
| 61 | 191 | PGa | temporal | area_PGa | CHARM | 1 | 1 | 1 | 1 | 0 | 1 | 1 | 1 | 0 | 1 | 0 | 1 | 0 | 1 | 1 | 1 | 0 | 1 | 1 | 1 | TRUE |
| 62 | 192 | FST | temporal | floor_of_the_superior_temporal_area | CHARM | 1 | 1 | 1 | 1 | 0 | 1 | 1 | 1 | 0 | 1 | 0 | 1 | 0 | 1 | 0 | 1 | 0 | 1 | 1 | 1 | TRUE |
| 63 | 196 | TPO | temporal | temporal_parietooccipital_associated_area | CHARM | 1 | 1 | 1 | 1 | 1 | 1 | 1 | 1 | 1 | 1 | 0 | 1 | 0 | 1 | 1 | 1 | 0 | 1 | 1 | 1 | TRUE |
| 64 | 197 | TAa | temporal | area_TAa | CHARM | 0 | 1 | 1 | 1 | 0 | 1 | 1 | 1 | 0 | 1 | 0 | 0 | 0 | 1 | 0 | 1 | 0 | 0 | 1 | 1 | TRUE |
| 65 | 198 | STGr | temporal | rostral_superior_temporal_gyrus | CHARM | 0 | 0 | 1 | 1 | 0 | 0 | 0 | 0 | 0 | 0 | 0 | 0 | 0 | 1 | 0 | 0 | 0 | 0 | 0 | 0 | TRUE |
| 66 | 200 | Tpt | temporal | temporo-parietal_area | CHARM | 0 | 1 | 1 | 1 | 0 | 0 | 1 | 1 | 0 | 0 | 0 | 1 | 0 | 1 | 0 | 0 | 0 | 1 | 0 | 0 | TRUE |
| 67 | 226 | Pi | temporal | parainsula | CHARM | 0 | 0 | 0 | 0 | 0 | 0 | 0 | 1 | 0 | 0 | 0 | 0 | 0 | 1 | 0 | 0 | 0 | 0 | 0 | 0 | TRUE |
| 68 | 228 | Ia-Id | temporal | agranular_and_dysgranular_insula | CHARM | 0 | 1 | 0 | 1 | 0 | 1 | 1 | 0 | 0 | 0 | 0 | 1 | 0 | 1 | 0 | 0 | 0 | 0 | 0 | 0 | TRUE |
| 69 | 229 | Ig | temporal | granular_insula | CHARM | 0 | 0 | 0 | 1 | 1 | 1 | 1 | 1 | 0 | 0 | 0 | 1 | 0 | 1 | 0 | 0 | 0 | 0 | 0 | 0 | TRUE |
| 70 | 230 | Ri | temporal | retroinsula | CHARM | 0 | 0 | 0 | 0 | 0 | 0 | 1 | 0 | 0 | 0 | 0 | 0 | 0 | 0 | 0 | 0 | 0 | 0 | 0 | 0 | TRUE |
| 71 | 232 | MT | temporal | middle_temporal_area | CHARM | 1 | 1 | 1 | 1 | 0 | 1 | 0 | 1 | 0 | 1 | 0 | 1 | 0 | 1 | 1 | 1 | 1 | 1 | 0 | 1 | TRUE |
| 72 | 235 | V4d | occipital | dorsal_visual_area_4 | CHARM | 1 | 1 | 1 | 1 | 1 | 1 | 1 | 1 | 1 | 1 | 1 | 1 | 1 | 1 | 1 | 1 | 1 | 1 | 1 | 1 | TRUE |
| 73 | 236 | V4v | occipital | ventral_visual_area_4 | CHARM | 1 | 1 | 1 | 1 | 0 | 0 | 1 | 1 | 0 | 1 | 1 | 1 | 1 | 0 | 0 | 1 | 0 | 1 | 1 | 1 | TRUE |
| 74 | 240 | V3A | occipital | visual_area_V3A | CHARM | 1 | 1 | 1 | 1 | 1 | 1 | 1 | 1 | 1 | 1 | 1 | 1 | 1 | 1 | 1 | 1 | 0 | 1 | 0 | 1 | TRUE |
| 75 | 241 | V3d | occipital | dorsal_visual_area_3 | CHARM | 1 | 1 | 1 | 1 | 0 | 1 | 1 | 1 | 0 | 1 | 1 | 1 | 1 | 1 | 1 | 1 | 0 | 1 | 0 | 1 | TRUE |
| 76 | 242 | V3v | occipital | ventral_visual_area_3 | CHARM | 1 | 1 | 1 | 1 | 0 | 1 | 1 | 1 | 0 | 1 | 1 | 1 | 1 | 1 | 1 | 1 | 1 | 1 | 1 | 1 | TRUE |
| 77 | 244 | possibly_V2 | occipital | possibly_part_of_visual_area_2 | CHARM | 0 | 1 | 0 | 1 | 0 | 1 | 1 | 1 | 0 | 0 | 0 | 1 | 0 | 1 | 0 | 0 | 0 | 1 | 0 | 0 | TRUE |
| 78 | 245 | clearly_V2 | occipital | clearly_part_of_visual_area_2 | CHARM | 1 | 1 | 1 | 1 | 1 | 1 | 1 | 1 | 0 | 1 | 1 | 1 | 1 | 1 | 1 | 1 | 1 | 1 | 1 | 1 | TRUE |
| 79 | 246 | V1 | occipital | primary_visual_cortex | CHARM | 1 | 1 | 1 | 1 | 1 | 1 | 1 | 1 | 0 | 1 | 1 | 0 | 1 | 1 | 1 | 1 | 1 | 1 | 1 | 1 | TRUE |

*Table S2, continued*

| | | | | | | | | | | | | | | | | | | | | | | | | | | |
|---|---|---|---|---|---|---|---|---|---|---|---|---|---|---|---|---|---|---|---|---|---|---|---|---|---|---|
| 80 | 19 | BM | subcorical | basomedial_amygdaloid_nucleus | SARM | 0 | 0 | 0 | 0 | 0 | 0 | 0 | 1 | 0 | 0 | 0 | 0 | 0 | 1 | 0 | 1 | 0 | 0 | 0 | 0 | TRUE |
| 81 | 20 | VCo | subcorical | ventral_cortical_amygdaloid_nucleus | SARM | 0 | 0 | 0 | 0 | 0 | 0 | 0 | 1 | 0 | 0 | 0 | 0 | 0 | 1 | 0 | 0 | 0 | 0 | 0 | 0 | TRUE |
| 82 | 21 | AHi | subcorical | amygdalohippocampal_area | SARM | 0 | 0 | 0 | 0 | 0 | 0 | 0 | 0 | 0 | 0 | 0 | 0 | 0 | 1 | 1 | 0 | 0 | 0 | 0 | 0 | TRUE |
| 83 | 24 | LaD | subcorical | lateral_dorsal_amygdaloid_nucleus | SARM | 0 | 0 | 1 | 0 | 0 | 0 | 1 | 1 | 0 | 0 | 0 | 0 | 0 | 1 | 0 | 0 | 0 | 0 | 0 | 0 | TRUE |
| 84 | 25 | LaV | subcorical | lateral_ventral_amygdaloid_nucleus | SARM | 0 | 0 | 0 | 0 | 0 | 0 | 1 | 1 | 0 | 0 | 0 | 0 | 0 | 1 | 0 | 0 | 0 | 0 | 0 | 0 | TRUE |
| 85 | 26 | APir | subcorical | amygdalopiriform_transition_area | SARM | 0 | 0 | 0 | 0 | 0 | 0 | 0 | 0 | 0 | 0 | 0 | 0 | 0 | 0 | 0 | 0 | 0 | 0 | 0 | 0 | FALSE |
| 86 | 28 | BLD | subcorical | basolateral_dorsal_amygdaloid_nucleus | SARM | 0 | 0 | 0 | 0 | 0 | 0 | 0 | 1 | 0 | 0 | 0 | 0 | 0 | 1 | 0 | 0 | 0 | 0 | 0 | 0 | TRUE |
| 87 | 29 | BLI | subcorical | basolateral_intermediate_amygdaloid_nucleus | SARM | 0 | 0 | 0 | 0 | 0 | 0 | 1 | 1 | 0 | 0 | 0 | 0 | 0 | 1 | 0 | 0 | 0 | 0 | 0 | 0 | TRUE |
| 88 | 30 | BLV | subcorical | basolateral_ventral_amygdaloid_nucleus | SARM | 0 | 0 | 0 | 0 | 0 | 0 | 0 | 1 | 0 | 0 | 0 | 0 | 0 | 1 | 0 | 0 | 0 | 0 | 0 | 0 | TRUE |
| 89 | 31 | PaL | subcorical | paralaminar_amygdaloid_nucleus | SARM | 0 | 0 | 0 | 0 | 0 | 0 | 0 | 0 | 0 | 0 | 0 | 0 | 0 | 0 | 0 | 0 | 0 | 0 | 0 | 0 | FALSE |
| 90 | 34 | Ce | subcorical | central_amygdaloid_nucleus | SARM | 0 | 0 | 0 | 1 | 0 | 0 | 0 | 0 | 0 | 0 | 0 | 0 | 0 | 0 | 0 | 0 | 0 | 0 | 0 | 0 | TRUE |
| 91 | 36 | Me | subcorical | medial_amygdaloid_nucleus | SARM | 0 | 0 | 0 | 1 | 0 | 0 | 0 | 1 | 0 | 0 | 0 | 0 | 0 | 0 | 0 | 0 | 0 | 0 | 0 | 0 | TRUE |
| 92 | 37 | AA | subcorical | anterior_amygdaloid_area | SARM | 0 | 0 | 0 | 0 | 0 | 0 | 0 | 0 | 0 | 0 | 0 | 0 | 0 | 0 | 0 | 0 | 0 | 0 | 0 | 0 | FALSE |
| 93 | 39 | EA | subcorical | extended_amygdala | SARM | 0 | 0 | 0 | 0 | 0 | 0 | 0 | 0 | 0 | 0 | 0 | 0 | 0 | 0 | 0 | 0 | 0 | 0 | 0 | 0 | FALSE |
| 94 | 46 | CdH | subcorical | caudate_head | SARM | 0 | 1 | 0 | 1 | 0 | 1 | 1 | 1 | 0 | 0 | 0 | 0 | 1 | 0 | 0 | 0 | 0 | 0 | 0 | 0 | TRUE |
| 95 | 47 | CdT | subcorical | caudate_tail | SARM | 0 | 0 | 0 | 1 | 0 | 0 | 0 | 0 | 0 | 0 | 0 | 0 | 0 | 1 | 0 | 1 | 0 | 0 | 0 | 0 | TRUE |
| 96 | 48 | Pu | subcorical | putamen | SARM | 1 | 1 | 0 | 1 | 0 | 1 | 1 | 1 | 0 | 0 | 0 | 0 | 0 | 1 | 0 | 1 | 0 | 1 | 0 | 0 | TRUE |
| 97 | 52 | Acb | subcorical | accumbens | SARM | 0 | 0 | 0 | 0 | 0 | 1 | 1 | 0 | 0 | 0 | 0 | 0 | 1 | 0 | 0 | 1 | 0 | 0 | 0 | 0 | TRUE |
| 98 | 119 | MD | subcorical | mediodorsal_thalamus | SARM | 0 | 1 | 0 | 0 | 0 | 0 | 1 | 1 | 0 | 0 | 0 | 1 | 0 | 1 | 0 | 0 | 0 | 0 | 0 | 0 | TRUE |
| 99 | 121 | IMD | subcorical | intermediodorsal_thalamus | SARM | 0 | 0 | 0 | 0 | 0 | 0 | 1 | 0 | 0 | 0 | 0 | 0 | 0 | 0 | 0 | 0 | 0 | 0 | 0 | 0 | TRUE |
| 100 | 122 | PT-PV-sm | subcorical | paratenial-paraventral-thalamus | SARM | 0 | 0 | 0 | 0 | 0 | 0 | 0 | 0 | 0 | 0 | 0 | 0 | 0 | 0 | 0 | 0 | 0 | 0 | 0 | 0 | FALSE |
| 101 | 123 | PVP | subcorical | paraventricular_posterior_thalamus | SARM | 0 | 0 | 0 | 0 | 0 | 0 | 0 | 0 | 0 | 0 | 0 | 0 | 0 | 0 | 0 | 0 | 0 | 0 | 0 | 0 | FALSE |
| 102 | 124 | PVA | subcorical | paraventricular_anterior_thalamus | SARM | 0 | 0 | 0 | 0 | 0 | 0 | 0 | 0 | 0 | 0 | 0 | 0 | 0 | 0 | 0 | 0 | 0 | 0 | 0 | 0 | FALSE |
| 103 | 125 | IAM | subcorical | interanteromedial_thalamus | SARM | 0 | 0 | 0 | 0 | 0 | 0 | 0 | 0 | 0 | 0 | 0 | 0 | 0 | 0 | 0 | 0 | 0 | 0 | 0 | 0 | FALSE |
| 104 | 126 | CM | subcorical | central_medial_thalamus | SARM | 0 | 0 | 0 | 0 | 0 | 0 | 1 | 0 | 0 | 0 | 0 | 0 | 0 | 0 | 0 | 0 | 0 | 0 | 0 | 0 | TRUE |
| 105 | 130 | VA | subcorical | ventral_anterior_thalamus | SARM | 0 | 0 | 0 | 0 | 0 | 0 | 1 | 0 | 0 | 0 | 0 | 0 | 0 | 0 | 0 | 0 | 0 | 0 | 0 | 0 | TRUE |
| 106 | 131 | VLA | subcorical | ventral_lateral_anterior_thalamus | SARM | 0 | 0 | 0 | 0 | 0 | 0 | 1 | 0 | 0 | 0 | 0 | 0 | 0 | 0 | 0 | 0 | 0 | 0 | 0 | 0 | TRUE |
| 107 | 133 | VLX | subcorical | ventral_lateral_x_thalamus | SARM | 0 | 0 | 0 | 0 | 0 | 0 | 0 | 0 | 0 | 0 | 0 | 0 | 0 | 0 | 0 | 0 | 0 | 0 | 0 | 0 | FALSE |
| 108 | 134 | VLPV | subcorical | ventral_lateral_posteroventral_thalamus | SARM | 0 | 0 | 0 | 0 | 0 | 0 | 0 | 1 | 0 | 0 | 0 | 0 | 0 | 0 | 0 | 0 | 0 | 0 | 0 | 0 | TRUE |
| 109 | 135 | VLPD | subcorical | ventral_lateral_posterodorsal_thalamus | SARM | 0 | 0 | 0 | 0 | 0 | 0 | 0 | 0 | 0 | 0 | 0 | 1 | 0 | 0 | 0 | 0 | 0 | 0 | 0 | 0 | TRUE |
| 110 | 137 | VPM-VPL | subcorical | ventroposterior_medial_and_lateral_thalamus | SARM | 0 | 0 | 0 | 0 | 0 | 0 | 0 | 0 | 0 | 0 | 0 | 0 | 0 | 0 | 0 | 0 | 0 | 0 | 0 | 0 | FALSE |
| 111 | 138 | VMPo-VMB | subcorical | ventromedial_posterior+basal_thalamus | SARM | 0 | 0 | 0 | 1 | 0 | 0 | 0 | 0 | 0 | 0 | 0 | 0 | 0 | 0 | 0 | 0 | 0 | 0 | 0 | 0 | TRUE |
| 112 | 139 | VPI | subcorical | ventroposterior_inferior_thalamus | SARM | 0 | 0 | 0 | 1 | 0 | 0 | 0 | 0 | 0 | 0 | 0 | 0 | 0 | 0 | 0 | 0 | 0 | 0 | 0 | 0 | TRUE |
| 113 | 140 | VM | subcorical | ventral_medial_thalamus | SARM | 0 | 0 | 0 | 0 | 0 | 0 | 1 | 0 | 0 | 0 | 0 | 0 | 0 | 0 | 0 | 0 | 0 | 0 | 0 | 0 | TRUE |
| 114 | 142 | LP | subcorical | lateral_posterior_thalamus | SARM | 0 | 1 | 0 | 1 | 1 | 0 | 1 | 0 | 0 | 0 | 0 | 1 | 0 | 1 | 0 | 0 | 0 | 0 | 0 | 0 | TRUE |
| 115 | 143 | LD | subcorical | lateral_dorsal_thalamus | SARM | 0 | 1 | 0 | 0 | 0 | 0 | 0 | 0 | 0 | 0 | 0 | 1 | 0 | 1 | 0 | 0 | 0 | 0 | 0 | 0 | TRUE |
| 116 | 145 | APul | subcorical | anterior_pulvinar | SARM | 0 | 1 | 0 | 1 | 0 | 0 | 1 | 0 | 0 | 1 | 0 | 1 | 0 | 1 | 0 | 0 | 0 | 0 | 0 | 1 | TRUE |
| 117 | 146 | MPul | subcorical | medial_pulvinar | SARM | 0 | 1 | 0 | 1 | 1 | 0 | 1 | 1 | 0 | 1 | 0 | 1 | 0 | 1 | 0 | 0 | 0 | 0 | 0 | 1 | TRUE |
| 118 | 147 | LPul | subcorical | lateral_pulvinar | SARM | 0 | 1 | 0 | 1 | 0 | 0 | 1 | 1 | 0 | 1 | 0 | 1 | 0 | 1 | 0 | 0 | 0 | 0 | 0 | 1 | TRUE |
| 119 | 148 | IPul | subcorical | inferior_pulvinar | SARM | 0 | 1 | 0 | 1 | 0 | 0 | 1 | 1 | 0 | 1 | 0 | 1 | 0 | 1 | 0 | 0 | 0 | 0 | 0 | 1 | TRUE |
| 120 | 151 | DLG | subcorical | dorsal_lateral_geniculate | SARM | 0 | 0 | 0 | 1 | 0 | 0 | 0 | 1 | 0 | 1 | 0 | 0 | 0 | 0 | 0 | 0 | 0 | 0 | 0 | 1 | TRUE |
| 121 | 152 | MG | subcorical | medial_geniculate | SARM | 0 | 1 | 0 | 1 | 0 | 0 | 0 | 1 | 0 | 1 | 0 | 1 | 0 | 1 | 0 | 0 | 0 | 0 | 0 | 1 | TRUE |

*Table S2, continued*

| | | | | | | | | | | | | | | | | | | | | | | | | | |
|---|---|---|---|---|---|---|---|---|---|---|---|---|---|---|---|---|---|---|---|---|---|---|---|---|---|
| 122 | 166 | SC | subcorical | superior_colliculus | SARM | 0 | 1 | 0 | 1 | 0 | 0 | 1 | 1 | 0 | 1 | 0 | 1 | 0 | 1 | 0 | 0 | 0 | 0 | 0 | 1 | TRUE |
| 123 | 240 | mcp | subcorical | medial_cerebellar_peduncle | SARM | 0 | 0 | 0 | 0 | 0 | 0 | 0 | 0 | 0 | 0 | 0 | 0 | 0 | 0 | 0 | 0 | 1 | 0 | 0 | 0 | TRUE |
| 124 | 243 | icp | subcorical | inferior_cerebellar_peduncle | SARM | 0 | 0 | 0 | 0 | 0 | 0 | 0 | 0 | 0 | 0 | 0 | 0 | 0 | 0 | 0 | 0 | 0 | 0 | 0 | 0 | FALSE |
| 125 | 246 | Lat | subcorical | lateral_cerebellar_nucleus | SARM | 0 | 0 | 0 | 0 | 0 | 0 | 1 | 0 | 0 | 0 | 0 | 0 | 0 | 0 | 0 | 0 | 0 | 0 | 0 | 0 | TRUE |
| 126 | 248 | IntA | subcorical | anterior_interposed_cerebellar_nucleus | SARM | 0 | 0 | 0 | 0 | 0 | 0 | 1 | 0 | 0 | 0 | 0 | 0 | 0 | 0 | 0 | 0 | 0 | 0 | 0 | 0 | TRUE |
| 127 | 249 | IntP | subcorical | posterior_interposed_cerebellar_nucleus | SARM | 0 | 0 | 0 | 0 | 0 | 0 | 1 | 0 | 0 | 0 | 0 | 0 | 0 | 0 | 0 | 0 | 0 | 0 | 0 | 0 | TRUE |
| 128 | 250 | Fas | subcorical | fastigial_cerebellar_nucleus | SARM | 0 | 0 | 0 | 0 | 0 | 0 | 0 | 0 | 0 | 0 | 0 | 0 | 0 | 0 | 0 | 0 | 0 | 0 | 0 | 0 | FALSE |
| 129 | 253 | Fl | subcorical | flocculus | SARM | 0 | 0 | 0 | 0 | 0 | 0 | 0 | 0 | 0 | 0 | 0 | 0 | 0 | 0 | 0 | 0 | 0 | 0 | 0 | 0 | FALSE |
| 130 | 254 | PFl | subcorical | paraflocculus | SARM | 0 | 0 | 0 | 0 | 0 | 0 | 1 | 0 | 0 | 0 | 0 | 0 | 0 | 0 | 0 | 0 | 0 | 0 | 0 | 0 | TRUE |
| 131 | 257 | Cb1V | subcorical | cerebellar_lobule_1_vermis | SARM | 0 | 0 | 0 | 0 | 0 | 0 | 0 | 0 | 0 | 0 | 0 | 0 | 0 | 0 | 0 | 0 | 0 | 0 | 0 | 0 | FALSE |
| 132 | 258 | Cb2V | subcorical | cerebellar_lobule_2_vermis | SARM | 0 | 0 | 0 | 0 | 0 | 0 | 0 | 0 | 0 | 0 | 0 | 0 | 0 | 0 | 0 | 0 | 0 | 0 | 0 | 0 | FALSE |
| 133 | 259 | Cb3V | subcorical | cerebellar_lobule_3_vermis | SARM | 0 | 0 | 0 | 0 | 0 | 0 | 1 | 1 | 0 | 0 | 0 | 0 | 0 | 0 | 0 | 0 | 0 | 0 | 0 | 0 | TRUE |
| 134 | 260 | Cb4V | subcorical | cerebellar_lobule_4_vermis | SARM | 0 | 0 | 0 | 0 | 0 | 0 | 1 | 1 | 0 | 0 | 0 | 1 | 0 | 0 | 0 | 0 | 0 | 1 | 0 | 0 | TRUE |
| 135 | 261 | Cb5V | subcorical | cerebellar_lobule_5_vermis | SARM | 0 | 1 | 1 | 1 | 0 | 1 | 1 | 1 | 0 | 0 | 0 | 0 | 0 | 1 | 1 | 1 | 0 | 1 | 0 | 0 | TRUE |
| 136 | 263 | Cb6V | subcorical | cerebellar_lobule_6_vermis | SARM | 0 | 0 | 0 | 0 | 0 | 0 | 1 | 1 | 0 | 0 | 0 | 0 | 0 | 0 | 1 | 1 | 0 | 0 | 0 | 0 | TRUE |
| 137 | 264 | Cb7V | subcorical | cerebellar_lobule_7_vermis | SARM | 0 | 0 | 0 | 0 | 0 | 0 | 0 | 0 | 0 | 0 | 0 | 0 | 0 | 0 | 1 | 1 | 0 | 0 | 0 | 0 | TRUE |
| 138 | 265 | Cb8V | subcorical | cerebellar_lobule_8_vermis | SARM | 0 | 0 | 0 | 0 | 0 | 0 | 0 | 0 | 0 | 0 | 0 | 0 | 0 | 0 | 0 | 0 | 0 | 0 | 0 | 0 | FALSE |
| 139 | 266 | Cb9V | subcorical | cerebellar_lobule_9_vermis | SARM | 0 | 0 | 0 | 0 | 0 | 0 | 0 | 0 | 0 | 0 | 0 | 0 | 0 | 0 | 0 | 0 | 0 | 0 | 0 | 0 | FALSE |
| 140 | 267 | Cb10V | subcorical | cerebellar_lobule_10_vermis | SARM | 0 | 0 | 0 | 0 | 0 | 0 | 0 | 0 | 0 | 0 | 0 | 0 | 0 | 0 | 0 | 0 | 0 | 0 | 0 | 0 | FALSE |
| 141 | 269 | Cb3I | subcorical | cerebellar_lobule_3_intermediate | SARM | 0 | 0 | 0 | 0 | 0 | 0 | 0 | 0 | 0 | 0 | 0 | 0 | 0 | 0 | 0 | 0 | 0 | 0 | 0 | 0 | FALSE |
| 142 | 270 | Cb4I | subcorical | cerebellar_lobule_4_intermediate | SARM | 0 | 0 | 1 | 0 | 0 | 0 | 1 | 1 | 0 | 0 | 0 | 0 | 0 | 0 | 0 | 0 | 0 | 0 | 0 | 0 | TRUE |
| 143 | 271 | Cb5I | subcorical | cerebellar_lobule_5_intermediate | SARM | 0 | 0 | 1 | 0 | 0 | 0 | 1 | 1 | 0 | 0 | 0 | 0 | 0 | 0 | 0 | 0 | 1 | 0 | 0 | 0 | TRUE |
| 144 | 272 | Cb6I | subcorical | cerebellar_lobule_6_intermediate | SARM | 0 | 0 | 0 | 0 | 0 | 0 | 1 | 1 | 0 | 0 | 0 | 0 | 0 | 0 | 0 | 1 | 0 | 1 | 0 | 0 | TRUE |
| 145 | 273 | Cop | subcorical | copula_of_the_pyramis | SARM | 0 | 0 | 0 | 0 | 0 | 0 | 0 | 0 | 0 | 0 | 0 | 0 | 0 | 0 | 0 | 0 | 0 | 0 | 0 | 0 | FALSE |
| 146 | 274 | Sim | subcorical | simple_lobule | SARM | 0 | 0 | 1 | 1 | 0 | 0 | 1 | 1 | 0 | 0 | 0 | 0 | 0 | 0 | 0 | 1 | 0 | 1 | 0 | 0 | TRUE |
| 147 | 275 | PM | subcorical | paramedian_lobule | SARM | 0 | 0 | 0 | 0 | 0 | 0 | 0 | 0 | 0 | 0 | 0 | 0 | 0 | 0 | 0 | 0 | 0 | 0 | 0 | 0 | FALSE |
| 148 | 277 | Crus1 | subcorical | ansiform_lobule_crus1 | SARM | 0 | 0 | 1 | 0 | 0 | 0 | 1 | 0 | 0 | 1 | 0 | 0 | 0 | 1 | 0 | 1 | 0 | 0 | 0 | 0 | TRUE |
| 149 | 278 | Crus2 | subcorical | ansiform_lobule_crus2 | SARM | 0 | 0 | 0 | 0 | 0 | 0 | 0 | 0 | 0 | 0 | 0 | 0 | 0 | 0 | 0 | 0 | 0 | 0 | 0 | 0 | FALSE |

**Table S3.** ROIs selected for each dataset. Center voxel is the center-of-gravity.

| ROI | Hemisphere | N of voxels | Center voxel (mm from AC) | | | Mean t-value |
|---|---|---|---|---|---|---|
| | | | x | y | z | |
| **monkey C** | **dPul(a)** | | | | | |
| area-24b-prime | R | 34 | 1.8 | -7.6 | 12.1 | 3.57 |
| area-8Ad | R | 628 | 14.1 | 7.8 | 14.7 | 6.25 |
| area-8Av | R | 218 | 17.9 | 7.3 | 13.5 | 5.88 |
| area-8Bs | R | 1264 | 11.2 | 8.2 | 12.9 | 5.58 |
| area-9d | R | 52 | 11.0 | 14.8 | 15.3 | 3.75 |
| area-46d | R | 560 | 13.3 | 12.7 | 13.2 | 3.92 |
| area-46f | R | 183 | 11.2 | 12.4 | 10.2 | 3.48 |
| area-46v | R | 1280 | 15.1 | 13.1 | 11.8 | 4.20 |
| area-12r | R | 50 | 17.6 | 17.7 | 10.0 | 3.40 |
| area-45a | R | 506 | 18.7 | 13.5 | 10.6 | 4.02 |
| area-45b | R | 989 | 16.7 | 8.7 | 10.1 | 5.68 |
| area-44 | R | 388 | 15.2 | 8.2 | 7.6 | 4.27 |
| PMdc | R | 1431 | 10.4 | 5.0 | 15.9 | 4.62 |
| F4 | R | 130 | 16.0 | 4.4 | 14.6 | 4.09 |
| F5 | R | 718 | 15.4 | 4.8 | 11.0 | 4.02 |
| SMA | R | 103 | 2.8 | 6.2 | 20.1 | 3.26 |
| V6 | R | 169 | 2.3 | -30.4 | 7.4 | 3.95 |
| V6Av | R | 45 | 1.3 | -30.4 | 8.5 | 3.20 |
| V6Ad | R | 98 | 2.0 | -31.0 | 16.0 | 3.61 |
| PEc-PEci | R | 192 | 6.1 | -25.3 | 21.8 | 3.40 |
| PEa | R | 68 | 9.9 | -22.5 | 20.5 | 3.12 |
| MIP | R | 308 | 4.9 | -27.5 | 14.6 | 3.34 |
| VIP | R | 388 | 6.6 | -20.6 | 12.6 | 3.70 |
| LIPd | R | 204 | 10.9 | -22.6 | 18.5 | 3.48 |
| LIPv | R | 367 | 7.7 | -22.8 | 14.3 | 3.65 |
| MST | R | 1706 | 13.9 | -21.2 | 8.9 | 5.17 |
| area-7a | R | 158 | 12.4 | -24.3 | 17.5 | 3.45 |
| area-7m | R | 437 | 1.5 | -24.7 | 14.5 | 3.42 |
| area-31 | R | 411 | 1.8 | -22.6 | 12.9 | 4.05 |
| area-23a | R | 136 | 2.3 | -19.7 | 8.3 | 3.45 |
| area-23b | R | 585 | 1.5 | -16.4 | 13.5 | 3.82 |
| area-v23 | R | 415 | 2.1 | -21.6 | 7.1 | 3.89 |
| area-29 | R | 33 | 3.9 | -19.0 | 8.2 | 3.33 |
| area-30 | R | 84 | 2.9 | -18.9 | 7.9 | 3.39 |
| TEO | R | 169 | 22.7 | -15.8 | 3.1 | 3.90 |
| TEpv | R | 32 | 19.0 | -19.2 | -4.5 | 3.53 |
| TEpd | R | 38 | 26.0 | -8.9 | -6.6 | 3.19 |
| TEm | R | 339 | 24.5 | -7.2 | -6.3 | 3.76 |
| TEa | R | 138 | 21.4 | -7.5 | -7.0 | 3.44 |
| IPa | R | 610 | 18.1 | -8.5 | -5.3 | 3.94 |
| PGa | R | 1431 | 16.8 | -13.9 | 1.5 | 5.32 |
| FST | R | 613 | 16.5 | -19.9 | 4.1 | 4.30 |
| TPO | R | 1436 | 20.8 | -10.5 | -0.7 | 4.64 |
| TAa | R | 438 | 25.0 | -7.2 | -1.9 | 4.35 |
| Tpt | R | 39 | 19.9 | -18.8 | 10.1 | 3.21 |
| Ia-Id | R | 125 | 19.2 | 0.6 | -1.3 | 3.49 |
| MT | R | 486 | 14.5 | -24.3 | 10.3 | 4.44 |
| V4d | R | 251 | 14.6 | -26.7 | 15.4 | 3.60 |
| V4v | R | 266 | 17.9 | -20.5 | -3.5 | 3.81 |
| V3A | R | 427 | 12.3 | -28.6 | 12.0 | 3.94 |
| V3d | R | 400 | 11.6 | -32.6 | 11.3 | 3.93 |
| V3v | R | 73 | 12.7 | -23.7 | -3.3 | 3.20 |
| possibly-V2 | R | 219 | 5.4 | -18.9 | 0.7 | 4.27 |
| clearly-V2 | R | 1983 | 8.2 | -28.8 | 6.2 | 3.92 |
| V1 | R | 712 | 11.1 | -29.2 | 5.8 | 3.59 |
| CdH | R | 274 | 5.2 | -0.9 | 7.3 | 3.53 |
| Pu | R | 28 | 9.3 | 8.3 | 2.9 | 3.14 |
| MD | R | 98 | 2.8 | -11.8 | 4.5 | 3.69 |
| LP | R | 96 | 6.1 | -12.3 | 5.1 | 7.07 |
| LD | R | 65 | 4.4 | -11.1 | 5.9 | 6.21 |
| APul | R | 333 | 6.2 | -12.6 | 2.1 | 6.99 |
| MPul | R | 312 | 6.5 | -15.2 | 1.8 | 5.65 |
| LPul | R | 69 | 10.2 | -13.8 | 1.6 | 4.52 |
| IPul | R | 103 | 9.6 | -13.0 | -0.9 | 4.43 |
| MG | R | 120 | 7.5 | -12.7 | -1.7 | 4.43 |
| SC | R | 437 | 2.9 | -15.7 | -0.2 | 4.03 |
| Cb5V | R | 29 | 2.4 | -27.0 | 3.3 | 2.69 |
| area-8Ad | L | 116 | -13.0 | 4.3 | 14.1 | 3.39 |

*Table S3, continued*

| | | | | | | |
|---|---|---|---|---|---|---|
| area-8Bs | L | 90 | -11.6 | 4.3 | 12.5 | 3.58 |
| area-45b | L | 92 | -12.6 | 5.6 | 9.5 | 3.59 |
| area-44 | L | 34 | -12.4 | 4.9 | 8.4 | 3.25 |
| PMdc | L | 166 | -11.8 | 1.3 | 14.7 | 3.57 |
| F4 | L | 59 | -14.2 | 2.1 | 14.7 | 3.39 |
| F5 | L | 284 | -12.5 | 2.5 | 11.9 | 3.45 |
| V6Av | L | 56 | -0.7 | -29.8 | 9.5 | 3.17 |
| V6Ad | L | 143 | -1.4 | -30.8 | 16.3 | 3.78 |
| PEa | L | 141 | -9.2 | -17.9 | 17.7 | 3.35 |
| MIP | L | 103 | -3.2 | -27.2 | 16.3 | 3.46 |
| LIPd | L | 386 | -11.1 | -19.7 | 18.3 | 3.54 |
| LIPv | L | 150 | -9.0 | -19.3 | 15.2 | 3.37 |
| LOP | L | 59 | -9.0 | -28.6 | 16.2 | 3.54 |
| MST | L | 306 | -14.9 | -21.0 | 9.4 | 3.67 |
| area-7a | L | 566 | -11.4 | -23.7 | 20.2 | 3.84 |
| area-7m | L | 194 | -0.3 | -24.9 | 14.6 | 3.17 |
| area-31 | L | 113 | -0.3 | -20.7 | 16.7 | 3.86 |
| area-23b | L | 190 | 0.1 | -15.4 | 14.3 | 3.32 |
| TEO | L | 195 | -22.2 | -16.0 | 2.2 | 3.17 |
| TEpv | L | 159 | -18.4 | -18.2 | -5.5 | 3.35 |
| IPa | L | 160 | -20.3 | -16.4 | -1.5 | 3.58 |
| PGa | L | 204 | -16.5 | -17.4 | 5.0 | 3.96 |
| FST | L | 160 | -16.7 | -20.4 | 5.2 | 3.13 |
| TPO | L | 137 | -20.9 | -14.6 | 3.0 | 3.50 |
| MT | L | 493 | -16.0 | -23.8 | 12.1 | 3.83 |
| V4d | L | 625 | -17.5 | -26.6 | 14.4 | 3.70 |
| V4v | L | 170 | -23.0 | -28.3 | -2.5 | 3.35 |
| V3A | L | 454 | -12.2 | -28.0 | 15.3 | 3.52 |
| V3d | L | 115 | -10.9 | -31.1 | 14.2 | 3.12 |
| V3v | L | 168 | -21.9 | -29.6 | -2.3 | 3.43 |
| clearly-V2 | L | 1363 | -13.2 | -32.6 | 12.6 | 3.89 |
| V1 | L | 1948 | -14.5 | -35.4 | 10.1 | 3.92 |
| Pu | L | 111 | -12.6 | -1.1 | 4.2 | 3.46 |
| **monkey C** | **dPul** | | | | | |
| area-24b | R | 98 | 1.0 | 15.3 | 8.8 | 3.23 |
| area-24c | R | 31 | 0.3 | 20.7 | 9.1 | 2.95 |
| area-8Ad | R | 487 | 14.9 | 6.7 | 14.6 | 6.07 |
| area-8Av | R | 320 | 18.2 | 7.7 | 13.8 | 7.65 |
| area-8Bs | R | 553 | 12.3 | 6.8 | 12.5 | 4.97 |
| area-9m | R | 26 | 2.1 | 22.8 | 8.3 | 3.15 |
| area-46d | R | 639 | 14.4 | 11.3 | 13.6 | 4.51 |
| area-46f | R | 48 | 11.4 | 12.5 | 10.7 | 2.66 |
| area-46v | R | 1662 | 15.2 | 13.8 | 12.0 | 6.03 |
| area-12l | R | 84 | 18.8 | 16.2 | 7.4 | 4.33 |
| area-12r | R | 408 | 15.8 | 18.8 | 10.5 | 4.89 |
| area-45a | R | 965 | 18.4 | 14.5 | 9.8 | 5.77 |
| area-45b | R | 1104 | 16.9 | 8.9 | 10.0 | 7.53 |
| area-44 | R | 400 | 15.2 | 8.4 | 7.5 | 4.04 |
| PMdc | R | 607 | 11.7 | 3.5 | 19.6 | 3.30 |
| F4 | R | 215 | 16.6 | 4.5 | 14.5 | 3.91 |
| F5 | R | 685 | 15.7 | 5.1 | 10.7 | 4.39 |
| MIP | R | 131 | 5.9 | -27.5 | 15.1 | 3.13 |
| VIP | R | 117 | 6.7 | -21.5 | 12.0 | 3.10 |
| PIP | R | 40 | 10.1 | -26.7 | 9.9 | 2.82 |
| LIPd | R | 143 | 7.6 | -27.8 | 16.8 | 4.01 |
| LIPv | R | 359 | 7.8 | -23.8 | 15.4 | 3.55 |
| LOP | R | 50 | 8.4 | -30.1 | 16.4 | 3.38 |
| MST | R | 1389 | 14.2 | -21.5 | 9.1 | 4.56 |
| area-7a | R | 285 | 12.3 | -26.1 | 17.2 | 3.33 |
| area-v23 | R | 98 | 2.4 | -19.8 | 2.7 | 3.40 |
| TEO | R | 1085 | 23.8 | -18.8 | 1.9 | 4.18 |
| TEpd | R | 33 | 23.6 | -21.1 | -4.5 | 3.22 |
| TEm | R | 102 | 21.8 | -6.5 | -7.3 | 3.49 |
| TEa | R | 269 | 19.9 | -5.2 | -8.6 | 3.67 |
| IPa | R | 808 | 18.2 | -8.8 | -5.3 | 3.71 |
| PGa | R | 1417 | 17.1 | -13.6 | 1.2 | 5.77 |
| FST | R | 761 | 17.8 | -20.1 | 3.8 | 4.47 |
| TPO | R | 1994 | 20.5 | -11.3 | 0.1 | 5.48 |
| TAa | R | 572 | 24.9 | -11.6 | 2.9 | 4.50 |
| STGr | R | 78 | 21.5 | 5.6 | -8.3 | 3.75 |
| Tpt | R | 76 | 21.9 | -20.8 | 10.3 | 4.25 |
| Ia-Id | R | 118 | 19.3 | -3.6 | 1.5 | 3.08 |
| Ig | R | 87 | 17.9 | -8.6 | 0.6 | 3.67 |
| MT | R | 1180 | 17.7 | -23.7 | 9.3 | 4.47 |

*Table S3, continued*

| | | | | | | |
|---|---|---|---|---|---|---|
| V4d | R | 2705 | 20.1 | -24.9 | 11.1 | 4.29 |
| V4v | R | 915 | 19.8 | -23.7 | -2.9 | 3.71 |
| V3A | R | 754 | 15.7 | -28.4 | 9.6 | 4.12 |
| V3d | R | 953 | 17.5 | -28.9 | 8.0 | 4.02 |
| V3v | R | 714 | 17.7 | -24.2 | -1.8 | 3.73 |
| possibly-V2 | R | 107 | 5.2 | -18.5 | 1.0 | 3.30 |
| clearly-V2 | R | 2802 | 18.8 | -29.7 | 7.1 | 3.84 |
| V1 | R | 1066 | 15.2 | -34.5 | 8.8 | 3.59 |
| Ce | R | 32 | 11.0 | -4.1 | -5.0 | 3.53 |
| Me | R | 51 | 10.0 | -4.6 | -5.6 | 3.33 |
| CdH | R | 252 | 5.0 | 6.4 | 6.4 | 3.32 |
| CdT | R | 70 | 12.8 | -10.2 | -3.7 | 3.28 |
| Pu | R | 59 | 12.6 | -1.7 | -3.6 | 3.09 |
| VMPo-VMB | R | 43 | 6.3 | -10.5 | -0.6 | 5.25 |
| VPI | R | 38 | 8.7 | -9.9 | -0.7 | 5.35 |
| LP | R | 67 | 6.1 | -12.3 | 4.4 | 5.67 |
| APul | R | 400 | 6.3 | -12.4 | 2.1 | 7.63 |
| MPul | R | 413 | 6.9 | -15.4 | 1.5 | 6.20 |
| LPul | R | 89 | 10.8 | -14.7 | 0.3 | 4.02 |
| IPul | R | 251 | 10.0 | -13.7 | -1.5 | 4.53 |
| DLG | R | 35 | 9.4 | -9.2 | -4.5 | 3.25 |
| MG | R | 259 | 7.8 | -12.0 | -2.2 | 5.11 |
| SC | R | 264 | 3.0 | -15.1 | 0.3 | 3.92 |
| Cb5V | R | 46 | 0.4 | -28.9 | 2.6 | 3.73 |
| Sim | R | 188 | 10.1 | -28.0 | -5.9 | 3.22 |
| area-24b | L | 42 | -0.2 | 14.7 | 8.9 | 3.21 |
| area-24c | L | 34 | -0.4 | 20.9 | 9.4 | 2.85 |
| area-8Ad | L | 87 | -13.6 | 4.8 | 11.9 | 3.66 |
| area-8Av | L | 73 | -16.8 | 6.8 | 11.5 | 3.50 |
| area-8Bs | L | 53 | -12.1 | 4.7 | 10.9 | 3.82 |
| area-9m | L | 28 | -1.0 | 21.4 | 10.2 | 2.86 |
| area-46v | L | 24 | -15.0 | 8.4 | 12.0 | 3.18 |
| area-45b | L | 311 | -14.8 | 6.3 | 10.4 | 3.52 |
| PMdc | L | 127 | -10.8 | 0.1 | 13.6 | 3.39 |
| F5 | L | 154 | -12.3 | 1.4 | 12.8 | 3.15 |
| LIPd | L | 129 | -10.9 | -18.6 | 16.4 | 3.42 |
| LIPv | L | 197 | -8.5 | -20.1 | 15.7 | 3.87 |
| LOP | L | 55 | -7.6 | -28.6 | 18.4 | 3.22 |
| MST | L | 197 | -17.3 | -19.8 | 8.2 | 3.86 |
| TEO | L | 1260 | -23.7 | -17.2 | 2.8 | 4.11 |
| PGa | L | 189 | -17.6 | -16.5 | 4.8 | 4.04 |
| FST | L | 231 | -19.5 | -19.9 | 4.1 | 4.24 |
| TPO | L | 601 | -20.8 | -13.1 | 2.3 | 4.80 |
| TAa | L | 278 | -23.9 | -12.3 | 3.5 | 4.30 |
| STGr | L | 108 | -21.8 | 5.0 | -8.7 | 3.36 |
| Tpt | L | 63 | -18.6 | -20.4 | 12.4 | 3.13 |
| MT | L | 66 | -19.7 | -21.9 | 8.9 | 2.88 |
| V4d | L | 1698 | -20.4 | -23.8 | 8.8 | 4.42 |
| V4v | L | 277 | -21.6 | -22.9 | 0.4 | 4.93 |
| V3A | L | 161 | -18.6 | -27.8 | 9.3 | 5.58 |
| V3d | L | 553 | -19.9 | -27.0 | 7.1 | 5.01 |
| V3v | L | 359 | -19.8 | -24.8 | 1.1 | 4.56 |
| clearly-V2 | L | 2290 | -19.2 | -28.9 | 7.6 | 4.38 |
| V1 | L | 678 | -18.8 | -32.9 | 9.2 | 3.57 |
| LaD | L | 38 | -13.0 | -1.3 | -8.9 | 3.46 |
| Cb5V | L | 93 | -1.1 | -29.0 | 2.5 | 3.54 |
| Cb4I | L | 24 | -6.8 | -20.9 | -6.6 | 2.91 |
| Cb5I | L | 72 | -8.6 | -19.3 | -7.0 | 3.18 |
| Sim | L | 159 | -11.7 | -27.3 | -6.6 | 3.39 |
| Crus1 | L | 122 | -14.2 | -25.7 | -7.0 | 3.15 |
| **monkey C** | **LIPd-a** | | | | | |
| area-24b-prime | R | 28 | -0.1 | -7.4 | 12.4 | 3.33 |
| area-8Ad | R | 515 | 13.5 | 8.3 | 15.4 | 6.30 |
| area-8Bs | R | 1139 | 11.2 | 8.0 | 13.1 | 6.47 |
| area-46d | R | 597 | 12.5 | 12.4 | 12.7 | 5.34 |
| area-46f | R | 381 | 10.1 | 13.7 | 10.2 | 4.17 |
| area-46v | R | 707 | 13.1 | 13.3 | 10.9 | 4.78 |
| area-44 | R | 24 | 14.2 | 10.4 | 7.4 | 3.43 |
| PMdc | R | 750 | 10.4 | 5.3 | 13.9 | 4.64 |
| F5 | R | 200 | 13.7 | 3.5 | 11.4 | 3.47 |
| V6 | R | 159 | 2.5 | -30.7 | 7.6 | 3.19 |
| V6Av | R | 246 | 2.8 | -31.5 | 11.3 | 3.50 |
| V6Ad | R | 173 | 2.4 | -31.1 | 15.1 | 3.65 |
| PEc-PEci | R | 34 | 1.7 | -30.2 | 16.9 | 5.21 |

| | | | | | | |
|---|---|---|---|---|---|---|
| PEa | R | 465 | 8.8 | -18.9 | 15.6 | 6.10 |
| MIP | R | 1075 | 5.5 | -29.0 | 13.9 | 4.79 |
| VIP | R | 1063 | 7.5 | -19.3 | 12.1 | 7.48 |
| PIP | R | 190 | 9.3 | -26.5 | 11.2 | 3.49 |
| LIPd | R | 805 | 9.7 | -24.2 | 16.9 | 11.23 |
| LIPv | R | 1250 | 8.6 | -22.5 | 14.3 | 12.67 |
| LOP | R | 383 | 8.9 | -29.9 | 14.9 | 5.31 |
| MST | R | 2085 | 13.8 | -21.3 | 9.5 | 7.10 |
| area-7a | R | 793 | 11.9 | -25.4 | 16.7 | 6.94 |
| area-7m | R | 535 | 1.4 | -26.8 | 12.7 | 4.04 |
| area-31 | R | 283 | 1.7 | -23.7 | 10.2 | 4.78 |
| area-23a | R | 74 | 1.7 | -15.2 | 9.8 | 3.79 |
| area-23b | R | 260 | 1.0 | -15.3 | 13.5 | 3.79 |
| area-v23 | R | 467 | 2.6 | -23.4 | 6.5 | 4.27 |
| IPa | R | 481 | 17.3 | -11.7 | -2.3 | 5.19 |
| PGa | R | 1525 | 16.7 | -14.4 | 2.0 | 9.55 |
| FST | R | 363 | 16.0 | -19.7 | 4.2 | 4.94 |
| TPO | R | 1069 | 20.1 | -12.4 | 1.3 | 7.63 |
| TAa | R | 203 | 24.0 | -11.2 | 1.9 | 5.67 |
| Ia-Id | R | 61 | 18.4 | -4.6 | 0.6 | 4.53 |
| Ig | R | 207 | 16.4 | -7.6 | 2.6 | 4.97 |
| MT | R | 656 | 15.1 | -24.2 | 10.3 | 5.91 |
| V4d | R | 1047 | 12.1 | -29.4 | 16.7 | 5.44 |
| V3A | R | 673 | 11.3 | -28.9 | 11.6 | 3.81 |
| V3d | R | 316 | 9.7 | -32.1 | 12.4 | 3.60 |
| V3v | R | 176 | 10.3 | -20.2 | -4.0 | 3.57 |
| possibly-V2 | R | 94 | 5.6 | -19.8 | 1.2 | 4.15 |
| clearly-V2 | R | 1383 | 5.4 | -26.5 | 5.8 | 3.90 |
| V1 | R | 177 | 14.0 | -31.7 | 3.6 | 3.35 |
| CdH | R | 230 | 4.8 | -2.4 | 8.3 | 3.34 |
| Pu | R | 233 | 13.7 | -6.4 | 0.2 | 3.32 |
| Acb | R | 27 | 3.0 | 7.1 | 0.6 | 3.41 |
| Cb5V | R | 41 | 2.0 | -27.0 | 3.8 | 2.80 |
| area-24b-prime | L | 40 | 0.0 | -7.4 | 12.4 | 3.40 |
| PMdc | L | 159 | -6.1 | -3.6 | 21.1 | 3.30 |
| PE | L | 45 | -10.7 | -18.2 | 21.7 | 3.90 |
| PEa | L | 73 | -5.7 | -19.8 | 16.2 | 3.76 |
| PIP | L | 115 | -8.2 | -25.8 | 8.4 | 3.94 |
| LIPd | L | 987 | -8.8 | -22.5 | 19.3 | 5.35 |
| LIPv | L | 255 | -7.0 | -22.1 | 16.6 | 4.13 |
| LOP | L | 48 | -5.9 | -27.8 | 18.9 | 3.56 |
| area-7a | L | 733 | -9.8 | -23.9 | 21.1 | 5.62 |
| area-7m | L | 140 | 0.2 | -26.1 | 12.4 | 3.09 |
| area-31 | L | 71 | 0.1 | -23.9 | 10.3 | 3.45 |
| area-23b | L | 118 | 0.0 | -15.4 | 14.1 | 3.19 |
| area-23c | L | 69 | -2.0 | -17.2 | 16.0 | 3.31 |
| area-29 | L | 35 | -3.3 | -19.8 | 7.6 | 3.46 |
| TEO | L | 62 | -26.6 | -17.0 | 0.4 | 3.45 |
| TPO | L | 71 | -20.0 | -12.1 | 2.0 | 3.21 |
| Ig | L | 41 | -17.7 | -9.6 | 1.4 | 4.49 |
| V4d | L | 72 | -6.3 | -28.1 | 20.0 | 3.78 |
| V3A | L | 89 | -10.0 | -26.4 | 8.6 | 3.82 |
| clearly-V2 | L | 106 | -2.8 | -28.8 | 9.5 | 3.18 |
| V1 | L | 130 | -10.8 | -24.4 | 6.9 | 3.32 |
| LP | L | 46 | -7.8 | -11.1 | 5.3 | 3.32 |
| MPul | L | 31 | -6.5 | -14.6 | 3.2 | 3.20 |
| **monkey C** | **LIPd-p** | | | | | |
| area-24c | R | 128 | 5.7 | 15.3 | 12.8 | 3.08 |
| area-24b-prime | R | 79 | 1.2 | -7.0 | 13.1 | 3.53 |
| area-24c-prime | R | 30 | 3.5 | -5.7 | 14.0 | 2.98 |
| area-8Ad | R | 131 | 11.5 | 11.1 | 15.1 | 5.26 |
| area-8Bs | R | 997 | 10.0 | 10.0 | 14.1 | 6.35 |
| area-9d | R | 31 | 9.9 | 14.0 | 15.7 | 3.82 |
| area-46d | R | 57 | 11.4 | 12.8 | 11.2 | 3.13 |
| area-46f | R | 401 | 8.6 | 17.4 | 9.1 | 3.29 |
| area-46v | R | 356 | 10.4 | 17.6 | 9.0 | 3.79 |
| area-45b | R | 58 | 13.9 | 9.5 | 8.4 | 3.73 |
| PMdc | R | 492 | 9.5 | 6.5 | 14.2 | 4.84 |
| V6 | R | 231 | 1.7 | -30.7 | 7.4 | 4.23 |
| V6Av | R | 389 | 2.3 | -31.1 | 10.7 | 5.15 |
| V6Ad | R | 54 | 4.3 | -31.9 | 15.5 | 3.85 |
| PEc-PEci | R | 69 | 5.4 | -22.6 | 22.6 | 3.18 |
| PEa | R | 691 | 10.0 | -17.1 | 15.8 | 3.98 |
| MIP | R | 1182 | 5.7 | -28.8 | 13.6 | 6.53 |

*Table S3, continued*

| | | | | | | |
|---|---|---|---|---|---|---|
| VIP | R | 896 | 7.1 | -21.4 | 11.9 | 4.62 |
| PIP | R | 275 | 8.8 | -27.1 | 10.0 | 4.01 |
| LIPd | R | 751 | 9.9 | -23.8 | 16.8 | 10.48 |
| LIPv | R | 1253 | 8.4 | -23.1 | 14.4 | 9.12 |
| LOP | R | 363 | 9.0 | -29.9 | 14.6 | 7.51 |
| MST | R | 1972 | 13.9 | -21.0 | 10.2 | 5.31 |
| area-7a | R | 1331 | 13.0 | -24.8 | 16.3 | 7.72 |
| area-7op | R | 26 | 13.4 | -9.3 | 8.3 | 2.99 |
| area-7m | R | 689 | 1.9 | -27.0 | 10.2 | 4.52 |
| area-31 | R | 199 | 2.3 | -24.2 | 9.4 | 4.54 |
| area-23a | R | 164 | 1.8 | -13.5 | 10.3 | 3.40 |
| area-23b | R | 373 | 1.3 | -13.2 | 12.7 | 3.46 |
| area-23c | R | 142 | 5.4 | -16.6 | 17.1 | 3.04 |
| area-v23 | R | 306 | 2.2 | -24.4 | 7.3 | 3.76 |
| area-29 | R | 64 | 3.5 | -13.9 | 9.8 | 3.35 |
| area-30 | R | 38 | 2.5 | -14.4 | 9.5 | 3.07 |
| TEO | R | 626 | 23.6 | -17.5 | 1.2 | 3.65 |
| TEpv | R | 125 | 21.6 | -20.1 | -6.0 | 3.25 |
| TEpd | R | 126 | 23.1 | -19.4 | -6.3 | 4.01 |
| TEm | R | 26 | 20.5 | -1.0 | -9.6 | 2.81 |
| TEa | R | 86 | 19.6 | -2.7 | -9.2 | 3.06 |
| IPa | R | 607 | 17.9 | -8.1 | -5.5 | 4.79 |
| PGa | R | 1575 | 16.9 | -13.4 | 1.2 | 6.75 |
| FST | R | 517 | 17.2 | -19.6 | 3.1 | 4.42 |
| TPO | R | 1222 | 20.1 | -11.8 | 0.7 | 4.59 |
| TAa | R | 98 | 23.8 | -13.7 | 4.2 | 3.18 |
| Tpt | R | 31 | 19.1 | -19.2 | 9.8 | 3.16 |
| Pi | R | 63 | 19.3 | -0.6 | -4.8 | 3.79 |
| Ig | R | 151 | 14.3 | -7.5 | 5.7 | 3.33 |
| MT | R | 439 | 14.9 | -24.4 | 12.0 | 4.96 |
| V4d | R | 2197 | 16.1 | -27.3 | 12.9 | 5.79 |
| V4v | R | 985 | 17.0 | -21.6 | -4.0 | 3.23 |
| V3A | R | 1433 | 13.3 | -29.0 | 10.6 | 4.80 |
| V3d | R | 1551 | 14.9 | -30.1 | 9.2 | 5.01 |
| V3v | R | 438 | 15.5 | -22.0 | -2.6 | 3.37 |
| possibly-V2 | R | 64 | 5.4 | -18.5 | 0.2 | 3.33 |
| clearly-V2 | R | 3597 | 10.1 | -32.6 | 11.5 | 4.95 |
| V1 | R | 2338 | 11.7 | -32.4 | 10.1 | 3.90 |
| BM | R | 165 | 9.2 | -1.2 | -8.1 | 3.47 |
| VCo | R | 38 | 7.7 | -0.9 | -8.7 | 3.21 |
| LaD | R | 88 | 12.3 | -0.9 | -8.4 | 3.22 |
| LaV | R | 40 | 13.0 | -0.4 | -9.9 | 3.04 |
| BLD | R | 97 | 11.1 | -0.7 | -7.3 | 3.32 |
| BLI | R | 82 | 10.9 | -0.4 | -8.8 | 3.91 |
| BLV | R | 46 | 10.1 | -0.1 | -9.9 | 3.88 |
| Me | R | 38 | 8.7 | -3.1 | -5.5 | 3.04 |
| CdH | R | 177 | 6.3 | -4.6 | 8.7 | 3.23 |
| Pu | R | 377 | 13.2 | -3.9 | 2.6 | 3.08 |
| MD | R | 118 | 1.1 | -8.8 | 2.8 | 3.35 |
| VLPV | R | 35 | 9.4 | -8.0 | 2.9 | 3.09 |
| MPul | R | 225 | 7.4 | -15.7 | 1.0 | 3.58 |
| LPul | R | 106 | 9.9 | -14.8 | 1.8 | 3.28 |
| IPul | R | 282 | 10.2 | -14.1 | -2.1 | 3.42 |
| DLG | R | 125 | 10.1 | -10.1 | -5.4 | 3.76 |
| MG | R | 60 | 8.5 | -12.3 | -2.6 | 2.91 |
| SC | R | 115 | 3.1 | -17.6 | -0.5 | 3.53 |
| Cb3V | R | 70 | 1.2 | -21.5 | -5.6 | 3.41 |
| Cb4V | R | 159 | 1.0 | -23.8 | -3.4 | 3.54 |
| Cb5V | R | 210 | 1.8 | -27.6 | -2.7 | 3.03 |
| Cb6V | R | 162 | 2.8 | -28.0 | -5.8 | 3.01 |
| Cb4I | R | 97 | 6.7 | -22.0 | -5.8 | 3.06 |
| Cb5I | R | 552 | 7.6 | -23.1 | -4.6 | 3.42 |
| Cb6I | R | 330 | 7.7 | -26.1 | -4.6 | 3.21 |
| Sim | R | 129 | 10.3 | -27.1 | -5.5 | 3.12 |
| area-24a-prime | L | 64 | -0.6 | -6.8 | 11.6 | 3.25 |
| area-24b-prime | L | 98 | -0.5 | -6.7 | 13.2 | 3.40 |
| area-24c-prime | L | 128 | -4.1 | -1.2 | 14.7 | 3.38 |
| V6 | L | 45 | -3.6 | -30.9 | 9.4 | 2.80 |
| V6Av | L | 206 | -1.3 | -30.8 | 10.4 | 3.71 |
| PE | L | 111 | -8.8 | -17.9 | 21.9 | 2.90 |
| PEa | L | 369 | -10.5 | -16.0 | 16.8 | 3.16 |
| MIP | L | 279 | -4.5 | -29.6 | 14.8 | 3.77 |
| VIP | L | 376 | -8.6 | -15.2 | 12.5 | 3.04 |
| AIP | L | 34 | -17.2 | -10.5 | 12.3 | 3.06 |
| LIPd | L | 893 | -8.1 | -23.5 | 19.2 | 4.82 |

*Table S3, continued*

| | | | | | | |
|---|---|---|---|---|---|---|
| LIPv | L | 149 | -7.6 | -21.0 | 15.8 | 3.70 |
| LOP | L | 129 | -6.4 | -28.2 | 18.3 | 4.75 |
| MST | L | 311 | -13.1 | -21.0 | 12.0 | 3.08 |
| area-7a | L | 805 | -10.9 | -23.8 | 19.4 | 3.61 |
| area-7b | L | 34 | -21.9 | -15.4 | 13.8 | 2.96 |
| area-7op | L | 741 | -18.6 | -14.0 | 11.5 | 3.20 |
| area-7m | L | 307 | -0.8 | -26.3 | 11.5 | 3.19 |
| area-31 | L | 76 | -1.7 | -23.9 | 8.9 | 3.25 |
| area-23a | L | 110 | -1.1 | -15.7 | 10.6 | 3.32 |
| area-23b | L | 448 | -1.2 | -15.3 | 12.7 | 3.65 |
| area-23c | L | 167 | -4.7 | -16.9 | 18.7 | 2.87 |
| area-v23 | L | 127 | -2.3 | -23.5 | 6.1 | 2.99 |
| area-29 | L | 126 | -2.5 | -15.8 | 10.1 | 3.79 |
| area-30 | L | 63 | -1.9 | -16.9 | 9.3 | 3.17 |
| TEO | L | 628 | -24.6 | -19.2 | -0.7 | 3.21 |
| TEpv | L | 101 | -15.3 | -15.0 | -7.8 | 3.13 |
| TEpd | L | 215 | -25.4 | -13.6 | -4.8 | 3.08 |
| TEad | L | 34 | -24.9 | -9.4 | -8.3 | 2.97 |
| TEm | L | 37 | -23.4 | -7.6 | -7.5 | 3.52 |
| TEa | L | 195 | -21.7 | -6.7 | -8.6 | 3.62 |
| PGa | L | 232 | -17.9 | -14.8 | 4.4 | 3.24 |
| FST | L | 34 | -20.0 | -19.5 | 0.7 | 3.51 |
| TPO | L | 509 | -19.8 | -12.2 | 3.6 | 3.01 |
| TAa | L | 155 | -22.7 | -17.1 | 9.3 | 3.21 |
| Tpt | L | 159 | -19.9 | -19.7 | 11.2 | 3.12 |
| Ia-Id | L | 234 | -17.6 | -1.5 | 1.2 | 3.04 |
| Ig | L | 147 | -18.1 | -7.1 | 2.9 | 3.42 |
| Ri | L | 104 | -12.3 | -12.6 | 8.4 | 3.77 |
| V4d | L | 887 | -17.4 | -24.6 | 11.8 | 3.44 |
| V4v | L | 1355 | -16.7 | -22.1 | -3.2 | 3.42 |
| V3A | L | 212 | -17.0 | -29.0 | 10.4 | 3.39 |
| V3d | L | 507 | -14.3 | -29.5 | 9.6 | 3.31 |
| V3v | L | 382 | -15.5 | -24.3 | -0.6 | 3.02 |
| possibly-V2 | L | 104 | -4.7 | -18.1 | 1.4 | 2.94 |
| clearly-V2 | L | 2027 | -11.3 | -31.7 | 9.7 | 3.32 |
| V1 | L | 1400 | -14.1 | -29.8 | 8.1 | 3.58 |
| LaD | L | 122 | -12.6 | -1.6 | -8.3 | 3.31 |
| LaV | L | 178 | -12.6 | -0.2 | -10.0 | 3.63 |
| BLI | L | 24 | -10.4 | -0.1 | -9.3 | 3.25 |
| CdH | L | 265 | -4.0 | 1.3 | 4.6 | 3.31 |
| Pu | L | 187 | -11.8 | 0.0 | 2.0 | 2.91 |
| Acb | L | 59 | -3.7 | 6.4 | 1.6 | 3.05 |
| MD | L | 41 | -1.6 | -9.7 | 2.6 | 2.42 |
| IMD | L | 112 | -0.7 | -10.0 | 2.8 | 3.72 |
| CM | L | 42 | -0.6 | -8.5 | 0.5 | 3.26 |
| VA | L | 51 | -2.3 | -6.0 | 0.8 | 3.23 |
| VLA | L | 25 | -6.7 | -4.6 | 5.2 | 2.99 |
| VM | L | 41 | -2.7 | -7.3 | -0.1 | 3.16 |
| LP | L | 41 | -6.3 | -11.6 | 5.1 | 3.09 |
| APul | L | 73 | -6.2 | -12.2 | 3.5 | 3.07 |
| MPul | L | 47 | -9.0 | -14.7 | 2.2 | 3.28 |
| LPul | L | 134 | -10.8 | -14.8 | 2.0 | 3.43 |
| IPul | L | 28 | -12.8 | -13.9 | -0.2 | 2.78 |
| SC | L | 64 | -3.9 | -17.7 | -0.3 | 3.31 |
| Lat | L | 37 | -6.5 | -25.9 | -7.7 | 3.35 |
| IntA | L | 94 | -5.1 | -25.6 | -7.3 | 3.73 |
| IntP | L | 51 | -4.1 | -25.6 | -7.6 | 3.53 |
| PFl | L | 33 | -14.3 | -20.5 | -8.3 | 2.98 |
| Cb3V | L | 99 | -0.9 | -20.8 | -5.3 | 3.29 |
| Cb4V | L | 194 | -1.3 | -24.4 | -4.6 | 3.69 |
| Cb5V | L | 376 | -2.1 | -27.5 | -2.3 | 3.19 |
| Cb6V | L | 235 | -3.0 | -27.9 | -6.0 | 3.31 |
| Cb4I | L | 27 | -6.0 | -23.3 | -5.2 | 3.70 |
| Cb5I | L | 165 | -8.5 | -21.8 | -6.2 | 3.08 |
| Cb6I | L | 453 | -8.4 | -26.1 | -5.0 | 3.54 |
| Sim | L | 331 | -10.2 | -27.0 | -5.8 | 2.95 |
| Crus1 | L | 26 | -15.5 | -25.0 | -6.8 | 3.21 |
| **monkey C** | **vPul** | | | | | |
| area-8Ad | R | 478 | 14.9 | 6.9 | 14.9 | 5.08 |
| area-8Av | R | 299 | 18.2 | 7.6 | 13.8 | 8.52 |
| area-8Bs | R | 306 | 13.2 | 5.9 | 11.5 | 4.77 |
| area-46d | R | 204 | 16.1 | 8.1 | 15.6 | 5.60 |
| area-46v | R | 422 | 17.5 | 11.1 | 13.0 | 5.06 |
| area-45a | R | 156 | 18.9 | 12.5 | 11.0 | 4.01 |

*Table S3, continued*

| | | | | | | |
|---|---|---|---|---|---|---|
| area-45b | R | 839 | 16.8 | 8.4 | 10.5 | 6.76 |
| area-44 | R | 102 | 15.0 | 7.1 | 8.4 | 3.98 |
| PMdc | R | 326 | 12.0 | 6.0 | 19.4 | 3.45 |
| F4 | R | 72 | 18.4 | 5.5 | 14.0 | 4.01 |
| F5 | R | 309 | 16.2 | 5.8 | 10.6 | 4.01 |
| PEa | R | 42 | 17.1 | -9.8 | 14.3 | 3.31 |
| MIP | R | 56 | 6.0 | -25.4 | 17.8 | 3.50 |
| TEO | R | 368 | 24.6 | -16.8 | 2.9 | 3.97 |
| IPa | R | 234 | 19.3 | -13.1 | -2.4 | 3.72 |
| PGa | R | 66 | 18.6 | -9.0 | -4.6 | 4.40 |
| FST | R | 173 | 18.5 | -20.6 | 3.2 | 3.47 |
| TPO | R | 260 | 20.7 | -13.0 | 0.1 | 4.54 |
| TAa | R | 183 | 25.3 | -15.2 | 5.9 | 4.09 |
| MT | R | 70 | 20.4 | -22.5 | 7.1 | 3.46 |
| V4d | R | 772 | 24.1 | -21.2 | 7.7 | 4.22 |
| V4v | R | 486 | 22.3 | -24.4 | -1.8 | 4.70 |
| V3A | R | 24 | 21.0 | -26.7 | 7.1 | 5.30 |
| V3d | R | 515 | 19.4 | -28.5 | 6.2 | 4.33 |
| V3v | R | 1012 | 19.6 | -27.6 | -1.8 | 5.46 |
| clearly-V2 | R | 2925 | 20.0 | -30.0 | 2.5 | 4.56 |
| V1 | R | 335 | 22.7 | -30.3 | 4.9 | 3.78 |
| APul | R | 43 | 8.1 | -12.2 | 0.9 | 5.09 |
| MPul | R | 344 | 6.8 | -15.2 | 1.8 | 4.19 |
| LPul | R | 131 | 10.4 | -14.8 | 0.8 | 4.43 |
| IPul | R | 349 | 10.5 | -13.2 | -1.6 | 7.15 |
| DLG | R | 423 | 11.0 | -10.2 | -3.5 | 7.25 |
| MG | R | 280 | 8.0 | -11.9 | -2.3 | 7.58 |
| SC | R | 336 | 3.0 | -15.2 | 0.0 | 4.52 |
| Crus1 | R | 83 | 14.8 | -30.8 | -6.6 | 3.28 |
| area-8Ad | L | 159 | -13.9 | 4.7 | 13.6 | 3.75 |
| area-8Av | L | 168 | -17.0 | 7.3 | 13.2 | 3.47 |
| area-8Bs | L | 52 | -12.2 | 4.3 | 12.5 | 3.72 |
| area-46d | L | 96 | -15.1 | 5.8 | 15.5 | 3.87 |
| area-46v | L | 73 | -17.7 | 9.7 | 11.8 | 3.60 |
| area-45b | L | 72 | -13.1 | 5.7 | 9.9 | 3.76 |
| F4 | L | 27 | -14.4 | 2.6 | 14.5 | 3.44 |
| F5 | L | 26 | -13.3 | 4.1 | 10.3 | 2.83 |
| LIPd | L | 27 | -7.0 | -26.4 | 18.9 | 3.81 |
| area-7a | L | 65 | -9.2 | -26.2 | 18.3 | 3.83 |
| TEO | L | 149 | -25.1 | -15.9 | 4.8 | 3.46 |
| TPO | L | 111 | -20.2 | -13.5 | 1.4 | 3.50 |
| V4d | L | 327 | -18.9 | -23.0 | 13.3 | 3.51 |
| V3A | L | 33 | -11.7 | -27.9 | 17.5 | 3.30 |
| **monkey B** | **dPul(a)** | | | | | |
| area-8Bd | R | 26 | 9.5 | 13.8 | 17.6 | 3.49 |
| area-8Bs | R | 126 | 10.1 | 11.5 | 15.7 | 3.59 |
| area-9d | R | 54 | 10.0 | 13.5 | 16.9 | 4.00 |
| area-46v | R | 105 | 20.5 | 11.5 | 10.3 | 3.69 |
| area-45a | R | 451 | 20.7 | 12.7 | 8.5 | 3.74 |
| PEa | R | 161 | 7.7 | -22.5 | 14.8 | 4.28 |
| MIP | R | 932 | 6.1 | -29.4 | 12.6 | 4.01 |
| VIP | R | 302 | 7.7 | -22.8 | 12.2 | 3.65 |
| PIP | R | 188 | 9.1 | -30.1 | 8.5 | 4.47 |
| LIPd | R | 43 | 19.3 | -18.3 | 17.8 | 3.49 |
| LIPv | R | 430 | 9.4 | -24.7 | 13.1 | 3.95 |
| LOP | R | 150 | 9.0 | -32.0 | 12.3 | 4.93 |
| MST | R | 692 | 17.3 | -23.8 | 8.7 | 4.38 |
| area-7a | R | 72 | 20.5 | -20.3 | 17.6 | 3.69 |
| area-7op | R | 125 | 20.6 | -20.3 | 15.9 | 3.81 |
| area-7m | R | 133 | 2.3 | -28.8 | 9.1 | 4.14 |
| area-23a | R | 38 | 1.8 | -18.7 | 8.0 | 3.42 |
| area-23b | R | 35 | 1.5 | -19.9 | 9.2 | 3.41 |
| area-v23 | R | 373 | 3.3 | -23.9 | 2.8 | 4.10 |
| area-30 | R | 28 | 2.2 | -18.3 | 7.8 | 3.50 |
| TEO | R | 309 | 25.5 | -19.7 | 3.1 | 4.46 |
| TEm | R | 94 | 28.0 | -11.8 | -5.5 | 3.82 |
| IPa | R | 153 | 23.5 | -18.4 | -0.1 | 4.49 |
| PGa | R | 345 | 17.7 | -21.1 | 5.2 | 4.78 |
| FST | R | 466 | 19.0 | -22.8 | 4.8 | 3.94 |
| TPO | R | 323 | 23.7 | -18.2 | 3.8 | 4.45 |
| Tpt | R | 251 | 23.9 | -22.0 | 12.1 | 4.73 |
| Ia-Id | R | 249 | 21.4 | -5.6 | -0.1 | 3.83 |
| Ig | R | 43 | 20.5 | -5.8 | 1.7 | 4.29 |
| MT | R | 640 | 19.9 | -25.6 | 9.3 | 4.20 |

*Table S3, continued*

| | | | | | | |
|---|---|---|---|---|---|---|
| V4d | R | 455 | 25.6 | -23.4 | 10.7 | 4.84 |
| V4v | R | 385 | 18.0 | -25.7 | -4.9 | 3.81 |
| V3A | R | 43 | 9.3 | -32.4 | 8.7 | 3.40 |
| V3d | R | 583 | 12.1 | -34.9 | 10.6 | 4.02 |
| V3v | R | 506 | 15.1 | -27.3 | -4.9 | 4.16 |
| possibly-V2 | R | 262 | 6.6 | -21.3 | 0.4 | 3.98 |
| clearly-V2 | R | 779 | 9.4 | -35.7 | 1.9 | 3.73 |
| MD | R | 207 | 3.2 | -12.0 | 4.0 | 5.11 |
| VLPD | R | 29 | 7.5 | -10.9 | 6.9 | 3.85 |
| LP | R | 148 | 7.6 | -13.6 | 5.7 | 7.25 |
| LD | R | 91 | 5.4 | -11.9 | 6.6 | 5.39 |
| APul | R | 416 | 7.5 | -14.4 | 2.8 | 10.88 |
| MPul | R | 362 | 7.8 | -17.6 | 1.8 | 10.91 |
| LPul | R | 108 | 11.1 | -16.9 | 2.4 | 7.06 |
| IPul | R | 68 | 11.0 | -15.7 | 0.0 | 5.91 |
| MG | R | 96 | 8.6 | -15.4 | -0.6 | 6.42 |
| SC | R | 431 | 3.5 | -18.2 | -0.1 | 5.64 |
| Cb4V | R | 93 | 2.4 | -25.2 | 0.8 | 3.33 |
| area-23b | L | 37 | 0.0 | -20.0 | 11.7 | 3.61 |
| TEO | L | 330 | -28.7 | -23.3 | 1.7 | 4.25 |
| V4d | L | 359 | -27.1 | -23.6 | 3.5 | 4.74 |
| V4v | L | 176 | -26.4 | -27.2 | 0.2 | 4.64 |
| V3A | L | 52 | -21.5 | -30.4 | 9.5 | 4.24 |
| V3d | L | 27 | -23.6 | -29.8 | 5.6 | 4.15 |
| V3v | L | 234 | -24.7 | -28.2 | 1.0 | 5.32 |
| clearly-V2 | L | 1151 | -24.7 | -30.2 | 3.1 | 4.82 |
| V1 | L | 580 | -25.9 | -36.4 | 1.2 | 4.17 |
| **monkey B** | **dPul** | | | | | |
| area-8Ad | R | 399 | 13.7 | 11.0 | 15.6 | 4.10 |
| area-8Bs | R | 553 | 11.5 | 8.7 | 14.8 | 3.77 |
| area-9d | R | 114 | 12.8 | 13.4 | 16.4 | 3.81 |
| area-46d | R | 466 | 14.9 | 13.4 | 13.1 | 3.95 |
| area-46f | R | 253 | 12.2 | 11.7 | 8.3 | 4.60 |
| area-46v | R | 1355 | 16.7 | 13.2 | 10.4 | 4.57 |
| area-12l | R | 207 | 22.3 | 13.8 | 4.0 | 4.24 |
| area-12r | R | 428 | 16.1 | 20.2 | 7.5 | 3.96 |
| area-45a | R | 1301 | 21.0 | 13.0 | 7.4 | 5.96 |
| area-45b | R | 298 | 21.5 | 9.2 | 6.9 | 4.31 |
| area-44 | R | 86 | 21.6 | 10.5 | 3.6 | 3.56 |
| PMdc | R | 164 | 11.2 | 7.1 | 15.8 | 3.52 |
| PMdr | R | 53 | 3.3 | 14.5 | 19.5 | 3.34 |
| F4 | R | 126 | 27.4 | -1.2 | 6.0 | 3.32 |
| F5 | R | 141 | 23.2 | 8.7 | 1.4 | 4.14 |
| V6 | R | 40 | 4.0 | -35.5 | 8.0 | 3.32 |
| MIP | R | 781 | 5.7 | -30.1 | 13.5 | 3.75 |
| VIP | R | 206 | 6.0 | -25.2 | 11.7 | 3.63 |
| LIPd | R | 118 | 18.3 | -17.5 | 16.8 | 3.25 |
| LIPv | R | 136 | 6.9 | -25.8 | 14.1 | 4.14 |
| LOP | R | 80 | 8.9 | -32.6 | 12.5 | 4.17 |
| MST | R | 906 | 15.8 | -25.2 | 10.9 | 4.27 |
| area-7a | R | 976 | 16.3 | -26.1 | 16.4 | 4.39 |
| area-7op | R | 214 | 21.2 | -19.6 | 15.2 | 3.67 |
| area-v23 | R | 82 | 3.2 | -25.3 | 3.2 | 3.60 |
| TEO | R | 840 | 27.1 | -18.8 | 1.1 | 6.23 |
| TEpv | R | 25 | 20.3 | -17.9 | -5.9 | 3.16 |
| TEpd | R | 469 | 29.1 | -15.7 | -5.4 | 5.84 |
| TEad | R | 37 | 28.4 | -10.5 | -8.5 | 3.59 |
| TEm | R | 1988 | 25.4 | -9.3 | -7.5 | 7.70 |
| TEa | R | 1559 | 22.2 | -7.0 | -10.2 | 7.36 |
| IPa | R | 2591 | 20.6 | -9.5 | -7.1 | 6.77 |
| PGa | R | 1445 | 20.0 | -11.0 | -3.3 | 5.90 |
| FST | R | 986 | 19.8 | -22.7 | 4.0 | 4.51 |
| TPO | R | 2655 | 23.2 | -8.3 | -5.1 | 5.89 |
| TAa | R | 832 | 27.7 | -10.8 | -3.1 | 6.64 |
| STGr | R | 280 | 26.4 | -0.3 | -5.7 | 3.86 |
| Tpt | R | 580 | 22.7 | -23.0 | 13.4 | 4.39 |
| Pi | R | 326 | 20.6 | -2.8 | -4.0 | 5.29 |
| Ia-Id | R | 1467 | 21.0 | -2.8 | -1.1 | 5.42 |
| Ig | R | 53 | 20.3 | -5.5 | 1.8 | 3.74 |
| MT | R | 976 | 19.2 | -26.1 | 10.4 | 4.13 |
| V4d | R | 1905 | 22.0 | -26.7 | 13.5 | 4.44 |
| V3A | R | 129 | 12.9 | -31.7 | 11.2 | 3.35 |
| V3d | R | 858 | 11.8 | -35.2 | 10.6 | 3.91 |
| V3v | R | 52 | 17.0 | -31.4 | -5.6 | 3.67 |

*Table S3, continued*

| | | | | | | |
|---|---|---|---|---|---|---|
| possibly-V2 | R | 123 | 6.8 | -20.4 | 0.5 | 4.19 |
| clearly-V2 | R | 726 | 9.3 | -35.3 | 7.8 | 3.47 |
| V1 | R | 271 | 14.5 | -36.7 | 3.5 | 3.32 |
| BM | R | 147 | 12.4 | -1.6 | -8.8 | 3.56 |
| VCo | R | 33 | 11.0 | 0.0 | -9.4 | 3.22 |
| AHi | R | 53 | 10.0 | -5.0 | -8.7 | 3.48 |
| LaD | R | 119 | 14.9 | -2.9 | -8.6 | 4.06 |
| LaV | R | 94 | 14.8 | -2.2 | -10.7 | 3.89 |
| BLD | R | 55 | 14.1 | -2.7 | -6.8 | 3.93 |
| BLI | R | 100 | 13.7 | -2.6 | -8.3 | 4.19 |
| BLV | R | 32 | 13.2 | -2.9 | -9.9 | 2.99 |
| CdT | R | 127 | 14.8 | -14.4 | -1.8 | 3.86 |
| Pu | R | 63 | 17.4 | -2.1 | -1.7 | 2.97 |
| MD | R | 162 | 3.0 | -13.2 | 4.1 | 6.99 |
| LP | R | 102 | 7.7 | -14.4 | 5.5 | 6.75 |
| LD | R | 31 | 5.8 | -13.1 | 6.2 | 5.32 |
| APul | R | 381 | 7.5 | -14.6 | 2.7 | 11.15 |
| MPul | R | 316 | 7.7 | -17.7 | 1.6 | 14.05 |
| LPul | R | 130 | 11.4 | -17.0 | 2.3 | 4.78 |
| IPul | R | 85 | 10.7 | -16.1 | -0.3 | 6.53 |
| MG | R | 142 | 8.5 | -15.6 | -0.9 | 8.08 |
| SC | R | 379 | 3.5 | -17.6 | 0.1 | 7.28 |
| Cb5V | R | 51 | 2.4 | -30.3 | 3.1 | 3.15 |
| Crus1 | R | 116 | 17.1 | -35.5 | -8.1 | 3.41 |
| area-45a | L | 36 | -18.2 | 16.6 | 5.9 | 3.24 |
| TEO | L | 105 | -27.8 | -24.7 | 0.4 | 3.47 |
| V4d | L | 346 | -27.9 | -23.8 | 6.0 | 3.61 |
| V4v | L | 131 | -26.7 | -26.8 | -0.1 | 4.01 |
| V3A | L | 59 | -21.0 | -30.8 | 9.7 | 3.81 |
| V3d | L | 56 | -21.9 | -31.2 | 7.1 | 3.80 |
| V3v | L | 298 | -24.4 | -29.6 | 0.4 | 4.25 |
| clearly-V2 | L | 1841 | -24.0 | -31.5 | 5.0 | 4.19 |
| V1 | L | 1327 | -24.3 | -34.4 | 5.4 | 3.75 |
| CdH | L | 55 | -4.4 | 7.2 | 0.8 | 3.48 |
| Acb | L | 73 | -5.3 | 5.9 | -2.0 | 3.24 |
| **monkey B** | **LIPd-a** | | | | | |
| area-24a | R | 48 | 1.7 | 7.7 | 9.7 | 3.39 |
| area-24b-prime | R | 59 | -0.2 | -5.1 | 11.5 | 4.37 |
| area-8Ad | R | 780 | 14.8 | 8.9 | 15.8 | 5.34 |
| area-8Bs | R | 809 | 11.5 | 7.2 | 14.4 | 4.31 |
| area-46d | R | 976 | 15.2 | 11.9 | 12.8 | 4.71 |
| area-46f | R | 262 | 11.8 | 12.7 | 8.6 | 3.56 |
| area-46v | R | 575 | 13.8 | 13.4 | 9.3 | 4.72 |
| area-12r | R | 136 | 14.7 | 22.1 | 6.9 | 3.20 |
| PMdc | R | 852 | 12.5 | 4.9 | 15.7 | 3.85 |
| V6 | R | 97 | 3.4 | -34.0 | 6.1 | 3.41 |
| V6Av | R | 48 | 1.7 | -33.6 | 11.8 | 3.37 |
| V6Ad | R | 397 | 2.4 | -33.2 | 16.2 | 4.00 |
| PEc-PEci | R | 190 | 2.2 | -31.7 | 19.0 | 4.00 |
| PE | R | 48 | 15.3 | -18.8 | 21.8 | 3.22 |
| PEa | R | 725 | 8.1 | -24.0 | 17.2 | 7.49 |
| MIP | R | 1951 | 5.4 | -29.9 | 14.7 | 5.92 |
| VIP | R | 514 | 7.5 | -24.1 | 11.8 | 4.05 |
| LIPd | R | 1687 | 11.9 | -24.9 | 17.7 | 11.16 |
| LIPv | R | 1729 | 9.8 | -24.8 | 14.4 | 11.59 |
| LOP | R | 340 | 9.5 | -32.2 | 13.6 | 3.82 |
| MST | R | 775 | 13.7 | -25.6 | 13.1 | 5.99 |
| area-7a | R | 1388 | 13.4 | -26.8 | 17.8 | 6.52 |
| area-7m | R | 1161 | 1.2 | -29.2 | 12.0 | 4.40 |
| area-31 | R | 646 | 1.6 | -24.7 | 11.1 | 7.09 |
| area-23b | R | 181 | 1.9 | -23.4 | 9.5 | 7.35 |
| area-v23 | R | 298 | 2.2 | -25.5 | 7.1 | 6.16 |
| TEO | R | 487 | 28.6 | -22.0 | -1.6 | 3.73 |
| TEpv | R | 856 | 22.4 | -18.9 | -7.6 | 3.45 |
| TEpd | R | 747 | 28.3 | -18.9 | -5.6 | 3.62 |
| TEm | R | 548 | 27.1 | -11.8 | -5.7 | 4.98 |
| TEa | R | 361 | 23.2 | -10.9 | -7.8 | 4.56 |
| IPa | R | 753 | 21.4 | -12.0 | -6.3 | 6.01 |
| PGa | R | 605 | 20.0 | -15.2 | -0.3 | 5.40 |
| FST | R | 237 | 19.5 | -21.4 | 3.2 | 3.57 |
| TPO | R | 812 | 24.1 | -11.1 | -3.8 | 4.30 |
| TAa | R | 228 | 27.7 | -9.6 | -4.7 | 4.53 |
| MT | R | 216 | 14.8 | -27.8 | 14.7 | 4.47 |
| V4d | R | 1714 | 16.9 | -30.0 | 15.8 | 4.36 |

*Table S3, continued*

| | | | | | | |
|---|---|---|---|---|---|---|
| V4v | R | 928 | 22.9 | -25.1 | -4.3 | 3.63 |
| V3A | R | 789 | 13.5 | -32.1 | 12.9 | 4.02 |
| V3d | R | 388 | 11.6 | -34.8 | 10.7 | 3.46 |
| V3v | R | 567 | 23.1 | -33.8 | -2.9 | 3.54 |
| clearly-V2 | R | 2160 | 10.6 | -36.9 | 1.7 | 3.63 |
| V1 | R | 5190 | 14.1 | -42.1 | 4.3 | 3.40 |
| BM | R | 76 | 12.4 | -3.0 | -6.9 | 3.13 |
| CdT | R | 40 | 17.5 | -11.7 | -5.3 | 3.62 |
| Pu | R | 45 | 7.2 | 5.7 | 0.5 | 3.27 |
| Acb | R | 135 | 4.2 | 4.4 | -0.5 | 3.27 |
| Cb5V | R | 217 | 1.6 | -35.7 | 1.3 | 3.82 |
| Cb6V | R | 253 | 2.7 | -40.3 | -3.8 | 3.48 |
| Cb7V | R | 37 | 1.2 | -43.5 | -6.8 | 3.86 |
| Cb6I | R | 44 | 11.1 | -30.4 | -4.5 | 3.26 |
| Sim | R | 130 | 12.4 | -33.8 | -5.4 | 3.31 |
| Crus1 | R | 57 | 13.6 | -34.2 | -6.3 | 3.04 |
| area-24a | L | 32 | -1.5 | 8.4 | 9.9 | 3.38 |
| area-24b | L | 24 | -1.8 | 8.4 | 10.6 | 3.42 |
| area-24a-prime | L | 54 | -0.6 | -4.5 | 11.0 | 3.34 |
| area-24b-prime | L | 134 | -0.4 | -5.1 | 11.5 | 3.93 |
| area-12r | L | 58 | -14.2 | 21.7 | 5.7 | 3.22 |
| PMdc | L | 355 | -14.6 | -1.2 | 19.4 | 3.26 |
| V6Av | L | 53 | -0.5 | -32.9 | 12.0 | 3.21 |
| V6Ad | L | 190 | -0.8 | -32.4 | 14.8 | 3.54 |
| PEc-PEci | L | 26 | -0.4 | -31.5 | 17.2 | 4.02 |
| PEa | L | 52 | -22.0 | -11.3 | 16.7 | 3.52 |
| LIPd | L | 451 | -12.7 | -23.4 | 19.7 | 8.11 |
| area-7a | L | 736 | -14.7 | -24.8 | 19.8 | 4.85 |
| area-7b | L | 43 | -22.0 | -15.6 | 17.9 | 3.35 |
| area-7m | L | 696 | -0.5 | -28.8 | 12.9 | 4.02 |
| area-31 | L | 411 | -0.8 | -24.5 | 11.4 | 5.37 |
| area-23b | L | 58 | -0.6 | -16.5 | 10.2 | 3.41 |
| area-v23 | L | 88 | -0.6 | -26.3 | 6.8 | 3.76 |
| TEO | L | 159 | -28.3 | -21.8 | 1.1 | 3.30 |
| TEm | L | 58 | -19.1 | 0.5 | -12.7 | 2.81 |
| TEa | L | 67 | -18.5 | 0.0 | -12.5 | 3.15 |
| IPa | L | 80 | -18.8 | -4.8 | -9.5 | 3.40 |
| PGa | L | 42 | -18.3 | 0.6 | -11.6 | 3.75 |
| TPO | L | 93 | -19.2 | 2.5 | -12.4 | 3.56 |
| MT | L | 65 | -18.2 | -26.3 | 12.8 | 3.20 |
| V4d | L | 641 | -21.8 | -26.8 | 10.8 | 3.71 |
| V3A | L | 115 | -22.2 | -27.8 | 9.0 | 3.62 |
| V3d | L | 271 | -19.9 | -30.4 | 8.9 | 3.67 |
| V3v | L | 61 | -22.8 | -27.3 | 2.2 | 3.23 |
| clearly-V2 | L | 1002 | -12.2 | -36.5 | 5.7 | 3.31 |
| V1 | L | 816 | -10.8 | -41.4 | 6.0 | 3.20 |
| AHi | L | 61 | -8.9 | -6.3 | -8.4 | 3.50 |
| Cb5V | L | 118 | -0.6 | -36.3 | 1.7 | 3.61 |
| Cb6V | L | 32 | -0.4 | -43.0 | -5.0 | 3.72 |
| Cb7V | L | 25 | -0.6 | -43.1 | -6.9 | 3.90 |
| **monkey B** | **LIPd-p** | | | | | |
| area-8Ad | R | 49 | 14.4 | 5.7 | 15.7 | 3.55 |
| area-8Bs | R | 217 | 12.3 | 5.1 | 12.9 | 3.95 |
| area-46d | R | 71 | 17.2 | 7.5 | 12.8 | 3.90 |
| area-46v | R | 93 | 16.9 | 7.6 | 11.6 | 3.47 |
| PMdc | R | 105 | 10.9 | 3.8 | 12.2 | 3.76 |
| F4 | R | 59 | 26.5 | 1.9 | 3.1 | 3.24 |
| F5 | R | 91 | 27.4 | 2.0 | 1.9 | 3.46 |
| V6 | R | 267 | 3.2 | -34.2 | 6.5 | 4.58 |
| PEc-PEci | R | 134 | 2.2 | -29.8 | 21.0 | 3.73 |
| PEa | R | 303 | 7.0 | -26.5 | 18.2 | 8.17 |
| MIP | R | 1878 | 5.6 | -30.1 | 14.7 | 8.07 |
| VIP | R | 292 | 6.9 | -24.9 | 11.9 | 4.81 |
| LIPd | R | 838 | 9.9 | -27.5 | 17.5 | 8.36 |
| LIPv | R | 1218 | 8.4 | -26.6 | 14.4 | 10.93 |
| LOP | R | 548 | 9.2 | -32.3 | 14.5 | 7.75 |
| MST | R | 700 | 12.9 | -26.4 | 13.0 | 7.53 |
| area-7a | R | 839 | 12.0 | -28.1 | 17.9 | 7.28 |
| area-7m | R | 323 | 1.9 | -28.8 | 7.8 | 6.94 |
| area-31 | R | 291 | 1.6 | -25.6 | 9.0 | 7.25 |
| area-23a | R | 152 | 1.1 | -20.8 | 7.6 | 4.41 |
| area-23b | R | 312 | 1.5 | -22.0 | 8.8 | 5.42 |
| area-v23 | R | 580 | 2.6 | -25.8 | 5.0 | 6.16 |
| TEO | R | 55 | 25.3 | -18.1 | 0.4 | 3.87 |

*Table S3, continued*

| | | | | | | |
|---|---|---|---|---|---|---|
| TEa | R | 60 | 22.8 | -12.2 | -7.0 | 3.58 |
| IPa | R | 421 | 21.9 | -13.7 | -4.4 | 3.89 |
| PGa | R | 630 | 19.3 | -17.9 | 2.4 | 5.01 |
| FST | R | 400 | 19.1 | -21.8 | 3.9 | 4.42 |
| TPO | R | 289 | 22.5 | -17.6 | 3.4 | 5.68 |
| Tpt | R | 33 | 24.2 | -22.3 | 10.9 | 4.43 |
| MT | R | 299 | 15.4 | -27.4 | 13.7 | 7.46 |
| V4d | R | 1099 | 13.6 | -30.6 | 16.9 | 7.29 |
| V4v | R | 258 | 15.3 | -24.4 | -6.2 | 4.89 |
| V3A | R | 559 | 11.8 | -32.3 | 12.8 | 4.70 |
| V3d | R | 202 | 9.3 | -34.6 | 12.9 | 4.34 |
| V3v | R | 640 | 14.2 | -26.0 | -5.3 | 4.63 |
| possibly-V2 | R | 28 | 7.1 | -22.6 | -0.9 | 3.88 |
| clearly-V2 | R | 2967 | 7.4 | -31.8 | 2.3 | 4.63 |
| V1 | R | 1231 | 12.9 | -36.5 | 2.9 | 4.04 |
| Pu | R | 67 | 15.3 | -1.2 | 4.4 | 3.43 |
| Cb4V | R | 80 | 2.4 | -24.9 | 0.9 | 2.82 |
| Cb5V | R | 60 | 2.2 | -30.1 | 3.3 | 3.18 |
| Cb6l | R | 76 | 12.4 | -28.1 | -5.9 | 3.46 |
| Sim | R | 35 | 11.5 | -31.4 | -5.7 | 3.47 |
| F4 | L | 112 | -26.6 | 0.7 | 4.9 | 3.42 |
| F5 | L | 39 | -26.7 | 3.0 | 4.3 | 3.42 |
| V6 | L | 45 | -0.8 | -31.7 | 7.8 | 3.27 |
| V6Av | L | 37 | -0.3 | -32.0 | 8.5 | 3.60 |
| LIPd | L | 308 | -9.8 | -26.2 | 19.8 | 4.53 |
| LIPv | L | 27 | -7.5 | -26.0 | 16.9 | 3.28 |
| MST | L | 104 | -14.8 | -24.7 | 13.7 | 3.77 |
| area-7a | L | 362 | -13.4 | -25.6 | 17.9 | 4.09 |
| area-7m | L | 84 | -0.6 | -28.6 | 8.1 | 3.58 |
| area-31 | L | 111 | -0.5 | -25.5 | 9.0 | 4.55 |
| area-23a | L | 87 | -0.4 | -20.3 | 7.5 | 3.64 |
| area-23b | L | 76 | -0.4 | -20.6 | 8.9 | 4.24 |
| area-v23 | L | 86 | -0.4 | -26.2 | 6.3 | 3.63 |
| MT | L | 201 | -15.1 | -26.4 | 14.6 | 3.52 |
| V4d | L | 159 | -13.5 | -27.3 | 17.2 | 3.65 |
| V3v | L | 44 | -13.3 | -28.3 | -3.5 | 3.28 |
| clearly-V2 | L | 342 | -7.8 | -30.3 | 3.0 | 3.89 |
| V1 | L | 244 | -15.8 | -31.3 | 3.0 | 3.50 |
| mcp | L | 54 | -8.5 | -22.7 | -9.3 | 3.35 |
| Cb5l | L | 34 | -8.2 | -22.3 | -8.3 | 3.55 |
| **monkey B** | **vPul** | | | | | |
| area-46v | R | 134 | 19.2 | 11.9 | 11.3 | 3.29 |
| area-45a | R | 116 | 20.0 | 13.7 | 8.6 | 3.44 |
| MST | R | 415 | 15.0 | -25.9 | 9.7 | 4.77 |
| TEO | R | 2081 | 27.4 | -21.5 | 1.0 | 5.94 |
| TEpv | R | 62 | 23.7 | -17.1 | -10.4 | 3.19 |
| TEpd | R | 299 | 28.2 | -19.1 | -5.0 | 4.42 |
| TEm | R | 837 | 25.6 | -11.6 | -5.9 | 5.05 |
| TEa | R | 651 | 22.2 | -7.9 | -9.7 | 4.61 |
| IPa | R | 1423 | 21.9 | -14.4 | -4.1 | 5.63 |
| PGa | R | 651 | 18.7 | -19.2 | 3.4 | 5.70 |
| FST | R | 1439 | 20.3 | -23.1 | 3.8 | 7.67 |
| TPO | R | 390 | 23.6 | -15.2 | -0.8 | 6.63 |
| TAa | R | 61 | 28.0 | -17.5 | 2.7 | 7.29 |
| MT | R | 665 | 18.9 | -26.0 | 9.0 | 5.20 |
| V4d | R | 1502 | 26.0 | -25.6 | 6.7 | 6.72 |
| V4v | R | 488 | 26.0 | -28.1 | -0.5 | 7.05 |
| V3A | R | 421 | 21.6 | -30.2 | 9.1 | 5.87 |
| V3d | R | 1304 | 17.8 | -32.5 | 8.7 | 5.85 |
| V3v | R | 981 | 24.2 | -30.3 | -0.3 | 7.19 |
| clearly-V2 | R | 5196 | 22.6 | -33.6 | 4.3 | 5.39 |
| V1 | R | 3446 | 20.6 | -37.9 | 3.7 | 5.30 |
| APul | R | 38 | 10.5 | -15.4 | 1.9 | 5.20 |
| MPul | R | 63 | 10.0 | -18.0 | 0.7 | 3.35 |
| LPul | R | 142 | 11.8 | -17.4 | 1.2 | 4.74 |
| IPul | R | 190 | 12.0 | -16.5 | -0.7 | 5.29 |
| DLG | R | 309 | 13.1 | -13.1 | -3.3 | 6.75 |
| MG | R | 147 | 9.0 | -15.5 | -1.6 | 3.91 |
| SC | R | 152 | 3.2 | -17.3 | 0.5 | 3.82 |
| area-8Ad | L | 140 | -15.2 | 8.5 | 16.2 | 3.59 |
| area-46d | L | 112 | -16.3 | 8.5 | 15.8 | 3.63 |
| MST | L | 57 | -21.0 | -20.6 | 5.9 | 4.78 |
| TEO | L | 407 | -27.3 | -21.3 | 3.3 | 3.80 |
| TEpd | L | 51 | -29.4 | -14.2 | -2.5 | 2.98 |

*Table S3, continued*

| | | | | | | |
|---|---|---|---|---|---|---|
| TEm | L | 392 | -26.9 | -14.8 | -2.7 | 4.37 |
| TEa | L | 52 | -24.1 | -14.9 | -4.6 | 3.76 |
| IPa | L | 448 | -22.0 | -12.8 | -4.8 | 3.79 |
| PGa | L | 107 | -19.9 | -19.5 | 5.2 | 4.19 |
| FST | L | 369 | -22.0 | -22.4 | 4.0 | 4.14 |
| TPO | L | 87 | -22.4 | -13.6 | -2.6 | 4.14 |
| TAa | L | 40 | -28.4 | -12.7 | -1.5 | 5.38 |
| V4d | L | 608 | -27.3 | -24.4 | 5.2 | 5.14 |
| V4v | L | 111 | -25.4 | -25.9 | 1.4 | 4.77 |
| V3v | L | 232 | -24.5 | -27.5 | 1.3 | 4.92 |
| clearly-V2 | L | 1763 | -27.5 | -28.3 | 4.0 | 5.03 |
| V1 | L | 555 | -27.3 | -32.4 | 2.9 | 3.92 |

**Table S4.** Stimulation effects in different task conditions, *atlas ROIs*. Significant p-values ($p<0.05$) for the two-way repeated measures ANOVAs and t-tests are in bold font. p – p-value, F – F-statistics, t – t-value. Unlike in the stimulation effect ROIs (**Table 2**), here the stimulation effect in the left hemisphere of monkey B was negative - i.e. activity decrease – in dPul, LIPd-a, and vPul (F-value of main effect stimulation marked with $^{(-)}$ superscript).

| | Main effect task | Main effect stim. | Task × stim. | Contra vs ipsi | Contra vs fix | Ipsi vs fix | Main effect task | Main effect stim. | Task × stim. | Contra vs ipsi | Contra vs fix | Ipsi vs fix |
|---|---|---|---|---|---|---|---|---|---|---|---|---|
| Format | F<br>p | F<br>p | F<br>p | t<br>p | t<br>p | t<br>p | F<br>p | F<br>p | F<br>p | t<br>p | t<br>p | t<br>p |
| | Left hemisphere | | | | | | Right hemisphere (stimulated) | | | | | |
| Monkey C | | | | | | | | | | | | |
| dPul(a) | 9.73 | 28.74 | 4.68 | 1.21 | 4.44 | 1.51 | 17.47 | 123.34 | 3.39 | 2.31 | 1.87 | -1.13 |
| | **0.000** | **0.000** | **0.010** | 0.227 | **0.000** | 0.133 | **0.000** | **0.000** | **0.035** | **0.023** | 0.064 | 0.262 |
| dPul | 2.51 | 1.04 | 4.95 | 2.10 | 2.81 | 1.29 | 8.10 | 29.35 | 0.77 | 0.73 | 1.18 | 0.54 |
| | 0.083 | 0.309 | **0.008** | **0.037** | **0.006** | 0.198 | **0.000** | **0.000** | 0.464 | 0.468 | 0.239 | 0.593 |
| LIPd-a | 16.60 | 27.18 | 5.49 | 2.55 | 2.92 | 0.00 | 22.39 | 48.12 | 1.28 | 1.40 | 0.10 | -1.48 |
| | **0.000** | **0.000** | **0.005** | **0.012** | **0.004** | 0.997 | **0.000** | **0.000** | 0.278 | 0.162 | 0.920 | 0.140 |
| LIPd-p | 102.89 | 28.60 | 22.58 | 4.55 | -1.63 | -7.43 | 30.66 | 67.78 | 26.13 | 3.70 | -3.19 | -8.11 |
| | **0.000** | **0.000** | **0.000** | **0.000** | 0.106 | **0.000** | **0.000** | **0.000** | **0.000** | **0.000** | **0.002** | **0.000** |
| vPul | 47.03 | 2.46 | 19.99 | 2.72 | -4.11 | -5.29 | 26.39 | 4.90 | 14.03 | 3.97 | -1.90 | -4.38 |
| | **0.000** | 0.119 | **0.000** | **0.007** | **0.000** | **0.000** | **0.000** | **0.029** | **0.000** | **0.000** | 0.059 | **0.000** |
| Monkey B | | | | | | | | | | | | |
| dPul(a) | 7.45 | 0.16 | 2.65 | -2.20 | -0.37 | 1.89 | 10.77 | 18.44 | 5.78 | -3.28 | -1.95 | 1.32 |
| | **0.001** | 0.686 | 0.072 | **0.030** | 0.712 | 0.061 | **0.000** | **0.000** | **0.003** | **0.001** | 0.053 | 0.188 |
| dPul | 48.21 | $31.25^{(-)}$ | 17.33 | -1.94 | 4.23 | 6.41 | 24.12 | 11.47 | 22.56 | -0.16 | 5.97 | 6.53 |
| | **0.000** | **0.000** | **0.000** | 0.055 | **0.000** | **0.000** | **0.000** | **0.001** | **0.000** | 0.870 | **0.000** | **0.000** |
| LIPd-a | 45.95 | $7.33^{(-)}$ | 0.42 | -0.54 | 0.40 | 0.96 | 42.13 | 22.75 | 3.04 | -2.49 | -0.85 | 1.54 |
| | **0.000** | **0.008** | 0.657 | 0.589 | 0.690 | 0.340 | **0.000** | **0.000** | **0.050** | 0.014 | 0.398 | 0.125 |
| LIPd-p | 21.43 | 2.55 | 13.22 | 1.92 | -3.30 | -5.33 | 82.31 | 7.07 | 16.17 | 0.99 | -4.52 | -6.26 |
| | **0.000** | 0.113 | **0.000** | 0.056 | **0.001** | **0.000** | **0.000** | **0.009** | **0.000** | 0.326 | **0.000** | **0.000** |
| vPul | 13.28 | $32.05^{(-)}$ | 1.97 | -1.84 | -0.44 | 1.51 | 15.81 | 0.46 | 0.08 | -0.13 | -0.37 | -0.29 |
| | **0.000** | **0.000** | 0.141 | 0.068 | 0.663 | 0.134 | **0.000** | 0.499 | 0.919 | 0.898 | 0.711 | 0.772 |

**Table S5.** Relationship between stimulation effect, spatial selectivity and task condition, *separately for each hemisphere*, stimulation effect ROIs. Significant p-values ($p<0.05$) for ANOVAs and t-tests are in bold font. R – Pearson's linear correlation coefficient for the three different task conditions, n – number of ROIs after exclusion of outliers (see Methods), p – p-value, F(df1,df2) – F-statistics (two-way mixed ANOVA), t – t-value for the t-test on the difference of stimulation effect strength between the three task conditions (contra, ipsi, fix). The left hemisphere ANOVA data are missing in monkey B in the dPul stimulation dataset because there were no contraversive (i.e. ipsilateral) tuned ROIs.

| | Corr. contra | Corr. ipsi | Corr. fix | Main effect tuning | Main effect task | Tuning × task | Corr. contra | Corr. ipsi | Corr. fix | Main effect tuning | Main effect task | Tuning × task |
|---|---|---|---|---|---|---|---|---|---|---|---|---|
| Format | R<br>(n)<br>p | R<br>(n)<br>p | R<br>(n)<br>p | F<br>(df1,df2)<br>p | F<br>(df1,df2)<br>p | F<br>(df1,df2)<br>p | R<br>(n)<br>p | R<br>(n)<br>p | R<br>(n)<br>p | F<br>(df1,df2)<br>p | F<br>(df1,df2)<br>p | F<br>(df1,df2)<br>p |
| | Left hemisphere | | | | | | Right hemisphere (stimulated) | | | | | |
| Monkey C | | | | | | | | | | | | |
| dPul<br>(a) | -0.77<br>(32)<br>**0.000** | 0.80<br>(32)<br>**0.000** | -0.47<br>(32)<br>**0.007** | 0.07<br>(1,30)<br>0.799 | 4.87<br>(2,60)<br>**0.011** | 13.78<br>(2,60)<br>**0.000** | -0.59<br>(64)<br>**0.000** | 0.74<br>(64)<br>**0.000** | -0.03<br>(64)<br>0.793 | 3.36<br>(1,62)<br>0.072 | 3.22<br>(2,124)<br>**0.044** | 40.93<br>(2,124)<br>**0.000** |
| dPul | -0.47<br>(33)<br>**0.006** | 0.11<br>(33)<br>0.546 | -0.03<br>(33)<br>0.868 | 2.51<br>(1,31)<br>0.124 | 2.82<br>(2,62)<br>0.068 | 6.55<br>(2,62)<br>**0.003** | -0.47<br>(63)<br>**0.000** | 0.21<br>(63)<br>0.100 | 0.24<br>(63)<br>0.053 | 0.08<br>(1,61)<br>0.781 | 7.50<br>(2,122)<br>**0.001** | 13.56<br>(2,122)<br>**0.000** |
| LIPd-a | -0.42<br>(21)<br>0.061 | 0.13<br>(21)<br>0.567 | -0.20<br>(21)<br>0.389 | 0.88<br>(1,19)<br>0.361 | 4.55<br>(2,38)<br>**0.017** | 2.31<br>(2,38)<br>0.113 | -0.15<br>(45)<br>0.334 | -0.07<br>(45)<br>0.634 | -0.16<br>(45)<br>0.287 | 0.64<br>(1,43)<br>0.428 | 6.24<br>(2,86)<br>**0.003** | 3.87<br>(2,86)<br>0.025 |
| LIPd-p | -0.56<br>(75)<br>**0.000** | 0.36<br>(75)<br>**0.001** | -0.12<br>(75)<br>0.288 | 0.51<br>(1,73)<br>0.478 | 0.07<br>(2,146)<br>0.929 | 7.13<br>(2,146)<br>**0.001** | -0.20<br>(80)<br>0.073 | 0.36<br>(80)<br>**0.001** | 0.15<br>(80)<br>0.177 | 9.16<br>(1,78)<br>**0.003** | 3.25<br>(2,156)<br>**0.041** | 7.39<br>(2,156)<br>**0.001** |
| vPul | -0.66<br>(14)<br>**0.011** | -0.23<br>(14)<br>0.425 | 0.44<br>(14)<br>0.116 | 2.79<br>(1,12)<br>0.121 | 1.27<br>(2,24)<br>0.298 | 4.38<br>(2,24)<br>**0.024** | -0.22<br>(34)<br>0.210 | 0.61<br>(34)<br>**0.000** | 0.09<br>(34)<br>0.611 | 0.22<br>(1,32)<br>0.644 | 1.28<br>(2,64)<br>0.286 | 3.38<br>(2,64)<br>**0.040** |
| Monkey B | | | | | | | | | | | | |
| dPul<br>(a) | 0.17<br>(8)<br>0.684 | 0.26<br>(8)<br>0.529 | -0.01<br>(8)<br>0.986 | 4.12<br>(1,6)<br>0.089 | 2.82<br>(2,12)<br>0.099 | 2.58<br>(2,12)<br>0.117 | -0.30<br>(47)<br>**0.042** | 0.38<br>(47)<br>**0.009** | 0.04<br>(47)<br>0.767 | 0.61<br>(1,45)<br>0.440 | 4.05<br>(2,90)<br>**0.021** | 7.02<br>(2,90)<br>**0.001** |
| dPul | 0.14<br>(11)<br>0.691 | 0.61<br>(11)<br>**0.046** | -0.48<br>(11)<br>0.139 | -<br>-<br>- | -<br>-<br>- | -<br>-<br>- | -0.20<br>(68)<br>0.100 | 0.40<br>(68)<br>**0.001** | 0.09<br>(68)<br>0.471 | 5.13<br>(1,66)<br>**0.027** | 22.87<br>(2,132)<br>**0.000** | 11.26<br>(2,132)<br>**0.000** |
| LIPd-a | 0.15<br>(32)<br>0.410 | 0.59<br>(32)<br>**0.000** | -0.24<br>(32)<br>0.195 | 1.83<br>(1,30)<br>0.186 | 4.65<br>(2,60)<br>**0.013** | 1.44<br>(2,60)<br>0.245 | 0.67<br>(53)<br>**0.000** | 0.69<br>(53)<br>**0.000** | -0.47<br>(53)<br>**0.000** | 13.30<br>(1,51)<br>**0.001** | 26.62<br>(2,102)<br>**0.000** | 21.80<br>(2,102)<br>**0.000** |
| LIPd-p | -0.70<br>(20)<br>**0.001** | 0.45<br>(20)<br>**0.048** | 0.30<br>(20)<br>0.197 | 0.04<br>(1,18)<br>0.846 | 7.58<br>(2,36)<br>**0.002** | 4.76<br>(2,36)<br>**0.015** | -0.29<br>(39)<br>0.069 | 0.12<br>(39)<br>0.482 | -0.04<br>(39)<br>0.808 | 0.55<br>(1,37)<br>0.464 | 4.97<br>(2,74)<br>**0.009** | 2.35<br>(2,74)<br>0.102 |
| vPul | -0.69<br>(15)<br>**0.005** | 0.91<br>(15)<br>**0.000** | 0.31<br>(15)<br>0.254 | 2.12<br>(1,13)<br>0.169 | 0.36<br>(2,26)<br>0.700 | 4.23<br>(2,26)<br>**0.026** | -0.62<br>(27)<br>**0.001** | 0.05<br>(27)<br>0.810 | 0.02<br>(27)<br>0.910 | 0.01<br>(1,25)<br>0.911 | 3.20<br>(2,50)<br>**0.049** | 5.96<br>(2,50)<br>**0.005** |

**Table S6.** Relationship between stimulation effect, spatial selectivity and task condition, *atlas ROIs*. Significant p-values (p<0.05) for ANOVAs and t-tests are in bold font. For this analysis, ROIs from both hemispheres were combined. R – Pearson's linear correlation coefficient for the three different task conditions, n – number of ROIs, p – p-value, F(df1,df2) – F-statistics (two-way mixed ANOVA), t – t-value for the t-test on the difference of stimulation effect strength between the three task conditions (contra, ipsi, fix).

| | Corr. contra | Corr. ipsi | Corr. fix | Main effect tuning | Main effect task | Tuning × task | Contra vs ipsi CS<0 | Contra vs ipsi CS>0 | Contra vs fix CS<0 | Contra vs fix CS>0 | Ipsi vs fix CS<0 | Ipsi vs fix CS>0 |
|---|---|---|---|---|---|---|---|---|---|---|---|---|
| Format | R<br>(n)<br>p | R<br>(n)<br>p | R<br>(n)<br>p | F<br>(df1,df2)<br>p | F<br>(df1,df2)<br>p | F<br>(df1,df2)<br>p | t<br>p | t<br>p | t<br>p | t<br>p | t<br>p | t<br>p |
| Monkey C | | | | | | | | | | | | |
| dPul(a) | -0.38<br>(256)<br>**0.000** | 0.73<br>(256)<br>**0.000** | -0.19<br>(256)<br>**0.003** | 9.92<br>(1,254)<br>**0.002** | 7.16<br>(2,508)<br>**0.001** | 125.61<br>(2,508)<br>**0.000** | 12.31<br>**0.000** | -8.28<br>**0.000** | 5.12<br>**0.000** | 0.83<br>0.407 | -8.43<br>**0.000** | 7.30<br>**0.000** |
| dPul | -0.65<br>(275)<br>**0.000** | -0.08<br>(275)<br>0.213 | -0.19<br>(275)<br>**0.001** | 18.33<br>(1,273)<br>**0.000** | 2.68<br>(2,546)<br>0.069 | 30.10<br>(2,546)<br>**0.000** | 9.07<br>**0.000** | -4.38<br>**0.000** | 5.45<br>**0.000** | -1.66<br>0.099 | -0.90<br>0.371 | 2.09<br>**0.039** |
| LIPd-a | 0.09<br>(277)<br>0.133 | 0.06<br>(277)<br>0.302 | -0.06<br>(277)<br>0.320 | 0.07<br>(1,275)<br>0.793 | 1.79<br>(2,550)<br>0.168 | 4.19<br>(2,550)<br>**0.016** | 0.27<br>0.785 | 2.77<br>**0.006** | -1.46<br>0.147 | 2.89<br>**0.004** | -1.85<br>0.067 | 0.27<br>0.788 |
| LIPd-p | -0.59<br>(275)<br>**0.000** | -0.01<br>(275)<br>0.844 | -0.31<br>(275)<br>**0.000** | 30.37<br>(1,273)<br>**0.000** | 26.94<br>(2,546)<br>**0.000** | 25.85<br>(2,546)<br>**0.000** | 8.46<br>**0.000** | -4.03<br>**0.000** | -1.39<br>0.165 | -5.16<br>**0.000** | -10.76<br>**0.000** | -2.76<br>**0.008** |
| vPul | -0.13<br>(273)<br>**0.029** | 0.59<br>(273)<br>**0.000** | -0.16<br>(273)<br>**0.010** | 14.95<br>(1,271)<br>**0.000** | 19.52<br>(2,542)<br>**0.000** | 35.58<br>(2,542)<br>**0.000** | 10.55<br>**0.000** | -4.00<br>**0.000** | -2.72<br>**0.007** | -1.79<br>0.075 | -10.43<br>**0.000** | 0.47<br>0.636 |
| Monkey B | | | | | | | | | | | | |
| dPul(a) | -0.67<br>(281)<br>**0.000** | 0.15<br>(281)<br>**0.015** | -0.00<br>(281)<br>0.936 | 17.66<br>(1,279)<br>**0.000** | 2.53<br>(2,558)<br>0.081 | 60.21<br>(2,558)<br>**0.000** | 5.93<br>**0.000** | -9.93<br>**0.000** | 4.98<br>**0.000** | -6.48<br>**0.000** | 0.60<br>0.551 | 2.37<br>**0.019** |
| dPul | -0.41<br>(277)<br>**0.000** | 0.49<br>(277)<br>**0.000** | -0.01<br>(277)<br>0.898 | 8.40<br>(1,275)<br>**0.004** | 44.87<br>(2,550)<br>**0.000** | 37.95<br>(2,550)<br>**0.000** | 3.80<br>**0.000** | -6.62<br>**0.000** | 11.14<br>**0.000** | 0.07<br>0.942 | 5.69<br>**0.000** | 7.27<br>**0.000** |
| LIPd-a | -0.19<br>(279)<br>**0.001** | 0.61<br>(279)<br>**0.000** | -0.18<br>(279)<br>**0.003** | 10.30<br>(1,277)<br>**0.001** | 2.69<br>(2,554)<br>0.069 | 34.59<br>(2,554)<br>**0.000** | 4.61<br>**0.000** | -8.02<br>**0.000** | 0.09<br>0.929 | -0.59<br>0.555 | -3.69<br>**0.000** | 6.54<br>**0.000** |
| LIPd-p | -0.46<br>(285)<br>**0.000** | 0.56<br>(285)<br>**0.000** | 0.15<br>(285)<br>**0.012** | 7.57<br>(1,283)<br>**0.006** | 35.14<br>(2,566)<br>**0.000** | 65.30<br>(2,566)<br>**0.000** | 8.78<br>**0.000** | -5.88<br>**0.000** | 1.06<br>0.291 | -9.02<br>**0.000** | -8.65<br>**0.000** | -2.71<br>**0.007** |
| vPul | -0.43<br>(279)<br>**0.000** | 0.41<br>(279)<br>**0.000** | 0.02<br>(279)<br>0.755 | 2.02<br>(1,277)<br>0.156 | 0.42<br>(2,554)<br>0.656 | 24.67<br>(2,554)<br>**0.000** | 4.83<br>**0.000** | -6.03<br>**0.000** | 2.74<br>**0.007** | -2.86<br>**0.005** | -1.47<br>0.143 | 2.76<br>**0.006** |

**Table S7.** Stimulation effect model fitting. Adjusted R-squared values signifying the goodness of fit of Pearson's linear correlation between actual stimulation effects and fitted estimates derived from the three models: additive (add.), additive scaled (add. scaled) and multiplicative (mult.), and the significance of paired t-test comparisons between these values across hemispheres and datasets, separately for each monkey, for the two sets of ROIs.

| ROI dataset | Stimulation effect ROIs | | | Atlas ROIs | | |
|---|---|---|---|---|---|---|
| Model | additive | add.scaled | multiplicative | additive | add.scaled | multiplicative |
| | Monkey C | | | | | |
| Adjusted $R^2$ | 0.297 | 0.459 | -0.532 | 0.361 | 0.509 | 0.442 |
| Pair-wise t-test comparison between models | add. vs add.scaled **0.001** | add. vs mult. **0.000** | add.scaled vs mult. **0.000** | add. vs add.scaled **0.001** | add. vs mult. 0.092 | add.scaled vs mult. **0.000** |
| | Monkey B | | | | | |
| Adjusted $R^2$ | 0.369 | 0.479 | -0.899 | 0.445 | 0.532 | 0.492 |
| Pair-wise t-test comparison between models | add. vs add.scaled **0.001** | add. vs mult. **0.000** | add.scaled vs mult. **0.000** | add. vs add.scaled **0.007** | add. vs mult. 0.101 | add.scaled vs mult. **0.007** |

### 1 Supplementary Results

#### 1.1 Task performance and eye movements

##### 1.1.1 Overall hit rate and trial aborts during and after stimulation

In monkey C, dPul(a) stimulation neither affected hit rate nor the number of trials aborted in and after the stimulation period, respectively (all $\chi^2$s(3) ≤ 0.83, all *p*s ≥ 0.3624). Similar results were found for dPul stimulation (all $\chi^2$s(3) ≤ 1.31, all *p*s ≥ 0.2523). In monkey B, overall hit rate and the number of trials aborted in the stimulation period were also not affected by dPul(a) stimulation (both $\chi^2$s(3) ≤ 2.494, all *p*s ≥ 0.1143). However, dPul(a) stimulation led to an overall increase in the number of trials aborted *after* the stimulation period ($\chi^2(3) = 15.642$, $p < 0.001$), which was mainly driven by an impairment in making saccades to cued locations in the contraversive (left) hemifield ($\chi^2(3) = 29.03$, $p < 0.001$); dPul stimulation did not have an effect on the number of trials aborted after the stimulation period ($\chi^2(3) = 1.48$, $p = 0.2240$). Overall hit rate was not affected by dPul stimulation ($\chi^2(3) = 2.21$, $p = 0.1370$) but there was a significant effect on the number of trials aborted in the stimulation period ($\chi^2(3) = 3.67$, $p < 0.05$) with a significantly decreased number of aborted trials in the contraversive memory saccade task ($\chi^2(3) = 5.37$, $p < 0.05$).

The stimulation in vPul did not lead to any changes in the overall hit rate or the number of trials aborted during or after the stimulation period in monkey C (although there was a weak tendency of increased hit rates in all task conditions), but in monkey B, both the hit rate overall increased ($\chi^2$s(3)=8.91, p=0.003) and the trials were aborted less frequently during and after the stimulation ($\chi^2$s(3)=4.30, p<0.05 and $\chi^2$s(3)=8.14, p<0.01), significant for the contraversive memory saccade trials after the stimulation.

LIPd-a stimulation did not have a significant effect on hit rate or the number of aborted trials (monkey C: all $\chi^2$s(3) ≤ 1.11, all *p*s ≥ 0.2930; monkey B: all $\chi^2$s(3) ≤ 3.63, all *p*s ≥ 0.0568). LIPd-p stimulation led to a significantly lower overall hit rate in monkey C ($\chi^2(3) = 5.03$, $p < 0.05$), which was mainly driven by a lower number of successful trials in the ipsiversive memory saccade task with stimulation compared to the control condition ($\chi^2(3) = 4.48$, $p < 0.05$). However, there was no significant increase in the number of trials aborted during or after the stimulation period ($\chi^2(3) = 0.59$, $p = 0.4416$ and 2(3) = 3.18, $p = 0.0747$, respectively). In monkey B, LIPd-p stimulation did not affect overall hit rate $\chi^2(3) = 1.84$, $p = 0.1753$) or the number of trials aborted in the stimulation period ($\chi^2(3) = 1.37$, $p = 0.2424$) but led to a significant decrease in the overall number of trials aborted after the stimulation period ($\chi^2(3) = 4.07$, $p < 0.05$). None of the comparisons between stimulation and control trials for each task separately reached significance (all $\chi^2$s(3) ≤ 2.70, all *p*s ≥ 0.1005).

**Figure S11** and **S12** summarize these data as bar plots.

#### 1.1.2 Frequency of eye movements

In brief, the effect on the frequency of eye movements was not consistent across datasets and animals: the stimulation of dPul(a) in monkey C and LIPd-p in monkey B led to a significant increase in the number of small eye movements during the fixation in stimulation period in all three tasks, but other datasets showed no change or a small increase and/or decrease in saccade frequency.

For dPul(a) dataset in monkey C, the two-way ANOVA on the number of eye movements in the stimulation period revealed a significant main effect of stimulation ($F(1, 526) = 139.41$, $p < 0.001$) with an increased number of eye movements in stimulation trials compared to the control condition in all three tasks (all *p*s < 0.001). In contrast, there were no significant effects of dPul stimulation on the number of eye movements in the stimulation period (main effect stimulation: $F(1, 823) = 0.07$, $p = 0.7881$, task × stimulation interaction: $F(2, 823) = 0.46$, $p = 0.6324$). In monkey B dPul(a) stimulation also significantly affected the number of eye movements in the stimulation period (main effect stimulation: $F(1, 1122) = 18.49$, task × stimulation interaction: $F(2, 1122) = 9.50$, both *p*s < 0.001), with a significantly lower number of eye movements in the fixation and the contraversive memory saccade task ($t(392) = 2.80$ and $t(364) = 5.09$, respectively, both *p*s < 0.001). dPul stimulation also significantly influenced the number of eye movements in the stimulation period as shown by a significant task × stimulation interaction effect ($F(2, 1263) = 14.71$, $p < 0.001$): there was a significantly higher number of eye movements in the ipsiversive memory saccade task ($t(424) = 4.13$, $p < 0.001$) whereas the number of eye movements was decreased by dPul stimulation in the contraversive memory saccade task ($t(412) = 3.15$, $p < 0.001$).

The stimulation in vPul did not lead to any changes in the number of eye movements in the stimulation period in monkey C, but in monkey B there was a main effect of the stimulation on decreasing the number of eye movements ($F(1, 983)=29.33$, $p<0.000$), significant also separately in all tasks (*p*s < 0.01).

The ANOVA on the number of eye movements in the stimulation period revealed a significant main effect of stimulation in LIPd-a in monkey C ($F(1, 873) = 10.74$, $p < 0.01$). Further post-hoc t tests showed that LIPd-a stimulation led to a significantly higher number of eye movements only in the contraversive memory saccade task ($t(318) = 3.09$, $p < 0.01$). LIPd-p stimulation did not affect the number of eye movements in the stimulation period (main effect stimulation: $F(1, 882) = 0.06$, $p = 0.8063$, task × stimulation interaction: $F(2, 882) = 1.73$, $p = 0.1773$). In monkey B, LIPd-a stimulation also affected the number of eye movements in the stimulation period as shown by a significant main effect of stimulation ($F(1, 1503) = 22.89$, $p < 0.001$), but in contrast to monkey C, it led to less eye movements with the difference between stimulation and control trials reaching significance in the fixation ($t(495) = 3.24$, $p < 0.01$) and the ipsiversive memory saccade task ($t(504) = 3.24$, $p < 0.01$). Stimulation in LIPd-p led to a higher number of eye movements in the stimulation period as shown by a significant main effect of stimulation ($F(1, 1278) = 41.94$, $p < 0.001$)

and significant differences between stimulation and control trials for all three tasks (fixation: $t(423) = 2.25$, $p < 0.05$; memory saccade right: $t(432) = 4.69$, $p < 0.001$; memory saccade left: $t(423) = 4.26$, $p < 0.001$).

**Figure S13** summarizes these data as bar plots.

##### 1.1.3 Saccade latencies

In monkey C, the two-way ANOVA on saccade latencies did not reveal significant effects of dPul(a) stimulation ($F(1, 336) = 0.16$, $p = 0.69$), but a significant effects of dPul stimulation ($F(1, 514)$s $\geq$ 8.04, $p$s < 0.001) with significantly longer latencies for contraversive saccades following stimulation compared to the control condition ($t(244) = 4.17$, $p < 0.001$). In monkey B for dPul(a) stimulation the two-way ANOVA on saccade latencies revealed a significant main effect of stimulation ($F(1, 727) = 15.70$, $p < 0.001$). Further post-hoc t tests showed that saccades to cued locations in both the ipsiversive and the contraversive hemifield were significantly delayed compared to the control conditions ($t(366) = 2.95$ and $t(361) = 2.68$, respectively, both $p$s < 0.01). Similar to dPul(a) stimulation, the two-way ANOVA revealed a significant main effect of dPul stimulation with longer latencies of saccades to both the contraversive and the ipsiversive hemifield ($F(1, 833) = 4.95$, $p < 0.05$). However, for neither of the saccade tasks the difference in saccade latencies between stimulation and control trials reached significance in post-hoc t tests ($t(423) = 1.60$ and $t(410) = 1.54$, both $p$s $\geq$ 0.1048).

The stimulation in vPul did not lead to any changes in the saccade latency in monkey C, but in monkey B there was a main effect of the stimulation on increasing the latency ($F(1, 651) = 8.12$, $p < 0.01$), significant also separately for the contraversive task ($p < 0.01$).

Saccade latencies were not affected by LIPd-a stimulation in monkey C (main effect stimulation: $F(1, 581) = 1.05$, $p = 0.3068$, task × stimulation interaction: $F(1, 581) = 0.10$, $p = 0.7557$), but the two-way ANOVA on saccade latencies revealed a significant main effect of LIPd-p stimulation ($F(1, 584) = 10.42$, $p < 0.01$) mainly driven by an increased latency due to stimulation in the contraversive memory saccade task ($t(290) = 3.03$, $p < 0.01$). Saccade latencies were also affected by LIPd-a stimulation in monkey B as shown by a significant main effect of stimulation ($F(1, 1004) = 11.11$, $p < 0.001$) and a significant task × stimulation interaction effect ($F(1, 1004) = 4.59$, $p < 0.05$). Post-hoc t tests showed that saccade latencies were significantly longer due to stimulation only in the contraversive memory saccade task ($t(502) = 3.66$, $p < 0.001$). LIPd-p stimulation did not have significant effects on saccade latencies (main effect stimulation: $F(1, 854) = 0.01$, $p = 0.9190$, task × stimulation interaction: $F(1, 854) = 0.36$, $p = 0.5493$).

**Figure S14** summarizes these data as bar plots.
